# Allosteric remodelling of the Ebola virus glycoprotein underlies differences in entry mechanisms across species

**DOI:** 10.64898/2026.09.04.749277

**Authors:** Valeria Calvaresi, Jiří Kratochvíl, Anna Bekauri, Philipp Kukura, Weston B. Struwe

**Affiliations:** The Kavli Institute for Nanoscience Discovery, University of Oxford, Oxford OX1 3QU, UK; Department of Biochemistry, University of Oxford, Oxford OX1 3QU, UK; Department of Chemistry, University of Oxford, Oxford OX1 3TA, UK

## Abstract

During Ebola virus transit through the endo-lysosomal pathway, the glycoprotein GP undergoes a series of conformational rearrangements upon cathepsin cleavage and NPC1 receptor binding, culminating in host-virus membrane fusion. While these rearrangements underpin viral entry, their molecular mechanisms remain poorly understood. Here, we combine hydrogen/deuterium-exchange mass spectrometry and mass photometry to resolve structural dynamic and energetic transitions of GP of two major species, Zaire (EBOV) and Sudan (SUDV). We describe the allosteric axes that govern reorganization of the cleavage sites, opening of the receptor binding cavity and priming for fusion upon NPC1 engagement. We show that GP cathepsin cleavage and receptor binding exhibit different mechanisms and kinetics across species, and are allosterically coupled in a species-specific manner. We reveal that fusion priming is optimized via distinct routes in EBOV and SUDV GP. These data suggest that GP structural dynamics and allosteric remodelling underlie differences in entry mechanisms across Ebolavirus species.

## Introduction

Ebola viruses emerged in Africa five decades ago^1,2^ and have since caused recurrent epidemics. Among the species that infect humans, Zaire (EBOV) and Sudan (SUDV) are responsible for most outbreaks to date^3^. The EBOV Makona variant emerged from the original Mayinga strain^4^, giving rise to the 2013-2016 outbreak, the deadliest ever recorded^3^. The fusion glycoprotein GP is the only protein on the virus surface^5–7^ and mediates entry into host cells^8,9^. GP is a glycosylated homo-trimer where each protomer is composed of two subunits cleaved apart by furin^10^, but remaining linked via a disulphide bond^11^: GP1, the receptor binding subunit, and GP2, the fusion subunit. GP1 has two highly glycosylated domains that occlude the receptor binding cavity, termed the glycan cap (GC) and the mucin-like domain (MLD), the latter of which is structurally disordered^12,13^. Upon attachment to a host cell, the virus transitions by macropinocytosis into the endo-lysosomal compartment^14,15^, where GP is cleaved by two cysteine-proteases (cathepsins B and L)^16–18^ that remove the GC and MLD^19–21^, exposing the receptor binding region (RBR)^20,22^ and allowing for its engagement to the lysosomal receptor NPC1^21,23–26^. Whether receptor binding is sufficient to trigger membrane fusion, or other factors are necessary, is still debated^27–30^.

Cryo-EM and crystal structures have provided high-resolution structural information of individual GP domains and their organization^12,13,22,31,32^, which has been crucial to understand the trimer topology and identify its NPC1 binding site^24–26^. Nevertheless, as for other fusion glycoproteins^33–37^, what mainly defines the capability of GP to exert its functions are large-scale conformational dynamic rearrangements and more localized site-specific motions that progressively enable the trimer to modify its conformation^38^, thus driving virus entry. To date, our understanding of these structural dynamics is very limited. For instance, the GP conformational changes associated with the virus transition into the endo-lysosomes are largely unknown, although these define how pH primes GP for proteolytic cleavage. A comprehensive characterization of cleavage sites and kinetics, proteolytic intermediates and the cleavage-associated conformational changes that prime GP for receptor binding is lacking. Structures of the cleaved trimer bound to NPC1 are static snapshots of receptor binding and cannot fully uncover the intricated mechanism of the successive binding events within the trimer, or how receptor binding primes GP for the fusogenic transition. Furthermore, it is established that GP is responsible for critical pathogenic differences among viral species and strains^39^. Understanding the structural dynamic differences across the GP trimers of SUDV and EBOV, and their functional significance in the entry process, would uncover species- and strain-specific molecular divergences that contribute to their pathogenic differences and could underlie virus evolution.

To address these unknowns, we employed hydrogen/deuterium-exchange mass spectrometry (HDX-MS)^40^ and mass photometry (MP)^41,42^ on the GP trimers of Mayinga EBOV, Makona EBOV and SUDV under conditions recapitulating the various stages of entry. We resolve the allosteric mechanisms governing the pH-driven conformational shift, proteolytic cleavage, protomer opening, NPC1 receptor binding and fusion priming, and how these processes allosterically feedback into one another to promote viral entry. Our comparative kinetic and structural dynamic data on cathepsin cleavage and receptor engagement unveil intra- and inter-species divergences in entry profiles that we explain by species- and variant-specific structural dynamic adaptations of the fusion glycoprotein GP, delineating the possible evolutionary routes undertaken by the Ebola virus that enabled these major infectious species to establish.

## Results

### An RBR-fusion peptide allosteric axis is potentiated in Makona and Sudan GP

We first investigated the differences in structural dynamics between Mayinga, Makona and Sudan GP by comparing the peptide-level HDX of their uncleaved trimers. In this study, we utilized pre-fusion stabilized GP ectodomains carrying GC and MLD, and containing T577P and K588F stabilizing substitutions^43^ (**Figs. S1-S3**).

HDX data uncovered an allosteric axis linking the opening of the RBR to the dynamics of the fusion peptide, which is potentiated in Makona and Sudan GP compared to Mayinga GP (**Fig. 1c**). Specifically, for Makona GP, we observed increased HDX in the RBR α1 helix (residues 80-85, depicted as area 3 in Fig. 1) and in the IFL β-hairpin that clamps the fusion peptide (residues 518-524 and 543-547, areas 15 and 16). Those two elements are bridged by the β3-α1 loop, which lies beneath the RBR and contacts the IFL β-hairpin, and also displayed increased HDX (residues 71-79, area 3) (**Fig. 1a,c,e**). For Sudan GP, we detected the same structural dynamics changes, but HDX effects were more pronounced compared to Makona GP and resulted in a detectable mobilization of the fusion peptide itself (**Fig. 1b,c,f**). Makona GP carries A82V in its α1 helix and T544I in its fusion peptide. Sudan GP, however, contains a higher number of residue substitutions in its α1 helix (N73S, A76S and V79I) and several more in its IFL domain, including T544, which altogether determine the higher potentiation of this axis in Sudan compared to Makona GP (**Fig. S1**).

**Fig. 1.**
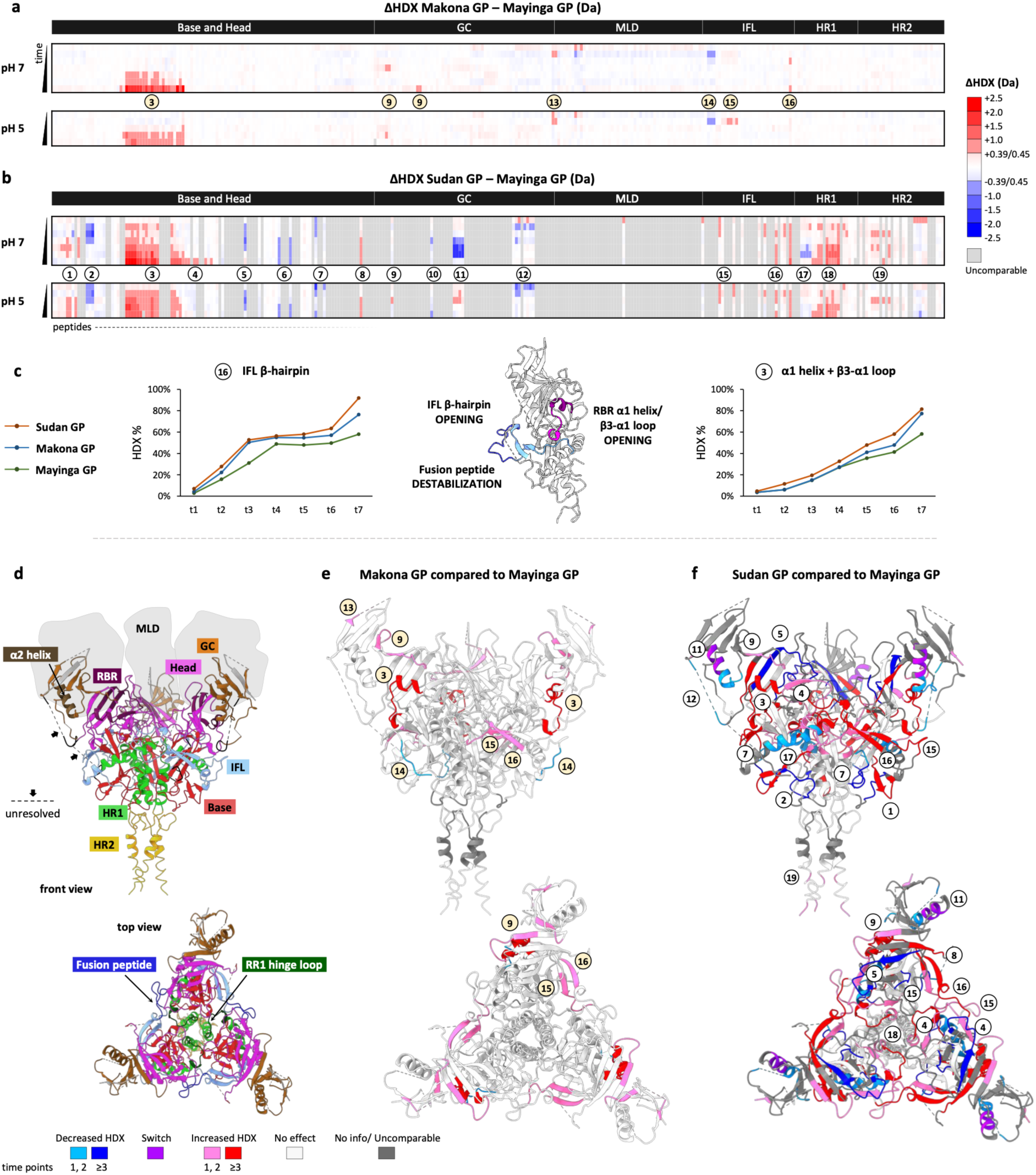
Differences in HDX (ΔHDX) at pH 7 and pH 5 among GP trimers. **a,b,** Plots illustrating ΔHDX profiles of Makona minus Mayinga GP (**a**) and Sudan minus Mayinga GP (**b**) at pH 7 and 5. X-axis: peptides whose HDX was followed ordered sequentially along the protein sequence. Y-axis: time points studied. pH 7: t1= 4 s (ice), t2= 4 s (23°C), t3=0.6 min (23°C), t4=6 min (23°C), t5=12 min (28°C), t6=1 h (23°C), t7=10 h (23°C). pH 5: t1= 400 s (ice), t2=400 s (23°C), t3=1 h (23°C), t4=10 h (23°C), t5=20 h (28°C). Positive and negative ΔHDX values (Da) surpassing the threshold of significance are colour-scaled in red and blue, respectively, based on the magnitude of the effect. Source data are provided within source data file. **c**, Allosteric axis RBB–fusion peptide illustrated on one protomer of GP structure (Mayinga GP, PDB 5JQ3): the opening of the α1 helix and of the IFL β-hairpin induces destabilization of the fusion peptide. Deuterium uptake plots (pH 7) of peptides 542-547 and 69-84 illustrate the axis potentiation for Makona and Sudan compared to Mayinga GP. **d**, Individual domains of GP (in PDB 5JQ3). GP1: Base and head, glycan cap (GC), and mucin-like domain (MLD). GP2: internal fusion loop (IFL), heptad repeat 1 (HR1), and heptad repeat 2 (HR2). The α1 helix lies in the RBR (head); the fusion peptide within the IFL; the RR1 hinge loop in the HR1, at the tip of the stalk. **e,f**, ΔHDX profiles of Makona minus Mayinga GP (**e**) and Sudan minus Mayinga GP (**f**) superimposed on GP structure.

Sudan and EBOV GP differ by 46% in their primary sequence (**Fig. S1**), and accordingly, we identified significant structural dynamics differences in several regions of the trimer, including the β-strands surrounding the α1 helix (areas 4, 5 and 8) and the HR1A (area 7) **(Fig. 1 b,f).** It is noteworthy that, in Sudan GP, the hinge loop of the refolding region 1 (RR1) in GP2 and the GP1 N-terminus (residues 572-585 and 33-43, areas 19 and 1, respectively) displayed increased HDX compared to Mayinga and Makona GP. These two areas include residues forming a small hydrophobic pocket directly responsible for the trimer stability^43^ and harbour F582Y and S583T substitutions in SUDV, which our data highlight reducing the degree of inter-protomer packing and the stability of the GP1-GP2 interface.

### Acidic pH induces rearrangements of the RBR, the α2 helix and the CatL cleavage site in a species-specific manner

Next, we compared the HDX effects that the pH change from 7 to 5 induces in each individual trimer (**Figs. S4-S6**). These HDX effects describe GP conformational changes upon the virus transition from the cytosolic to the endo-lysosomal compartment. All GP trimers appeared overall more dynamic at pH 5 than pH 7, with most regions showing comparable magnitude of HDX increase across trimers, except for four areas (**Fig. 2**). These areas include the RBR α1 helix (residues 69-84, area 1), the region spanning the β13-β14 loop (residues 185-212, area 2), and the α2 helix in the GC (residues 255-261, area 3), and displayed HDX effects that are remarkably different between EBOV and SUDV GP (**Fig. 2c**). The β13-β14 loop of SUDV and EBOV harbours several residue changes (**Fig. S1**). Three histidine substitutions in the SUDV α2 helix (F248H, Y261H and S263H) very likely underlie its marked sensitivity to acidic pH, leading to a greater helical opening than in EBOV. The pH-associated switch of the α2 helix in Sudan GP also appears to induce significant conformational changes in the region spanning residues 302-334 that connects the GC and MLD (area 4), whose structure and topology is unresolved, but likely neighbours the α2 helix. Although the GC α2 helix has yet to have an assigned function, the β13-β14 loop and the RBR α1 helix are known to play a role in cathepsin cleavage^12,19^ and receptor binding^24–26^, and show clear inter-species divergences in structural dynamics.

**Fig. 2.**
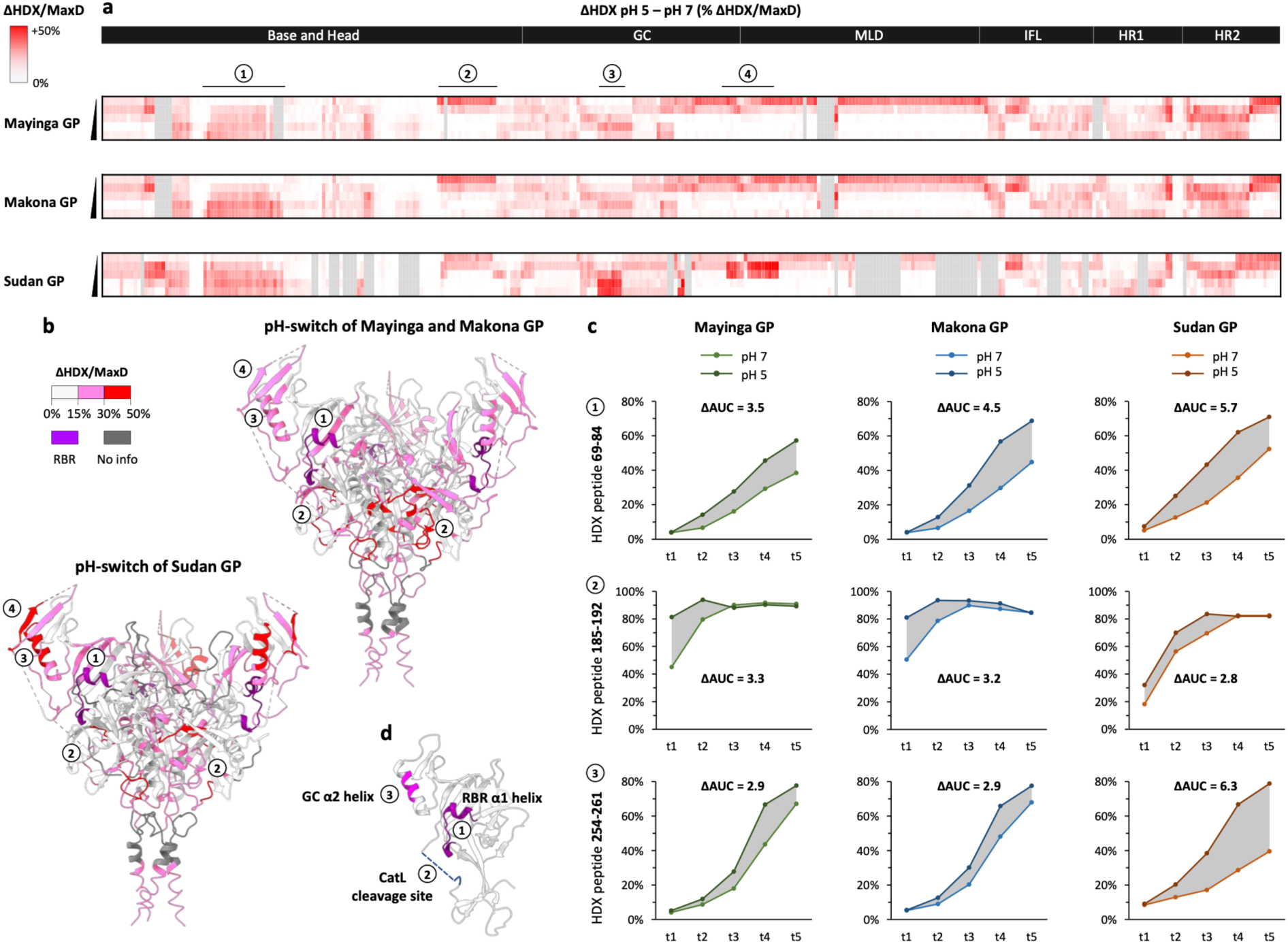
Differences in HDX (ΔHDX) between pH 7 and 5 for each individual trimer. **a,** Plots illustrating ΔHDX profiles of pH 5 minus pH 7 for Mayinga, Makona and Sudan GP. Positive ΔHDX values relative to the maximal deuteration (MaxD) are red colour-scaled based on the magnitude of the effect. **b**, ΔHDX profiles superimposed on GP structure illustrate areas of difference in ΔHDX upon change in pH across GP trimers. **c**, Deuterium uptake plots of reference peptides spanning areas of difference in ΔHDX across trimers upon change in pH. The ΔAUC values (areas between the deuterium uptake curves of pH 7 and pH 5) represent a cumulative magnitude of HDX effect. **d**, The areas of different ΔHDX across the three trimers are coloured on one protomer of the GP structure. Source data are provided within source data file.

### CatB and CatL cleave GP in multiple and distinct sites

Successively, by combining bottom-up MS and MP approaches, we comprehensively characterized the cleavage sites of CatB and CatL and their resulting GP intermediates. Conditions for optimal in-vitro cathepsin digestions were identified at pH ≤4.5 (**Figs. S7 and S8**, **Table S1**) and ratios 1:3 and 1:30 enzyme:GP trimer for CatB and CatL, respectively (herein, enzyme:GP ratios are referred to as 1x CatB/CatL). Under our experimental conditions, digestion of Makona GP with a mixture of 1x CatB plus 1x CatL yielded a product of ∼120 kDa, whereas individual digestions with 1x CatB and 1x CatL yielded products measuring ∼160 and ∼164 kDa, respectively (**Fig. 3c**). These mass measurements were conducted by MP and indicate the mass of whole cleaved trimers, while uncleaved GP trimer was measured at ∼300 kDa.

**Fig. 3.**
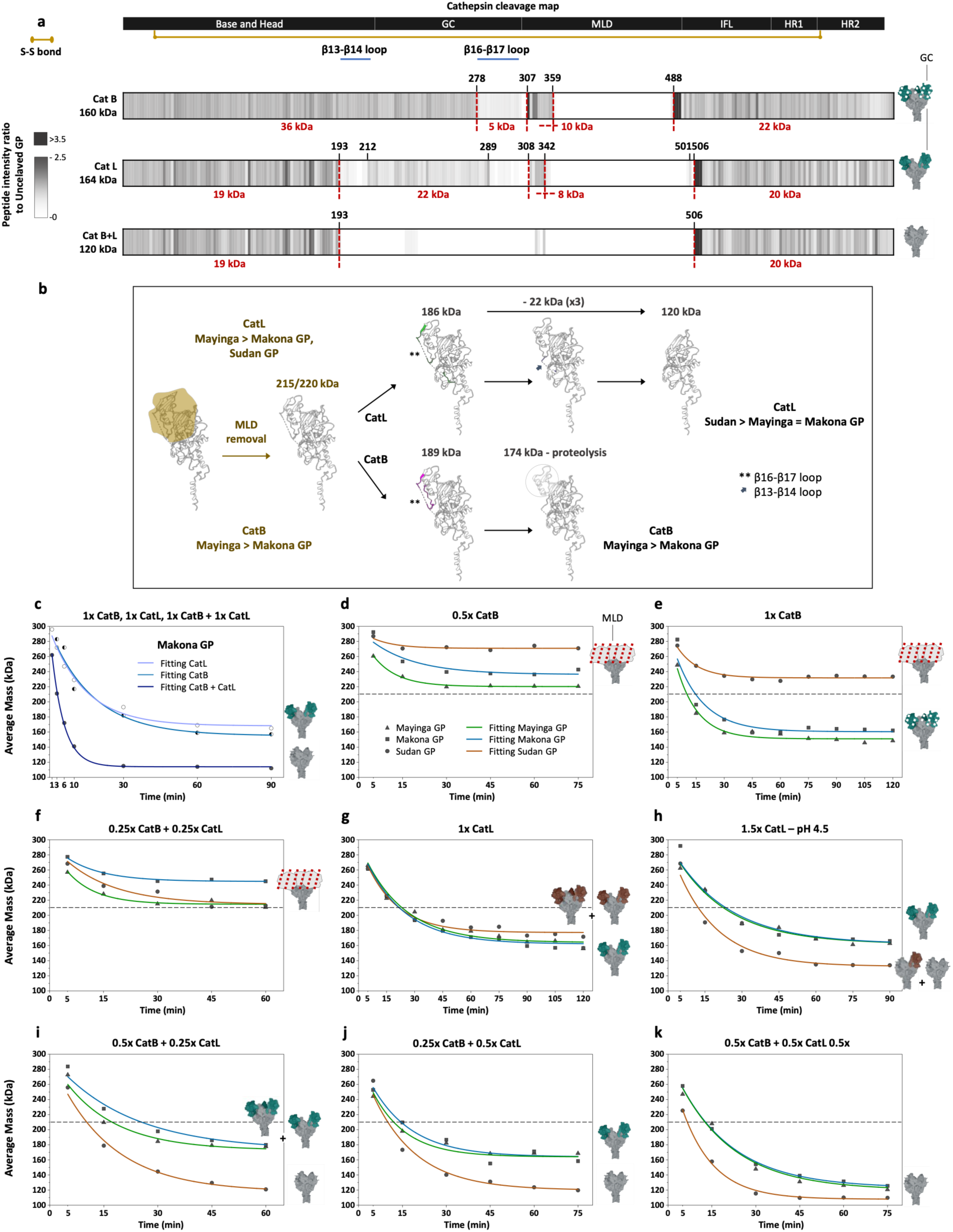
Cathepsin cleavage pathways. **a**, Peptide-level map of the sites cleaved by cathepsins. Numbers in black: residues after which cleavage occurs. Numbers in red: calculated mass (backbone + glycans) of specific segments (glycosylation assigned with nominal mass of 1.5 kDa). **b,** Schematic of the proteolytic pathways of CatB and CatL; cleaved segments and intermediates are represented on a single protomer of GP structure. **c-k,** Plots illustrating the mass of the three GP trimers measured by MP over the course of digestions under different cathepsin conditions. The dashed lines indicate the approximate mass of separation between the first and second digestion phases. Icons with GC in green: Mayinga and Makona GP. Icons with GC in brown: Sudan GP.

The MS analysis of the 160- and 164-kDa GP products identified not one unique cleavage site, but rather multiple sites susceptible to CatB and CatL cleavage (**Fig. 3a**). Both CatB and CatL can remove the MLD by cleaving GP between residues 359-488 and 342-501/506, respectively (**Figs 3a and S9**). The latter site cleaved by CatL is adjacent to the multibasic motif cleaved by furin, explaining how CatL can replace furin^44–46^. Furthermore, both CatB and CatL can digest the β16-β17 loop connecting GC and MLD, but CatL can also digest the β13-β14 loop connecting GC and head domain, unlike CatB (residue-level cleavage sites in **Fig. 3a**). The cleavage of both loops operated by CatL results in the removal of a 22-kDa segment comprising the GC and the β14 strand, which otherwise occlude the receptor binding cavity (PDB 5JQ3). Our data show that this 22-kDa segment is not only cleaved from the GP structure, but also further proteolyzed. The 120-kDa product, generated by mixing CatB and CatL, lacked all the three GC and MLD domains, due to cleavage between residues 193-506, and yields a monomeric mass for GP1 and GP2 of 19/20 kDa (**Figs. 3a and S10**).

The 120-kDa product represents the fully cleaved GP (GPcl), also referred to as 19-kDa GP^17,20^. The 164-kDa CatL product is thus the 120-kDa GPcl that still carries two uncleaved 22-kDa GC, while the 160-kDa CatB product is GP without the MLD, and proteolyzed in other parts of the structure that we could not unambiguously determine. Both products are long-live intermediates of the cathepsin digestion. Digestion with 2x and 3x CatL yielded a fast transition to 120-kDa GP, indicating that CatL alone can easily generate GPcl, unlike CatB (**Fig. S11, Table S1**), but the formation of 120-kDa GPcl is accelerated when CatL cleaves together with CatB (**Fig. 3c,e,g,k**). These observations suggest that CatL could be the necessary cathepsin and CatB the auxiliary cathepsin.

To understand the sequential order in which the two loops are cleaved by CatL, we digested Makona GP in the presence and absence of the therapeutic mAb REGN3470, which occludes an epitope encompassing the β16-β17 loop^47^ (**Fig. S12**). CatL digestion in the presence of this mAb yielded higher abundance of 120-kDa GPcl, indicating faster release of the 22-kDa GC. Therefore, under standard conditions, the cleavage of the β13-β14 loop follows the cleavage of the β16-β17 loop, and is thus the limiting proteolytic step. HDX data highlighted that the β13-β14 loop also displays different structural dynamics between EBOV and SUDV GP (**Fig. 2**).

### Cathepsin cleavage kinetics differ among species

To gain insight into the cleavage kinetics across Makona, Mayinga and Sudan GP, we digested the trimers under various cathepsin concentrations and combinations and monitored their decrease in mass over time by MP (**Figs. S13-15**, kinetic parameters in **Table S2)**. Our kinetic data and cleavage maps together reveal how cathepsin digestion proceeds in a stepwise fashion, and in two distinct phases. The first phase entails the removal of the MLD, leading to 215/220-kDa GP. The second phase entails the removal of the 22-kDa GC upon cleavage of the β16-β17 and β13-β14 loops and is only efficiently achieved by CatL (**Fig. 3b**). Varying the conditions of cathepsin digestion allowed us to distinguish the relative susceptibility of GP trimers in being cleaved at each phase.

We observed that for EBOV GPs, both CatB and CatL yielded efficient MLD removal, but Makona GP showed a lower cleavage rate than Mayinga GP under most conditions tested (**Figs. 3d-g**). Makona has 15 residues substitutions in the MLD compared to Mayinga GP, and those are seemingly linked to its reduced cleavage kinetics (**Fig. S1**). This disparity could also be due to a decreased cleavability by CatL of the 500-507 segment, which carries A503V and exhibited lower HDX in Makona than Mayinga GP (area 14 in **Fig. 1a,e**).

Strikingly, compared to EBOV, Sudan GP was significantly more resistant to MLD removal by CatB and strictly required CatL to fully digest the MLD (**Figs. 3d-g**). SUDV MLD differs from EBOV in 160 residues (**Fig. S1**) and evidently contains one or multiple sites recalcitrant to CatB. Additionally, the lower HDX observed in its 292-302 segment (area 12 in **Fig. 1b,f)** may underlie a decreased cleavability of the β16-β17 loop by CatB. However, when digestion was conducted under conditions of high cathepsin activity (mixture of CatB and CatL at high concentration or CatL alone at high concentration) (**Fig. 3h-k**), Sudan GP measured lower masses than EBOV GPs in the earliest stage of cleavage (5 min), indicating that SUDV MLD removal becomes faster than EBOV. This dual-mode kinetic behaviour reinforces the observation that SUDV MLD is recalcitrant to CatB specifically: its digestion is more favourable than EBOV when cleaved by CatL, and as well benefits more than EBOV from the CatB-CatL synergy.

The cleavage steps of the second phase, i.e. the removal of the three 22-kDa GC toward 120-kDa GPcl, occurred at significantly higher rate in Sudan than in EBOV GPs (k=0.9 and 0.5 min^-1^ respectively) (**Fig. 3k**). The kinetics of the second phase for Sudan GP was monophasic under most cathepsin conditions tested, meaning that the successive 22-kDa GC removal proceeded without significant energetic barriers (**Fig. 3g-j**). In contrast, this kinetics for both Makona and Mayinga GP was in most instances multiphasic, namely their digestion under several cathepsin conditions yielded a plateau at 164 kDa, corresponding to the removal of a single GC (**Fig. 3g-j and Fig. S16**). Our data for Makona GP cleaved with 2x CatL support a model whereby the removal of each GC requires twice as long for each successive cleaved protomer (**Fig. S17**). This suggests progressive rearrangement of the β13-β14 loop toward less accessible states over the course of cleavage occurring for EBOV GP, but not for Sudan GP.

### An allosteric axis linking the RBR and the CatL cleavage site via the GC α2 helix regulates cleavage

To gain insights on the conformational rearrangements during the cathepsin cleavage process, we compared the HDX of the 160-kDa CatB intermediate, the 164-kDa CatL intermediate and the 120-kDa GPcl with the uncleaved GP (Makona EBOV) at pH 5 (**Fig. 4a,b)**. First, we observed a marked increase in HDX in the α1 helix of the cleaved forms (area 1), which was the highest for 120-kDa GPcl, followed by 164-kDa and 160-kDa GP compared to uncleaved GP (ΔG_opening_=13.0, 16.4, 22.9, and 26.6 kJ/mol, respectively) (**Fig. 4a,b,e**, **Table S3**). This explicitly shows that cathepsin cleavage progressively opens the RBR, contradicting the conclusions of a previous report that examined cleavage effects at neutral pH^48^. The 164-kDa CatL intermediate was observed adopting two distinct conformations in the α1 helix: a more closed conformation that overlaps with the 160-kDa CatB intermediate and a more open conformation that overlaps with 120-kDa GPcl (**Figs. 4c and S18**), confirming that a fraction of its protomers is occluded by the 22-kDa GC and a fraction is not.

**Fig. 4.**
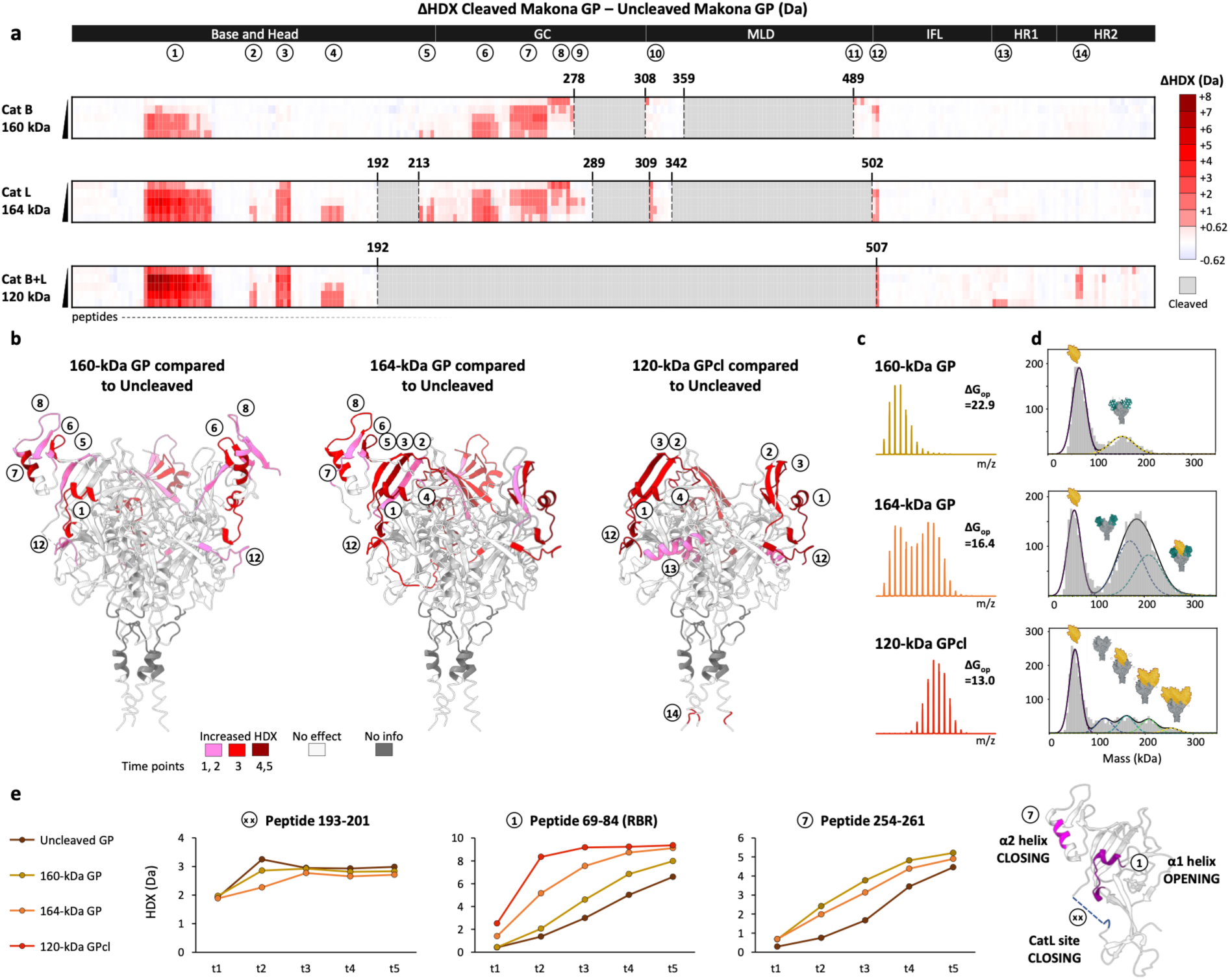
Differences in HDX (ΔHDX) and receptor binding across the cleaved forms (Makona EBOV). **a,** Plots illustrating ΔHDX profiles of the 160-kDa CatB intermediate, the 164-kDa CatL intermediate and 120-kDa GPcl minus uncleaved GP at pH 5. **b,** ΔHDX profiles are superimposed on GP structures (PDB 5JQ3) where cleaved segments have been removed. We note that HDX data are ensemble information and the effects of the intermediates result from an average of the three protomers (cleaved and uncleaved) within the trimer. **c,** HDX spectra of peptide 69-84 at time point 4 s exemplifies the structural dynamics and degree of opening of the RBR α1 helix for the 160- and 164-kDa intermediates and for 120-kDa GPcl. ΔG_opening_ (ΔG_op_) are indicated. **d,** MP spectra of the 160- and 164-kDa intermediates and 120-kDa GPcl in the presence of NPC1 at 1 µM (NPC1:cleaved GP 3:1) show different degree of binding. In the illustrations of the GP trimers, the yellow icons represent NPC1. **e,** Deuterium uptake plots of peptides spanning the allosteric triad α1 helix–α2 helix–CatL cleavage site exemplify the structural dynamics along this axis in the uncleaved GP, 160- and 164-kDa intermediates. **f,** The allosteric axis RBR α1 helix–α2 helix–CatL cleavage site superimposed on one protomer of GP structure. Source data are provided within source data file.

In addition to the RBR α1 helix, the GC α2 helix and the β13-β14 loop (CatL cleavage site) also exhibited differences in HDX across the cleaved products. Within Makona GP, the α2 helix of the 164-kDa intermediate (segment 255-261, area 7) adopted two distinct conformations (**Figs. 4e and S18**) with overall lower HDX than the 160-kDa intermediate, an effect that is opposite to what was displayed by the RBR helix (**Fig. 4e**). The 193-201 segment, which spans the β13-β14 loop, appeared more rigid (lower HDX) in the 164-kDa CatL intermediate than in the 160-kDa CatB intermediate, and significantly more rigid than in the uncleaved GP (**Fig. 4e**). Mechanistically, the opening of the RBR α1 helix thus occurs concertedly with the closing of the GC α2 helix and the rigidification of the β13-β14 loop. This signifies that this site in EBOV GP becomes increasingly less accessible for CatL as the cleavage progresses, which explains how the EBOV GC removal becomes progressively more difficult with each successive protomer of the trimer. In contrast, the HDX-MS analysis of a 140-kDa cleaved form of Sudan GP revealed that its 193-200 segment (β13-β14 loop) is more flexible than in uncleaved Sudan GP, and correspondingly its RBR α1 helix is more closed while its α2 helix is more open than in Makona 164-kDa GP (**Fig. S19**). This demonstrates how Sudan, unlike Makona GP, does not encounter energetic barriers in the successive removal of its GC domains. HDX data clearly identified a functional triad axis formed by α1 helix–α2 helix–CatL cleavage site in the GP trimer. This axis shows marked differences in structural dynamics between SUDV and EBOV GP (**Fig. 2**) and significantly influences their cleavage kinetics.

### Cleavage of the β13-β14 loop is necessary for receptor binding

The 160-kDa CatB intermediate and the 164-kDa CatL intermediate, despite being very similar in mass, have different degrees of cleavage (**Fig. 3a**). Both exhibited increased HDX in their GC (areas 6-9), resulting from the MLD removal. However, the 160-kDa intermediate displayed HDX characteristics denoting that its receptor binding cavities, formed by the α1 helix and the juxtaposed β7 and β9 strands^24–26^ (areas 1, 2 and 3), are all occluded by the GC, unlike in the 164-kDa intermediate and 120-kDa GPcl (**Figs. 4a,c**). This suggests that cleavage of the β16-β17 loop only, operated by CatB, does not induce a relevant shift of the GC from its original position. Accordingly, MP experiments conducted at pH 5 in the presence of NPC1 (domain C) indicated that the 160-kDa intermediate is not receptor binding competent, whereas the 164-kDa intermediate and 120-kDa GPcl can bind one and three NPC1 per trimer, respectively (**Fig. 4d**). This indicates that the cleavage of the β13-β14 loop, which occurs in 120-kDa GPcl and in one protomer of 164-kDa GP, but not in 160-kDa GP, is a critical requirement for receptor binding, confirming the idea that CatL is the necessary cathepsin.

### Makona GPcl exhibits the most open binding cavity and SUDV GPcl the most dynamic fusion peptide

When fully cleaved, GP trimers maximize the opening of their binding cavity, priming this site for receptor binding. The HDX of the RBR α1 helix and the β7 and β9 strands (areas 4, 5 and 6) informs on the degree of cavity opening (ΔG_opening_), which was the highest for Makona, followed by Sudan and Mayinga GPcl. Makona GP also displayed the greatest shift in opening upon transitioning from uncleaved GP to GPcl (**Fig. 5a, Table S3**). For the fully cleaved trimers, we still observed the potentiation of the RBR–fusion peptide axis for both Makona and Sudan GPcl compared to Mayinga GPcl (**Figs. 5b,c,d and S20-S22**). Unlike in EBOV, the transition from uncleaved GP to GPcl in SUDV showed to significantly mobilize its fusion peptide, pre-priming it for fusion (**Fig. S23**). This, together with the potentiation of the RBR-fusion peptide axis, resulted in Sudan GPcl having the most dynamic fusion peptide, followed by Makona and Mayinga GPcl (**Fig. 4b**).

**Fig. 5.**
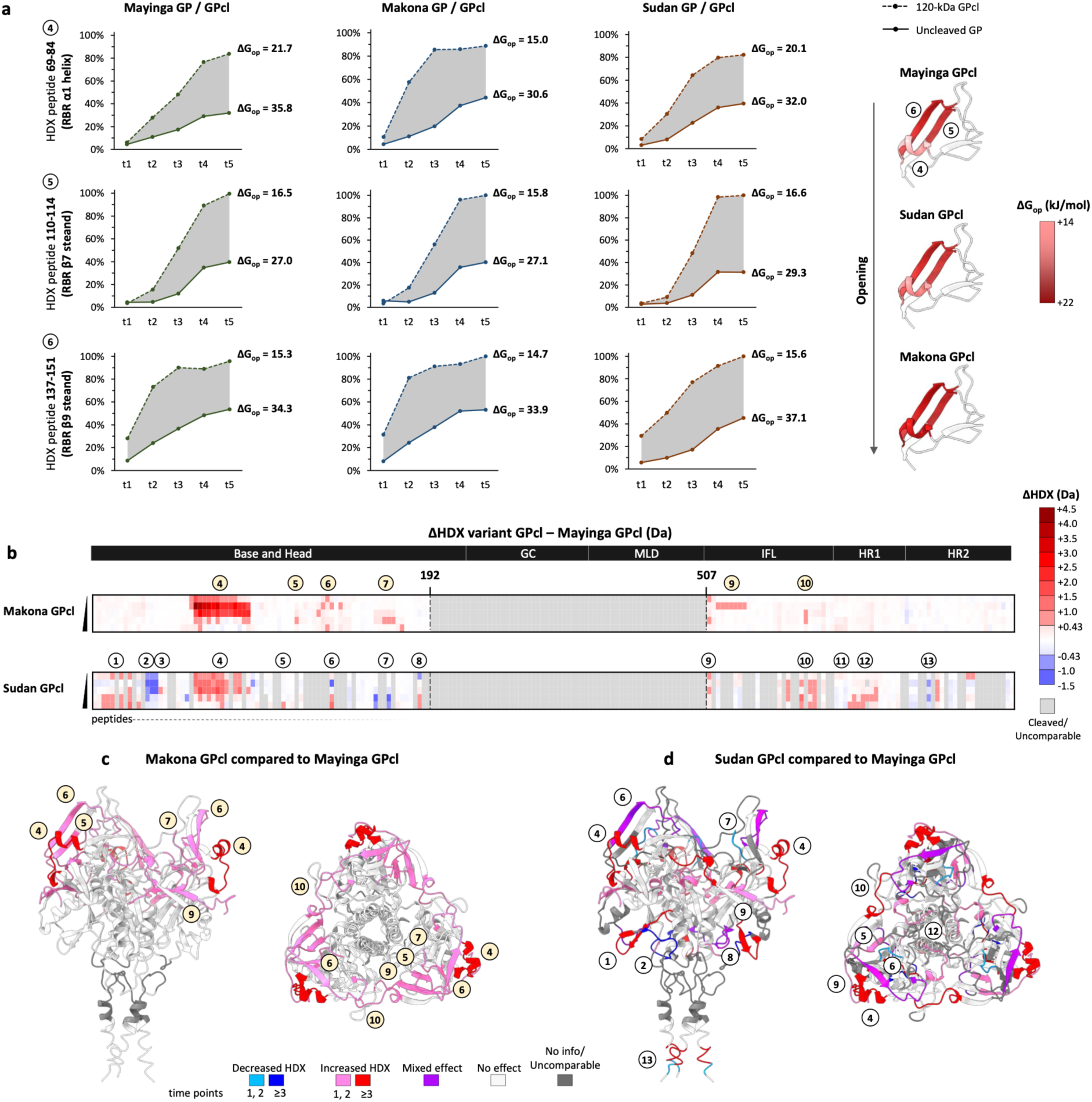
Differences in HDX (ΔHDX) across GP_cl_ trimers at pH 5. **a,** Deuterium uptake plots of model peptides spanning the RBR helix and β-strands for the three trimers illustrate their shift in opening of the binding cavity upon GP transitioning from uncleaved to fully cleaved (120-kDa GPcl). The calculated ΔG_opening_ are red colour-scaled and superimposed onto the binding cavity of the three trimers. **b,** Plots illustrating ΔHDX profiles of Makona minus Mayinga GPcl (top) and Sudan minus Mayinga GPcl (bottom). **c-d**, ΔHDX profiles Makona minus Mayinga GPcl (**c**) and Sudan minus Mayinga GPcl (**d**) superimposed on the GP structure where cleaved segments have been removed. Source data are provided within source data file.

### NPC1 binding is stronger but less cooperative in SUDV than in EBOV GPcl

Upon NPC1 receptor binding, all GPcl trimers displayed a marked decrease in HDX in the α1 helix as well as in the β7 and β9 strands and associated loops (residues 76-85, 138-151 and 111-122, areas 2, 4 and 5), in agreement with the published structures^24–26^ (**Figs. 6a,b and S24-S26**). In the RBR, Mayinga and Makona GP only differ in A82V in the α1 helix (**Fig. S1**). The difference in HDX between NPC1-bound and apo GPcl (ΔHDX) was greater in Makona than Mayinga (**Figs. 6a-c and S27**), yielding a ΔG_binding_ of 11.3 and 4.6 kJ/mol respectively, in agreement with the degree of opening of their binding cavity (**Table S3**). Sudan GPcl showed a greater ΔHDX than Makona GPcl (ΔG_binding_=23.9 kJ/mol, **Table S3**), despite its cavity being less open. Therefore, this increase in ΔG_binding_ stems from favourable residue substitutions in the SUDV binding cavity^26^; the highest increase was displayed by the β9 strand, which harbours V141A, S142Q and A148P (**Figs. S1 and S27**), and is thus the main driver of SUDV increased binding affinity.

**Fig. 6.**
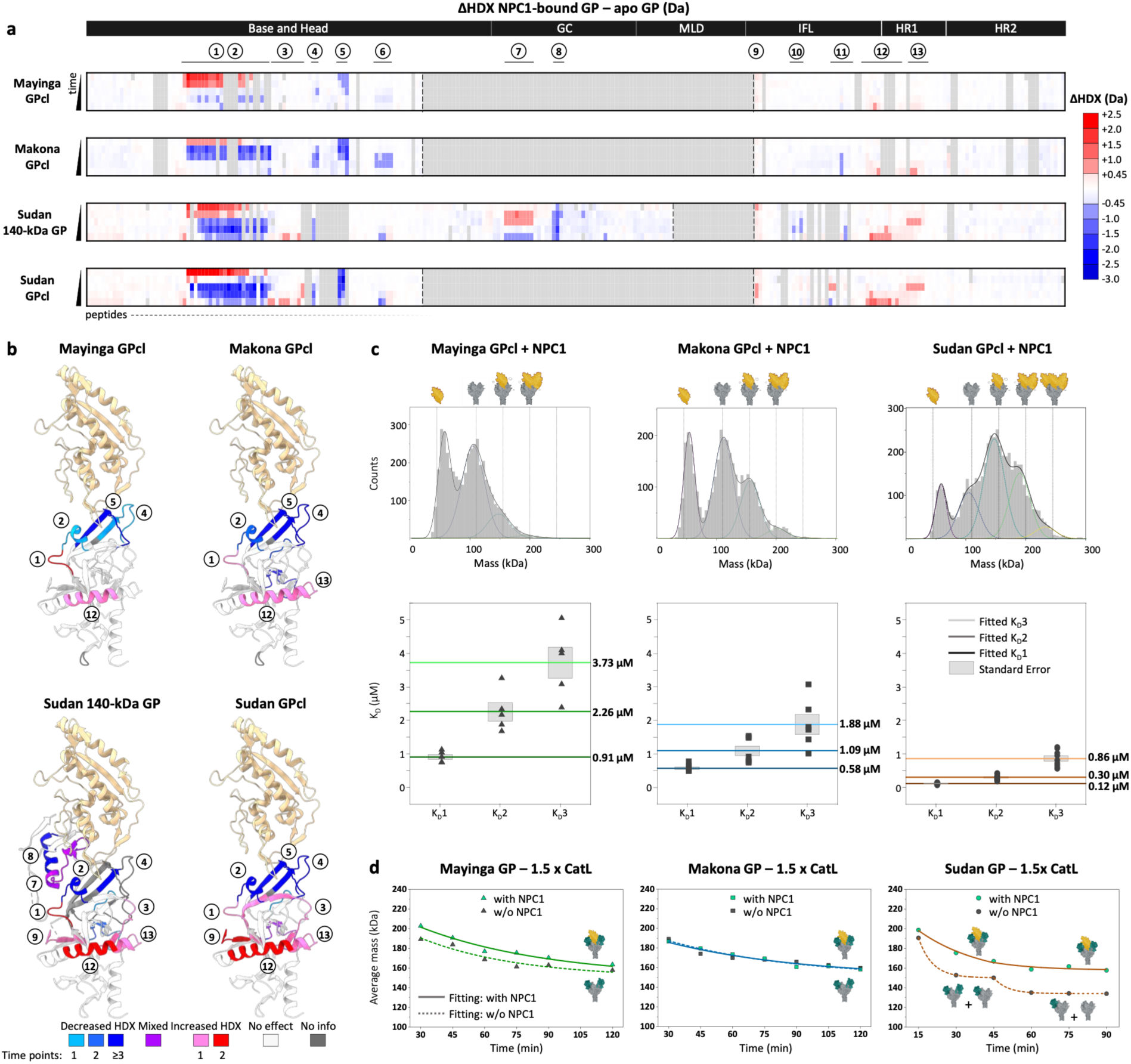
Differences in NPC1 binding across GP_cl_ trimers at pH 5. **a,** Plots illustrating ΔHDX profiles of Mayinga, Makona, Sudan 120-kDa GPcl and 140-kDa Sudan bound to NPC1 minus apo. **b**, ΔHDX profiles NPC1 bound minus apo superimposed on the structure of a bound protomer (PDB 5F1B). MP spectra of Mayinga, Makona, Sudan GPcl in the presence of NPC1 at 0.5 µM (NPC1:GPcl 3:1) show differences in binding across trimers. **c**, Plots illustrating the three KDs of binding for Mayinga, Makona, Sudan GPcl, fitted from MP experiments conducted in the presence of NPC1 (NPC1:GPcl 3:1). The average K_D_n/K_D_n+1 ratio is defined as the average of K_D_1/K_D_2 ratio and K_D_2/K_D_3 ratio. **d,** Plots illustrating the mass of GP trimers measured by MP over the course of the cathepsin cleavage reactions conducted in the presence and absence of NPC1. Source data are provided within source data file.

MP interaction experiments^42^ of GPcl with NPC1 at pH 5 enabled the calculation of three distinct K_D_s per trimer (one for each binding event), which yielded an average K_D_ at 2.3 µM for Mayinga, 1.8 µM for Makona and 0.42 µM for Sudan GPcl (individual K_D_ values in **Fig. 6c, Fig. S28**). These values are in strong agreement with the ΔG_binding_ measured by HDX, unambiguously indicating that Sudan GPcl has the strongest receptor binding affinity, followed by Makona and Mayinga GPcl. We note that the K_D_s measured at pH 5 for GPcl generated by mixing CatB and CatL, or generated by CatL alone, were comparable (**Fig. S29**), and significantly lower than the previously reported K_D_ sampled under neutral conditions (pH 7 and 6)^24–26^. Accordingly, the K_D_s of GPcl-NPC1 decreased on average 2.5 times from pH 7 to 5, indicating a pH-dependent interaction with the greatest boost in affinity at pH 5 (**Fig. S30**).

Our individual K_D_ values (reporting on the relative probability of the three distinct binding events within each trimer) uncovered positive cooperativity for receptor binding in EBOV GPcl. The average K_D_n/K_D_n+1 ratio for Mayinga and Makona GPcl were respectively 0.48 and 0.55 (**Fig. 6c**), hence greater than the K_D_n/K_D_n+1 ratio of 0.33 expected for a trimer with three equivalent binding sites^49^. This signifies that, for EBOV GP, NPC1 binding to a protomer facilitates the binding of the subsequent one/-s by promoting the opening of their binding cavity. In contrast, NPC1 binding to Sudan GPcl exhibited only minimal cooperativity (average K_D_n/K_D_n+1=0.37) (**Fig. 6d**). Protomers’ dynamics and binding events are thus inter-dependent in EBOV GP but largely independent in Sudan GP. This is potentially due to the lower degree of inter-protomer packing of Sudan GP and GPcl compared to EBOV GP and GPcl (area 13 in **Fig. 5b,c** and area 16 in **Fig. 1b,e**).

Additionally, we investigated the role of the 22-kDa GC in the NPC1 binding of partially cleaved trimers (**Fig. S31**). The K_D_1 of 164-kDa Makona GP (which carries two protomers occluded by the GC) was double (0.97 µM) than that determined for Makona 120-kDa GPcl, which suggests that the fully cleaved protomer is subjected to a closing effect induced by the two uncleaved GCs and confirms that the binding events are inter-dependent in EBOV. In contrast, the K_D_1 of a 155-kDa Sudan GP was equivalent (0.11 µM) to that of Sudan 120-kDa GPcl, indicating again that the receptor binding events are independent in Sudan GP.

### NPC1 binding hinders cathepsin cleavage to different extents across GP trimers

MP experiments revealed that full cleavage to 120-kDa GPcl is not required for receptor binding. As such, a partially cleaved form of Sudan GP of 140 kDa, which carries one of the three RBRs occluded by the GC, displayed ΔHDX binding effects in the same areas as the fully cleaved trimers, but expectedly of lower magnitude than Sudan 120-kDa GPcl (-30% ΔG_binding_) (**Figs. 6a,b**, **S32 and S33a**). Strikingly, the GC that remains on one of the protomers displayed significant dynamic changes upon binding, including a marked decrease in HDX in the α2 helix (residues 255-265, area 8). These HDX effects suggest that the α1 helix–α2 helix–CatL cleavage site axis is altered upon receptor binding, possibly influencing the kinetics of cleavage.

To explore this, we digested GP trimers with 1.5x CatL in the presence and absence of NPC1 and measured their decrease in mass over time by MP. The cleavage rates of Mayinga and Sudan GP were reduced by the presence of NPC1, but in Sudan GP this reduction corresponded to 22 kDa of mass difference, while in Mayinga GP it was minor and approximated 9 kDa. (**Fig. 6d and S34**). However, the cleavage rate of Makona GP were equivalent in the presence and absence of the receptor when digestion was conducted with 1.5x CatL (**Fig. 6d**). A reduction in cleavage rate for Makona GP was only clearly observable when digestion occurred at a lower rate (0.75x CatL) and minimally when it occurred at a higher rate (2x CatL) (**Fig. S34**). Taken together, our data show that cathepsin cleavage and receptor binding can occur simultaneously and that their kinetics and efficacy are inter-dependant. We observed that NPC1 engagement is hindered in partially cleaved Makona GP, but it is fully enabled in partially cleaved Sudan GP (**Fig. S31**). Therefore, Sudan GP can strongly engage the receptor while cleavage is occurring, which in turn reduces cleavage, but Makona GP cannot. Based on our data, in the presence of NPC1, Sudan GP is more readily cleaved (k=0.8 min^-1^) than EBOV GPs and has the strongest receptor engagement, while the cleavage of Makona and Mayinga GP is very similar (k=0.4 and 0.3 min^-1^ respectively) but Makona has stronger receptor engagement than Mayinga GP (**Figs. S35 and 6c**). Therefore, the transition to the fully cleaved/fully receptor bound state appears most favourable for Sudan, followed by Makona then Mayinga GP.

### Fusion priming is potentiated in SUDV compared to EBOV GP

GP trimers analysed in this study are pre-fusion stabilized via a proline substitution in the RR1 hinge loop, therefore their full fusogenic transition is inhibited. Nevertheless, upon NPC1 binding, they exhibited conformational fluctuations that are hallmarks of fusion priming, displayed as increased dynamics (increased HDX) in their fusion machinery elements: HR1 and the fusion peptide (**Fig. 7a**). Two allosteric axes were identified based on the effects captured for Sudan GP, for which the full set of dynamic changes were unambiguously observed (**Fig. 7b**). An intra-protomer axis transmits the binding energy from the RBR to the HR1, via the destabilization of the β3-α1 loop (residues 70-75, area 1) that bridges the α1 helix and the 507-517 GP2 segment (area 9), which abuts and in turn mobilizes the HR1 helices and hence the RR1 hinge loop (residues 555-581, areas 12 and 13). This pathway is shared with the RBR-fusion peptide axis, which instead transmits effects of rigidification from the 507-517 GP2 segment to the fusion peptide (residues 530-542, area 11). Secondly, an inter-protomer axis transmits effects of mobilization from the RBR to the fusion peptide via the interposed β4 strand and β4-β5 loop (residues 86-109, area 3). At the structural level, these conformational dynamic changes reflect a decrease of inter-protomer stability via the mobilization of the RR1 hinge loop (trimer opening), the unleashing of the HR1 and the unclamping of the fusion peptide. These are in line with the structural rearrangements known to occur during the fusogenic transition of fusion glycoproteins^50^ and previously visualized by HDX-MS for the SARS-CoV-2 spike^33^.

**Fig. 7.**
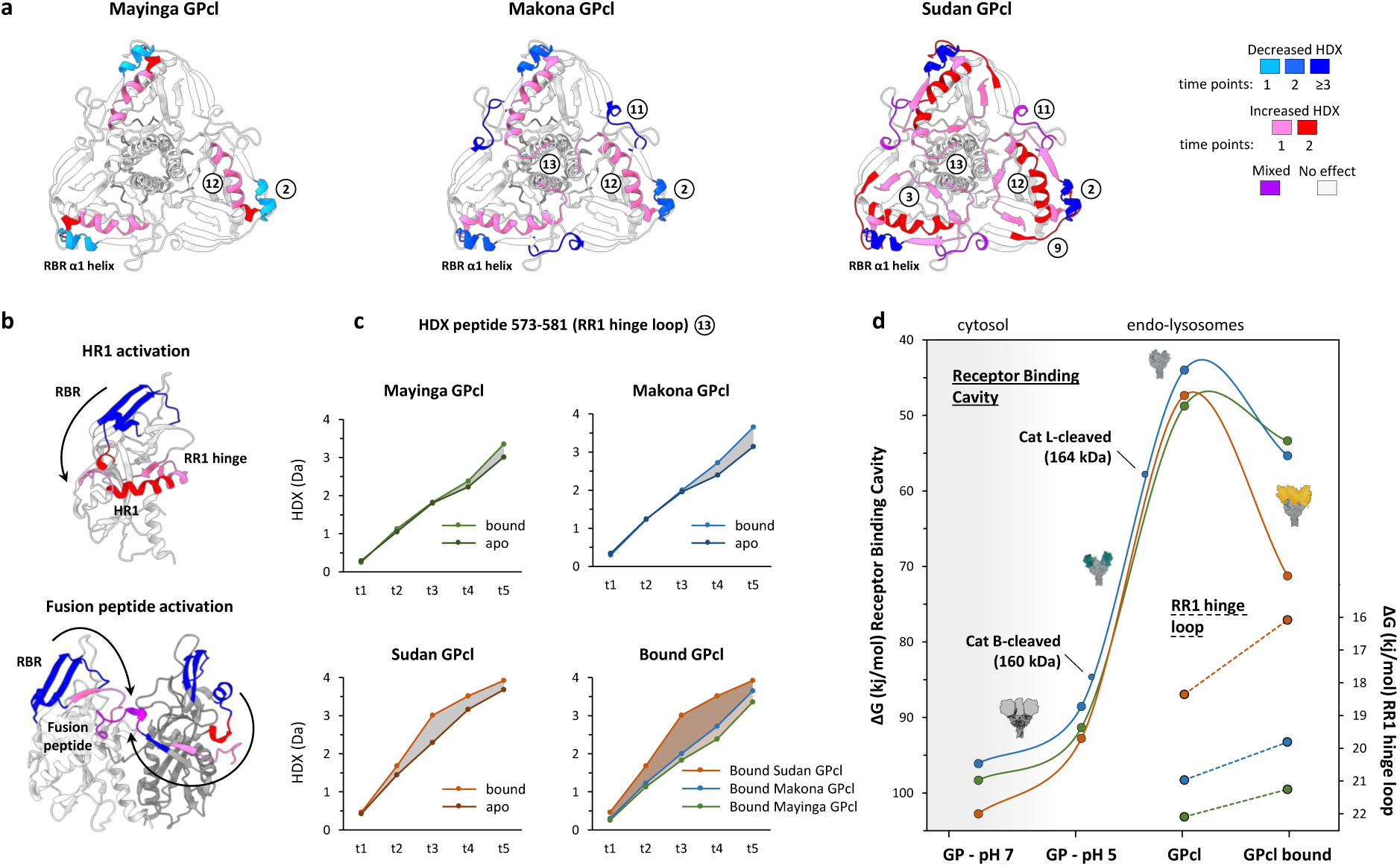
HDX effects of fusion priming. **a,** ΔHDX profiles of NPC1 bound minus apo superimposed on the GP structure (PDB 5JQ3) show increased HDX in the elements of the trimer fusion machinery. **b,** Allosteric axes of fusion priming are illustrated along one (top) or two protomers (bottom) of the structure of GP. **c,** Deuterium uptake plots of a model peptides spanning the RR1 hinge loop exemplify the magnitude of the priming effect across different trimers. **d,** Plot illustrating the energetic transitions of the binding cavity and the RR1 hinge refolding loop for Makona, Mayinga and Sudan GP in terms of ΔG_opening_ along different stage of the entry pathway. Source data are provided within source data file.

While these conformational changes are qualitatively equivalent in every trimer, the magnitude of the observed HDX effects scaled with Sudan>Makona>Mayinga GPcl. Sudan 140-kDa GP, despite not being fully cleaved, exhibited these effects at higher magnitude than Makona GPcl (**Fig. S33b,c**). In **Fig. 7d**, we illustrate the conformational transitions along the entry pathway of the binding cavity and of the RR1 hinge loop in terms of ΔG_opening_, demonstrating that the priming effect on the RR1 hinge loop correlates with the ΔG_binding_ (**Table S3**). Additionally, the RR1 hinge loop and the fusion peptide are baseline more dynamic (lower ΔG_opening_) in Sudan GPcl than EBOV GPcl (area 11 and 13 in **Fig. 5b,c,d**), altogether unveiling that Sudan GP is more adapted for fusion than the EBOV GP trimers.

## Discussion

By integrating HDX-MS and mass photometry, we reveal a direct quantitative link between the structural dynamics of the Ebola virus GP trimer and its three functional transitions occurring during entry: proteolytic cleavage by endosomal cathepsins, engagement of the NPC1 receptor and fusion priming. By examining Makona, Mayinga and Sudan GPs, we discover not only that these steps are mechanistically coupled by allosteric axes, but also that differences in GP sequences tune the structural dynamics of these axes, redistributing the energetic and kinetic bottlenecks of entry.

We here unveil that cleavage and receptor binding are not strictly sequential events, but can occur simultaneously and continuously feedback on one another – and this allosteric feedback is species-and variant-specific. This model is founded upon three findings. First, we show that progressive cathepsin cleavage is accompanied by a stepwise opening of the RBR α1 helix, in contrast to the previously reported model^48^. The α1 helix opening is mechanistically coupled, via the GC α2 helix, with a structural reorganization of the CatL cleavage site, where each round of proteolysis affects the accessibility of this site for the subsequent cleavage. This coupling makes the GC removal progressively harder for EBOV GP as cleavage proceeds, but progressively easier for SUDV GP. Second, we demonstrate that partially cleaved trimers are legitimate receptor-binding competent intermediates, rather than requiring full cleavage for NPC1 engagement. Third, we found that the degree of GP cleavage can negatively affect NPC1 affinity (an effect that is more pronounced in EBOV), and NPC1 binding can reciprocally slow further cleavage (an effect that is more pronounced in SUDV). Under this model, increasing NPC1 affinity is only advantageous if matched by a compensatory loss of cleavage–binding coupling, as achieved by Makona GP, or by a gain in cleavage efficiency, as achieved by Sudan GP. This interdependence may constrain the evolutionary trajectories available to Ebola virus GP, requiring cleavage and receptor engagement to co-adapt for a new variant or species to establish.

As for other fusion glycoproteins^33,51,52^, we found that receptor binding is allosterically coupled to fusion priming in Ebola virus GP. Compared to Mayinga, both Makona and Sudan GP potentiate an RBR–fusion peptide allosteric axis that couples the opening of the receptor-binding cavity to the destabilization of the internal fusion loop, improving both receptor engagement and fusion peptide mobility. A structurally analogous axis links opening of the SARS-CoV-2 spike trimer to mobilization of its fusion-peptide-proximal region, and the potentiation of this axis by D614G^33,53,54^ contributed to the emergence of variants of concern. The A82V substitution that defines Makona GP offers a direct test of this mechanism against an independent, orthogonal dataset based on molecular dynamics simulations and single-molecule FRET^55^. Durham et al.^55^ showed that A82V unwinds the α1 helix, re-orients W86 and destabilizes the fusion loop, driving GP out of the prefusion conformation hence increasing GP-membrane engagement. Our multi-species study shows that this destabilization strategy is not unique to A82V in Makona GP, as SUDV GP, which lacks this substitution but carries its own cluster in the α1 helix (N73S, A76S, V79I) and in the IFL (including T544 shared with Makona GP), potentiates this axis even more strongly, to the point of directly mobilizing the fusion peptide itself. Ebola virus GP has therefore converged on the same destabilization mechanism at least twice, once through the A82V substitution acquired by the EBOV Makona variant, which gave rise to the 2013–2016 epidemic, and independently through several substitutions present in the Sudan species. Furthermore, SUDV is equipped with higher mobility in its RR1 hinge loop, which allows for extensive priming of its stalk helices upon receptor engagement, in a manner that is similar to SARS-CoV-2 omicron spike^33^. The greatest mobility of the fusion peptide and RR1 hinge loop together make SUDV GP the trimer most adapted for fusion among those examined.

We also show that GP proteolytic priming is far more complex than previously understood. We characterized multiple sites cleaved by cathepsins beyond the β13-β14 loop and residue 502^19^ so far identified, and show that cleavage at these additional sites results in the generation of proteolytic intermediates of masses higher than GPcl and capable of NPC1 binding, thus functional for virus entry. The structural analyses of these intermediates indicates that CatL is the necessary cathepsin and CatB the auxiliary cathepsin, in apparent contrast to Chandran et al.^16^ but in agreement with Schornberg et al.^17^ We also elucidate how EBOV and SUDV diverge at the level of cathepsin cleavage kinetics and susceptibility. While EBOV GPs differ only modestly in their kinetics, largely through a slightly reduced MLD cleavability in Makona compared to Mayinga GP, Sudan GP is markedly more resistant to CatB and its MLD removal depends almost entirely on CatL. SUDV compensates for this via substitutions along the α1 helix–α2 helix–CatL axis that keeps the CatL cleavage site accessible, so that once cleavage begins, SUDV GP sheds its GC domains and engages NPC1 more readily than EBOV GP. SUDV thus trades a higher barrier to initiate cleavage for a lower barrier to complete it and to bind NPC1.

Altogether, these data identify the allosteric axes governing specific steps of Ebolavirus entry and show how sequence divergence among circulating species/strains affect a defined number of structural nodes that couple dynamics to function. These axes thus represent tractable and mechanistically defined targets for antibody-based therapeutics and for rational structure-guided design of vaccine immunogens, against both circulating Ebolavirus species and strains that have yet to emerge.

## Materials and methods

### Protein production

The cDNA encoding for Mayinga, Makona and Sudan GP were based on a version of GP ectodomain (residues 1-647) with two stabilizing substitutions (T577P and K588F)^43^ and a C-terminal 6His-tag. The cDNA encoding for domain C of NPC1 (residues 374–620)^25^ were cloned into a pcDNA3.1(+) vector with a C-terminal 6His-tag (Genscript). Proteins were expressed in human embryonic kidney (HEK) 293F cells (FreeStyleTM, ThermoFisher Scientific) growing in suspension at 37 °C in an 8% CO2 atmosphere in FreeStyleTM expression medium, transfected at confluency 10^6^ cells/mL with OPTIMEMTM (Gibco) and 1 mg of DNA per litre of culture. The supernatant (containing the proteins) was harvested by centrifugation on the fourth day post-transfection, and clarified with 0.45 µm pore-size filters (Merck). His-tagged proteins were purified using a HisTrap HP 5 mL column (Cytiva) connected to an AKTA pure protein purification system (Cytiva) and by size-exclusion chromatography (SEC) on a Superose 6 Increase 10/300 GL column (GE Healthcare) equilibrated with DPBS pH 7.4 (ThermoFisher Scientific). Proteins were concentrated to a final concentration of ∼10 mg/mL for the GP trimers and 4.8 mg/mL for NPC1, in PBS.

### HDX-MS experiments

For peptide mapping, Mayinga, Makona and Sudan GP trimers were diluted in PBS and subjected to the same digestion protocol and liquid chromatographic gradient as detailed below. MSE analysis was performed with a Synapt G2-Si mass spectrometer (Waters), applying collision energy ramping from 20 to 30 kV. Sodium iodide was used for calibration and leucine enkephalin was applied for mass-accuracy correction. MSE runs were analysed with ProteinLynx Global Server (PLGS) 3.0 (Waters), and peptides identified in 4 out of 5 runs, with at least 0.2 fragments per amino acid and at least 2 fragments in total, and with mass error below 10 ppm, were selected in DynamX 3.0 (Waters). The GP trimers were mapped with integration of glycoproteomics data (source data) to identify glycosylated peptides. HDX experiments were conducted with GP trimers (Makona, Mayinga, Sudan GP) at 5.7 µM (trimer concentration) in PBS (pH 7.4) or acetic buffer (pH 5), diluted 1:10 in a deuterated PBS buffer (pHread=7) or deuterated acetic buffer (10 mM acetate, 150 mM NaCl, pHread=5) to start the exchange reaction. The time points studied were the following: pH 7: t1=4 s (ice), t2=4 s (23°C), t3=0.6 min (23°C), t4=6 min (23°C), t5=12 min (28°C), t6=1 h (23°C), t7=10 h (23°C). pH 5: t1=400 s (ice), t2=400 s (23°C), t3=1 h (23°C), t4=10 h (23°C), t5=20 h (28°C). Following the time window expansion method^34,56,57^, the time points at pH 5 were adjusted of a factor x100 relative to the time points at pH 7, enabling their comparison. For the analysis of cleavage products, the uncleaved Makona GP, the 164-kDa CatL intermediate, the 160-kDa CatB intermediated and the 120-kDa GPcl, all at 5.7 µM, were diluted 1:10 in the deuterated acetic buffer (pHread=5) and the HDX reaction were conducted for the following time points: t1=400 s (ice), t2=400 s (23°C), t3=1 h (23°C), t4=10 h (23°C), t5=20 h (28°C). For interaction experiments, GP trimers (Mayinga, Makona, Sudan GPcl and 140-kDa Sudan GP) were mixed at ratio 1:3 with NPC1 C-domain (final concentrations of GP trimers at 5.7 µM and NPC1 at 17 µM in the acetic buffer at pH 5) and let equilibrate for 1 h. In parallel, GP trimers were incubated alone in the acetic buffer. The exchange reactions were conducted upon 1:10 dilution in the deuterated acetic buffer (pHread=5) and the time points studied were the following: t1=400 s (ice), t2=400 s (23°C), t3=1 h (23°C), t4=10 h (23°C), t5=20 h (28°C). All exchange reactions were quenched by a 1:1 dilution (vol/vol) with an ice-cold 100 mM phosphate buffer at 3 M urea and 200 mM TCEP (pHread=2.3). Samples were held for 30 s on ice and snap-frozen in liquid nitrogen. Maximally-labelled (MaxD) controls were performed by diluting the GP trimers in a deuterated buffer at 3M Urea and 5 mM TCEP, yielding a final D_2_O fractions at 90%, as for the other deuterated samples. The exchange reaction was conducted for 10 h and quenched 1:1 with an ice-cold 100 mM phosphate buffer (pHread=2.3). MaxD samples were held for 30 s on ice and snap-frozen in liquid nitrogen. Frozen samples were quickly thawed and injected into an Acquity UPLC M-Class System with HDX Technology (Waters). Injected samples first passed at 120 µL/min through a home-made pepsin column at 1.5°C and then through a Nepentesin-2 column (AffiPro) at 15°C for on-line digestion. Peptides were trapped/desalted with solvent A (0.23% formic acid in water, pH 2.5) for 4 min at 120 μL/min and at 1.5°C through an Acquity BEH C18 VanGuard pre-column (1.7 μm, 2.1 mm × 5 mm, Waters). Peptides were eluted into an Acquity UPLC BEH C18 analytical column (1.7 μm, 2.1 mm × 50 mm, Waters) with a 7-min linear gradient rising from 8% to 35% solvent B (0.23% formic acid in acetonitrile) at a flow rate of 90 μL/min, at 1.5°C. Peptides were then subjected to electrospray ionization in positive mode and MS analysis with ion-mobility separation. Triplicates were conducted at all time points and in every state. Peptide-level deuterium uptakes were calculated with DynamX 3.0 (Waters), after data were visually inspected and curated. The difference in deuterium incorporation between peptides carrying residue substitutions across GP trimers were calculated as previously described^33^, selecting Mayinga GP as reference state. The threshold for statistically significant differences in HDX was established at the 99% confidence level, based on an approach described previously^58^. As per community-based recommendations^59^, in tables S4-S10 a summary of the HDX-MS experiments is provided.

### Derivatization of ΔG_opening_ from HDX data

Peptides spanning the receptor binding cavity and the RR1 hinge loop were amened to determination of the ΔG_opening_ across GP trimers (Table S3). These peptides are 69-84 (α1 helix), 77-88 (α1 helix), 110-114 (β7 strand), 137-151 (β9 strand), 573-581 and 574-581 (RR1 hinge loop). Their deuterium incorporation, expressed as a percentage of MaxD (D%), was fitted across time points t1-t5 with the stretched-exponential function D%(t)=1−exp[−(kt)^β^], where β is a stretching factor that captures the range of individual amide exchange rates that are averaged at the peptide level, hence the resulting deviations from a simple exponential behaviour. Time points t1 (ice) and t5 (28°C) were converted into time points at 23°C, multiplying them by 0.1 and 2, respectively^56^. The k_HX_ was determined as k_HX_=k(ln2)^1−1/β^. Peptide-level protection factors (pf) were calculated as pf=k_ch_/k_HX,_ where the k_ch_ was calculated based on the reference parameters for protein hydrogen exchange rates^60^. The ΔG_opening_ was calculated as ΔG_op_=RTln(pf)^61^, where T=296.15°K.

### Optimization of CatB and CatL digestion conditions

Recombinant human CatB (2.5 U/µg) and human CatL (25 U/µg) were purchased from Bio-techne (United Kingdom). Both CatB and CatL were activated for 25 min at 37°C in acidic buffer before starting the digestions of the GP trimers, which were always conducted at 37°C. First, digestions of Makona GP at pH 5, 4.5 and 4 (acetic buffers) were conducted to test the activity of CatB and CatL depending on pH. An aliquot of the reaction mixture was removed at defined time points, ranging between 1 and 120 min, quickly diluted in an acetic buffer (pH 5) and analysed by MP at a final concentration of ∼8 nM GP trimer. MP measurements and data analysis were conducted as detailed in *Kinetic MP experiments.* Optimal pH was determined at pH=4 and pH≤4.5 for CatB and CatL, respectively (Fig. S7). To quantify the variation in cathepsin cleavage efficiency over time (pH 4), Makona GP was digested for 5 min after the cathepsins incubated alone for different time periods (from 5 to 120 min) at 37°C. MP measurement were conducted as detailed above and the ratio between the mass of GP digested with the enzymes incubated for x min and 5 min was used to determine the cleavage efficiency. This remained significant (>55%) up to 60 and 120 min for CatB and CatL, respectively (Fig. S8). Baseline digestion conditions were established at 1:3 CatB:GP trimer and 1:30 CatL:GP trimer, where GP trimer is at 3.3 µM in the reaction mixture, referred to as 1x CatB and 1x CatL, respectively. These ratios were selected as they correlate with the relative activity of CatB and CatL and yielded measurable cleavage in the timeline of cathepsins retaining appreciable activity.

### Identification of cathepsin cleavage sites

The 160-kDa intermediate digested by 1x CatB, the 164-kDa intermediate digested by 1x CatL and the 120-kDa product digested by a mixture of CatB and CatL were subjected by bottom-up mass spectrometry analysis upon digestion with non-specific proteases (pepsin, nepenthesin) as performed for HDX-MS experiments and analysed by PLGS and DynamX (Waters). The use of non-specific proteases for protein digestion, combined with open-termini PLGS software analysis, enabled to accurately identify the peptide termini, hence the cathepsin cleavage sites. This approach avoided the constraints of pre-determined sites of cleavages given by the more specific proteases used in standard bottom-up proteomics, which confound the information on peptide termini. The peptide signal intensity of the 164-kDa, the 160-kDa and the 120-kDa GP were compared to the signal intensity in the uncleaved GP, to quantify the extent by which segments are cleaved. The peptide signals in fifteen MS runs for each state were averaged and compared. To confirm the CatB and CatL cleavage sites, the reaction mixtures containing the 160-kDa and the 120-kDa products were also subjected to SEC purification; the SEC fractions of the main peak were pooled, de-glycosylated with PNGase F, and then analysed as detailed above.

### Investigation of mAb REGN3470 effect on cathepsin cleavage

Makona GP trimer and mAb REGN3470 were mixed at ratio 1:3 and let incubate for 30 min in PBS at concentrations 4.8 µM and 14.4 µM, respectively. In parallel, Makona GP was incubated alone in PBS. The HDX reaction was initiated upon 1:10 dilution in deuterated PBS buffer (pHread=7) and allowed to proceed for 4 s, 0.6 min, 6 min, 1 h and 10 h at 23°C. The exchange reactions were quenched and samples analysed as detailed above. HDX-MS data analysis was conducted as detailed above. Makona GP trimer was digested at 37°C with 0.75x CatB and 0.75x CatL in an acetic buffer at pH 4, in the presence and absence of mAb REGN3470 (ratio 1:3 GP trimer:mAb). At selected time points (5, 15, 30, 45 and 60 min), an aliquot was withdrawn from the reaction mixture and subjected to non-reducing SDS-page.

### Kinetic MP experiments

To determine cleavage kinetics, Makona, Mayinga and Sudan GP and cathepsins were mixed at various concentrations and combinations of CatB and CatL. These conditions were: 0.5x CatB, 1x CatB, 1x CatL, 1.5x CatL, 0.25x CatB + 0.25 CatL, 0.5x CatB + 0.25x CatL, 0.25x CatB + 0.5x CatL, 0.5x CatB + 0.5x CatL; all conducted at pH 4, except for 1.5x CatL that was conducted at pH 4.5. Makona GP was also digested with 2x and 3x CatB and 2x and 3x CatL (pH 4). An aliquot of the reaction mixture was withdrawn at defined time points, ranging between 1 and 120 min, quickly diluted in an acidic buffer (pH 5) and analysed by MP at a final concentration of ∼8 nM GP trimer. For testing the inhibition of cathepsin cleavage upon receptor binding, Makona, Mayinga and Sudan GP were also digested with 1.5x CatL (pH 4.5) in the presence and absence of 10 µM NPC1 (1:3 GP trimer:NPC1). The GP trimers and NPC1 were not pre-incubated but directly mixed together upon initiation of the digestion reaction. Makona GP was also digested in the presence and absence of NPC1 with 0.75x and 2x CatL (pH 4.5). An aliquot of the reaction mixture was withdrawn at defined time points, ranging between 1 and 150 min, quickly diluted 1:20 in solution at 3.5 M urea to induce dissociation of GP and NPC1, further diluted 1:20 in an acetic buffer (pH 5) and analysed by MP at a final concentration of 8 nM GP trimer. MP measurements were conducted using a Refeyn TwoMP system (Refeyn Ltd) in a similar manner as previously described^62^, using protein Dynamin-1 as calibration standard^63^. MP movies were acquired in the regular field of view for 60 s, and processed using DiscoverMP (version 2024 R2.1, Refeyn Ltd) to generate a list of individual landing events. The average mass of the uncleaved trimer was measured at ∼300 kDa and the smallest cleaved form was measured at ∼110 kDa. Therefore, events yielding masses <80 kDa and >500 kDa, arising from small traces of impurities or aggregates, were discarded. To further remove outliers, an initial round of gaussian fit was applied to the mass histograms, and signals exceeding the average mass with ±3 σ were discarded, as they likely belong to small traces of (cleaved) aggregates or (cleaved) free monomers in solution. The selected MP signals were then used to calculate the average mass (arithmetic mean) and standard deviation, and as well subjected to a second and final round of gaussian fit, which enabled deriving peak centre, σ, and mean absolute deviation (L1 error). For the kinetic experiments in the presence of NPC1 (and the corresponding unbound series), the same analysis workflow was applied, with the difference that events yielding masses <105 kDa were all discarded, in order to remove any signal arising from NPC1. Kinetic parameters (Tables S1 and S2), namely plateau P (expressed in kDa) and cleavage rate k (expressed as 1/min) were determined in OriginPro (version 2026) by fitting an exponential function to the calculated average masses over the time (x, in min), as per following equation: (296-P)*exp(-k*x)+P, where 296 corresponds to the mass in kDa of uncleaved GP.

### Interaction MP experiments

The interaction of GP-NPC1, of predicted affinity in the µM order, is hardly measurable with standard MP, due to its particle concentration limit of 50 nM^63^. Therefore, we employed nano-patterned passivated surfaces^42^ that enabled measuring samples at concentrations up to 1.33 µM. GP trimers (of different degree of cleavage) and NPC1 (C-domain) were incubated for 1 h in acetic buffer (pH 5) at ratio 1:3, and at concentrations in serial dilution. Samples were deposited on gaskets at final concentrations of GP trimers and NPC1 of 0.33 and 1 µM, 0.165 and 0.5 µM, 0.083 and 0.25 µM, 0.042 and 0.125 µM, respectively. Movies were promptly recorded in the regular field of view, in triplicates for each condition. The histograms of 60-s movies of these binding experiments were fitted with 5-Gaussian peak model with independent amplitudes. The first peak (representing NPC1) was fitted with independent mass and σ (σ_NPC1_). The second peak (representing cleaved GP) were fitted with independent mass. The peaks of the bound species were fitted with a mass equivalent to GP+nΔ, where n is number of bound NPC1 and Δ is the mass shift caused by NPC1 binding. The GP peaks, unbound and NPC1-bound, were fitted with a shared σ (σ_GP_). To calculate binding K_D_s, the molar concentrations of unbound GP and GP with 1–3 bound NPC1 were directly derived from their respective peak amplitudes, as previously described^64^. An elevated low-mass noise caused higher variability in NPC1 amplitude. Therefore, to achieve better accuracy, the concentration of free NPC1 was mathematically calculated by subtracting the bound NPC1 molar fraction from the known total molar concentration of NPC1 present in the sample. K_D_1 was defined as follows: [freeGP]*[freeNPC1]/[GP+1NPC1]. K_D_2 was defined as follows: [GP+1NPC1]*[freeNPC1]/[GP+2NPC1]. K_D_3 was defined as follows: [GP+2NPC1]*[freeNPC1]/[GP+3NPC1]. The K_D_ calculated at different GP:NPC1 concentrations were averaged to yield a final K_D_ value, and the standard error was calculated. The average of K_D_1/K_D_2 ratio and K_D_2/K_D_3 ratio, defined as average K_D_n/K_D_n+1 ratio, was used to infer the degree of binding cooperativity, as previously described^49^. In a trimer with three identical, independent binding sites, K_D_n+1 is equivalent to 3x K_D_n, because of statistical degeneracy^49^. Therefore, K_D_n/K_D_n+1>0.33 indicates non-equivalent binding sites, with positive cooperativity in NPC1 receptor binding.

## Data availability

According to the community-based recommendations^59^ and to allow access to the HDX data from this study, the HDX summary tables are included as Supplementary Tables S4–S10 and the HDX data tables are provided as source data within this paper. All MP histograms are included in the Supplementary Dataset within this paper.

## Acknowledgments

The authors thank Dr. Francesca Donnellan and Dr. Pramila Rijal for the generous donation of mAb REGN3470. The authors also thank Dr. Antoni G. Wrobel and Tia Hawkins for helpful discussions and comments on the manuscript draft.

## Author contribution

V.C. and W.B.S. conceptualized the study and supervised the work. V.C. expressed and purified proteins. V.C. conceptualized, performed, and analysed all HDX-MS and MS experiments and interpretated the results. V.C., J.K. and W.B.S. conceptualized MP experiments. V.C. performed all kinetic MP experiments. V.C. and J.K. analysed kinetic MP experiments and V.C. interpreted the data. V.C. and J.K. performed, analysed and interpreted interaction MP experiments. V.C. and A.B. performed and analysed the HDX-MS epitope mapping with REGN3470, and performed SDS-page of GP cleavage in the presence and absence of REGN3470. V.C., P.K. and W.B.S. provided reagents and/or instrumentations. V.C., J.K. and W.B.S. wrote the original draft and all authors reviewed and edited the final draft.

## Supplementary Information

**Fig. S1.**
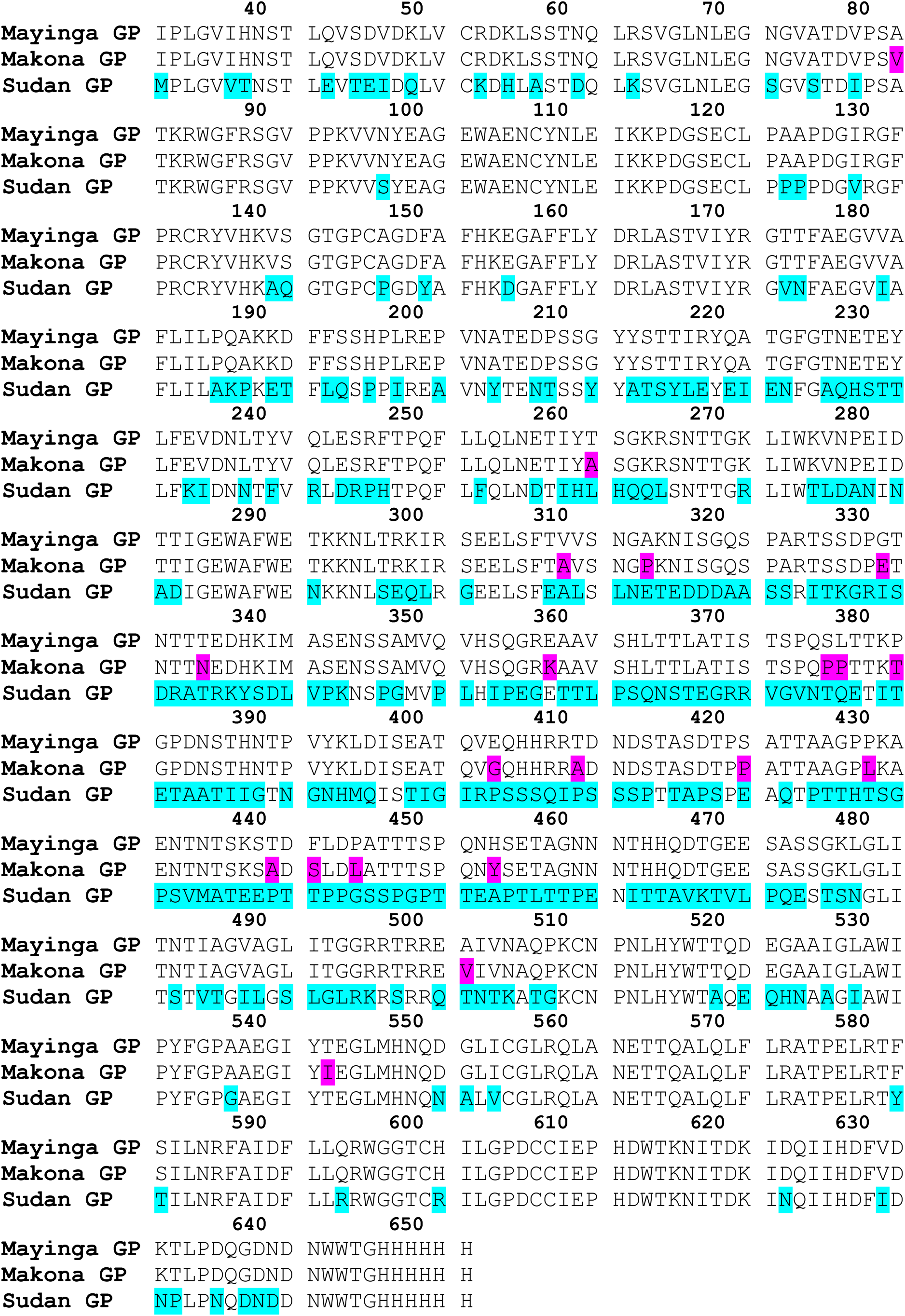
Sequence alignment of Mayinga, Makona and Sudan GP. Residues that are different in Makona compared to Mayinga GP and in Sudan compared to Mayinga GP are highlighted.

**Fig. S2.**
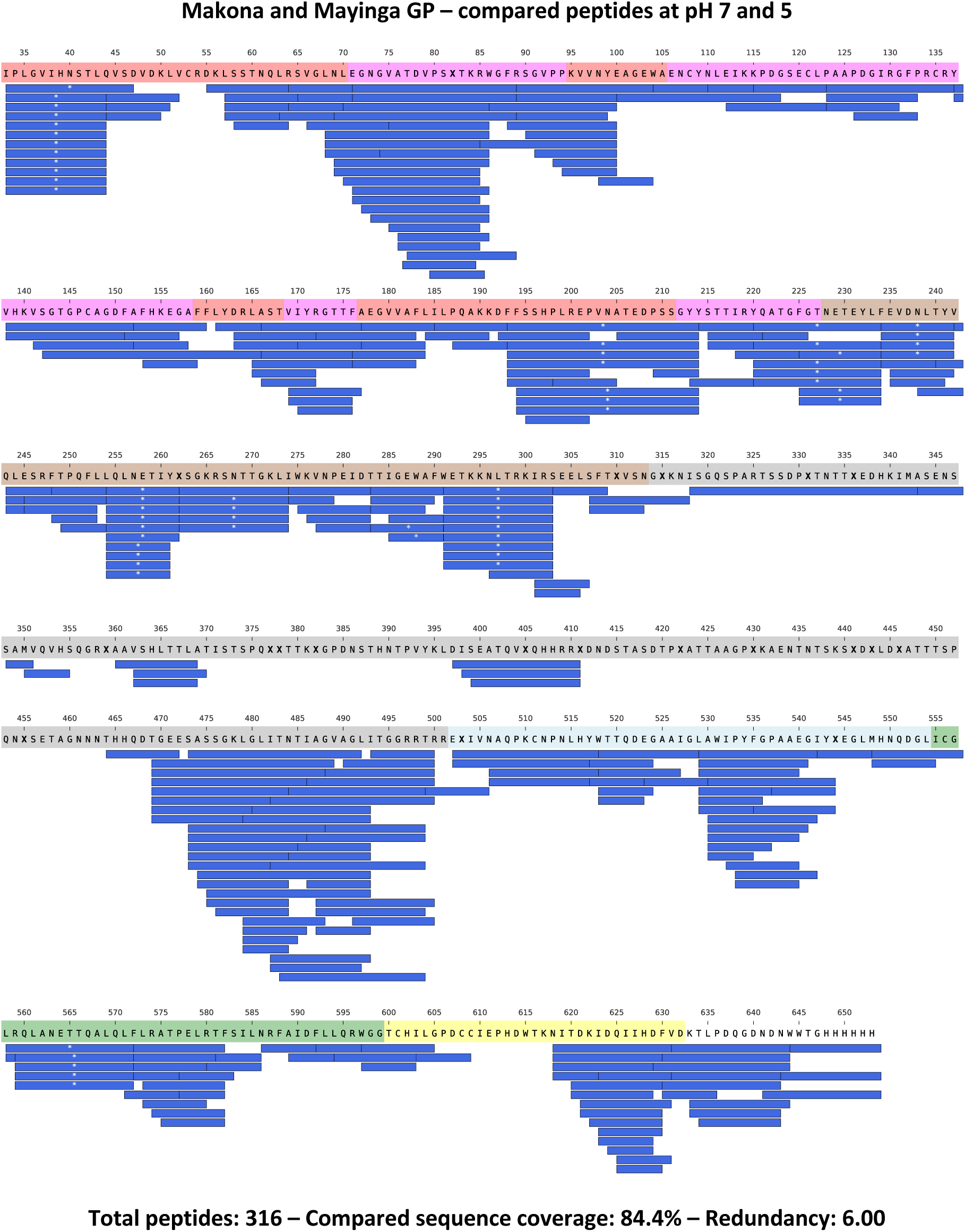
Peptides whose HDX was compared between Makona and Mayinga GP at pH 7 and 5 are illustrated along the protein sequence with blue bars. Blue bars with white asterisks (*) indicate glycosylated peptides. Residues marked as X are different between Makona and Mayinga GP. The individidual domains of GP are color-coded along the protein sequence as follow: red: Base; purple: Head domain; brown: Glycan Cap (GC); gray: Mucin-Like Domain (MLD); light blue: Internal Fusion Loop (IFL); green: Heptad Repeat 1 (HR1); yellow: Heptad Repeat 2 (HR2).

**Fig. S3.**
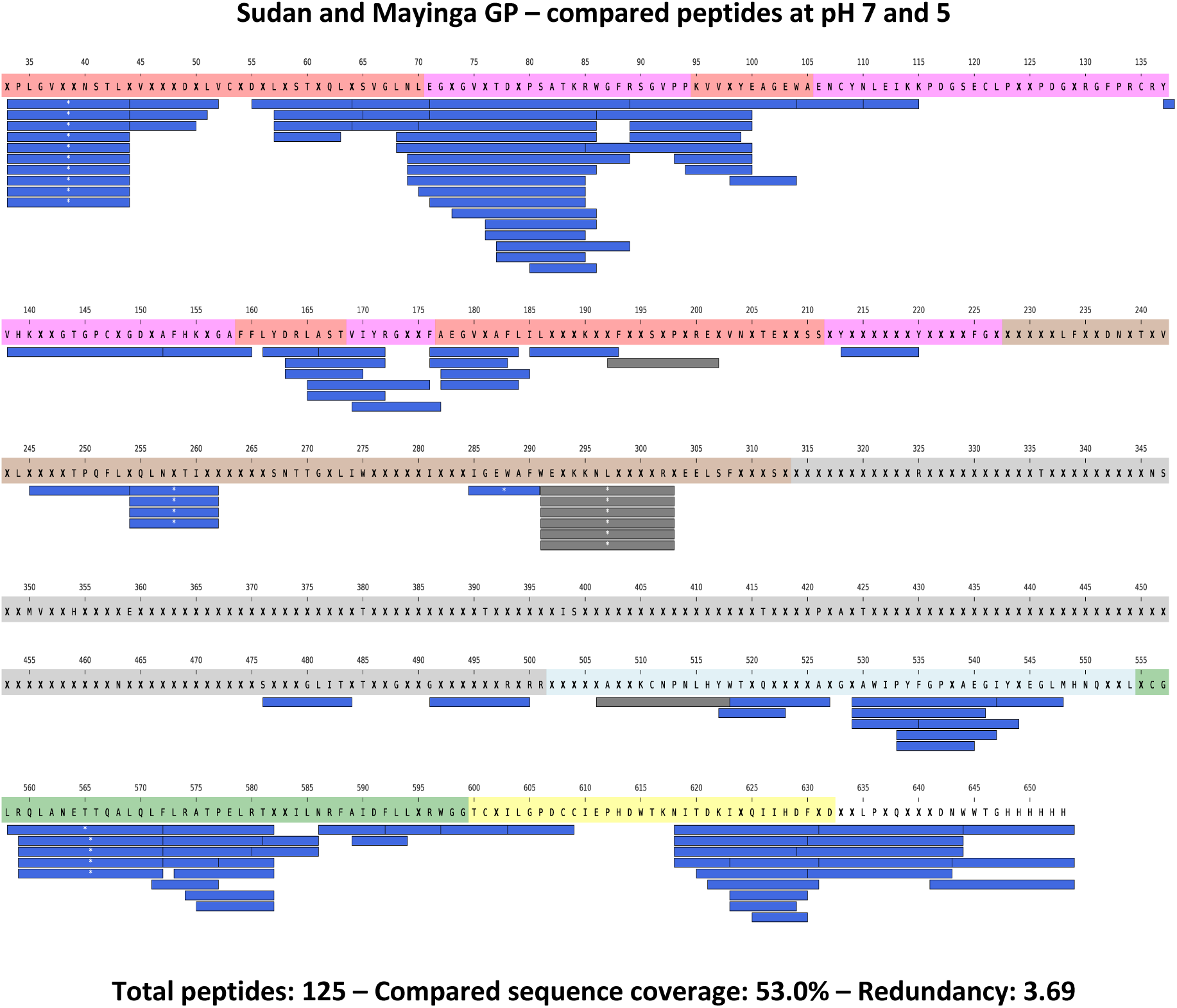
Peptides whose HDX was compared between Sudan and Mayinga GP at pH 7 and 5 are illustrated along the protein sequence with blue bars. Blue bars with white asterisks (*) indicate glycosylated peptides. Gray bars indicate peptides that have been compared by their deuterium uptake normalized by MaxD as they differ in their termini between Mayinga and Sudan GP for one residue. Residues marked as X are different between Sudan and Mayinga GP. The individidual domains of GP are color-coded along the protein sequence as in Fig. S1.

**Fig. S4.**
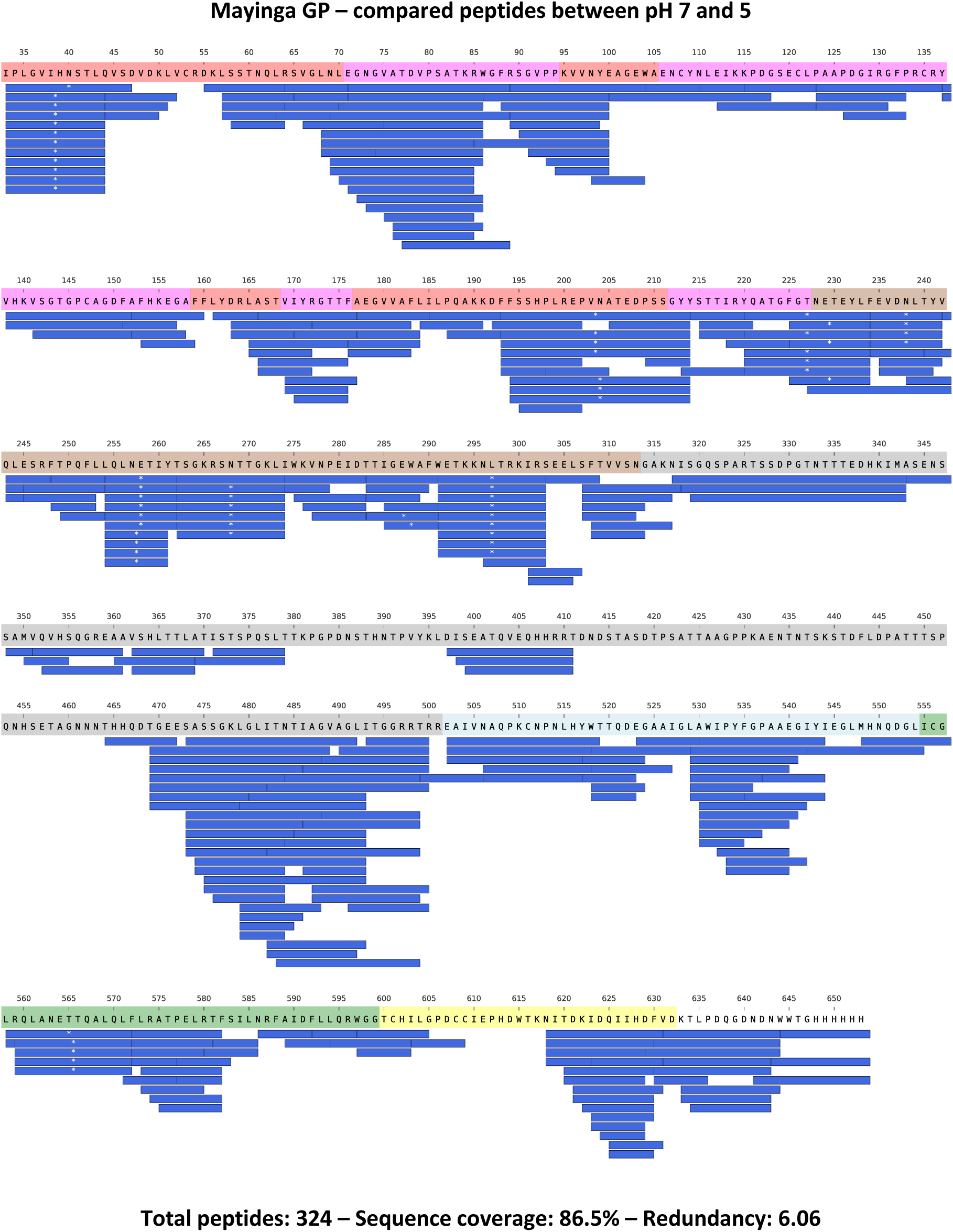
Peptides whose HDX was followed for Mayinga GP at pH 7 and 5 are illustrated along the protein sequence with blue bars. Blue bars with white asterisks (*) indicate glycosylated peptides. The individidual domains of GP are color-coded along the protein sequence as in Fig. S1.

**Fig. S5.**
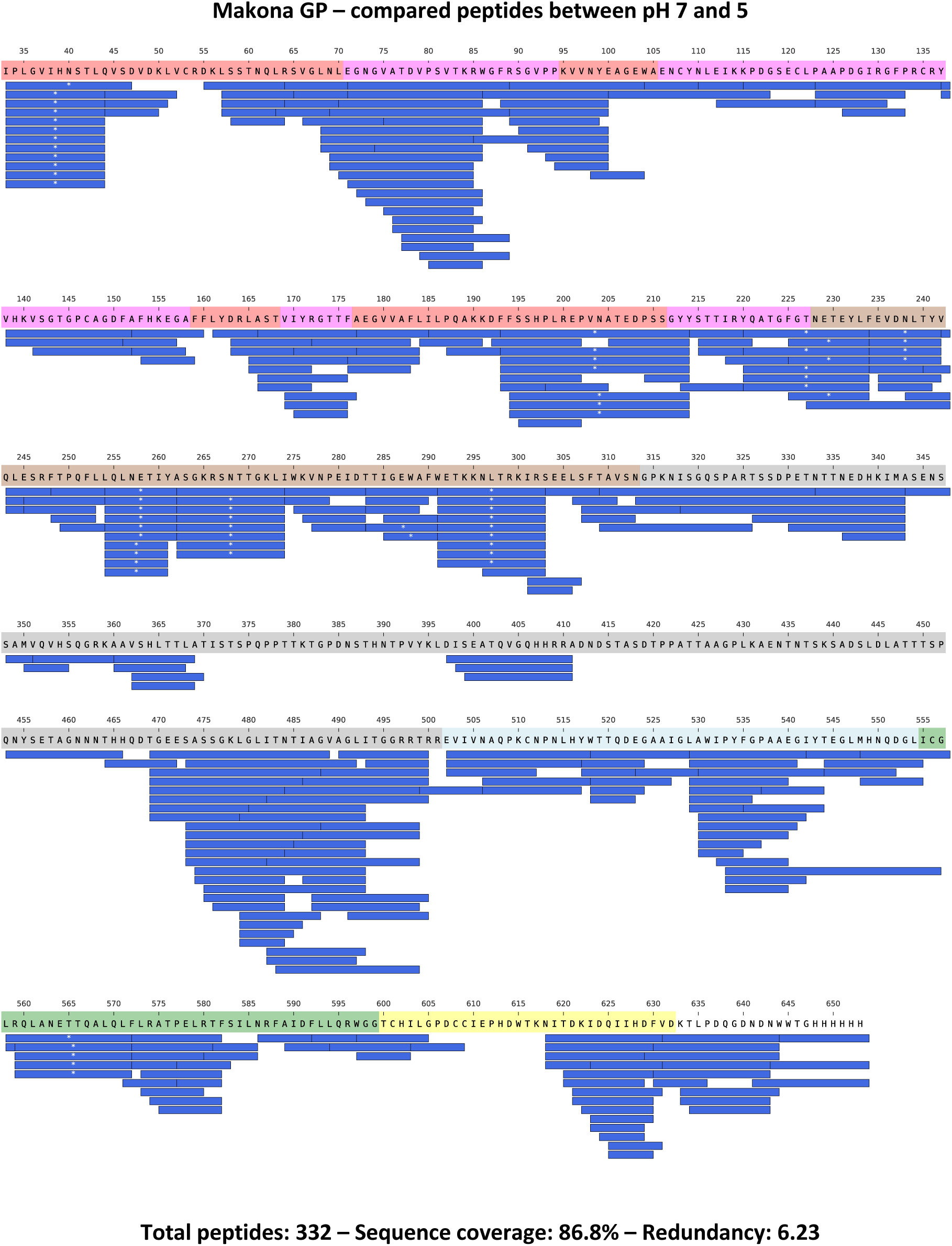
Peptides whose HDX was followed for Makona GP at pH 7 and 5 are illustrated along the protein sequence with blue bars. Blue bars with white asterisks (*) indicate glycosylated peptides. The individidual domains of GP are color-coded along the protein sequence as in Fig. S1.

**Fig. S6.**
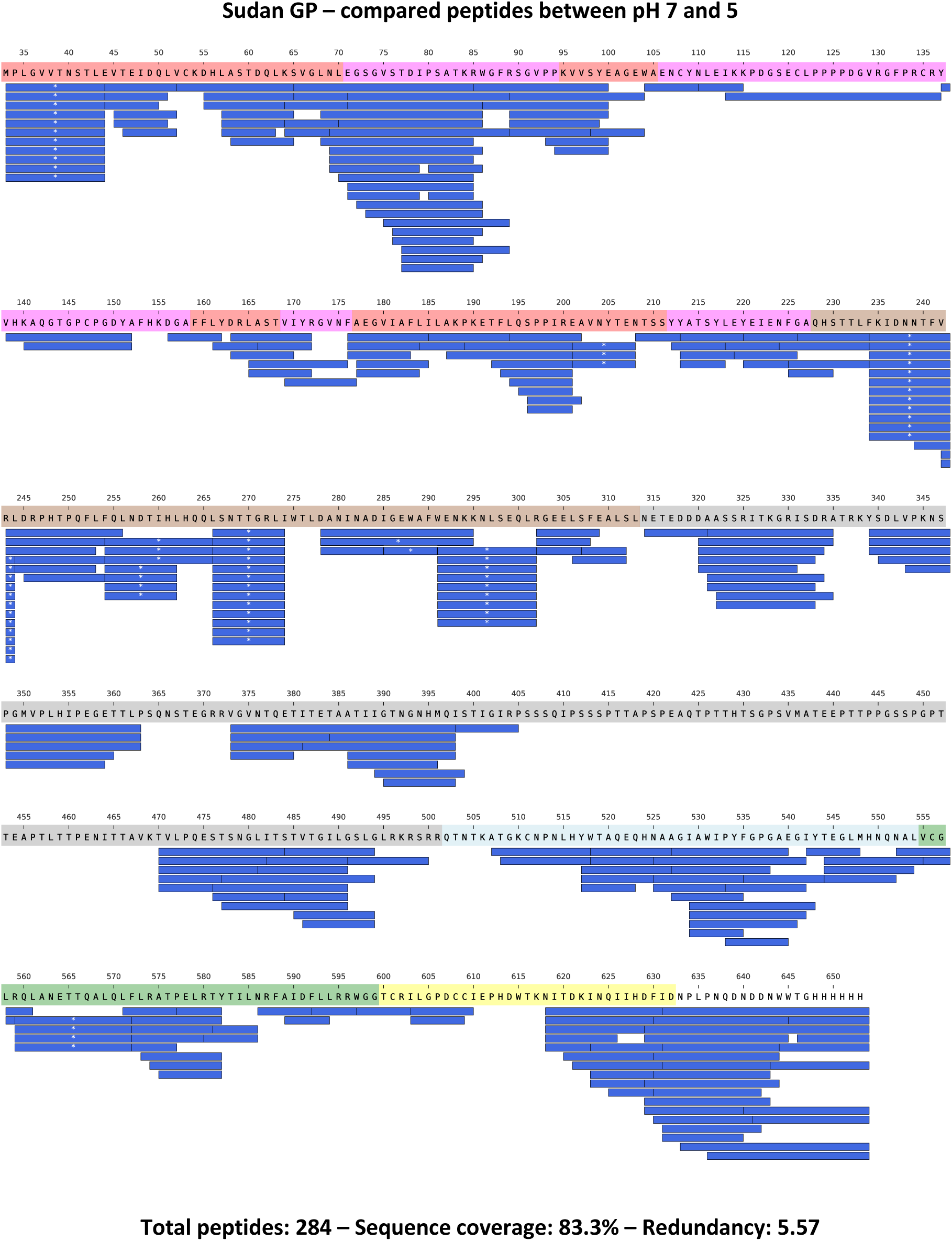
Peptides whose HDX was followed for Sudan GP at pH 7 and 5 are illustrated along the protein sequence with blue bars. Blue bars with white asterisks (*) indicate glycosylated peptides. The individidual domains of GP are color-coded along the protein sequence as in Fig. S1.

**Fig. S7.**
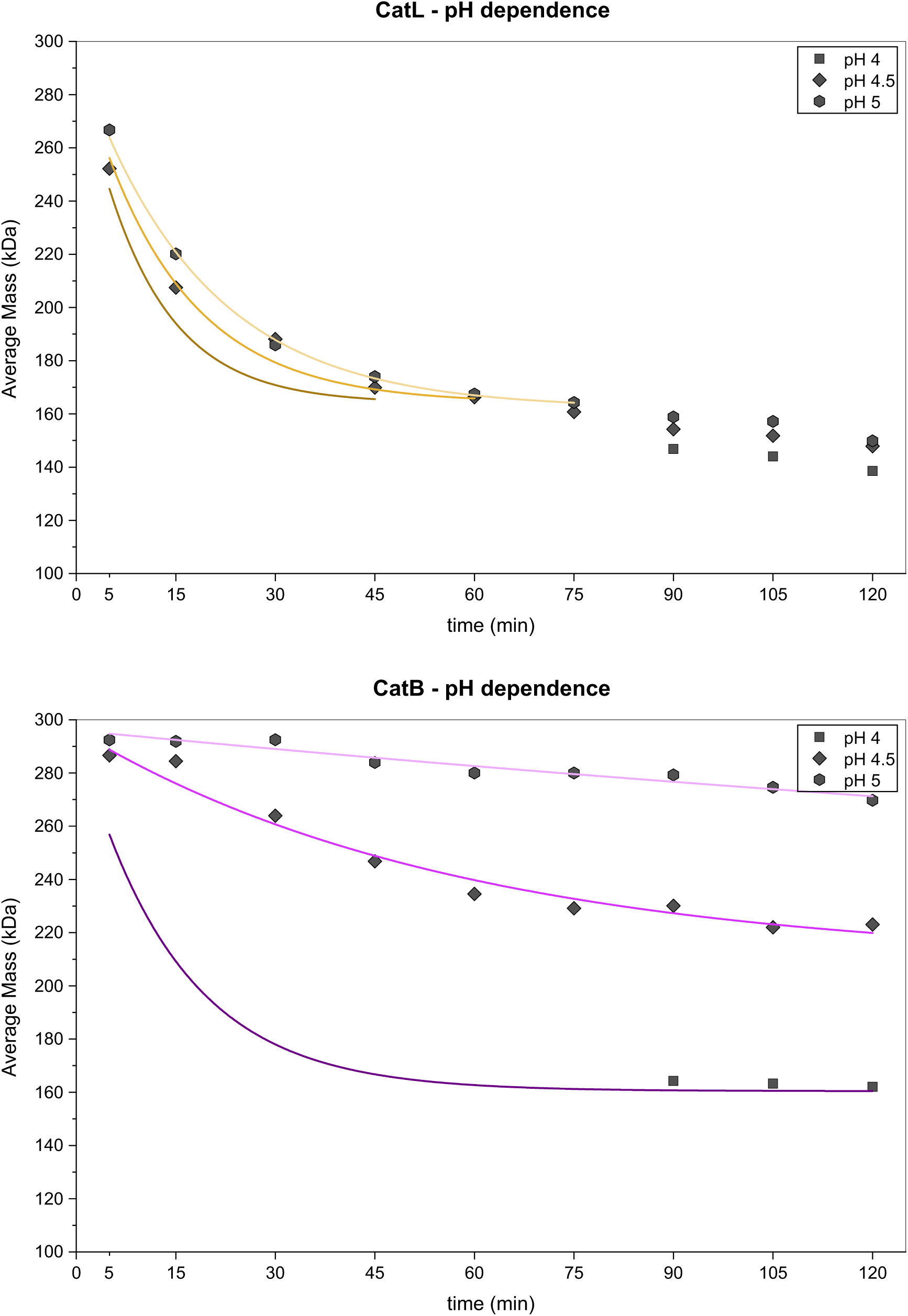
Dependence of CatL **(top)** and CatB **(bottom)** activity on pH measured by MP upon cleavage of Makona GP. Concentrations of enzymes: CatL 2x and CatB 1x. Fitted curves are coloured as follow: CatL, pH 5: yellow, CatL; pH 4.5: orange; CatL, pH 4: brown; CatB, pH 5: pink; CatB, pH 4.5: magenta; CatB, pH 5: purple. Kinetic parameters in Table S1. Source data are included in source data files.

**Fig. S8.**
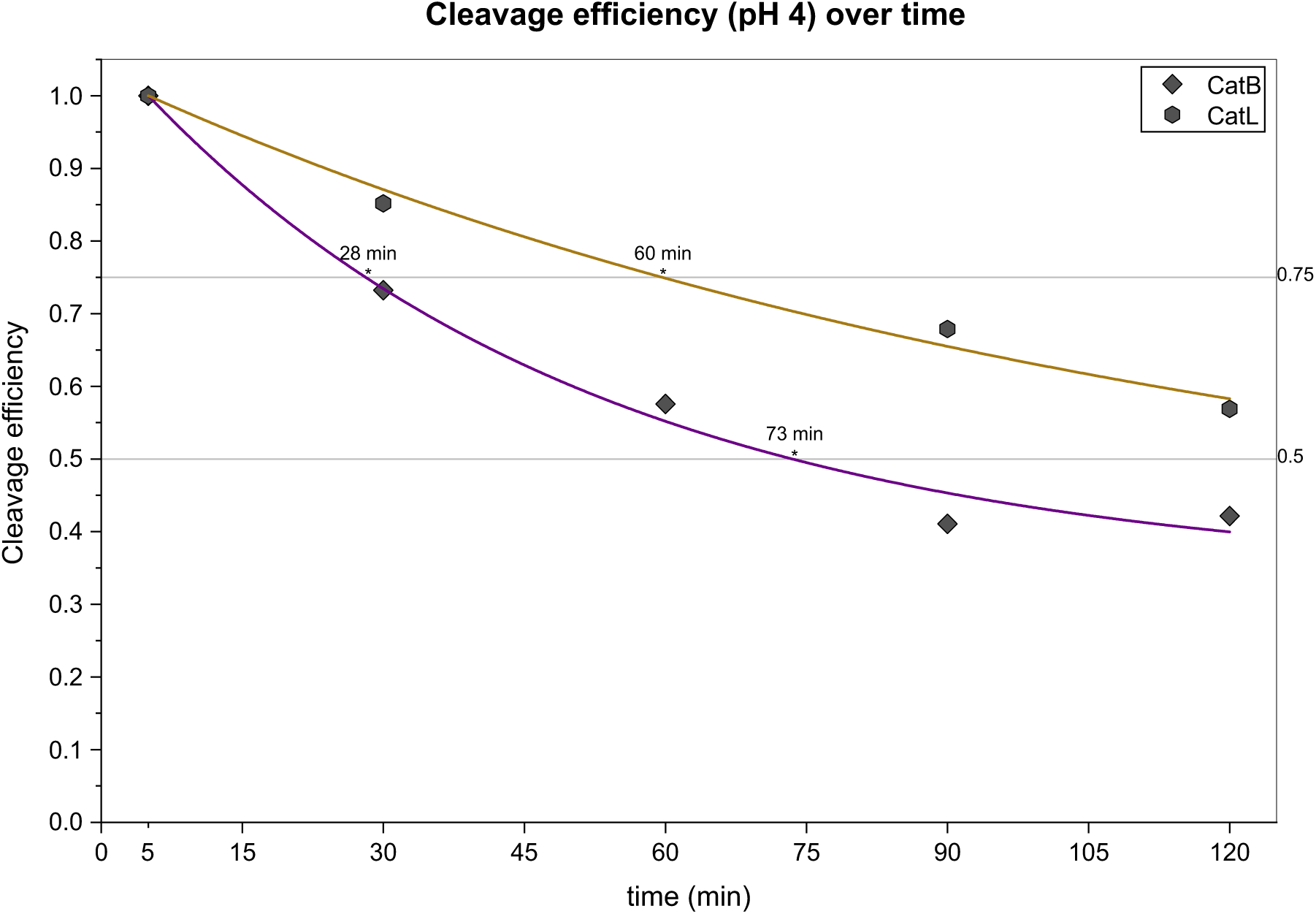
Cleavage efficiency at pH 4 of CatB and CatL over time measured by MP. Makona GP was digested for five minutes after the enzymes incubated alone for different time periods (from 5 to 120 min). The ratio between the mass of GP digested with the enzyme incubated for x min and 5 min is plotted as cleavage efficiency. Fitted curves are coloured as follow: CatL: yellow; CatB: purple. Enzyme activity remains significant (>55%) up to 60 and 120 min for CatB and CatL, respectively. Kinetic parameters in Table S1. Source data are included in source data files.

**Fig. S9.**
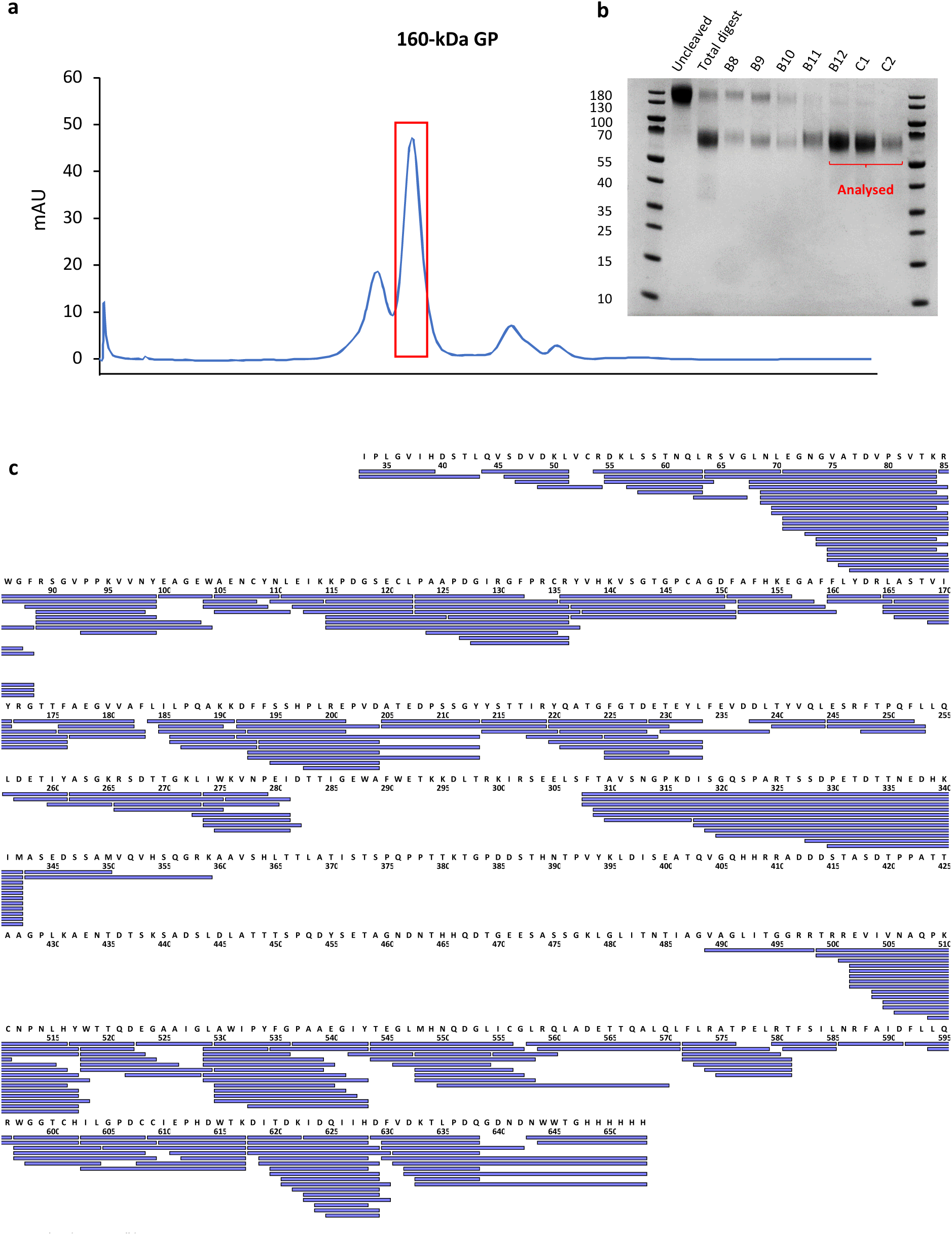
Peptide identification of 160-kDa product of Makona GP. **a)** SEC trace of the digest containing the 160-kDa product. **b)** Non-reducing SDS-page showing the signal of uncleaved GP, the non-purified digest and various SEC fractions, including those pooled and subjected to bottom-up MS to analysis. **c)** Sequence coverage of the SEC-purified and de-glycosylated 160-kDa product contained in the pool of selected fractions.

**Fig. S10.**
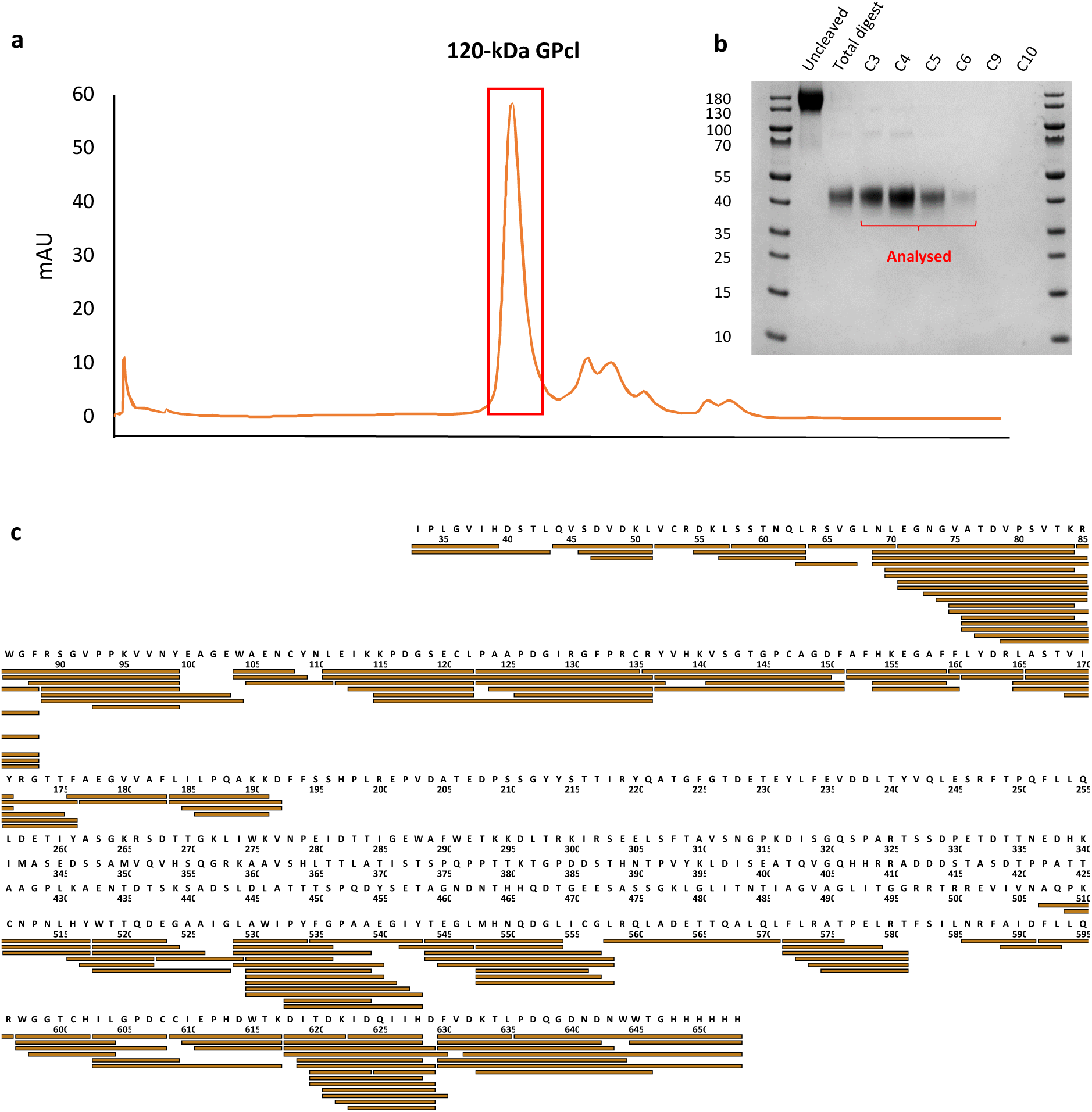
Peptide identification of 120-kDa product of Makona GP. **a)** SEC trace of the digest containing the 120-kDa product. **b)** Non-reducing SDS-page showing the signal of uncleaved GP, the non-purified digest, and various SEC fractions, including those pooled and subjected to bottom-up MS to analysis. **c)** Sequence coverage of the SEC-purified and de-glycosylated 120-kDa product contained in the pool of selected fractions.

**Fig. S11.**
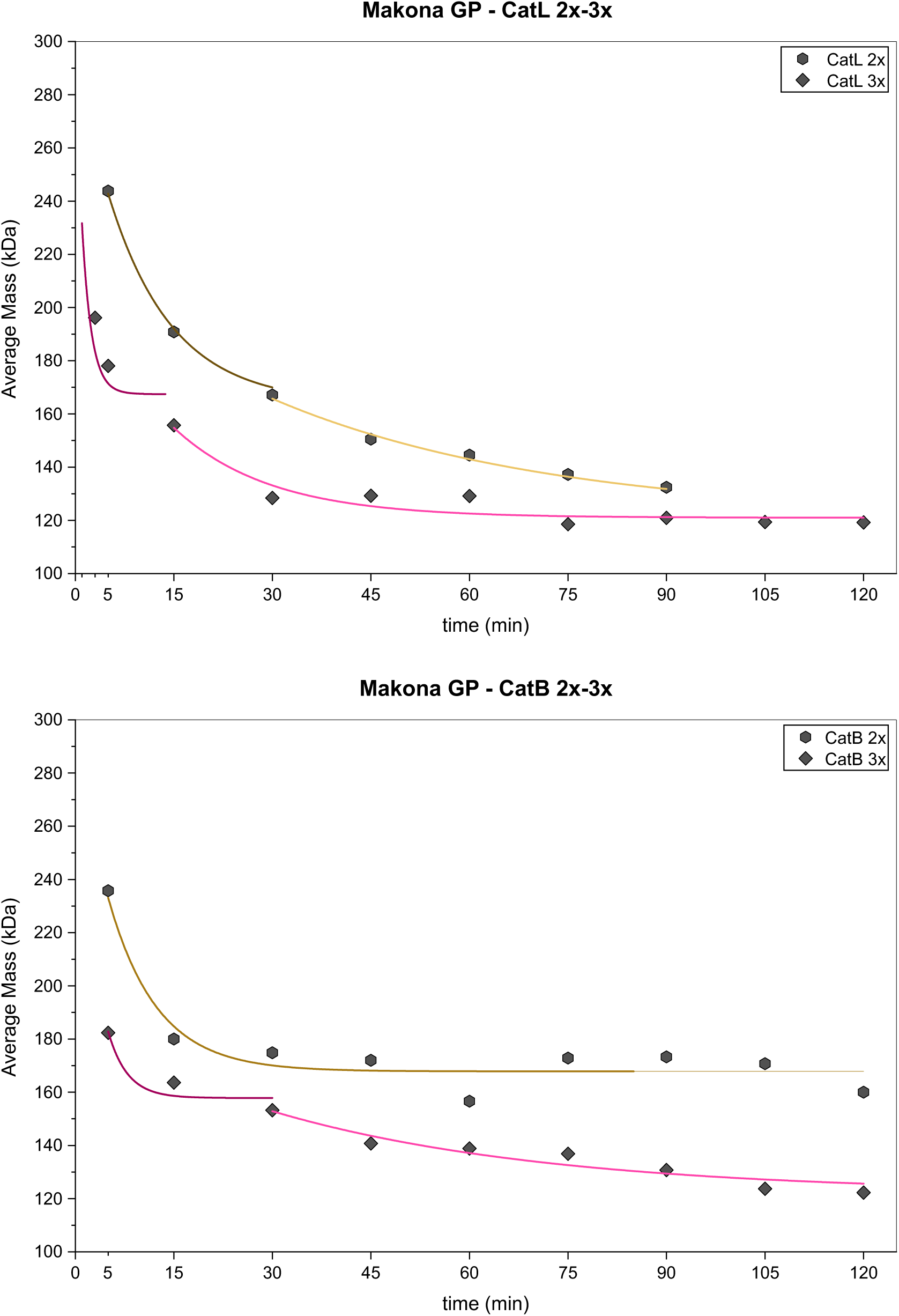
Cleavage of Makona GP with CatL **(top)** and CatB **(bottom)** at concentrations 2x and 3x measured by MP. Fitted curves are coloured as follow: CatL 2x to 164 kDa: brown; CatL 2x 164-120 kDa: yellow; CatL 3x to 164 kDa: purple; CatL 3x 164-120 kDa: magenta; CatB 2x to 160 kDa: brown; CatB 3x to 160 kDa: purple; CatB 3x 160-120 kDa: magenta. Kinetic parameters in Table S1. Source data are included in source data files.

**Fig. S12.**
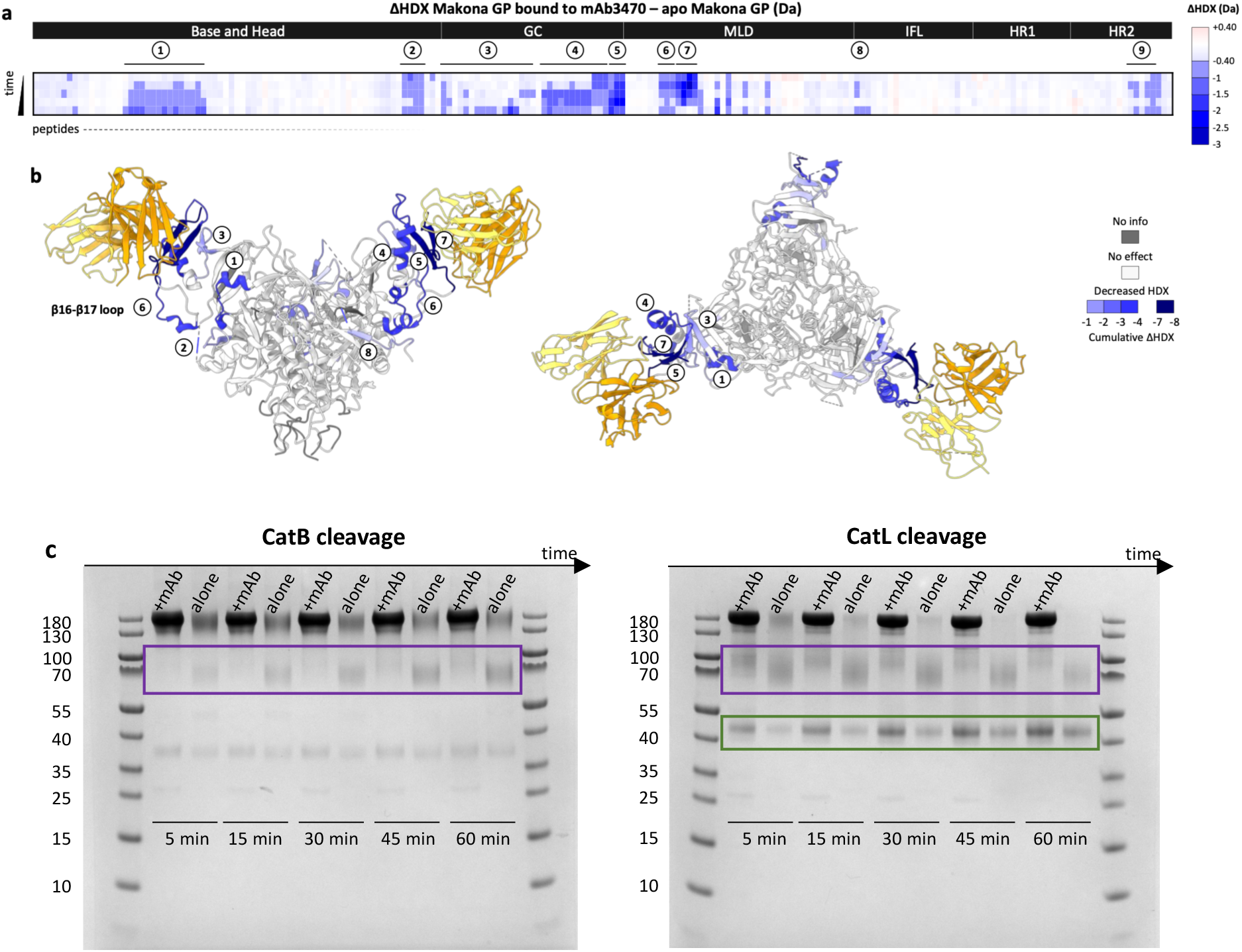
Effect of mAb3470 on the cleavage of Makona GP with CatB and CatL. **a)** Plot illustrating ΔHDX profile of bound GP – apo GP indicates several areas of decreased HDX upon binding. **b)** ΔHDX effects are superimposed on the structure of GP bound to mAb3470 (PDB: 7TN9). Segments spanning residues 274-282 and 303-312 (areas 5 and 7, respectively) show the greatest decrease in HDX and align with the mAb epitope. Segment 291-302 (area 6) and 71-85 (RBR helix, area 1) also show a significant decrease in HDX (rigidification) upon binding. Areas 5, 6 and 7 span the GC-MLD loop cleaved by CatB; area 6 and 7 span the GC-MLD loop cleaved by CatL. **c)** Cleavage of Makona GP over time (5, 15, 30, 45 and 60 min) with CatB and CatL in the presence and absence of mAb monitored by SDS-page. With CatB, no band corresponding to cleaved forms was detected in the presence of mAb. With CatL, two bands corresponding to cleaved forms were detected both in the presence and absence of mAb. Bands framed in purple: the band in the presence of mAb is at higher mass and fainter compared to the band in the absence of mAb, likely because the cleavage of the GC-MLD loop is skipped in the presence of mAb, yielding a longer protein segment. Band framed in green: it corresponds to 120-kDa GPcl and is darker in the presence of mAb, likely because the final product formation is facilitated by the GC-head domain loop becoming the favourite cleavage site, resulting in a faster liberation of 22-kDa GC (as cleavage of the GC-MLD loop is in fact unnecessary for removing the GC). Source data are included in source data files.

**Fig. S13.**
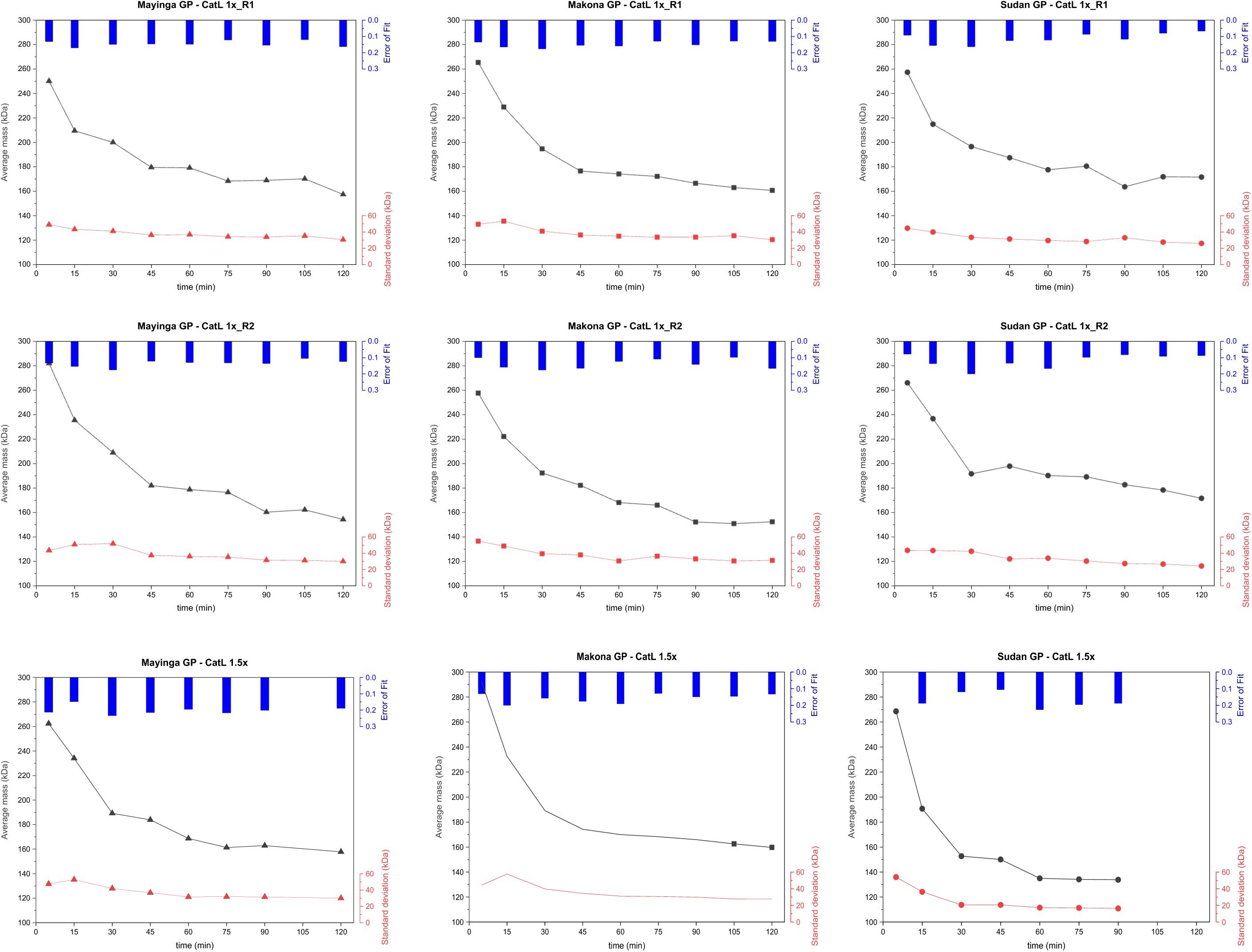
Extended charts of kinetic experiments of GP cleavage with 1x CatL and 1.5x CatL. For each MP signal across the time points studied, the graphs show average mass, standard deviation from the average mass, and L1 error (error of fit); the higher the L1 error, the higher deviation from gaussian fit, indicating the presence of multiple populations. Source data are included in source data files.

**Fig. S14.**
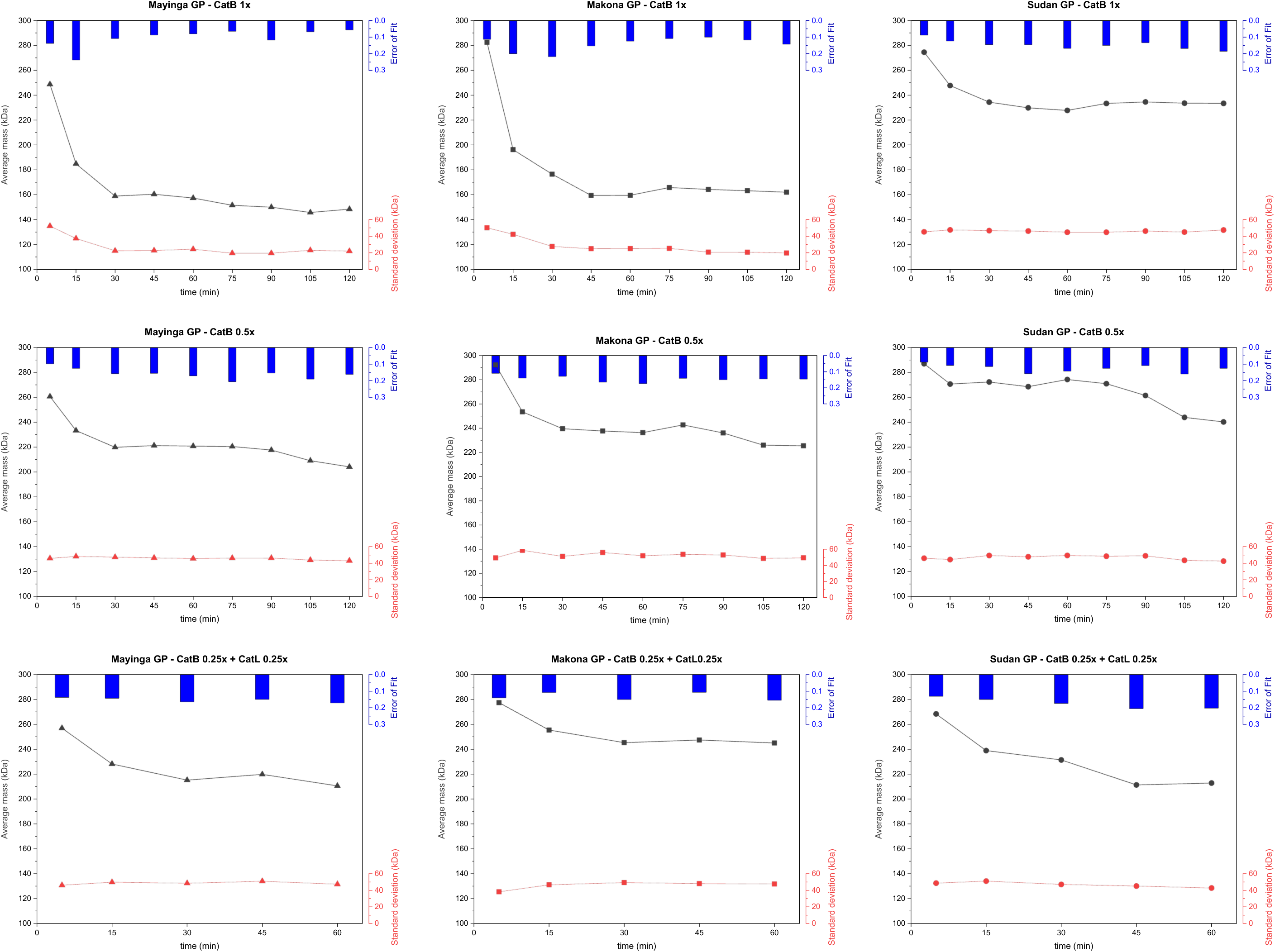
Extended charts of kinetic experiments of GP cleavage with 0.5x CatB, 1x CatB, and 0.25x CatB + 0.25x CatL. For each MP signal across the time points studied, the graphs show average mass, standard deviation from the average mass, and L1 error (error of fit); the higher the L1 error, the higher deviation from gaussian fit, indicating the presence of multiple populations. Source data are included in source data files.

**Fig. S15.**
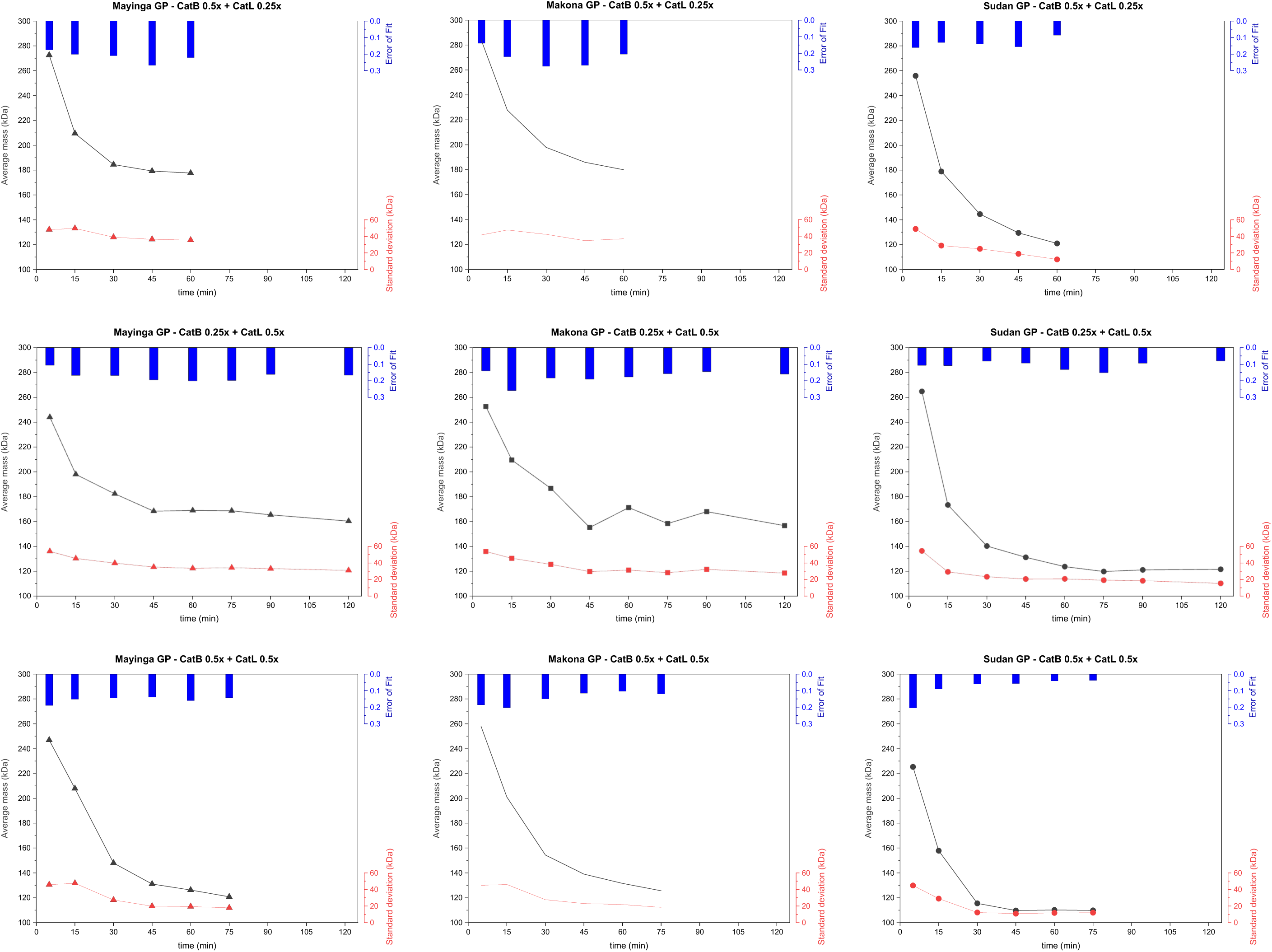
Extended charts of kinetic experiments of GP cleavage with 0.5x CatB + 0.25x CatL, 0.25x CatB + 0.5x CatL, and 0.5x CatB + 0.5x CatL. For each MP signal across the time points studied, the graphs show average mass, standard deviation from the average mass, and L1 error (error of fit); the higher the L1 error, the higher deviation from gaussian fit, indicating the presence of multiple populations. Source data are included in source data files.

**Fig. S16.**
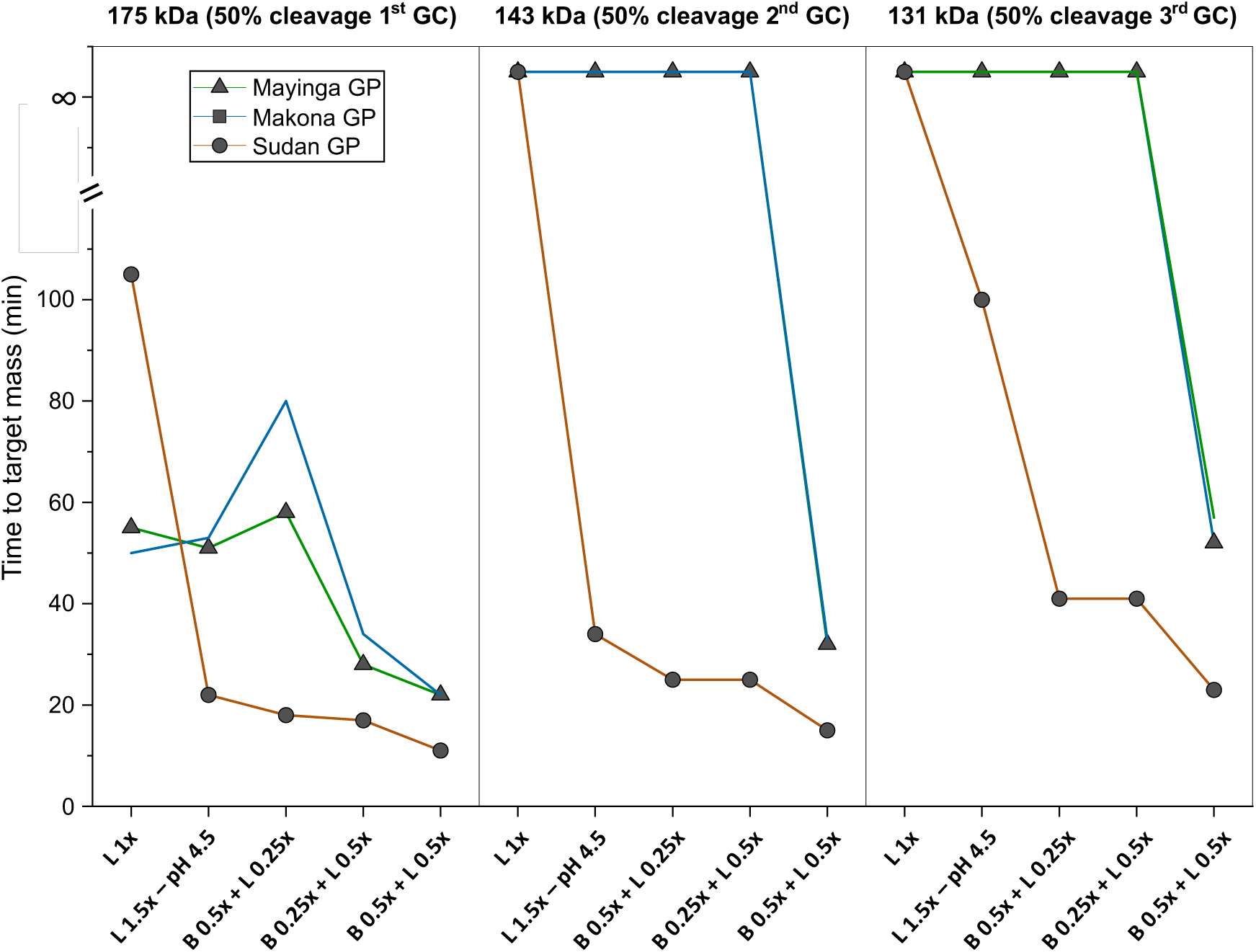
Plot illustrating the time required to cleave out each individual GC from Mayinga, Makona and Sudan GP under the various cathepsin digestion conditions tested (B=CatB, L=CatL).

**Fig. S17.**
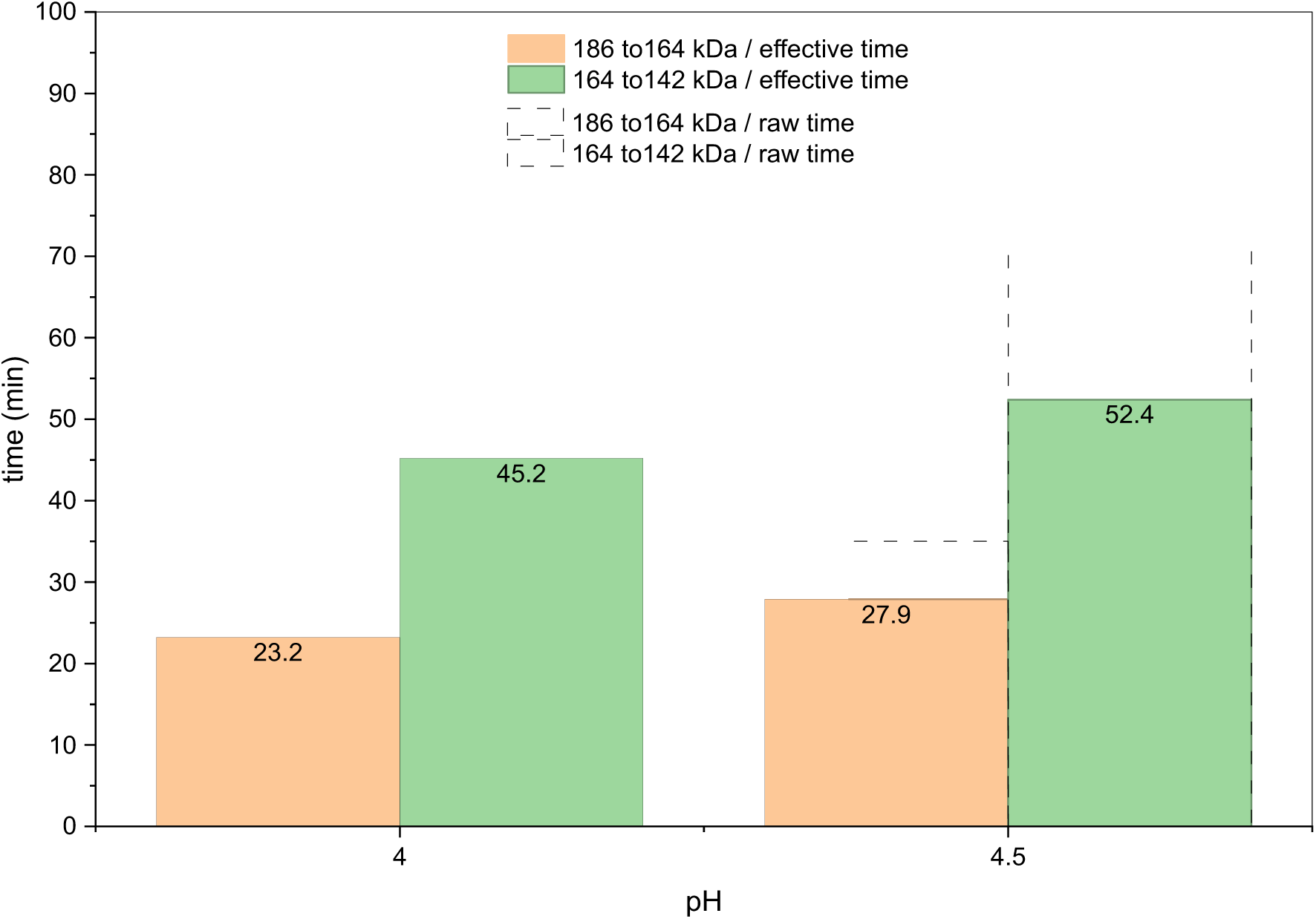
Plot illustrating the time required to cleave out each individual 22-kDa GC in Makona GP at pH 4 and at pH 4.5 with 2x CatL (data derived from plot in Fig. S7). The labels indicate the time (in min) for the 1^st^ and 2^nd^ 22-kDa GC removal. The data show that the 2^nd^ 22-kDa GC removal (transition from 164 to 142 kDa) requires approximately double of time than the 1^st^ 22-kDa GC removal (transition from 186 to 164 kDa). The effective times (solid columns) have been derived by correcting the raw times (dashed transparent columns) by the loss of CatL activity over time (plot in Fig. S8). As digestion with 2x CatL did not yield full removal of the 3^rd^ 22-kDa GC, we cannot exactly calculate the time required for its completion, but the data suggest that this takes even longer than the 2^nd^ removal.

**Fig. S18.**
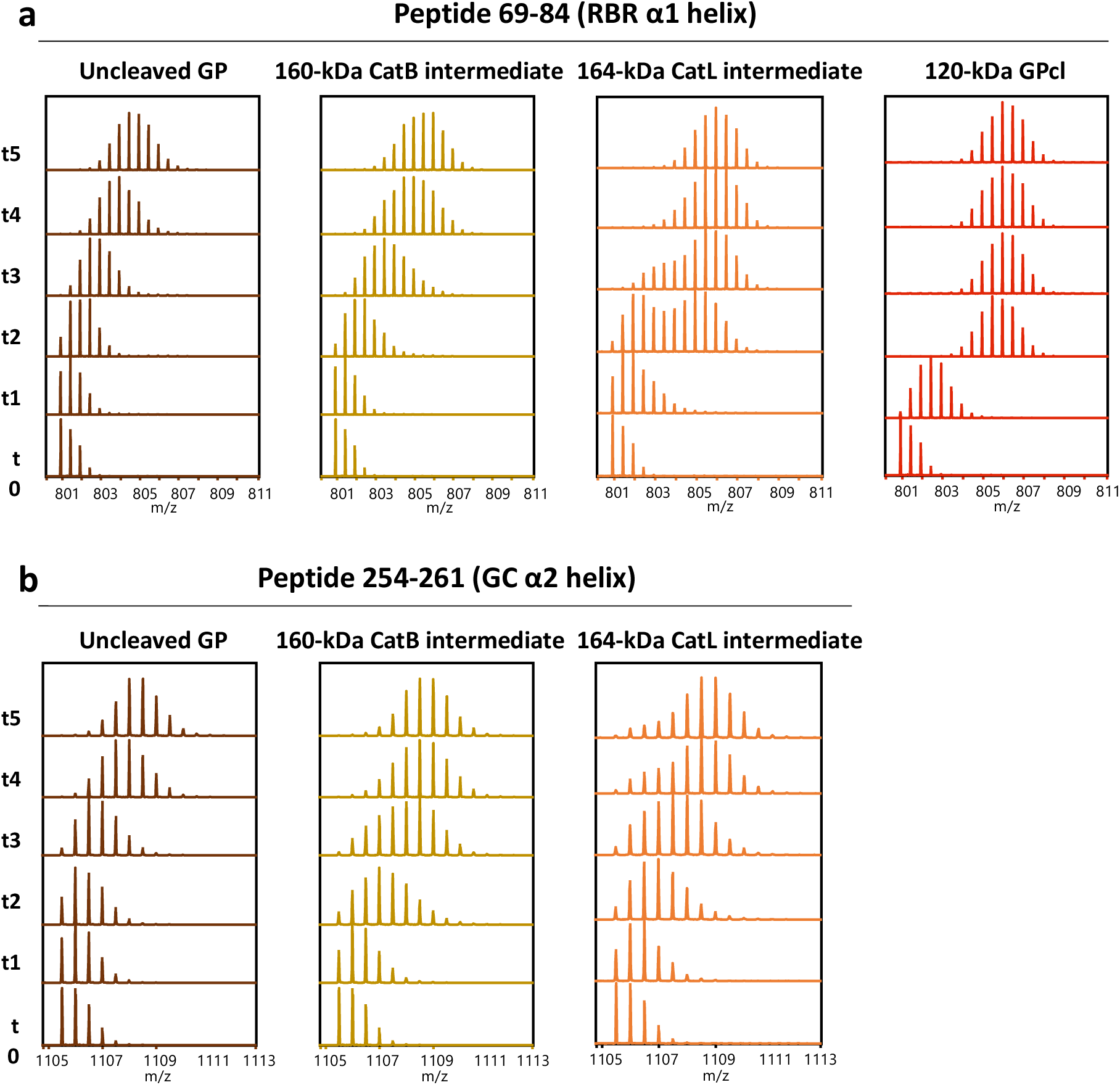
Spectra of model peptides spanning the RBR α1 helix **(a)** and the GC α2 helix **(b)** at the time points studied (t1 = 400 s on ice; t2 = 400 s at 23 °C; t3 = 1 h at 23 °C; t4 = 10 h at 23 °C; t5 = 20 h at 28 °C) and under the four states investigated: Uncleaved Makona GP, 160-kDa CatB intermediate, 164-kDa CatL intermediate and 120-kDa GPcl. Peptide 254-261 is absent for the state 120-kDa GPcl, as the α2 helix is completely digested. The spectra indicate a shift toward higher average masses for the RBR helix, scaling with Uncleaved < 160-kDa < 164-kDa < 120-kDa GP. **a)** The spectra indicate a shift toward higher average masses for the α2 helix, scaling with Uncleaved < 164-kDa < 160-kDa GP. **b)** Bimodal HDX spectra are visible for the peptide spanning the RBR helix under the state 164-kDa GP; and for the peptide spanning the α2 helix under the state 160-kDa and 164-kDa GP, the latter being of lower average masses.

**Fig. S19.**
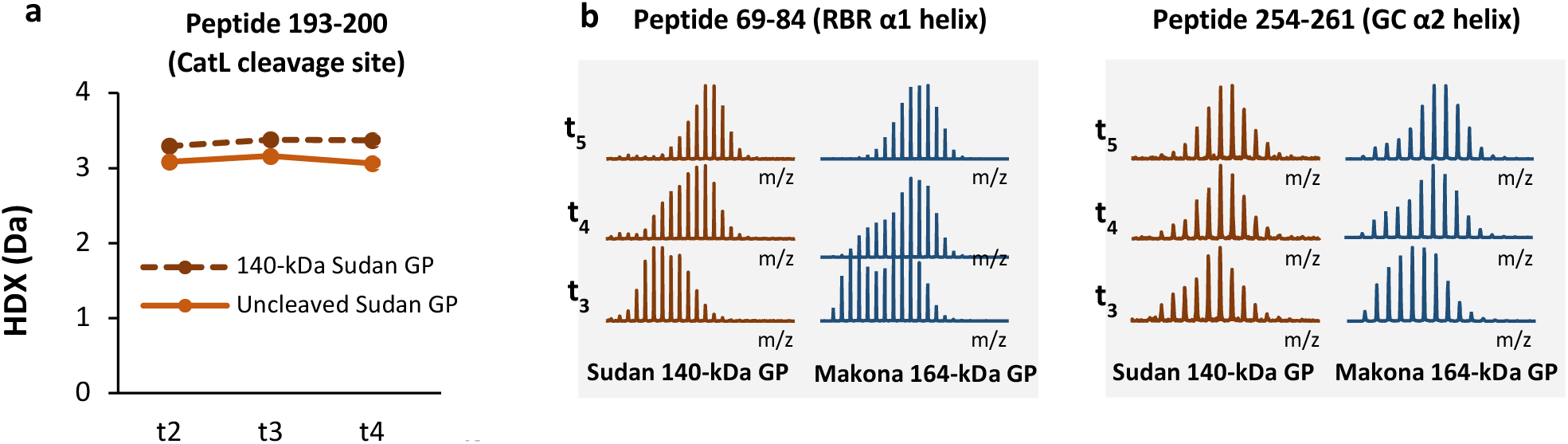
**a)** Deuterium uptake plot of peptide 193-200 (spanning the CatL cleavage site) illustrates that this segment is more dynamic in 140-kDa Sudan GP than in the uncleaved Sudan GP. **b)** Spectra of peptides 69-84 (spanning the RBR helix) and 254-261 (spanning the α2 helix) in Sudan 140-kDa GP and Makona 164-kDa GP illustrate differences in structural dynamics of the RBR helix–α2 helix–CatL cleavage site axis between the two trimers. Sudan 140-kDa GP, despite being more cleaved that Makona 164-kDa GP, displays a more closed RBR helix than Makona 164-kDa GP, exemplified by the lower average mass of both populations. Accordingly, Sudan 140-kDa GP displays a more open α2 helix than Makona 164-kDa GP, exemplified by the lower abundance of the low-mass population compared to Makona 164-kDa GP. Time points: t2 = 400 s at 23 °C; t3 = 1 h at 23 °C, t4 = 10 h at 23 °C.

**Fig. S20.**
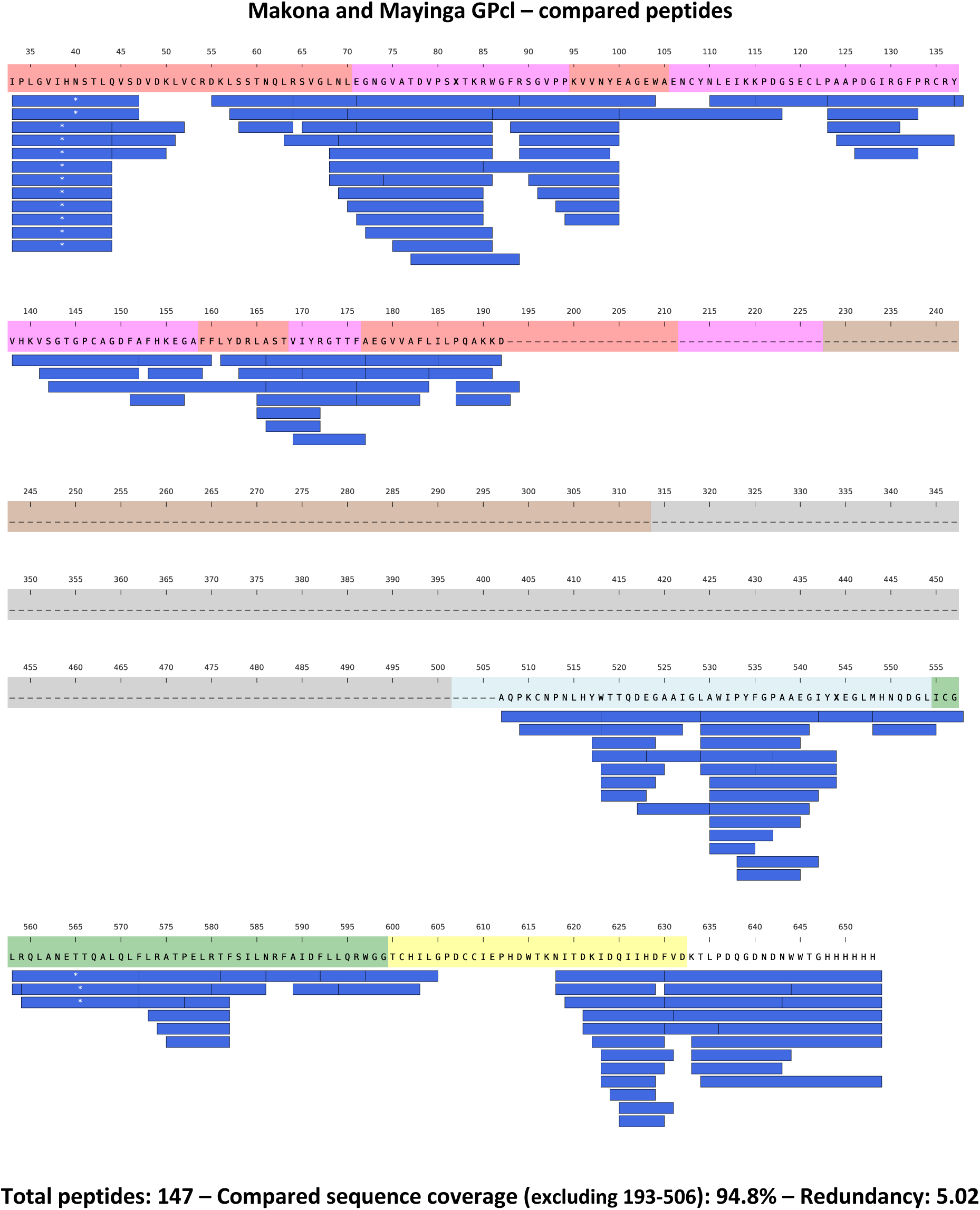
Peptides whose HDX was compared between Makona and Mayinga GPcl at pH 5 are illustrated along the protein sequence with blue bars. Blue bars with white asterisks (*) indicate glycosylated peptides. Residues marked as X are different between Makona and Mayinga GP. The dashed line indicates the cleaved portion of GP, whose peptides are absent in the dataset. The individidual domains of GP are color-coded along the protein sequence as in Fig. S1.

**Fig. S21.**
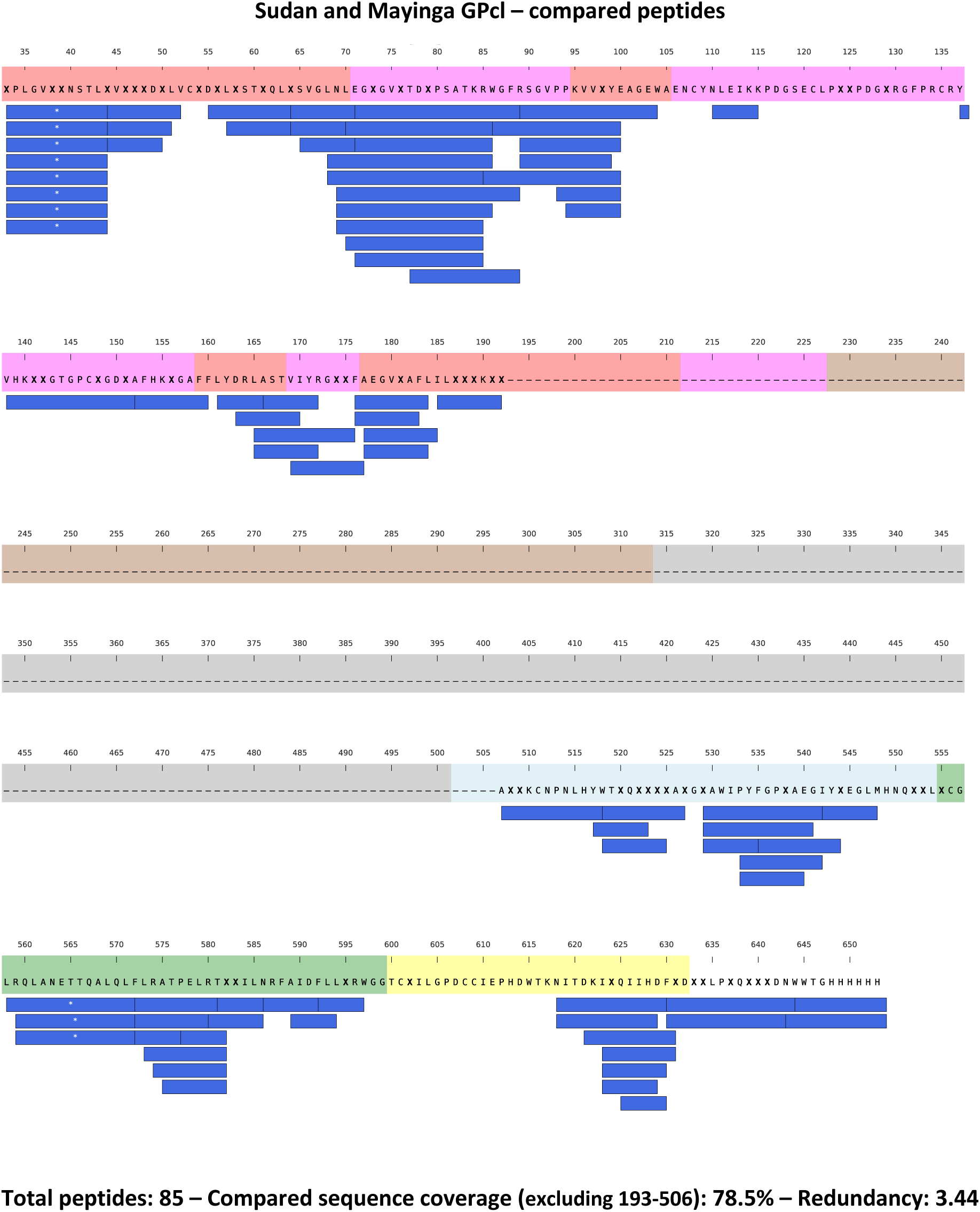
Peptides whose HDX was compared between Sudan and Mayinga GPcl at pH 5 are illustrated along the protein sequence with blue bars. Blue bars with white asterisks (*) indicate glycosylated peptides. Residues marked as X are different between Sudan and Mayinga GP. The dashed line indicates the cleaved portion of GP, whose peptides are absent in the dataset. The individidual domains of GP are color-coded along the protein sequence as in Fig. S1.

**Fig. S22.**
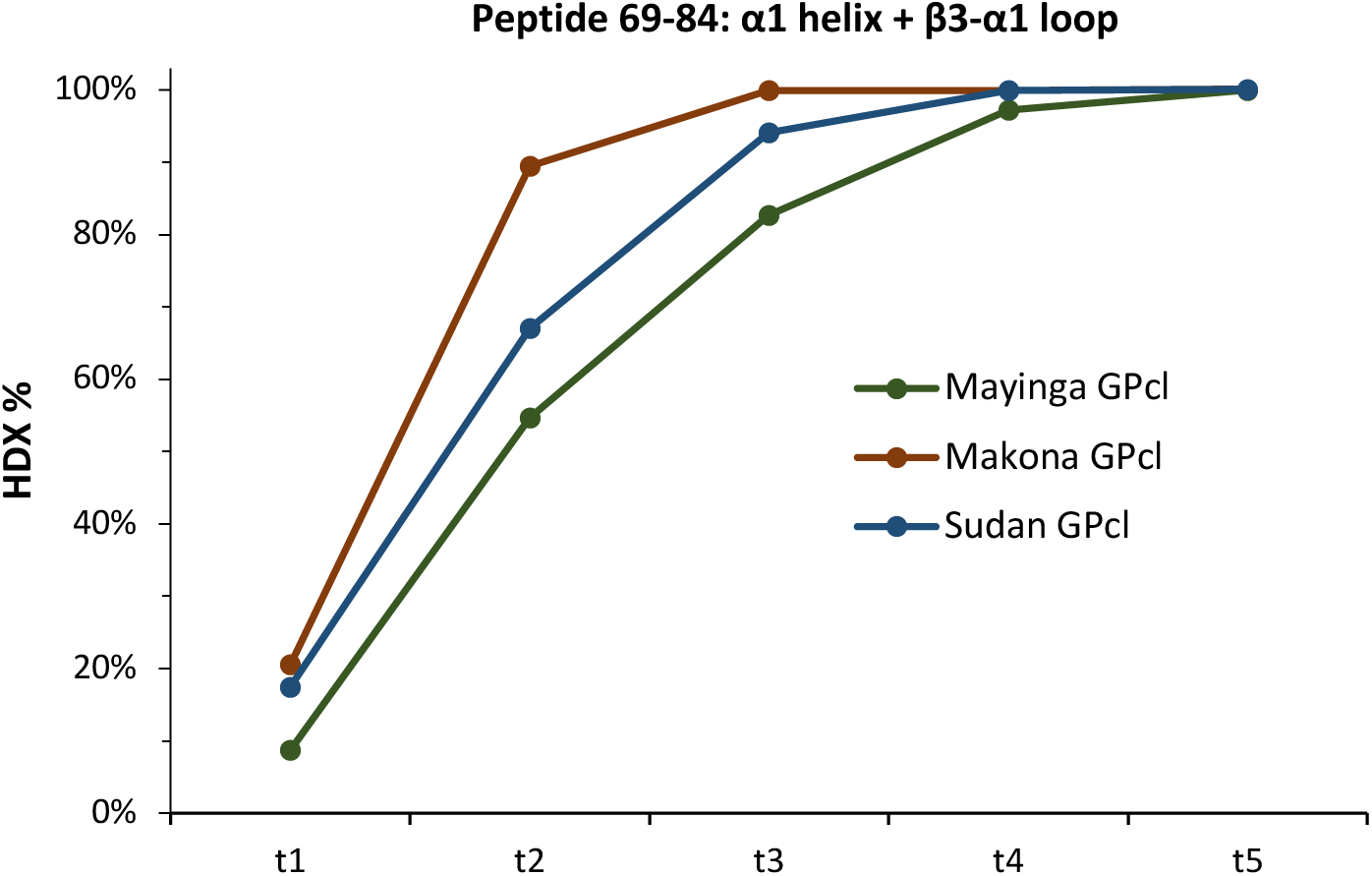
Deuterium uptake plot of peptide 69-84 (spanning the α1 helix and β3-α1 loop) exemplifies the potentiation of the RBR-fusion peptide axis of Makona and Sudan GPcl compared to Mayinga GPcl. Source data are included in source data files.

**Fig. S23.**
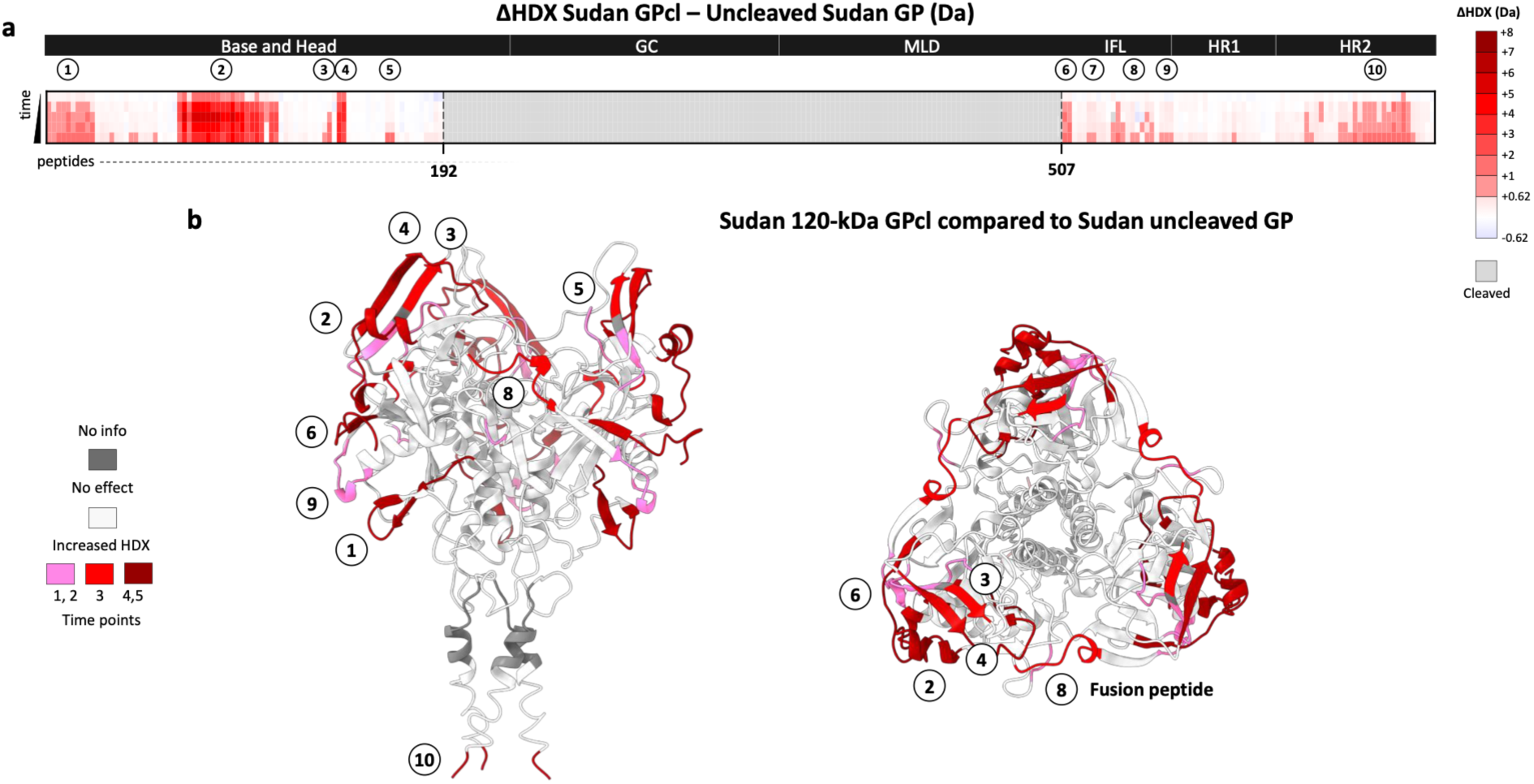
Differences in HDX (ΔHDX) between Sudan GPcl and Sudan uncleaved GP at pH 5. **a)** Plot illustrating ΔHDX profile along the protein sequence. **b)** ΔHDX profile superimposed on GP structures (PDB 5JQ3) where cleaved segments have been removed. Source data are included in source data files.

**Fig. S24.**
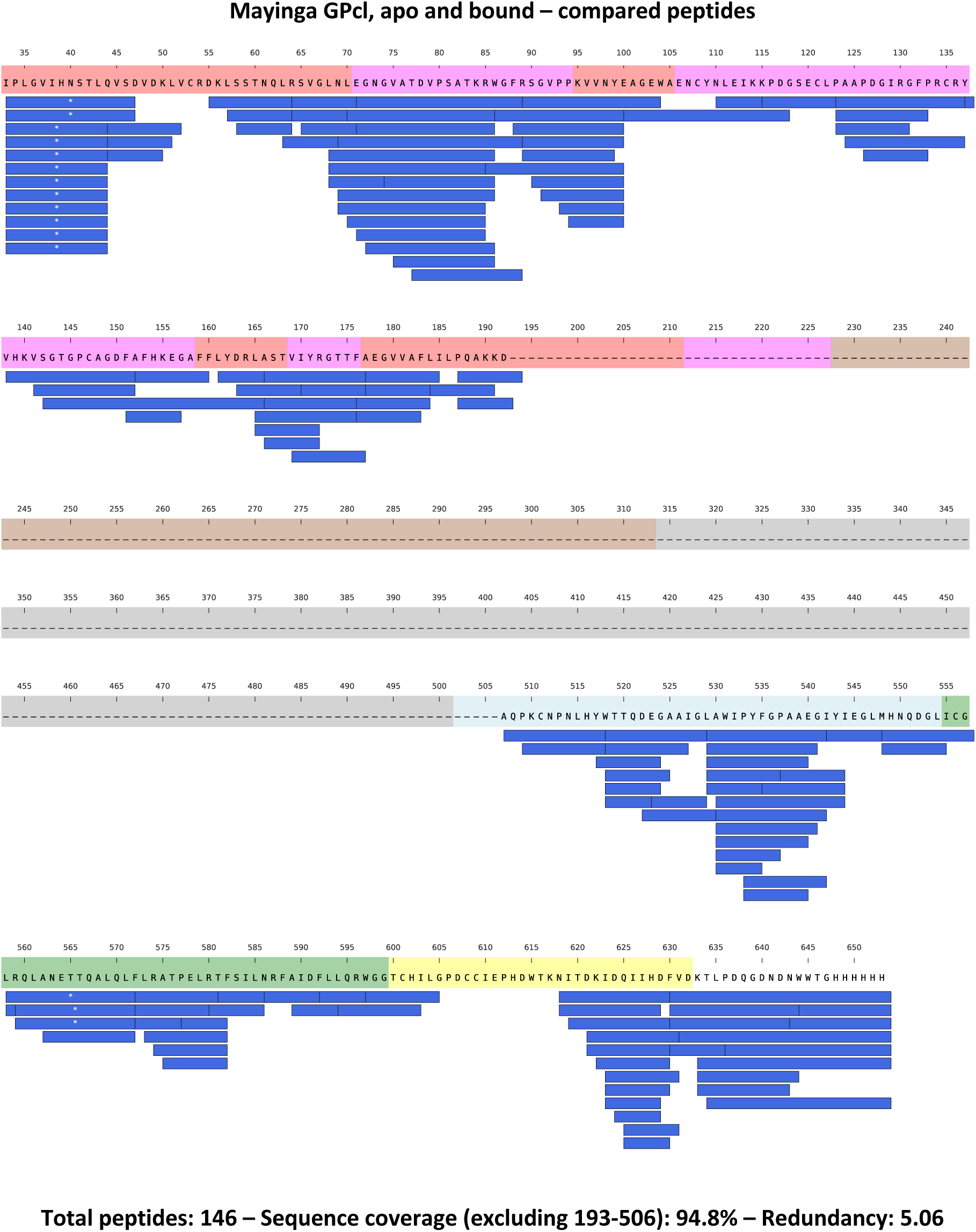
Peptides whose HDX was followed for Mayinga GPcl (apo and NPC1 bound) are illustrated along the protein sequence with blue bars. Blue bars with white asterisks (*) indicate glycosylated peptides. The dashed line indicates the cleaved portion of GP, whose peptides are absent in the dataset. The individidual domains of GP are color-coded along the protein sequence as in Fig. S1.

**Fig. S25.**
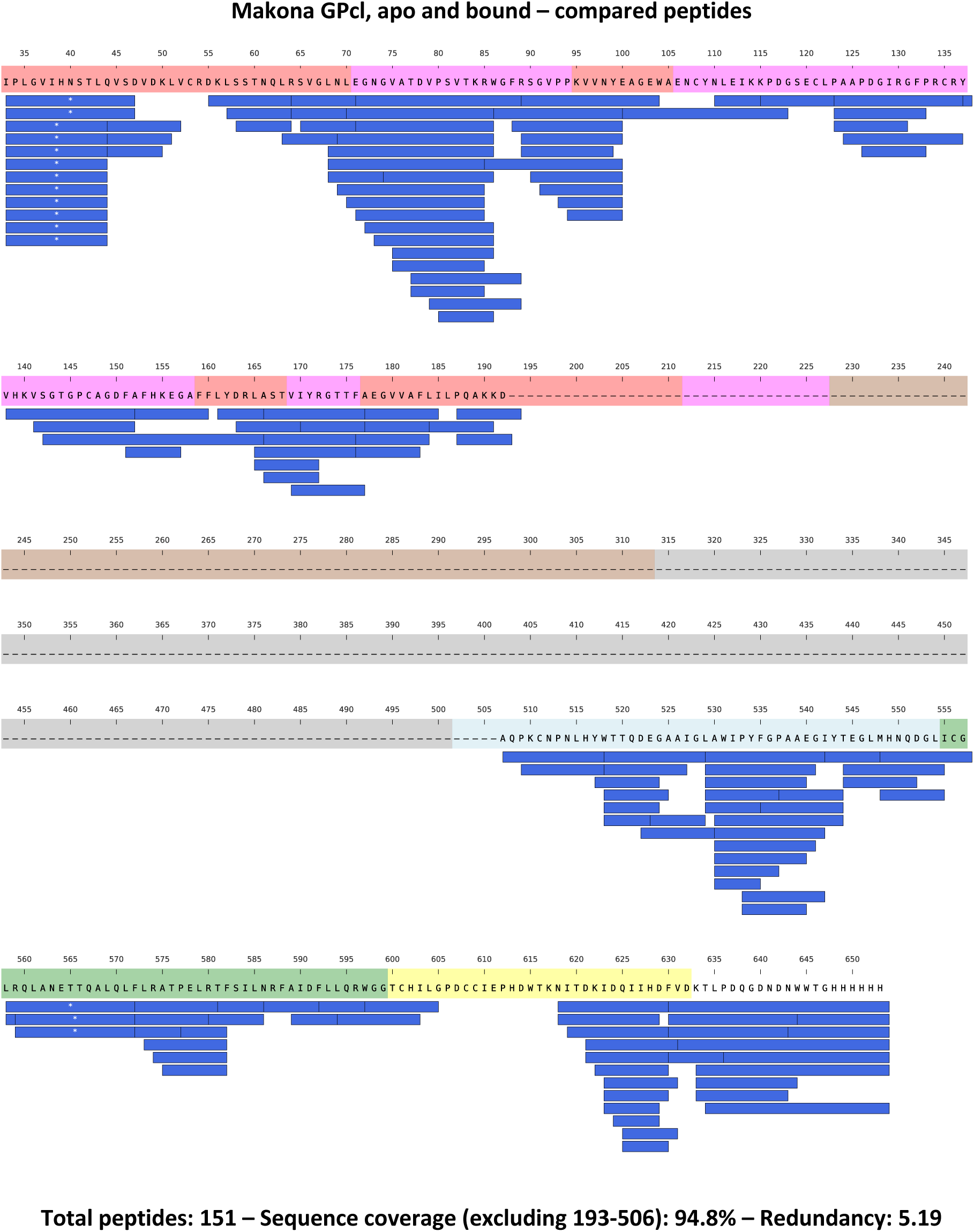
Peptides whose HDX was followed for Makona GPcl (apo and NPC1 bound) are illustrated along the protein sequence with blue bars. Blue bars with white asterisks (*) indicate glycosylated peptides. The dashed line indicates the cleaved portion of GP, whose peptides are absent in the dataset. The individidual domains of GP are color-coded along the protein sequence as in Fig. S1.

**Fig. S26.**
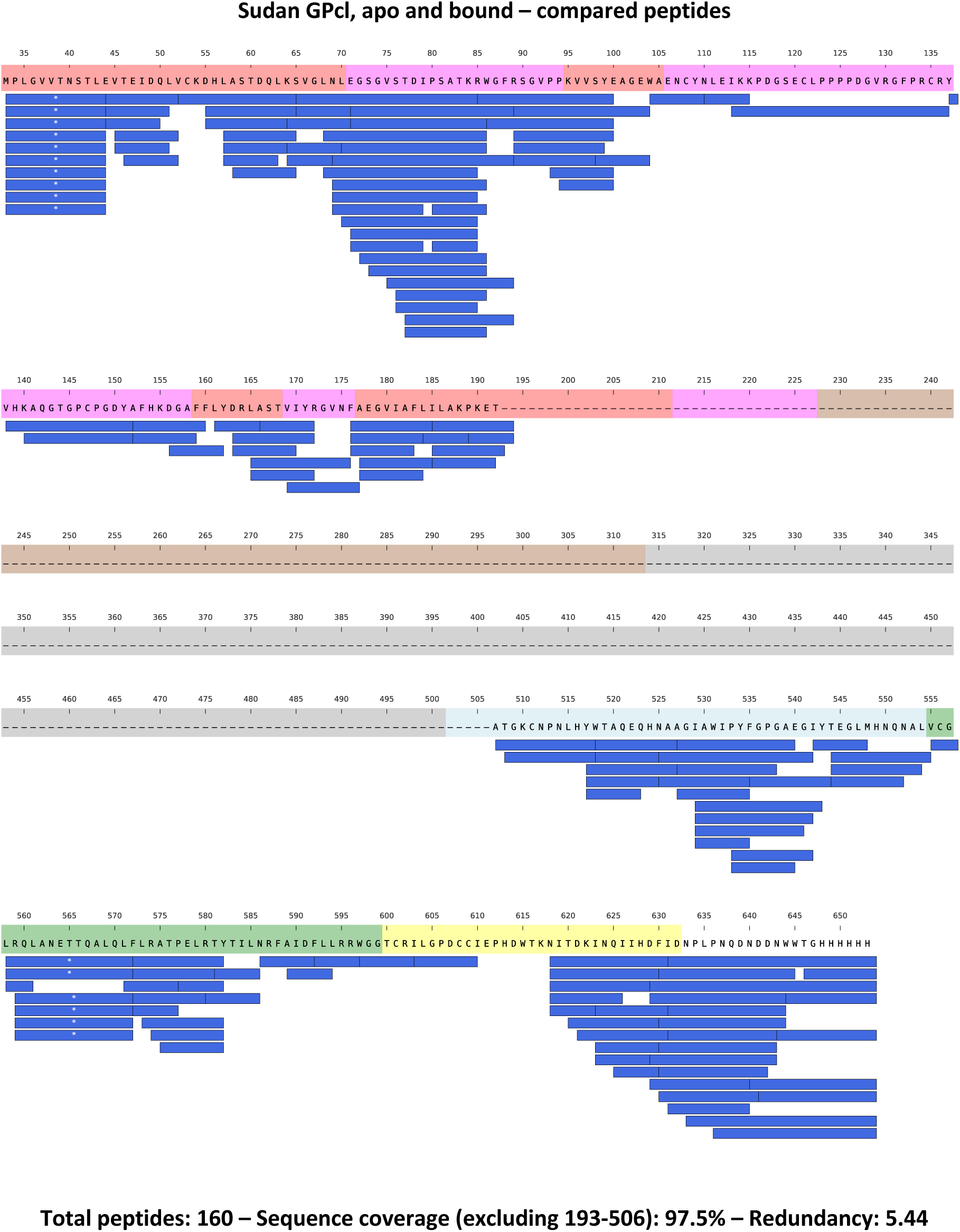
Peptides whose HDX was followed for Sudan GPcl (apo and NPC1 bound) are illustrated along the protein sequence with blue bars. Blue bars with white asterisks (*) indicate glycosylated peptides. The dashed line indicates the cleaved portion of GP, whose peptides are absent in the dataset. The individidual domains of GP are color-coded along the protein sequence as in Fig. S1.

**Fig. S27.**
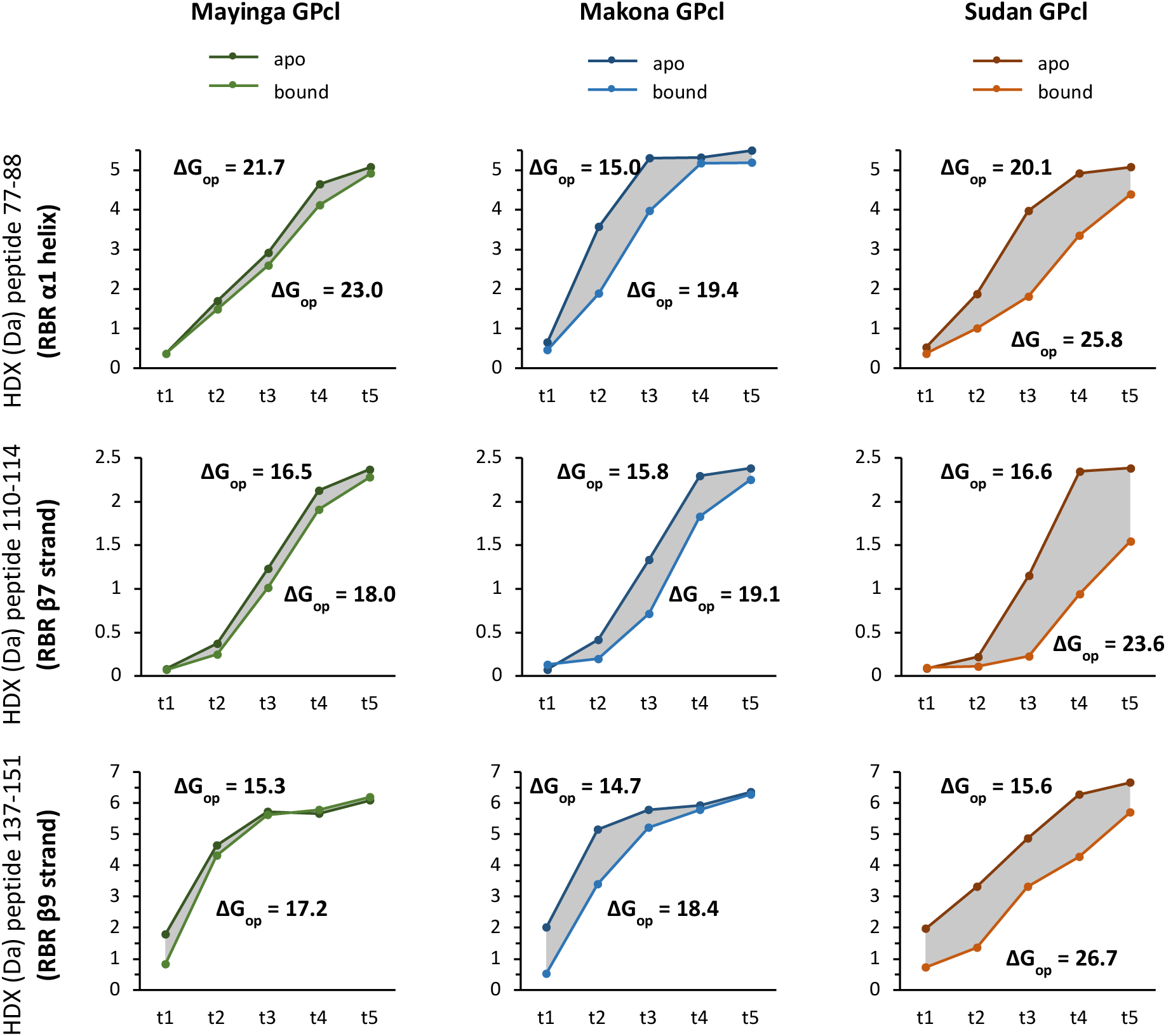
Deuterium uptake plots of model peptides spanning the binding cavity in the apo and bound GPcl across trimers illustrate that receptor binding is stronger for Sudan, followed by Makona and Mayinga GPcl. ΔG_opening_ (ΔG_op_) calculated from HDX data for the apo and bound state are included in each plot. The differences between ΔGopening of apo and bound state yields the ΔGbinding. Source data are included in source data files.

**Fig. S28.**
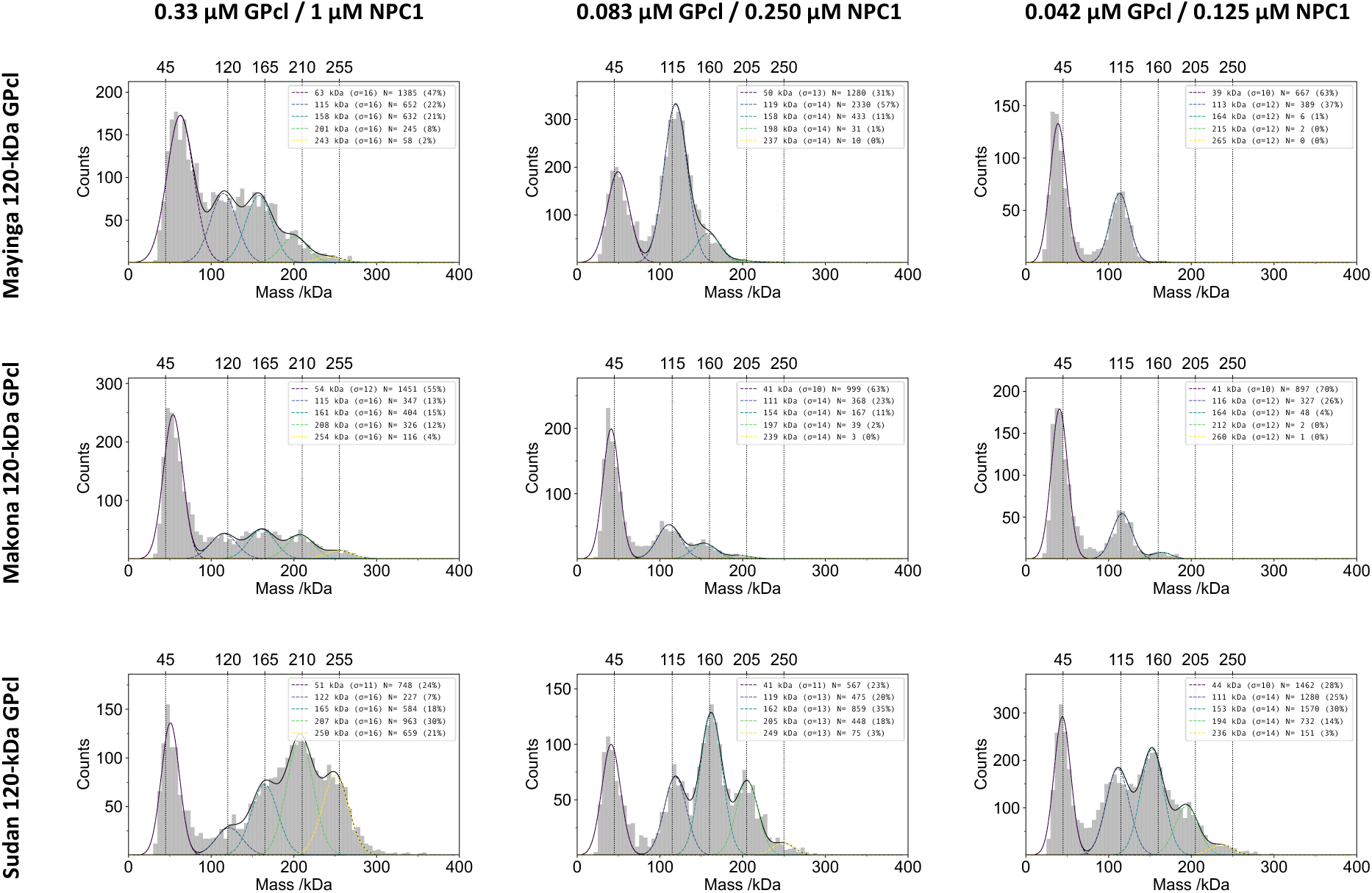
Examples of MP histograms of 120-kDa GPcl (Mayinga, Makona and Sudan) bound to NPC1 at concentrations 0.33 µM GPcl / 1 µM NPC1, 0.083 µM GPcl / 0.250 µM NPC1, and 0.042 µM GPcl / 0.125 µM NPC1. Concentration 0.167 µM GPcl / 0.5 µM NPC1 is in Fig. 6c of the main text. First peak: free NPC1, second peak: free GPcl, third peak: GPcl bound to 1 NPC1, fourth peak: GPcl bound to 2 NPC1, fifth peak: GPcl bound to 3 NPC1. The histograms report the relative % of the five species in solution. To note, the peak of free NPC1 exhibited significant low-mass noise, causing a high variability in its amplitude that does not reflect its effective abundance in solution, therefore it was not considered for the K_D_ calculations.

**Fig. S29.**
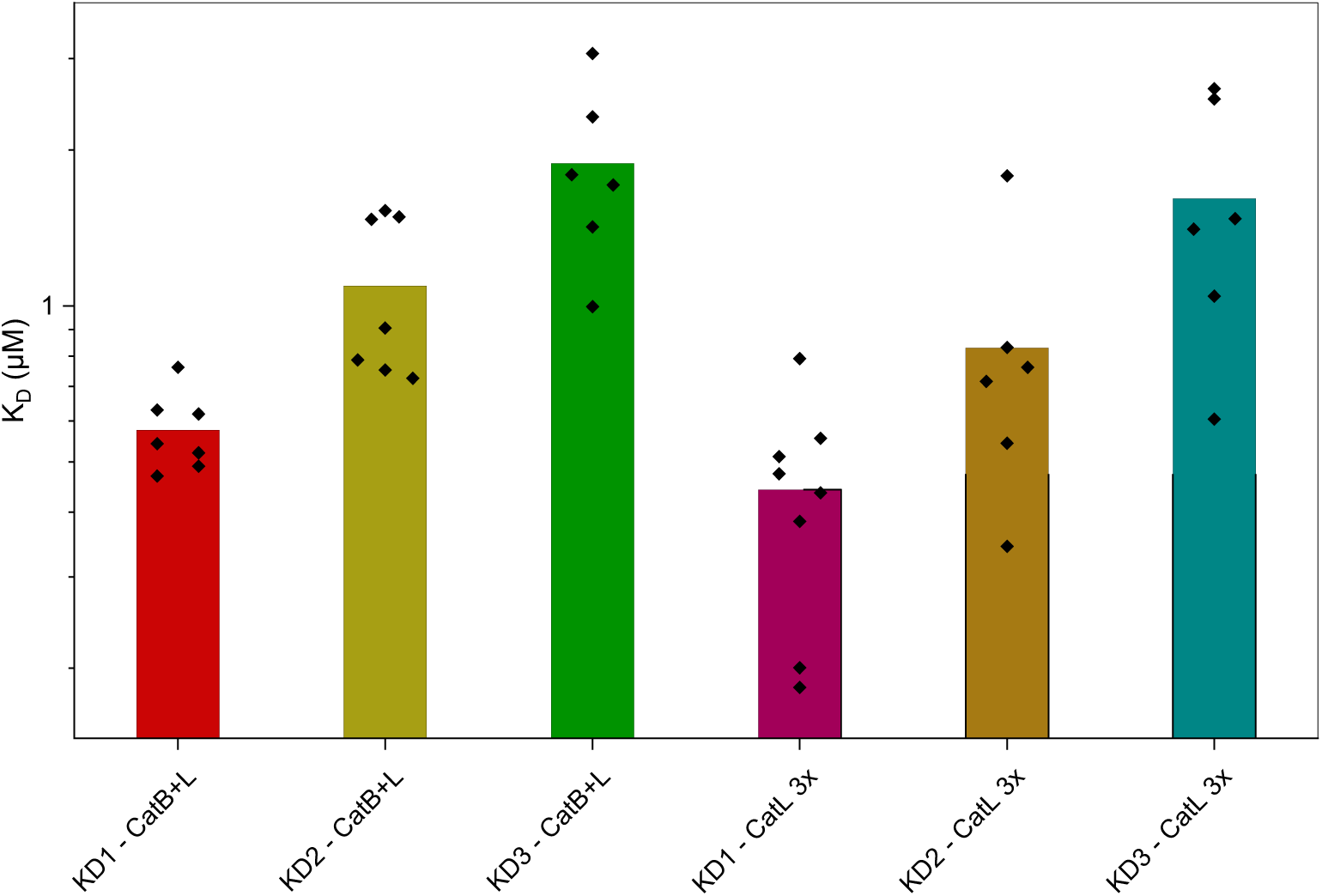
Binding affinity measurements of Makona 120-kDa GP to NPC1 generated upon digestion with CatL alone (3x) or CatL mixed with CatB. The data show that the K_D_1, K_D_2 and K_D_3 are comparable. Data were acquired by MP. Calculated K_D_ values (µM) are labelled on top of the bars. Source data are included in source data files.

**Fig. S30.**
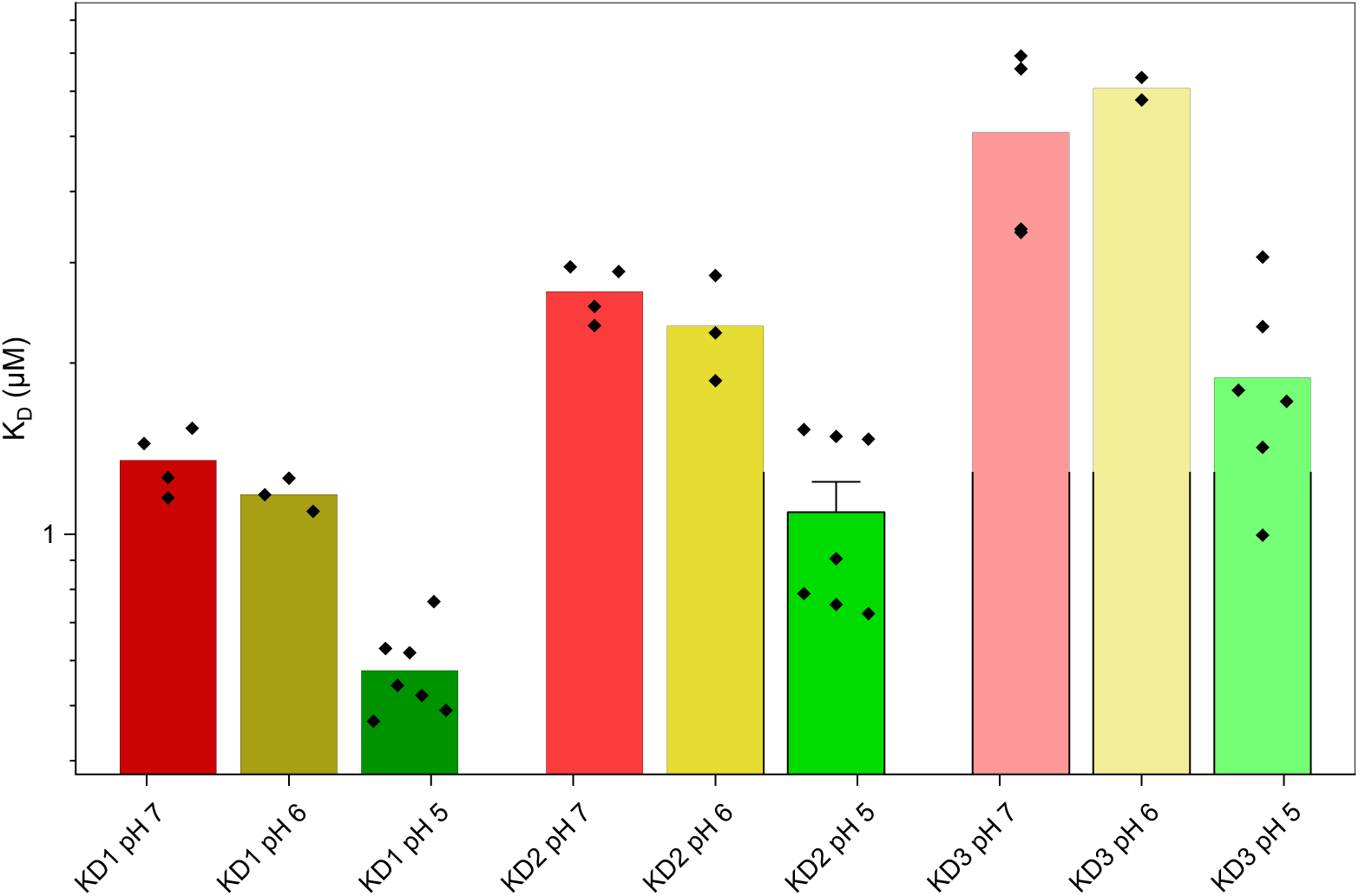
Dependence of GPcl-NPC1 binding on pH. Binding affinity measurements of Makona 120-kDa GP to NPC1 at pH 7, 6 and 5. The data show that the K_D_1, K_D_2 and K_D_3 increase with the pH increasing. Data were acquired by MP. Calculated K_D_ values (µM) are labelled on top of the bars. Source data are included in source data files.

**Fig. S31.**
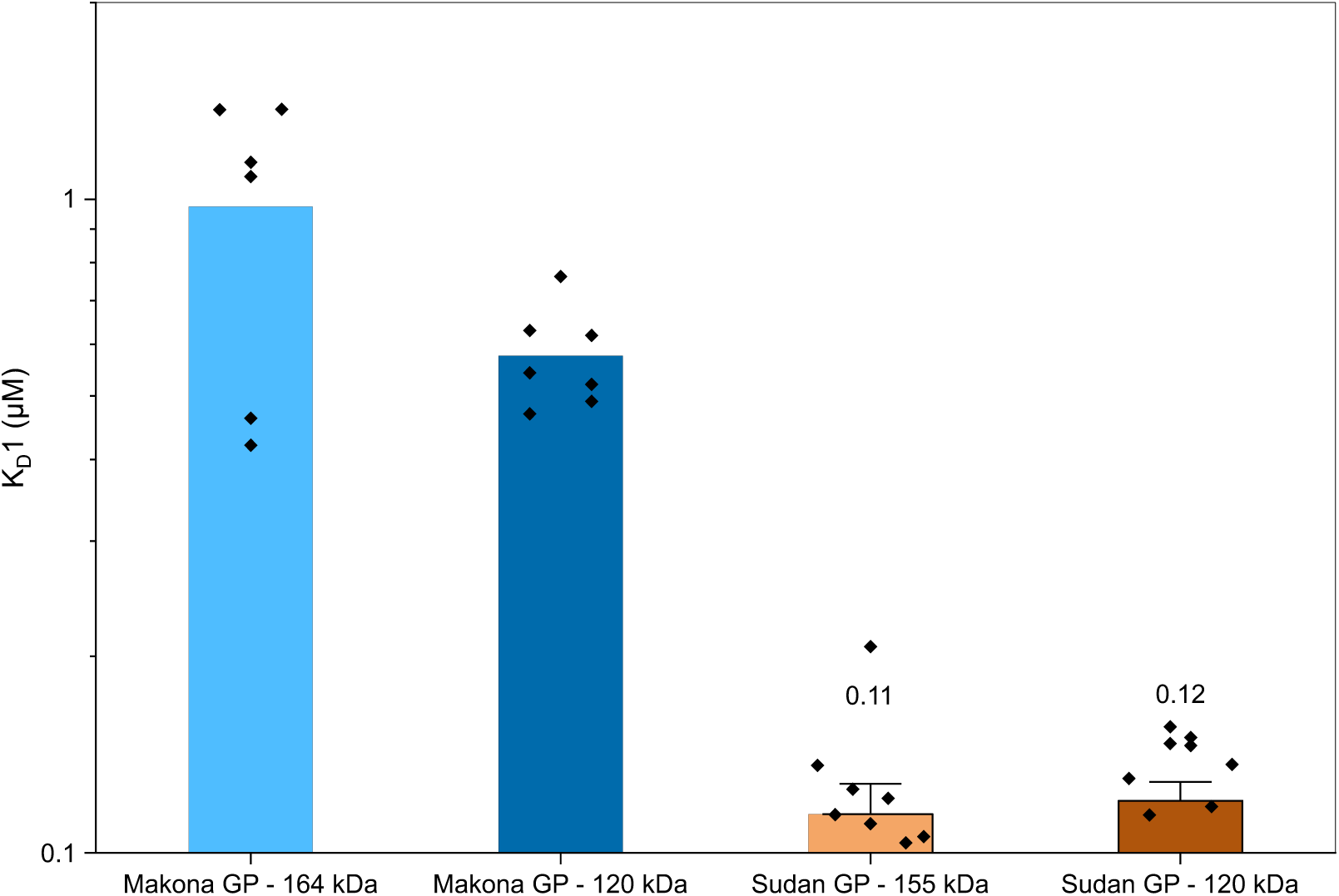
Binding affinity measurements (K_D_1) of Makona 164-kDa GP and Sudan 155-kDa GP compared to Makona and Sudan 120-kDa GP. The KD1 values show that binding affinity is lower in Makona 164-kDa compared to 120-kDa GP, but it is equal in Sudan 155-kDa compared to 120-kDa GP. Calculated K_D_1 values (µM) are labelled on top of the bars. Source data are included in source data files.

**Fig. S32.**
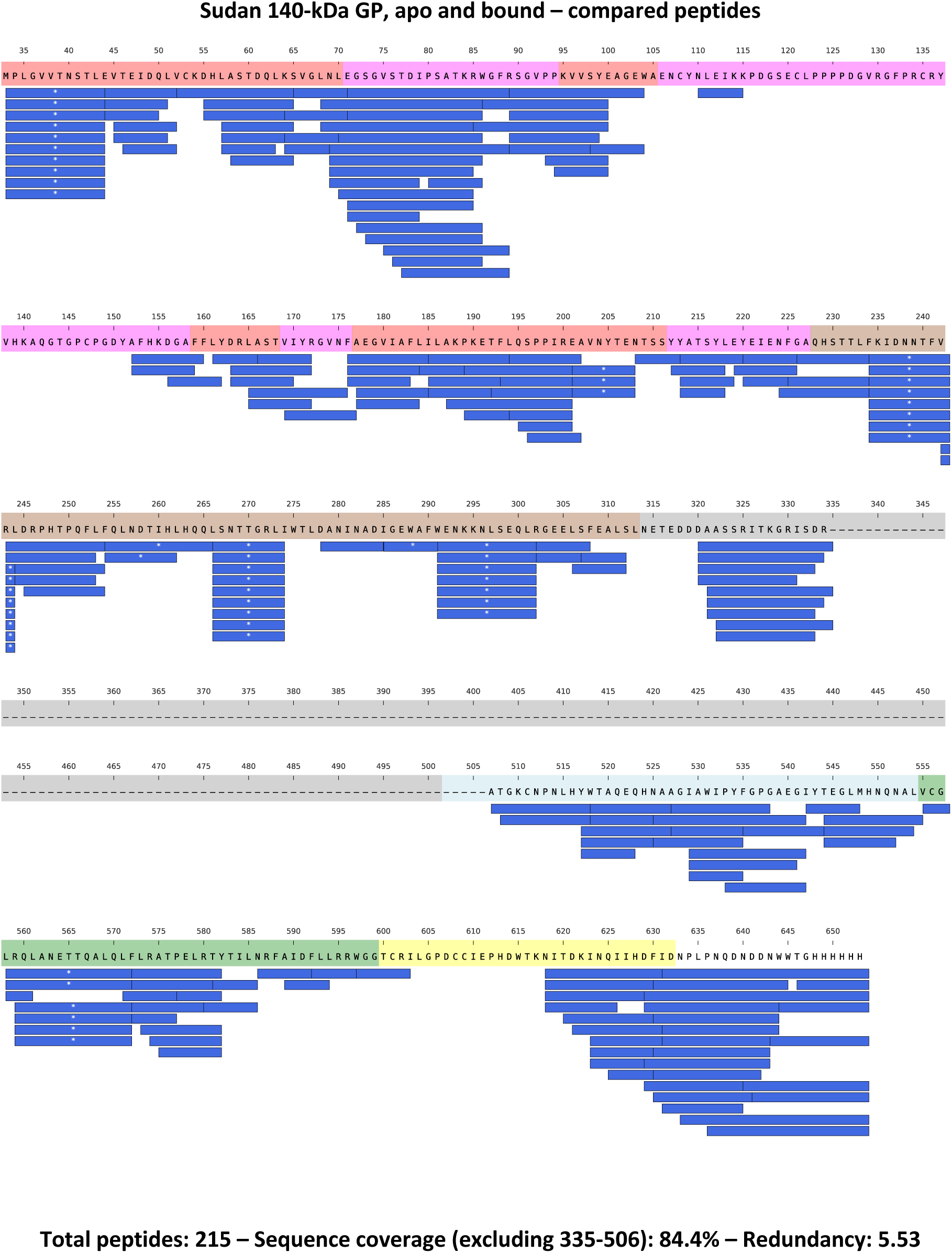
Peptides whose HDX was followed for Sudan 140-kDa GP (apo and NPC1 bound) are illustrated along the protein sequence with blue bars. Blue bars with white asterisks (*) indicate glycosylated peptides. The dashed line indicates the cleaved portion of GP, whose peptides are absent in the dataset. The individidual domains of GP are color-coded along the protein sequence as in Fig. S1.

**Fig. S33.**
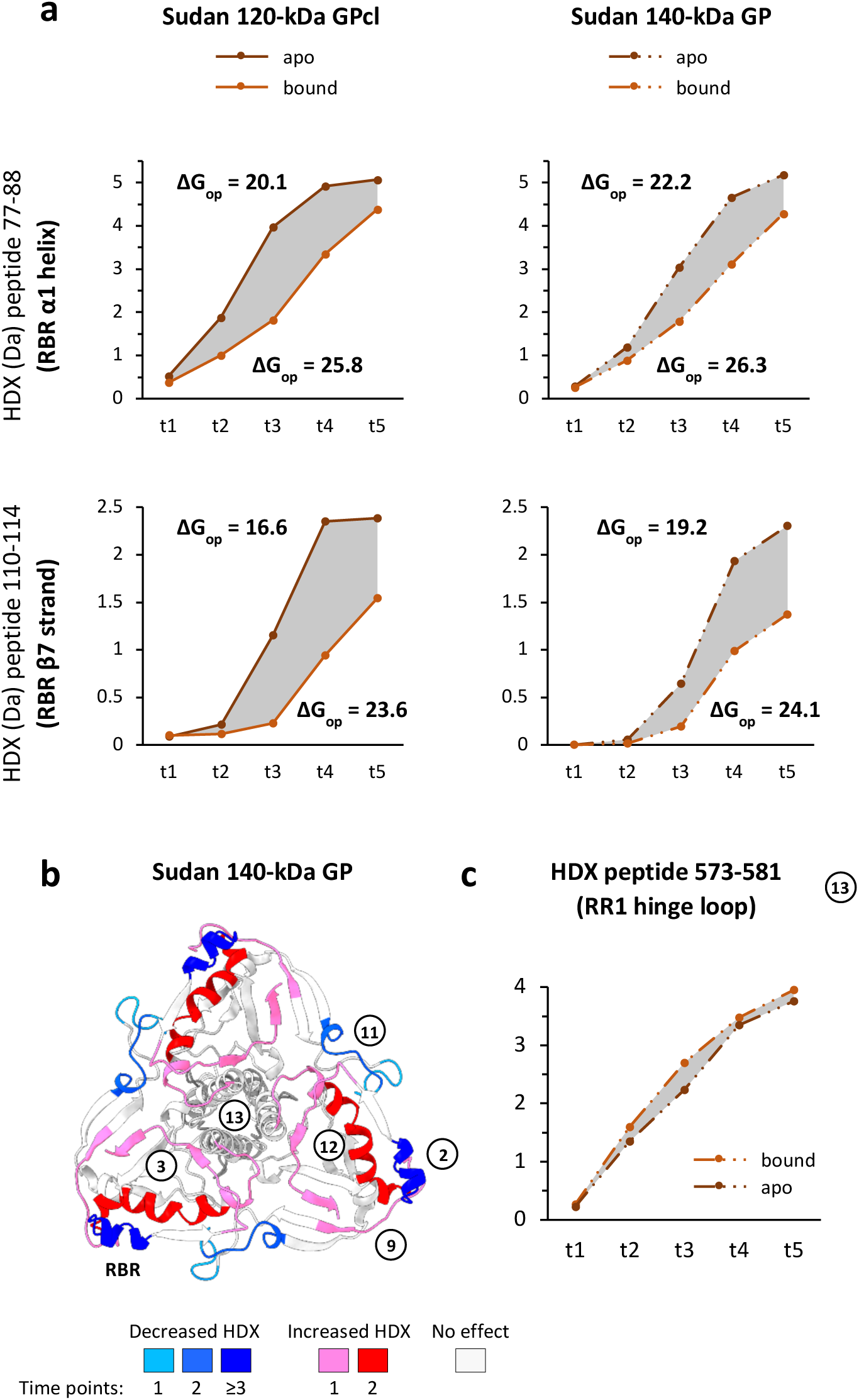
**a)** Deuterium uptake plots of a model peptide spanning the binding cavity in the apo and bound Sudan 140-kDa GP and 120-kDa GPcl across illustrate that receptor binding is stronger for Sudan GPcl compared to 140-kDa GP. ΔG_opening_ (ΔG_op_) calculated from HDX data for the apo and bound state are included in each plot. The differences between ΔGopening of apo and bound state yields the ΔGbinding. **b)** The ΔHDX profile NPC1 bound - apo Sudan 140-kDa GP superimposed on the GP structure (PDB 5JQ3) shows increased HDX in the elements of the trimer fusion machinery. **c)** Deuterium uptake plot of a model peptides spanning the RR1 hinge loop exemplifies the magnitude of the priming effect in Sudan 140-kDa GP. Source data are included in source data files.

**Fig. S34.**
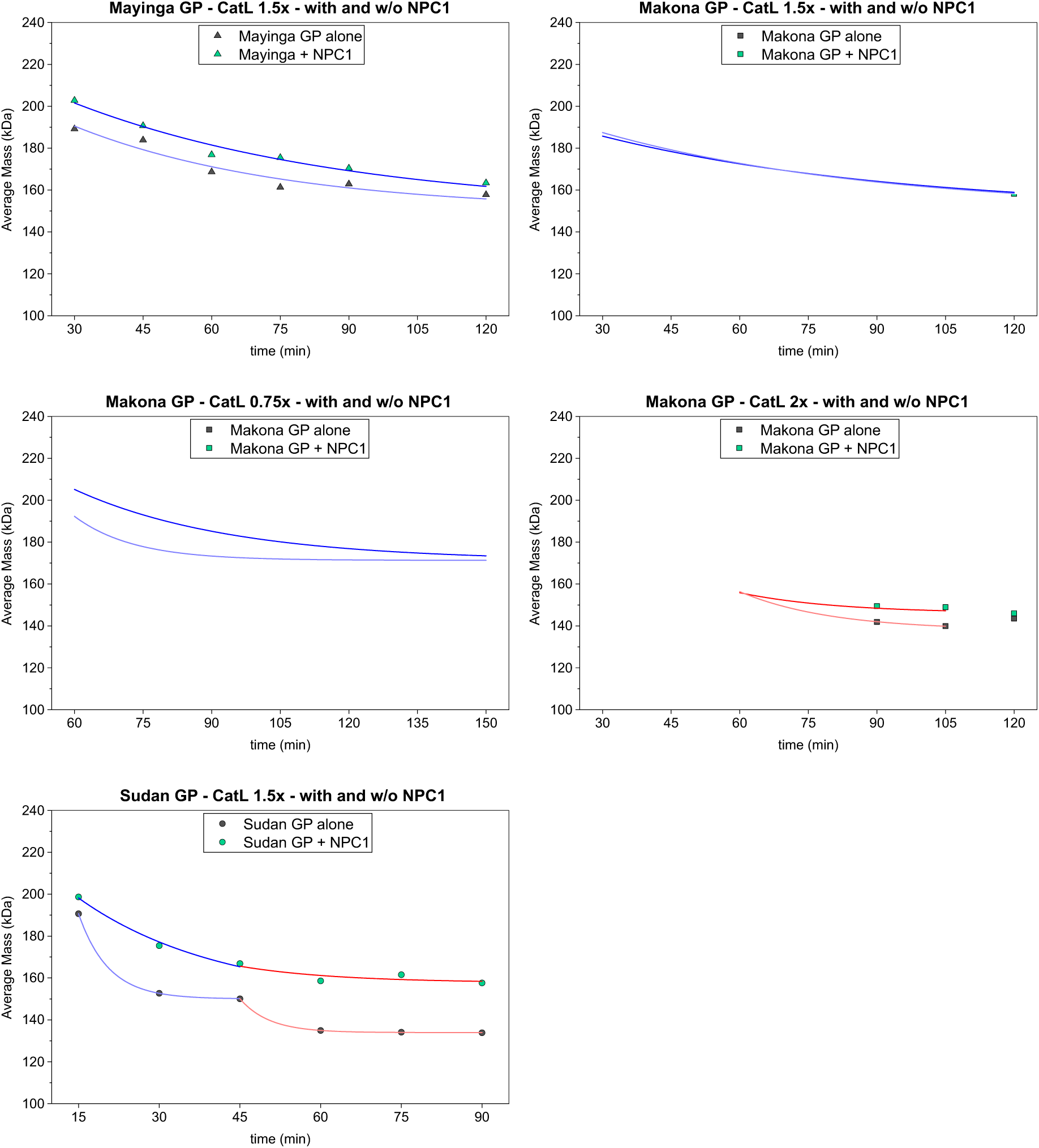
CatL cleavage kinetics for apo and receptor-bound GP trimers. Plots illustrating the mass of GP trimers measured by MP over the course of the cathepsin cleavage reactions conducted in the presence and absence of NPC1. For Mayinga and Sudan GP, digestions were conducted with 1.5x CatL (pH 4.5); for Makona GP, digestions were conducted with 0.75, 1.5 and 2x (pH 4.5). Blue and light blue lines: fitted curves in the range of ∼190-150 kDa (range 3-1.5 GC) of bound and apo GP, respectively; red and pink lines: fitted curves in the range of ∼150-130 kDa (range 1.5-0 GC) of bound and apo GP, respectively. Tables: ΔAUC: area between the curves of the apo and bound GP. ΔAUC/ Δtime: average mass difference between the two curves indicating the approximate fraction of 22-kDa GC that is still carried by bound GP compared to apo GP, hence the extent of cleavage rate reduction. Source data are included in source data files.

**Fig. S35.**
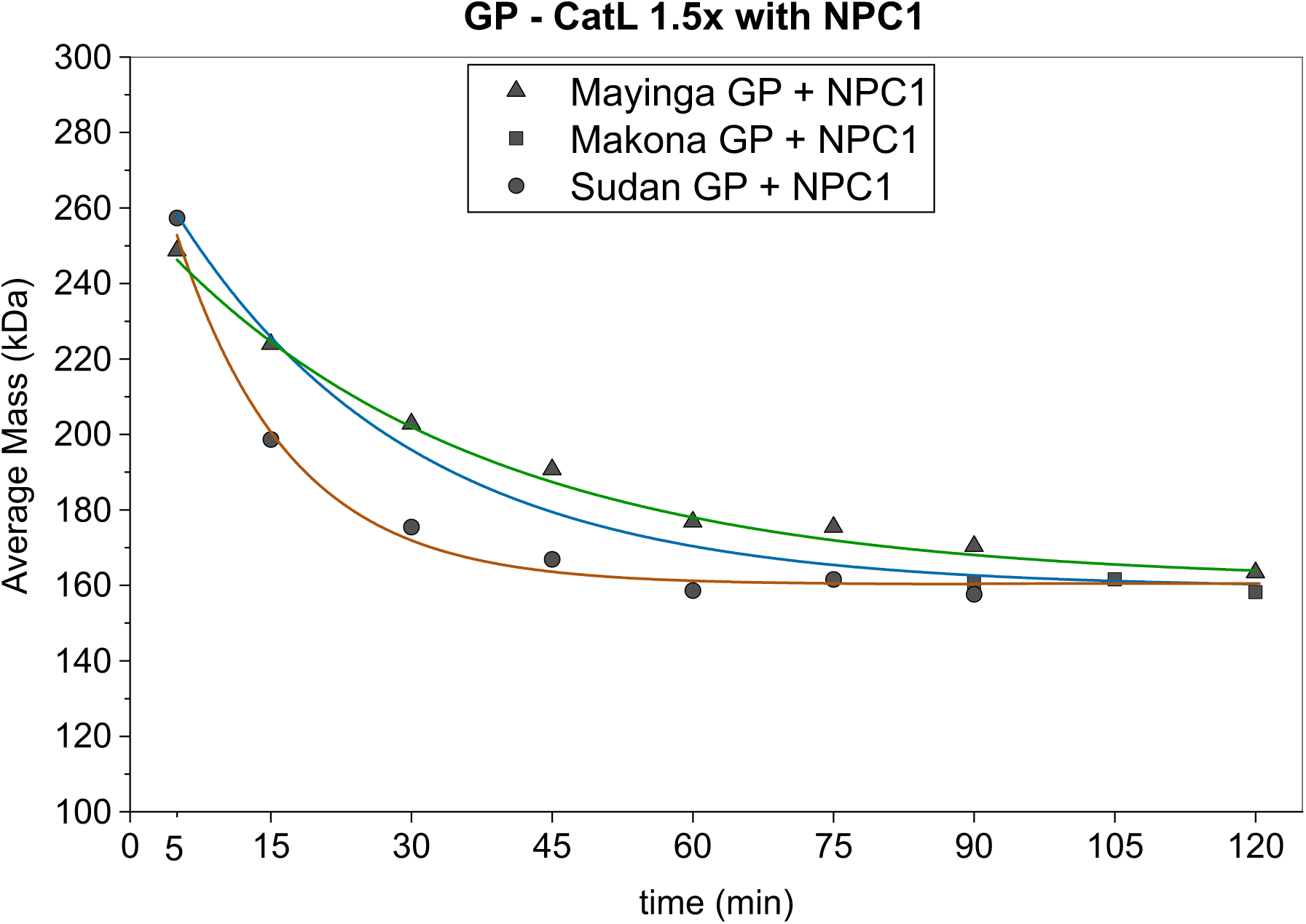
Cleavage kinetics for the receptor-bound GP trimers digested with CatL 1.5x. Green line: fitted curve of Mayinga GP; blue line: fitted curve of Makona GP; brown line: fitted curve of Sudan GP. The table indicate the plateaus reached, the cleavage rate and the precision of the fitting. Source data are included in source data files.

**Table S1.** Kinetic parameters of the various digestion conditions tested on Makona GP (optimization).

| Digestion conditions | Kinetic parameters | Makona GP |
| --- | --- | --- |
| 1x CatB<br>pH 5 | Plateau (kDa)<br>k (1/min)<br>Adj. R-Square | NA<br>NA<br>NA |
| 1x CatB<br>pH 4.5 | Plateau (kDa)<br>k (1/min)<br>Adj. R-Square | 209 ± 8.04<br>0.017 ± 0.003<br>0.96745 |
| 1x CatB<br>pH 4 | Plateau (kDa)<br>k (1/min)<br>Adj. R-Square | 160 ± 5.18<br>0.068 ± 0.013<br>0.91579 |
| 2x CatL<br>pH 5<br>fitted until ~164 kDa | Plateau (kDa)<br>k (1/min)<br>Adj. R-Square | 162 ± 1.50<br>0.055 ± 0.002<br>0.99797 |
| 2x CatL<br>pH 4.5<br>fitted until ~164 kDa | Plateau (kDa)<br>k (1/min)<br>Adj. R-Square | 164 ± 4.86<br>0.072 ± 0.009<br>0.9738 |
| 2x CatL<br>pH 4<br>fitted until ~164 kDa | Plateau (kDa)<br>k (1/min)<br>Adj. R-Square | 164 ± 5.41<br>0.099 ± 0.014<br>0.96949 |
| 2x CatB<br>pH 4 | Plateau (kDa)<br>k (1/min)<br>Adj. R-square | 168 ± 2.60<br>0.135 ± 0.018<br>0.91459 |
| 3x CatB<br>pH 4 | Plateau 1 (kDa)<br>k1 (1/min)<br>Adj. R-Square 1<br>Plateau 2 (kDa)<br>k2 (1/min)<br>Adj. R-Square 2 | 158± 5.03<br>0.342 ± 0.066<br>0.78667<br>122 ± 0<br>0.024 ± 0.004<br>0.90534 |
| 2x CatL<br>pH 4 | Plateau 1 (kDa)<br>k1 (1/min)<br>Adj. R-Square 1<br>Plateau 2 (kDa)<br>k2 (1/min)<br>Adj. R-Square 2 | 164 ± 4.62<br>0.103 ± 0.010<br>0.99312<br>121 ± 9.73<br>0.024 ± 0.010<br>0.97532 |
| 3x CatL<br>pH 4 | Plateau 1 (kDa)<br>k1 (1/min)<br>Adj. R-Square 1<br>Plateau 2 (kDa)<br>k2 (1/min)<br>Adj. R-Square 2 | 167 ± 11.88<br>0.693 ± 0.247<br>0.68402<br>121 ± 2.26<br>0.069 ± 0.025<br>0.87257 |

**Table S2.** Kinetic parameters of the various digestion conditions tested for Mayinga, Makona and Sudan GP to infer cleavage kinetics. Source data are included in the source data file.

| Digestion conditions | Kinetic parameters | Mayinga GP | Makona GP | Sudan GP |
| --- | --- | --- | --- | --- |
| 0.5x CatB<br>pH 4<br>fitted until 75 min | Plateau (kDa)<br>k (1/min)<br>Adj. R-square | 220 ± 0.73<br>0.123 ± 0.006<br>0.99266 | 236 ± 5.69<br>0.067 ± 0.024<br>0.85841 | 271 ± 1.69<br>0.130 ± 0.044<br>0.76547 |
| 1x CatB<br>pH 4 | Plateau (kDa)<br>k (1/min)<br>Adj. R-square | 151 ± 1.99<br>0.089 ± 0.007<br>0.97854 | 160 ± 5.18<br>0.068 ± 0.013<br>0.91579 | 232 ± 1.11<br>0.091 ± 0.009<br>0.96576 |
| 1x CatL<br>pH 4 | Plateau (kDa)<br>k (1/min)<br>Adj. R-square | 164 ± 3.36<br>0.045 ± 0.005<br>0.97038 | 162 ± 2.51<br>0.047 ± 0.004<br>0.98312 | 177 ± 2.45<br>0.059 ± 0.006<br>0.96991 |
| 1.5x CatL<br>pH 4.5 | Plateau (kDa)<br>k (1/min)<br>Adj. R-square | 162 ± 4.43<br>0.046 ± 0.005<br>0.97879 | 162 ± 9.11<br>0.044 ± 0.010<br>0.94407 | 132 ± 5.42<br>0.061 ± 0.008<br>0.96833 |
| 0.25x CatB + 0.25x CatL<br>pH 4 | Plateau (kDa)<br>k (1/min)<br>Adj. R-square | 215 ± 2.39<br>0.126 ± 0.016<br>0.95569 | 245 ± 1.33<br>0.100 ± 0.010<br>0.97965 | 215 ± 5.52<br>0.072 ± 0.017<br>0.92322 |
| 0.5x CatB + 0.25x CatL<br>pH 4 | Plateau (kDa)<br>k (1/min)<br>Adj. R-square | 173 ± 7.75<br>0.069 ± 0.015<br>0.95218 | 173 ± 12.37<br>0.047 ± 0.012<br>0.95516 | 118 ± 6.27<br>0.064 ± 0.007<br>0.98555 |
| 0.25x CatB + 0.5x CatL<br>pH 4 | Plateau (kDa)<br>k (1/min)<br>Adj. R-square | 164 ± 0<br>0.088 ± 0.006<br>0.96979 | 164 ± 0<br>0.073 ± 0.008<br>0.95023 | 120 ± 4.62<br>0.067 ± 0.008<br>0.96634 |
| 0.5x CatB + 0.5x CatL<br>pH 4 | Plateau (kDa)<br>k (1/min)<br>Adj. R-square | 120 ± 6.55<br>0.053 ± 0.006<br>0.97835 | 123 ± 1.93<br>0.054 ± 0.002<br>0.99807 | 108 ± 1.69<br>0.093 ± 0.004<br>0.99615 |
| 1.5x CatL + NPC1<br>pH 4.5 | Plateau (kDa)<br>k (1/min)<br>Adj. R-square | 161 ± 2.99<br>0.029 ± 0.003<br>0.99135 | 159 ± 3.74<br>0.040 ± 0.006<br>0.97395 | 160 ± 2.22<br>0.083 ± 0.010<br>0.98786 |

**Table S3.**
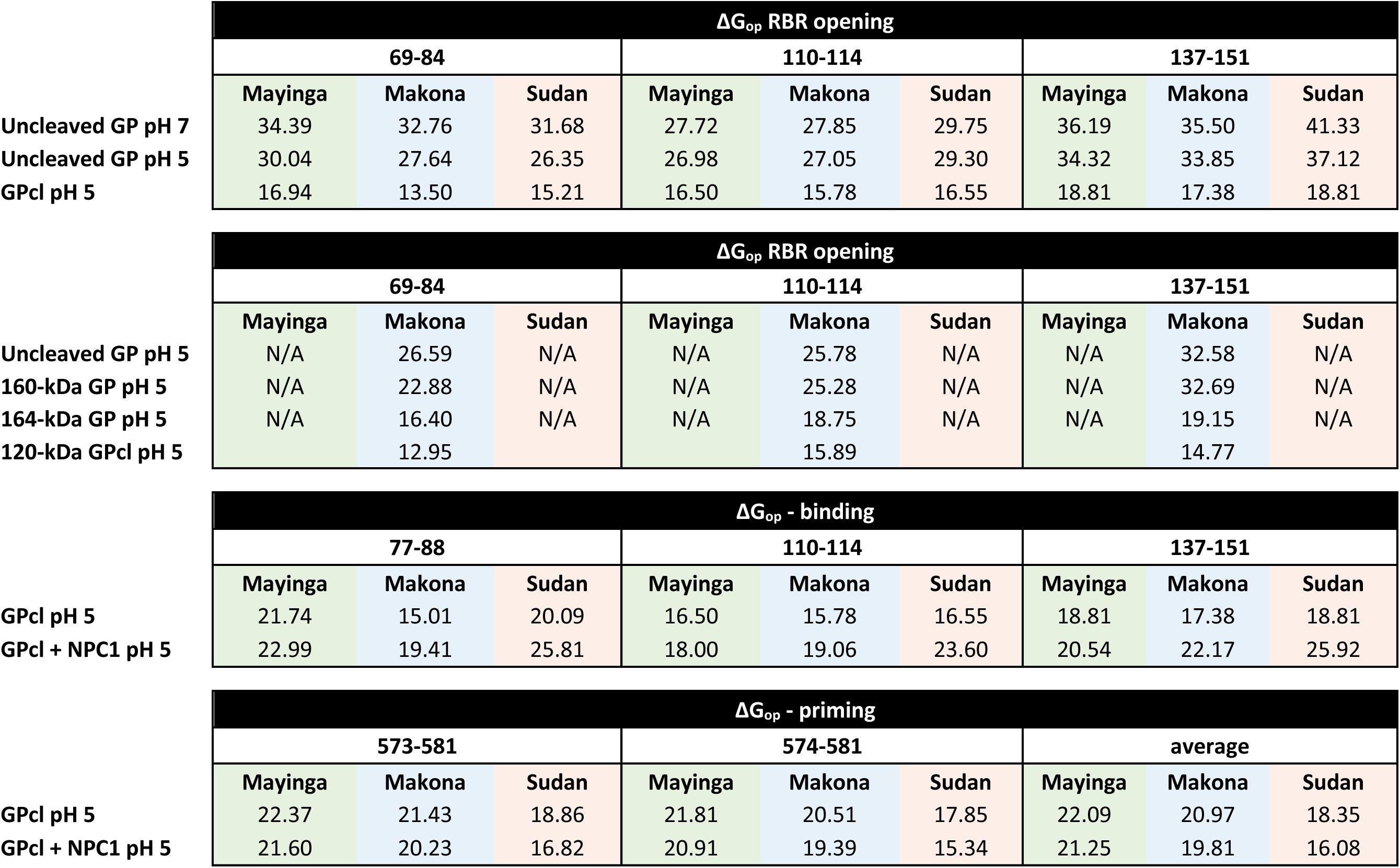
ΔG_opening_ (ΔG_op_ - kJ/mol) of the backbone amides calculated from HDX data of selected regions indicate the degree of opening of the receptor binding cavity, the strength of NPC1 binding and the extent of priming, across Mayinga, Makona and Sudan GP. Source data are included in the source data file.

**Table S4.**
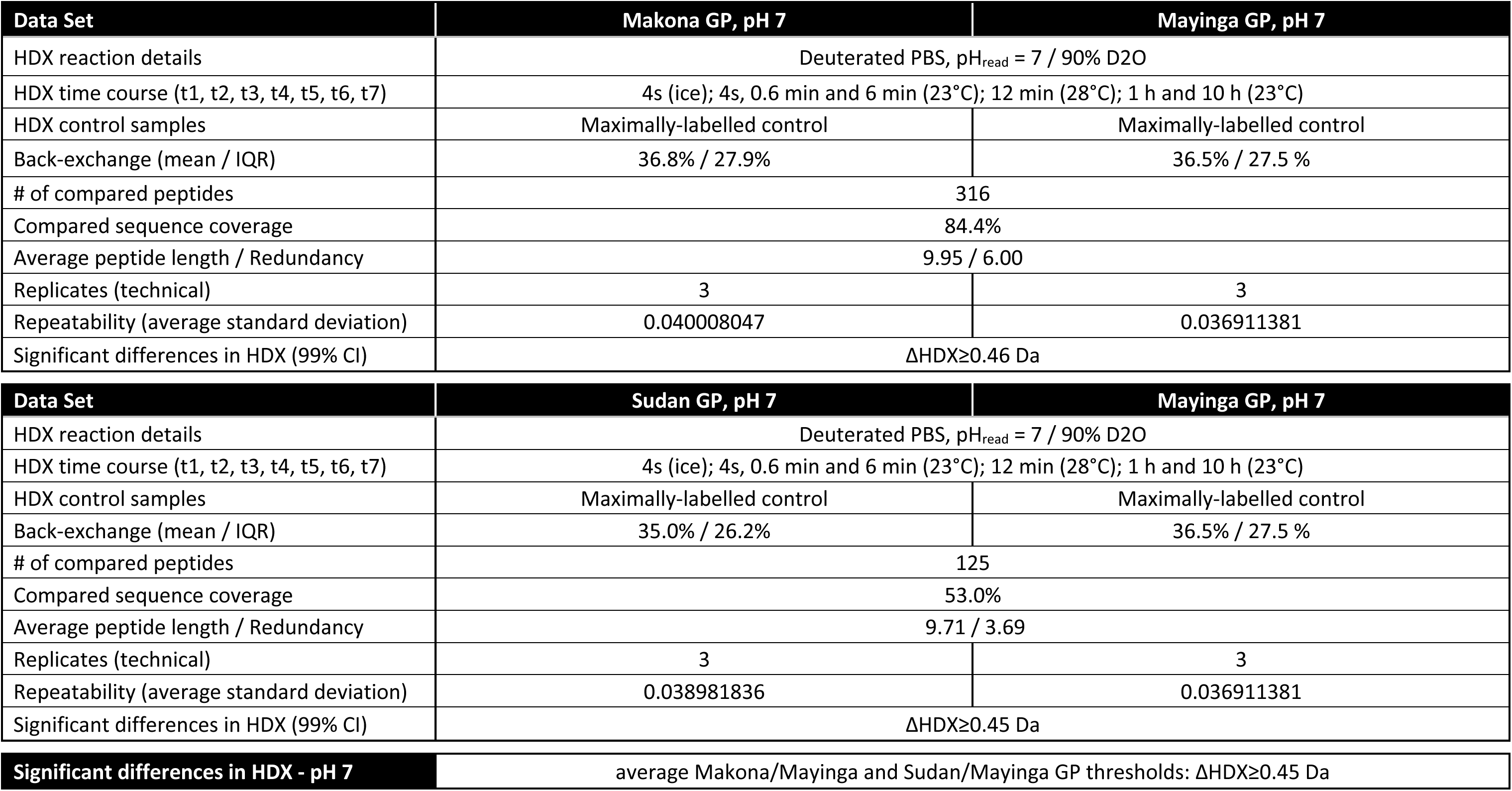
Summary of the experimental details of the HDX comparison across GP trimers at pH 7.

| Data Set | Makona GP, pH 7 | Mayinga GP, pH 7 |
| --- | --- | --- |
| HDX reaction details | Deuterated PBS, pH<sub>read</sub> = 7 / 90% D2O |  |
| HDX time course (t1, t2, t3, t4, t5, t6, t7) | 4s (ice); 4s, 0.6 min and 6 min (23°C); 12 min (28°C); 1 h and 10 h (23°C) |  |
| HDX control samples | Maximally-labelled control | Maximally-labelled control |
| Back-exchange (mean / IQR) | 36.8% / 27.9% | 36.5% / 27.5 % |
| # of compared peptides | 316 |  |
| Compared sequence coverage | 84.4% |  |
| Average peptide length / Redundancy | 9.95 / 6.00 |  |
| Replicates (technical) | 3 | 3 |
| Repeatability (average standard deviation) | 0.040008047 | 0.036911381 |
| Significant differences in HDX (99% CI) | $\Delta\text{HDX} \geq 0.46$ Da | |

| Data Set | Sudan GP, pH 7 | Mayinga GP, pH 7 |
| --- | --- | --- |
| HDX reaction details | Deuterated PBS, pH<sub>read</sub> = 7 / 90% D2O |  |
| HDX time course (t1, t2, t3, t4, t5, t6, t7) | 4s (ice); 4s, 0.6 min and 6 min (23°C); 12 min (28°C); 1 h and 10 h (23°C) |  |
| HDX control samples | Maximally-labelled control | Maximally-labelled control |
| Back-exchange (mean / IQR) | 35.0% / 26.2% | 36.5% / 27.5 % |
| # of compared peptides | 125 |  |
| Compared sequence coverage | 53.0% |  |
| Average peptide length / Redundancy | 9.71 / 3.69 |  |
| Replicates (technical) | 3 | 3 |
| Repeatability (average standard deviation) | 0.038981836 | 0.036911381 |
| Significant differences in HDX (99% CI) | $\Delta\text{HDX} \geq 0.45$ Da | |

|  |  |
| --- | --- |
| Significant differences in HDX - pH 7 | average Makona/Mayinga and Sudan/Mayinga GP thresholds: $\Delta\text{HDX} \geq 0.45$ Da |

**Table S5.**
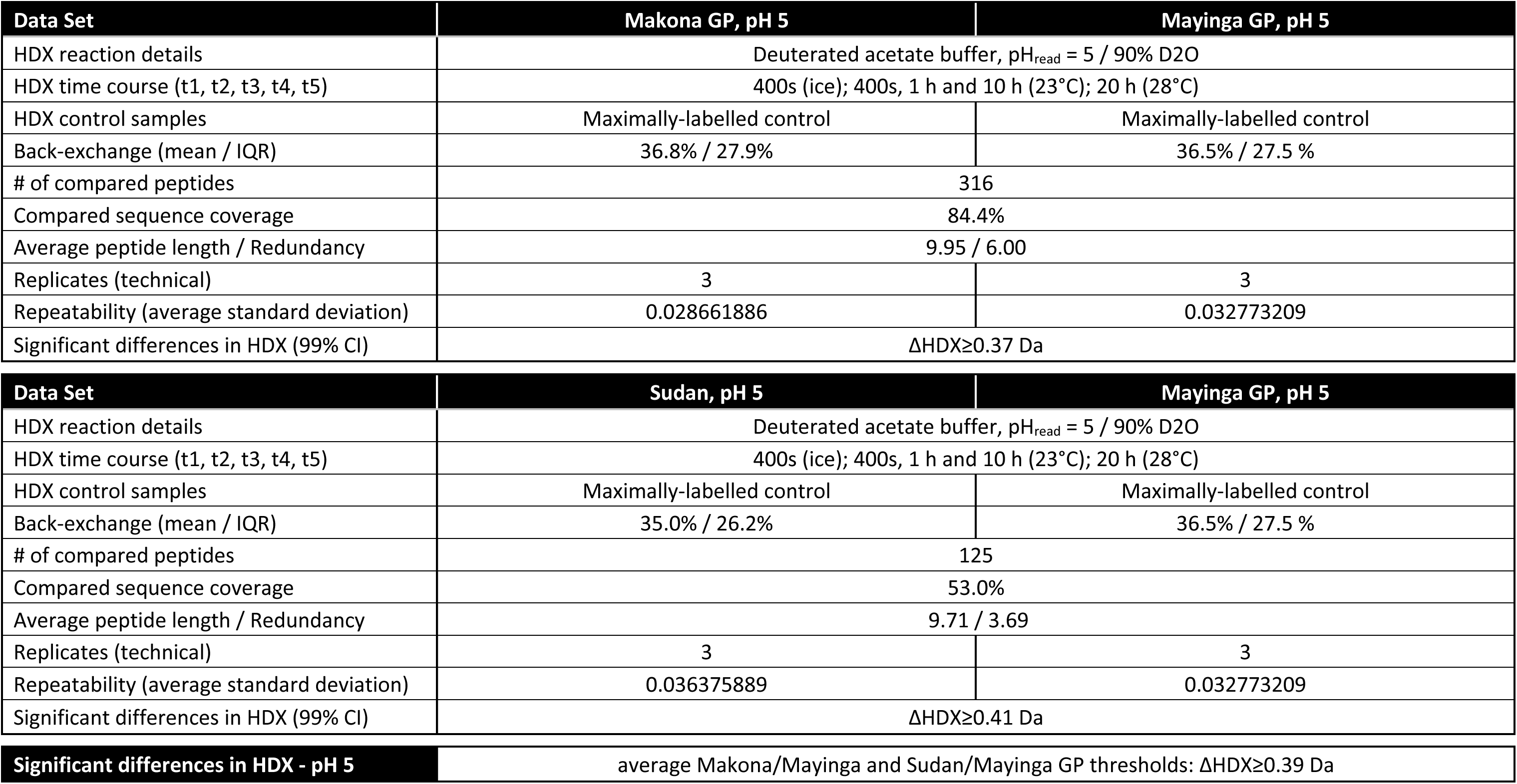
Summary of the experimental details of the HDX comparison across GP trimers at pH 5.

| Data Set | Makona GP, pH 5 | Mayinga GP, pH 5 |
| --- | --- | --- |
| HDX reaction details | Deuterated acetate buffer, pH<sub>read</sub> = 5 / 90% D2O |  |
| HDX time course (t1, t2, t3, t4, t5) | 400s (ice); 400s, 1 h and 10 h (23°C); 20 h (28°C) |  |
| HDX control samples | Maximally-labelled control | Maximally-labelled control |
| Back-exchange (mean / IQR) | 36.8% / 27.9% | 36.5% / 27.5 % |
| # of compared peptides | 316 |  |
| Compared sequence coverage | 84.4% |  |
| Average peptide length / Redundancy | 9.95 / 6.00 |  |
| Replicates (technical) | 3 | 3 |
| Repeatability (average standard deviation) | 0.028661886 | 0.032773209 |
| Significant differences in HDX (99% CI) | $\Delta\text{HDX} \geq 0.37$ Da | |

| Data Set | Sudan, pH 5 | Mayinga GP, pH 5 |
| --- | --- | --- |
| HDX reaction details | Deuterated acetate buffer, pH<sub>read</sub> = 5 / 90% D2O |  |
| HDX time course (t1, t2, t3, t4, t5) | 400s (ice); 400s, 1 h and 10 h (23°C); 20 h (28°C) |  |
| HDX control samples | Maximally-labelled control | Maximally-labelled control |
| Back-exchange (mean / IQR) | 35.0% / 26.2% | 36.5% / 27.5 % |
| # of compared peptides | 125 |  |
| Compared sequence coverage | 53.0% |  |
| Average peptide length / Redundancy | 9.71 / 3.69 |  |
| Replicates (technical) | 3 | 3 |
| Repeatability (average standard deviation) | 0.036375889 | 0.032773209 |
| Significant differences in HDX (99% CI) | $\Delta\text{HDX} \geq 0.41$ Da | |

|  |  |
| --- | --- |
| **Significant differences in HDX - pH 5** | average Makona/Mayinga and Sudan/Mayinga GP thresholds: $\Delta\text{HDX} \geq 0.39$ Da |

**Table S6.** Summary of the experimental details of the HDX comparison of each individual GP trimers at pH 5 and pH 7.

| Data Set | Mayinga GP, pH 7 | Mayinga GP, pH 5 |
| --- | --- | --- |
| HDX reaction details | Deuterated PBS, pH<sub>read</sub> = 7 / 90% D2O | Deuterated acetate buffer, pH<sub>read</sub> = 5 / 90% D2O |
| HDX time course (t1, t2, t3, t4, t5) | 4s (ice); 4s, 0.6 min and 6 min (23°C); 12 min (28°C) | 400s (ice); 400s, 1 h and 10 h (23°C); 20 h (28°C) |
| HDX control samples | Maximally-labelled control |  |
| # of peptides | 324 |  |
| Sequence coverage | 86.5% |  |
| Average peptide length / Redundancy | 10.04 / 6.06 |  |
| Replicates (technical) | 3 | 3 |
| Repeatability (average standard deviation) | 0.036911381 | 0.032773209 |

| Data Set | Makona GP, pH 7 | Makona GP, pH 5 |
| --- | --- | --- |
| HDX reaction details | Deuterated PBS, pH<sub>read</sub> = 7 / 90% D2O | Deuterated acetate buffer, pH<sub>read</sub> = 5 / 90% D2O |
| HDX time course (t1, t2, t3, t4, t5) | 4s (ice); 4s, 0.6 min and 6 min (23°C); 12 min (28°C) | 400s (ice); 400s, 1 h and 10 h (23°C); 20 h (28°C) |
| HDX control samples | Maximally-labelled control |  |
| # of peptides | 332 |  |
| Sequence coverage | 86.8% |  |
| Average peptide length / Redundancy | 10.12 / 6.23 |  |
| Replicates (technical) | 3 | 3 |
| Repeatability (average standard deviation) | 0.040008047 | 0.028661886 |

| Data Set | Sudan GP, pH 7 | Sudan GP, pH 5 |
| --- | --- | --- |
| HDX reaction details | Deuterated PBS, pH<sub>read</sub> = 7 / 90% D2O | Deuterated acetate buffer, pH<sub>read</sub> = 5 / 90% D2O |
| HDX time course (t1, t2, t3, t4, t5) | 4s (ice); 4s, 0.6 min and 6 min (23°C); 12 min (28°C) | 400s (ice); 400s, 1 h and 10 h (23°C); 20 h (28°C) |
| HDX control samples | Maximally-labelled control |  |
| # of peptides | 284 |  |
| Sequence coverage | 83.3% |  |
| Average peptide length / Redundancy | 10.15 / 5.57 |  |
| Replicates (technical) | 3 | 3 |
| Repeatability (average standard deviation) | 0.038981836 | 0.036375889 |

**Table S7.** Summary of the experimental details of the HDX comparison of the cleaved forms of Makona and Sudan GP and uncleaved Makona and Sudan GP.

| Data Set | Makona uncleaved GP | Makona 160-kDa GP (CatB-cleaved) |
| --- | --- | --- |
| HDX reaction details | Deuterated acetate buffer, pH<sub>read</sub> = 5 / 90% D2O |  |
| HDX time course (t1, t2, t3, t4, t5) | 400s (ice); 400s, 1 h and 10 h (23°C); 20 h (28°C) |  |
| # of peptides | 244 |  |
| Compared sequence coverage | 97.1% (excluding cleaved areas) |  |
| Average peptide length / Redundancy | 9.98 / 5.09 |  |
| Replicates (technical) | 3 | 3 |
| Repeatability (average standard deviation) | 0.060667015 | 0.06211289 |
| Significant differences in HDX (99% CI) | $\Delta\text{HDX} \geq 0.68$ | |
| Data Set | Makona uncleaved GP | Makona 164-kDa GP (CatL-cleaved) |
| HDX reaction details | Deuterated acetate buffer, pH<sub>read</sub> = 5 / 90% D2O |  |
| HDX time course (t1, t2, t3, t4, t5) | 400s (ice); 400s, 1 h and 10 h (23°C); 20 h (28°C) |  |
| # of peptides | 226 |  |
| Compared sequence coverage | 96.4% (excluding cleaved areas) |  |
| Average peptide length / Redundancy | 9.84 / 4.87 |  |
| Replicates (technical) | 3 | 3 |
| Repeatability (average standard deviation) | 0.060667015 | 0.045079013 |
| Significant differences in HDX (99% CI) | $\Delta\text{HDX} \geq 0.59$ | |
| Data Set | Makona uncleaved GP | Makona 120-kDa GPcl (CatB+L-cleaved) |
| HDX reaction details | Deuterated acetate buffer, pH<sub>read</sub> = 5 / 90% D2O |  |
| HDX time course (t1, t2, t3, t4, t5) | 400s (ice); 400s, 1 h and 10 h (23°C); 20 h (28°C) |  |
| # of peptides | 144 |  |
| Compared sequence coverage | 94.8% (excluding cleaved areas) |  |
| Average peptide length / Redundancy | 8.76 / 4.32 |  |
| Replicates (technical) | 3 | 3 |
| Repeatability (average standard deviation) | 0.060667015 | 0.044846413 |
| Significant differences in HDX (99% CI) | $\Delta\text{HDX} \geq 0.60$ | |
| Significant differences in HDX - cleavage | average uncleaved/160-kDa, uncleaved/164-kDa, uncleaved/120-kDa: $\Delta\text{HDX} \geq 0.62$ Da | |

| Data Set | Sudan uncleaved GP | Sudan 120-kDa GPcl (CatB+L-cleaved) |
| --- | --- | --- |
| HDX reaction details | Deuterated acetate buffer, pH<sub>read</sub> = 5 / 90% D2O |  |
| HDX time course (t1, t2, t3, t4, t5) | 400s (ice); 400s, 1 h and 10 h (23°C); 20 h (28°C) |  |
| # of compared peptides | 159 |  |
| Compared sequence coverage | 97.4% (excluding cleaved areas) |  |
| Average peptide length / Redundancy | 10.15 / 5.38 |  |
| Replicates (technical) | 3 | 3 |
| Repeatability (average standard deviation) | 0.036375889 | 0.03773772 |
| Significant differences in HDX (99% CI) | $\Delta\text{HDX} \geq 0.43$ Da --> **0.62 Da** to align to Makona GP cleavage analyses | |

**Table S8.** Summary of the experimental details of the HDX comparison across GPcl trimers.

| Data Set | Makona GPcl, pH 5 | Mayinga GPcl, pH 5 |
| --- | --- | --- |
| HDX reaction details | Deuterated acetate buffer, pH<sub>read</sub> = 5 / 90% D2O |  |
| HDX time course (t1, t2, t3, t4, t5) | 400s (ice); 400s, 1 h and 10 h (23°C); 20 h (28°C) |  |
| # of compared peptides | 147 |  |
| Compared sequence coverage | 94.8% |  |
| Average peptide length / Redundancy | 9.95 / 5.02 |  |
| Replicates (technical) | 3 | 3 |
| Repeatability (average standard deviation) | 0.038894453 | 0.036814172 |
| Significant differences in HDX (99% CI) | $\Delta\text{HDX} \geq 0.43$ | |

| Data Set | Sudan GPcl, pH 5 | Mayinga GPcl, pH 5 |
| --- | --- | --- |
| HDX reaction details | Deuterated acetate buffer, pH<sub>read</sub> = 5 / 90% D2O |  |
| HDX time course (t1, t2, t3, t4, t5) | 400s (ice); 400s, 1 h and 10 h (23°C); 20 h (28°C) |  |
| # of compared peptides | 85 |  |
| Compared sequence coverage | 78.5% |  |
| Average peptide length / Redundancy | 9.75 / 3.44 |  |
| Replicates (technical) | 3 | 3 |
| Repeatability (average standard deviation) | 0.03773772 | 0.036814172 |
| Significant differences in HDX (99% CI) | $\Delta\text{HDX} \geq 0.44$ | |
| **Significant differences in HDX - GPcl** | average Makona/Mayinga and Sudan/Mayinga GPcl thresholds: $\Delta\text{HDX} \geq 0.43$ Da | |

**Table S9.** Summary of the experimental details of the HDX comparison between apo GPcl and NPC1 bound GPcl.

| Data Set | Mayinga GPcl, pH 5 | Mayinga GPcl + NPC1, pH 5 |
| --- | --- | --- |
| HDX reaction details | Deuterated acetate buffer, pH<sub>read</sub> = 5 / 90% D2O |  |
| HDX time course (t1, t2, t3, t4, t5) | 400s (ice); 400s, 1 h and 10 h (23°C); 20 h (28°C) |  |
| # of peptides | 146 |  |
| Compared sequence coverage | 94.8% (excluding cleaved areas) |  |
| Average peptide length / Redundancy | 10.09 / 5.06 |  |
| Replicates (technical) | 3 | 3 |
| Repeatability (average standard deviation) | 0.036814172 | 0.03635922 |
| Significant differences in HDX (99% CI) | $\Delta\text{HDX} \geq 0.42$ | |

| Data Set | Makona GPcl, pH 5 | Makona GPcl + NPC1, pH 5 |
| --- | --- | --- |
| HDX reaction details | Deuterated acetate buffer, pH<sub>read</sub> = 5 / 90% D2O |  |
| HDX time course (t1, t2, t3, t4, t5) | 400s (ice); 400s, 1 h and 10 h (23°C); 20 h (28°C) |  |
| # of peptides | 151 |  |
| Compared sequence coverage | 94.8% (excluding cleaved areas) |  |
| Average peptide length / Redundancy | 9.99 / 5.19 |  |
| Replicates (technical) | 3 | 3 |
| Repeatability (average standard deviation) | 0.038894453 | 0.042470522 |
| Significant differences in HDX (99% CI) | $\Delta\text{HDX} \geq 0.45$ | |

| Data Set | Sudan GPcl, pH 5 | Sudan GPcl + NPC1, pH 5 |
| --- | --- | --- |
| HDX reaction details | Deuterated acetate buffer, pH<sub>read</sub> = 5 / 90% D2O |  |
| HDX time course (t1, t2, t3, t4, t5) | 400s (ice); 400s, 1 h and 10 h (23°C); 20 h (28°C) |  |
| # of peptides | 160 |  |
| Compared sequence coverage | 97.5% (excluding cleaved areas) |  |
| Average peptide length / Redundancy | 10.17 / 5.44 |  |
| Replicates (technical) | 3 | 3 |
| Repeatability (average standard deviation) | 0.03773772 | 0.054172447 |
| Significant differences in HDX (99% CI) | $\Delta\text{HDX} \geq 0.53$ | |

| Data Set | Sudan 140-kDa GP, pH 5 | Sudan 140-kDa GP + NPC1, pH 5 |
| --- | --- | --- |
| HDX reaction details | Deuterated acetate buffer, pH<sub>read</sub> = 5 / 90% D2O |  |
| HDX time course (t1, t2, t3, t4, t5) | 400s (ice); 400s, 1 h and 10 h (23°C); 20 h (28°C) |  |
| # of peptides | 215 |  |
| Compared sequence coverage | 84.4% (excluding cleaved areas) |  |
| Average peptide length / Redundancy | 9.75 / 5.53 |  |
| Replicates (technical) | 3 | 3 |
| Repeatability (average standard deviation) | 0.037190034 | 0.03807275 |
| Significant differences in HDX (99% CI) | $\Delta\text{HDX} \geq 0.42$ | |
| **Significant differences in HDX - +/- NPC1** | average +/- NPC1 thresholds: $\Delta\text{HDX} \geq 0.45$ Da | |

**Table S10.**
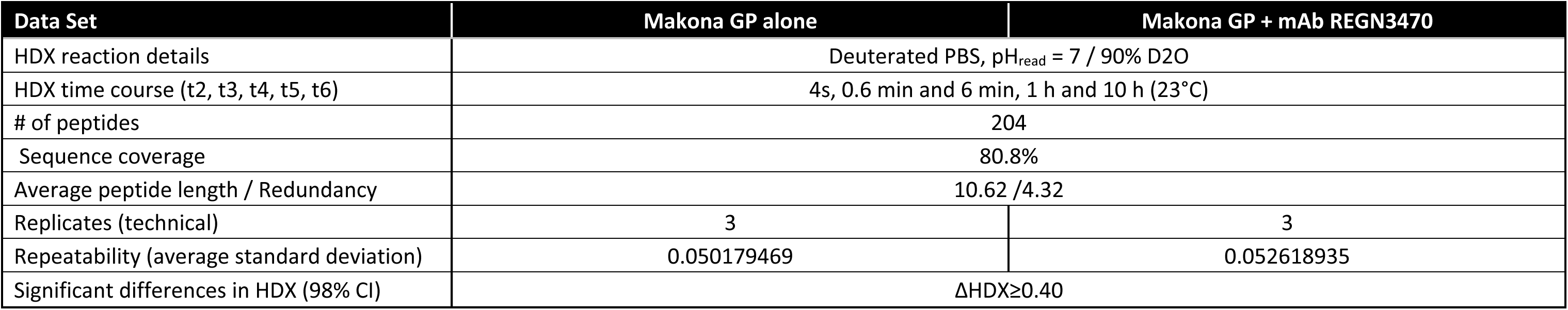
Summary of the experimental details of the HDX comparison between apo Makona GP and mAb REGN3470 bound Makona

| Data Set | Makona GP alone | Makona GP + mAb REGN3470 |
| --- | --- | --- |
| HDX reaction details | Deuterated PBS, pH<sub>read</sub> = 7 / 90% D2O |  |
| HDX time course (t2, t3, t4, t5, t6) | 4s, 0.6 min and 6 min, 1 h and 10 h (23°C) |  |
| # of peptides | 204 |  |
| Sequence coverage | 80.8% |  |
| Average peptide length / Redundancy | 10.62 / 4.32 |  |
| Replicates (technical) | 3 | 3 |
| Repeatability (average standard deviation) | 0.050179469 | 0.052618935 |
| Significant differences in HDX (98% CI) | $\Delta\text{HDX} \geq 0.40$ | |

