## Supplementary Dataset for "Allosteric remodelling of the Ebola virus glycoprotein underlies differences in entry mechanisms across species"

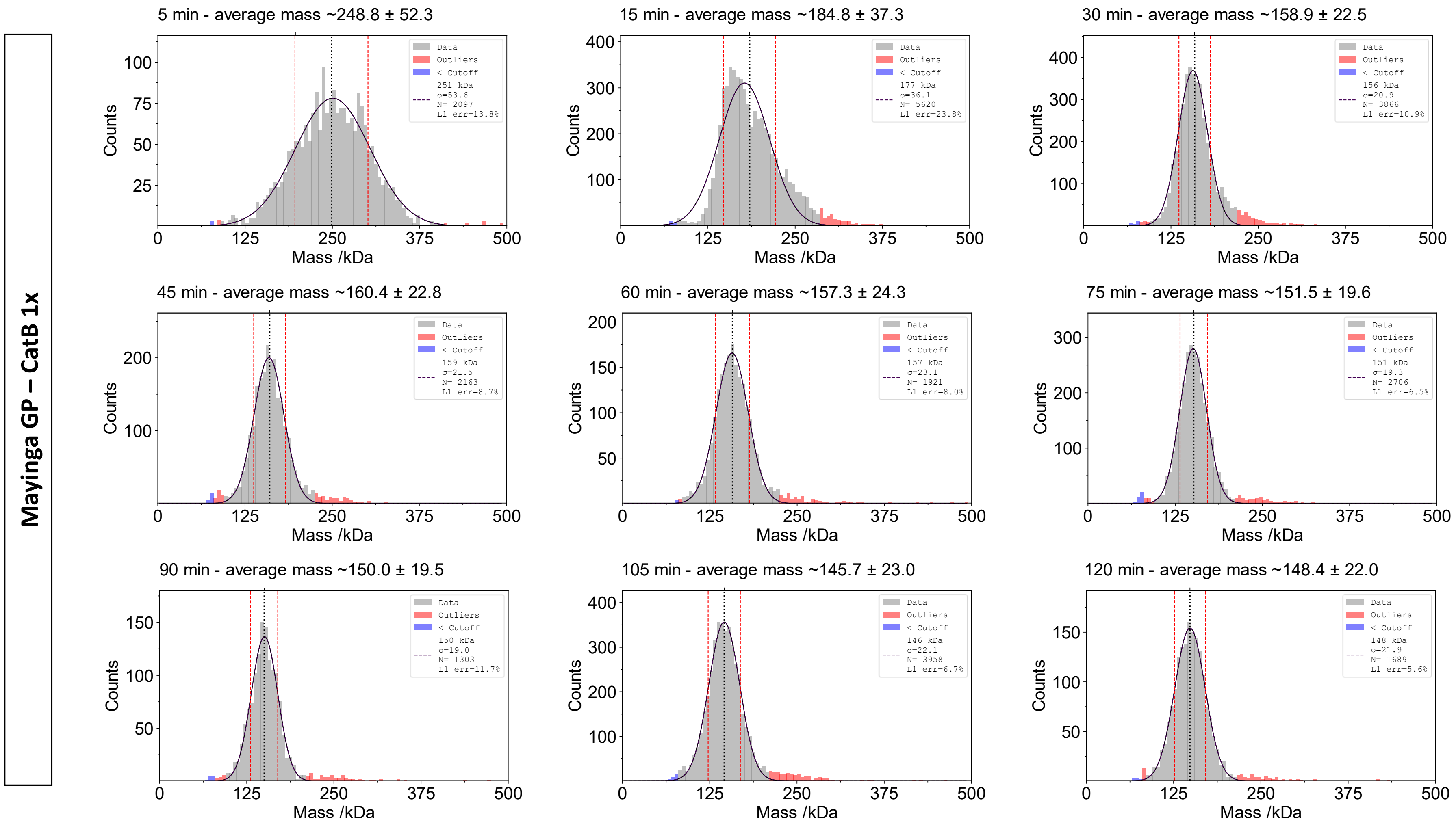
Mayinga GP – CatB 1x
5 min - average mass ~248.8 ± 52.3
Data
Outliers
< Cutoff
251 kDa
σ=53.6
N= 2097
L1 err=13.8%
15 min - average mass ~184.8 ± 37.3
177 kDa
σ=36.1
N= 5620
L1 err=23.8%
30 min - average mass ~158.9 ± 22.5
156 kDa
σ=20.9
N= 3866
L1 err=10.9%
45 min - average mass ~160.4 ± 22.8
159 kDa
σ=21.5
N= 2163
L1 err=8.7%
60 min - average mass ~157.3 ± 24.3
157 kDa
σ=23.1
N= 1921
L1 err=8.0%
75 min - average mass ~151.5 ± 19.6
151 kDa
σ=19.3
N= 2706
L1 err=6.5%
90 min - average mass ~150.0 ± 19.5
150 kDa
σ=19.0
N= 1303
L1 err=11.7%
105 min - average mass ~145.7 ± 23.0
146 kDa
σ=22.1
N= 3958
L1 err=6.7%
120 min - average mass ~148.4 ± 22.0
148 kDa
σ=21.9
N= 1689
L1 err=5.6%
Counts
Mass /kDa

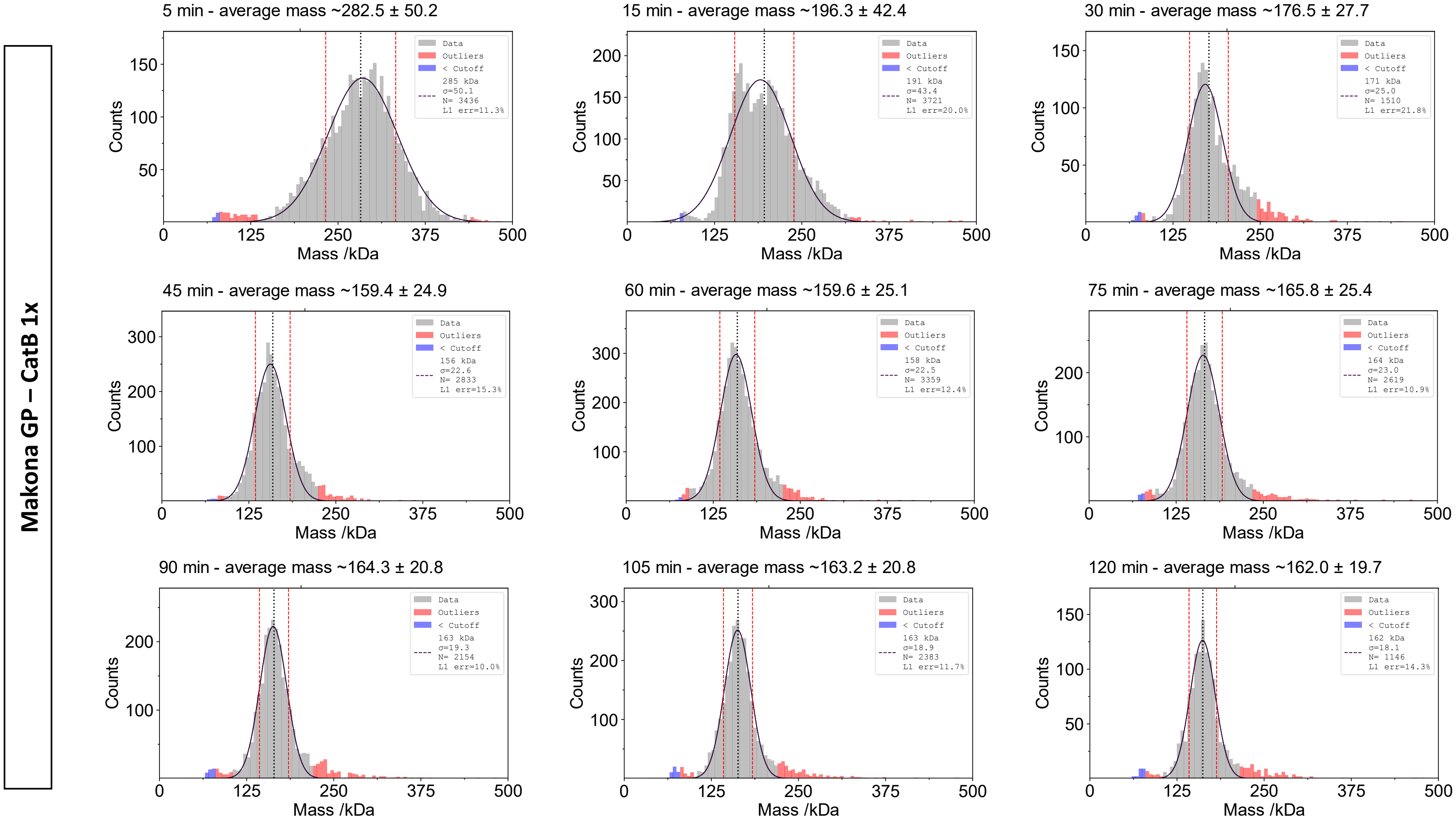
Makona GP – CatB 1x
5 min - average mass ~282.5 ± 50.2
15 min - average mass ~196.3 ± 42.4
30 min - average mass ~176.5 ± 27.7
45 min - average mass ~159.4 ± 24.9
60 min - average mass ~159.6 ± 25.1
75 min - average mass ~165.8 ± 25.4
90 min - average mass ~164.3 ± 20.8
105 min - average mass ~163.2 ± 20.8
120 min - average mass ~162.0 ± 19.7
Data
Outliers
< Cutoff
285 kDa σ=50.1 N= 3436 L1 err=11.3%
191 kDa σ=43.4 N= 3721 L1 err=20.0%
171 kDa σ=25.0 N= 1510 L1 err=21.8%
156 kDa σ=22.6 N= 2833 L1 err=15.3%
158 kDa σ=22.5 N= 3359 L1 err=12.4%
164 kDa σ=23.0 N= 2619 L1 err=10.9%
163 kDa σ=19.3 N= 2154 L1 err=10.0%
163 kDa σ=18.9 N= 2383 L1 err=11.7%
162 kDa σ=18.1 N= 1146 L1 err=14.3%
Counts
Mass /kDa
0
125
250
375
500

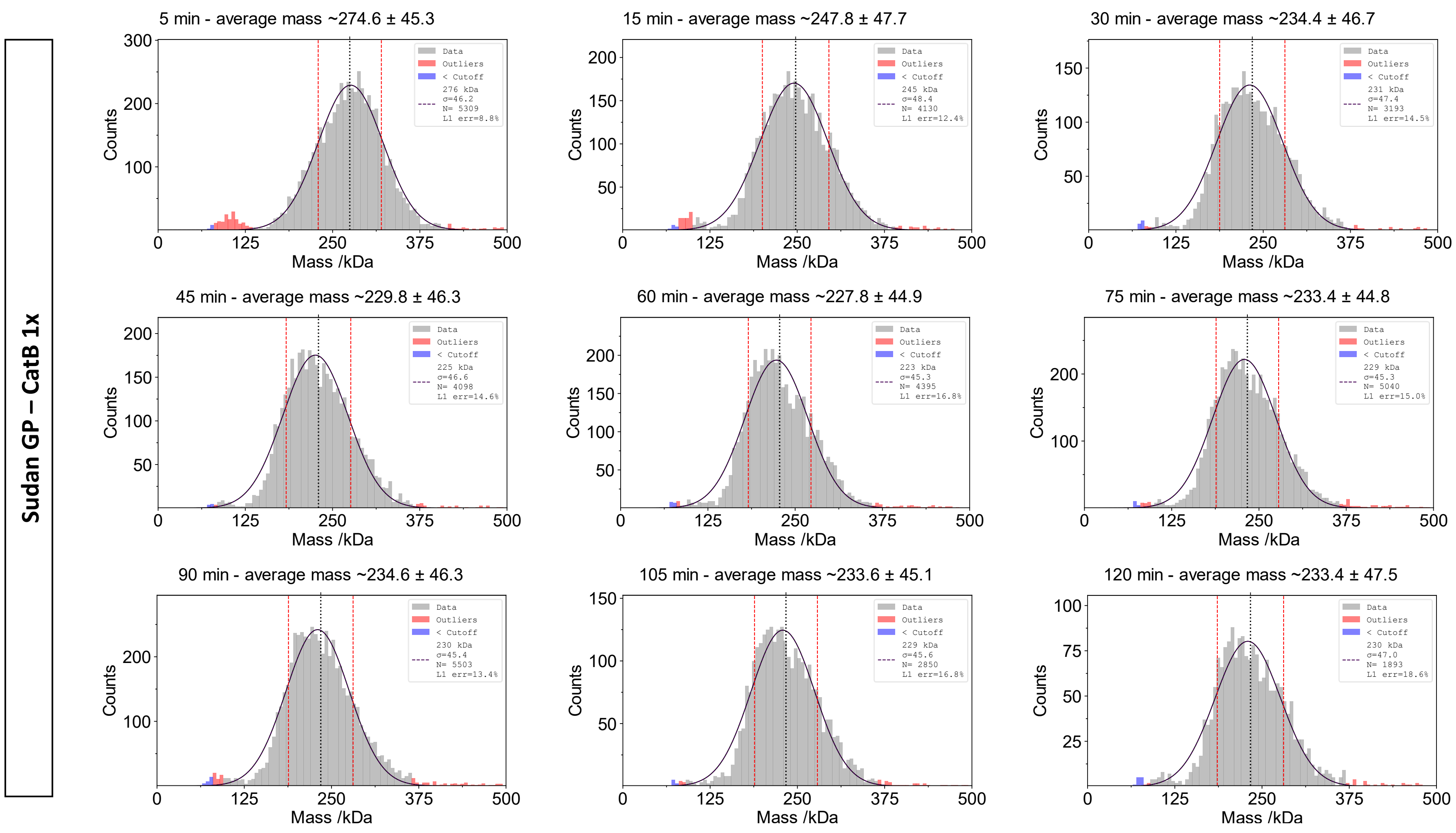
Sudan GP – CatB 1x
5 min - average mass ~274.6 ± 45.3
276 kDa σ=46.2 N= 5309 L1 err=8.8%
15 min - average mass ~247.8 ± 47.7
245 kDa σ=48.4 N= 4130 L1 err=12.4%
30 min - average mass ~234.4 ± 46.7
231 kDa σ=47.4 N= 3193 L1 err=14.5%
45 min - average mass ~229.8 ± 46.3
225 kDa σ=46.6 N= 4098 L1 err=14.6%
60 min - average mass ~227.8 ± 44.9
223 kDa σ=45.3 N= 4395 L1 err=16.8%
75 min - average mass ~233.4 ± 44.8
229 kDa σ=45.3 N= 5040 L1 err=15.0%
90 min - average mass ~234.6 ± 46.3
230 kDa σ=45.4 N= 5503 L1 err=13.4%
105 min - average mass ~233.6 ± 45.1
229 kDa σ=45.6 N= 2850 L1 err=16.8%
120 min - average mass ~233.4 ± 47.5
230 kDa σ=47.0 N= 1893 L1 err=18.6%
Data
Outliers
< Cutoff
Counts
Mass /kDa

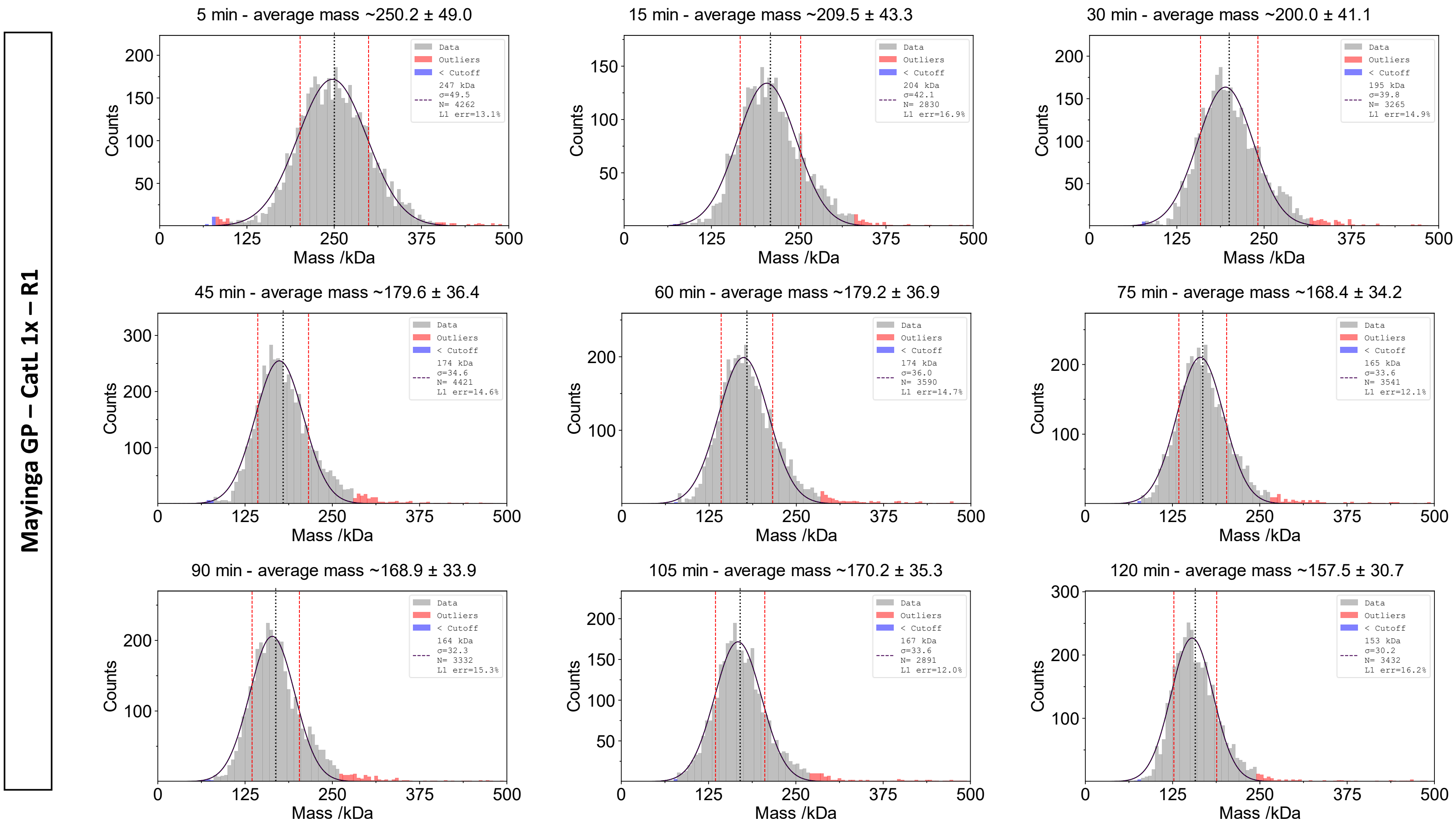

Mayinga GP – CatL 1x – R1
5 min - average mass ~250.2 ± 49.0
Data
Outliers
< Cutoff
247 kDa
σ=49.5
N= 4262
L1 err=13.1%
15 min - average mass ~209.5 ± 43.3
204 kDa
σ=42.1
N= 2830
L1 err=16.9%
30 min - average mass ~200.0 ± 41.1
195 kDa
σ=39.8
N= 3265
L1 err=14.9%
45 min - average mass ~179.6 ± 36.4
174 kDa
σ=34.6
N= 4421
L1 err=14.6%
60 min - average mass ~179.2 ± 36.9
174 kDa
σ=36.0
N= 3590
L1 err=14.7%
75 min - average mass ~168.4 ± 34.2
165 kDa
σ=33.6
N= 3541
L1 err=12.1%
90 min - average mass ~168.9 ± 33.9
164 kDa
σ=32.3
N= 3332
L1 err=15.3%
105 min - average mass ~170.2 ± 35.3
167 kDa
σ=33.6
N= 2891
L1 err=12.0%
120 min - average mass ~157.5 ± 30.7
153 kDa
σ=30.2
N= 3432
L1 err=16.2%
Counts
Mass /kDa

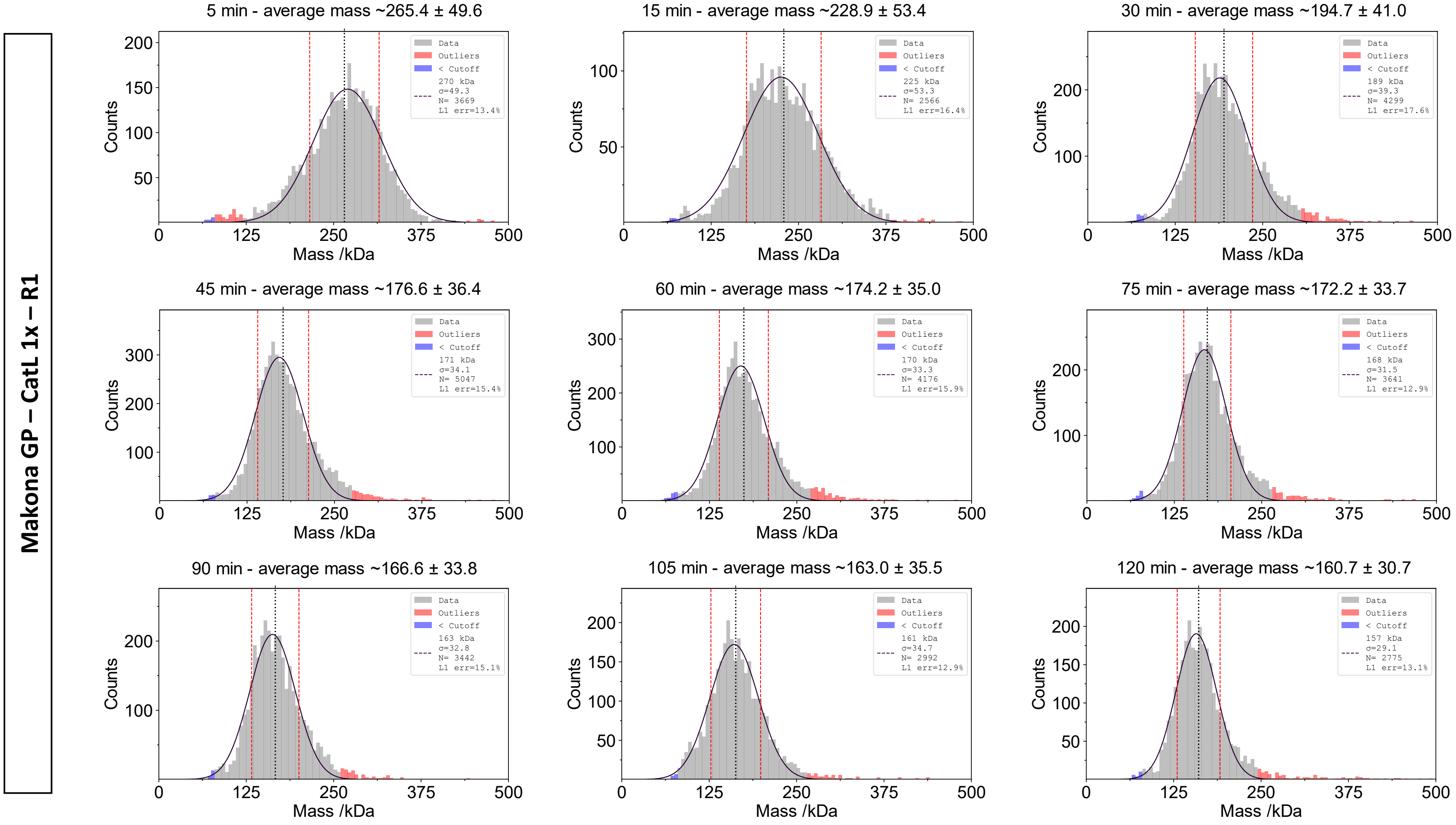
Makona GP – CatL 1x – R1
5 min - average mass ~265.4 ± 49.6
270 kDa
σ=49.3
N= 3669
L1 err=13.4%
15 min - average mass ~228.9 ± 53.4
225 kDa
σ=53.3
N= 2566
L1 err=16.4%
30 min - average mass ~194.7 ± 41.0
189 kDa
σ=39.3
N= 4299
L1 err=17.6%
45 min - average mass ~176.6 ± 36.4
171 kDa
σ=34.1
N= 5047
L1 err=15.4%
60 min - average mass ~174.2 ± 35.0
170 kDa
σ=33.3
N= 4176
L1 err=15.9%
75 min - average mass ~172.2 ± 33.7
168 kDa
σ=31.5
N= 3641
L1 err=12.9%
90 min - average mass ~166.6 ± 33.8
163 kDa
σ=32.8
N= 3442
L1 err=15.1%
105 min - average mass ~163.0 ± 35.5
161 kDa
σ=34.7
N= 2992
L1 err=12.9%
120 min - average mass ~160.7 ± 30.7
157 kDa
σ=29.1
N= 2775
L1 err=13.1%
Data
Outliers
< Cutoff
Counts
Mass /kDa

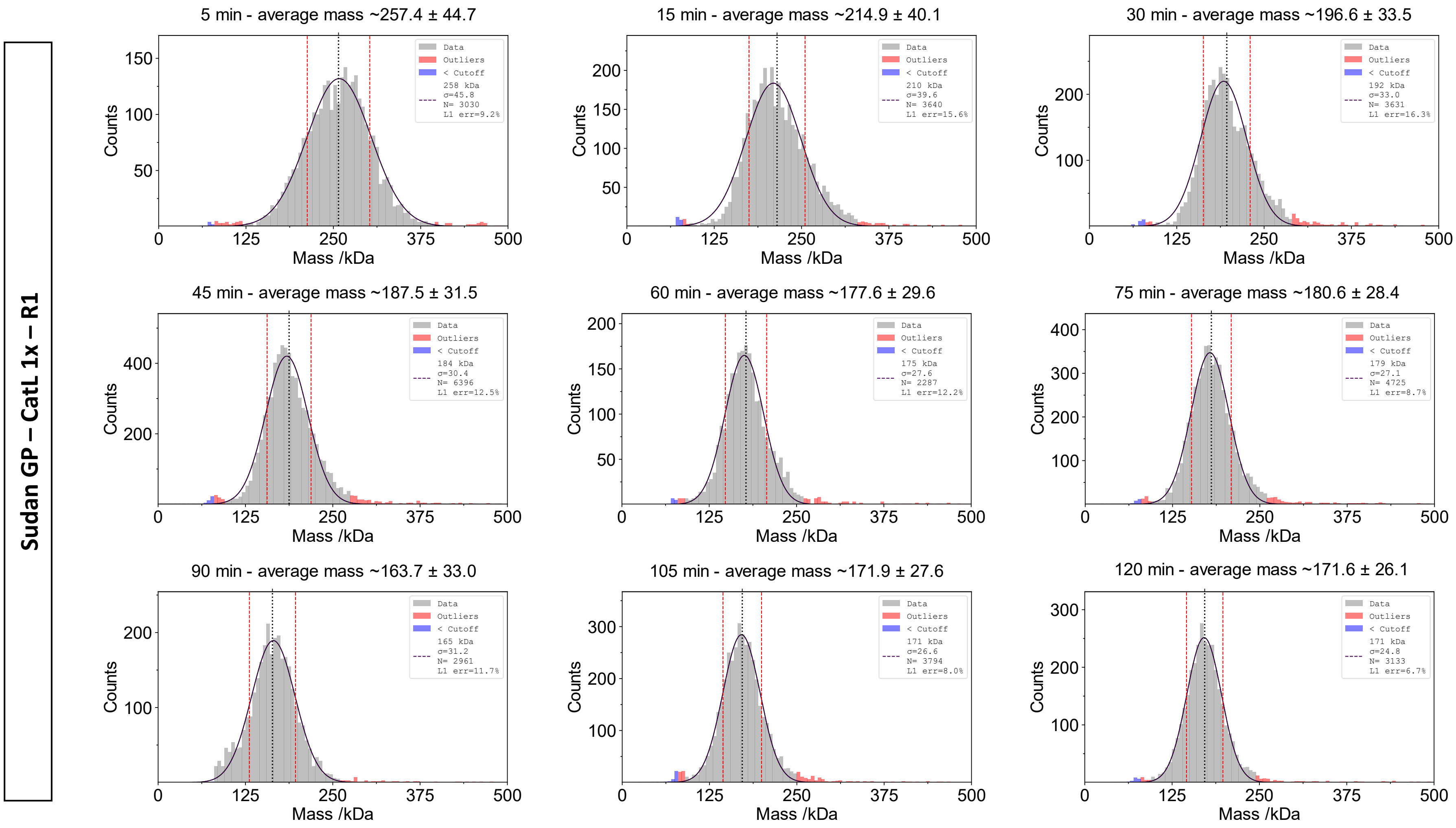
Sudan GP – CatL 1x – R1
5 min - average mass ~257.4 ± 44.7
Data
Outliers
< Cutoff
258 kDa
σ=45.8
N= 3030
L1 err=9.2%
15 min - average mass ~214.9 ± 40.1
Data
Outliers
< Cutoff
210 kDa
σ=39.6
N= 3640
L1 err=15.6%
30 min - average mass ~196.6 ± 33.5
Data
Outliers
< Cutoff
192 kDa
σ=33.0
N= 3631
L1 err=16.3%
45 min - average mass ~187.5 ± 31.5
Data
Outliers
< Cutoff
184 kDa
σ=30.4
N= 6396
L1 err=12.5%
60 min - average mass ~177.6 ± 29.6
Data
Outliers
< Cutoff
175 kDa
σ=27.6
N= 2287
L1 err=12.2%
75 min - average mass ~180.6 ± 28.4
Data
Outliers
< Cutoff
179 kDa
σ=27.1
N= 4725
L1 err=8.7%
90 min - average mass ~163.7 ± 33.0
Data
Outliers
< Cutoff
165 kDa
σ=31.2
N= 2961
L1 err=11.7%
105 min - average mass ~171.9 ± 27.6
Data
Outliers
< Cutoff
171 kDa
σ=26.6
N= 3794
L1 err=8.0%
120 min - average mass ~171.6 ± 26.1
Data
Outliers
< Cutoff
171 kDa
σ=24.8
N= 3133
L1 err=6.7%
Counts
Mass /kDa
0
125
250
375
500

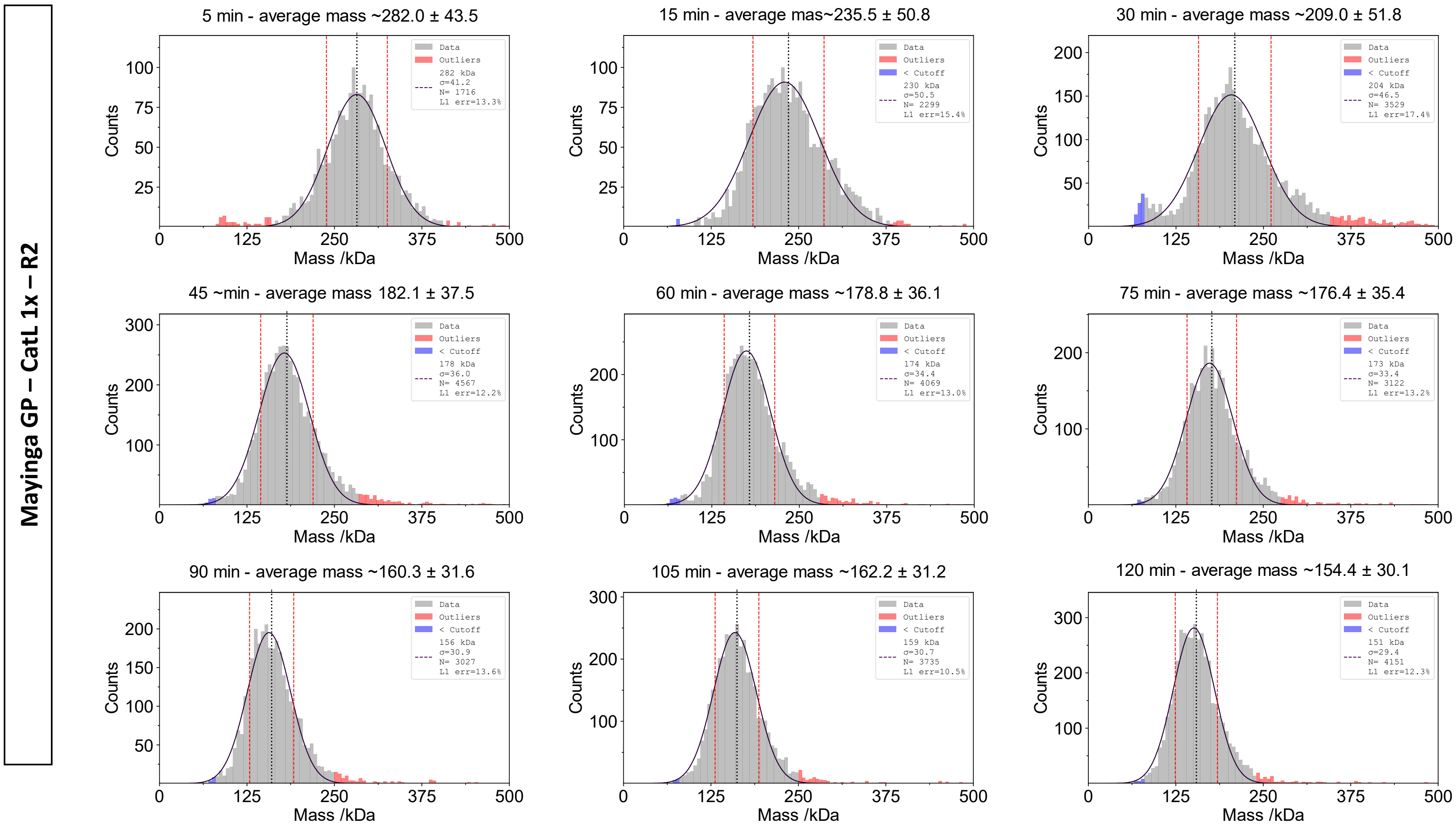
Mayinga GP – CatL 1x – R2
5 min - average mass ~282.0 ± 43.5
Data
Outliers
282 kDa σ=41.2 N= 1716 L1 err=13.3%
15 min - average mas~235.5 ± 50.8
Data
Outliers
< Cutoff
230 kDa σ=50.5 N= 2299 L1 err=15.4%
30 min - average mass ~209.0 ± 51.8
Data
Outliers
< Cutoff
204 kDa σ=46.5 N= 3529 L1 err=17.4%
45 ~min - average mass 182.1 ± 37.5
Data
Outliers
< Cutoff
178 kDa σ=36.0 N= 4567 L1 err=12.2%
60 min - average mass ~178.8 ± 36.1
Data
Outliers
< Cutoff
174 kDa σ=34.4 N= 4069 L1 err=13.0%
75 min - average mass ~176.4 ± 35.4
Data
Outliers
< Cutoff
173 kDa σ=33.4 N= 3122 L1 err=13.2%
90 min - average mass ~160.3 ± 31.6
Data
Outliers
< Cutoff
156 kDa σ=30.9 N= 3027 L1 err=13.6%
105 min - average mass ~162.2 ± 31.2
Data
Outliers
< Cutoff
159 kDa σ=30.7 N= 3735 L1 err=10.5%
120 min - average mass ~154.4 ± 30.1
Data
Outliers
< Cutoff
151 kDa σ=29.4 N= 4151 L1 err=12.3%
Counts
Mass /kDa

**Makona GP – CatL 1x – R2**

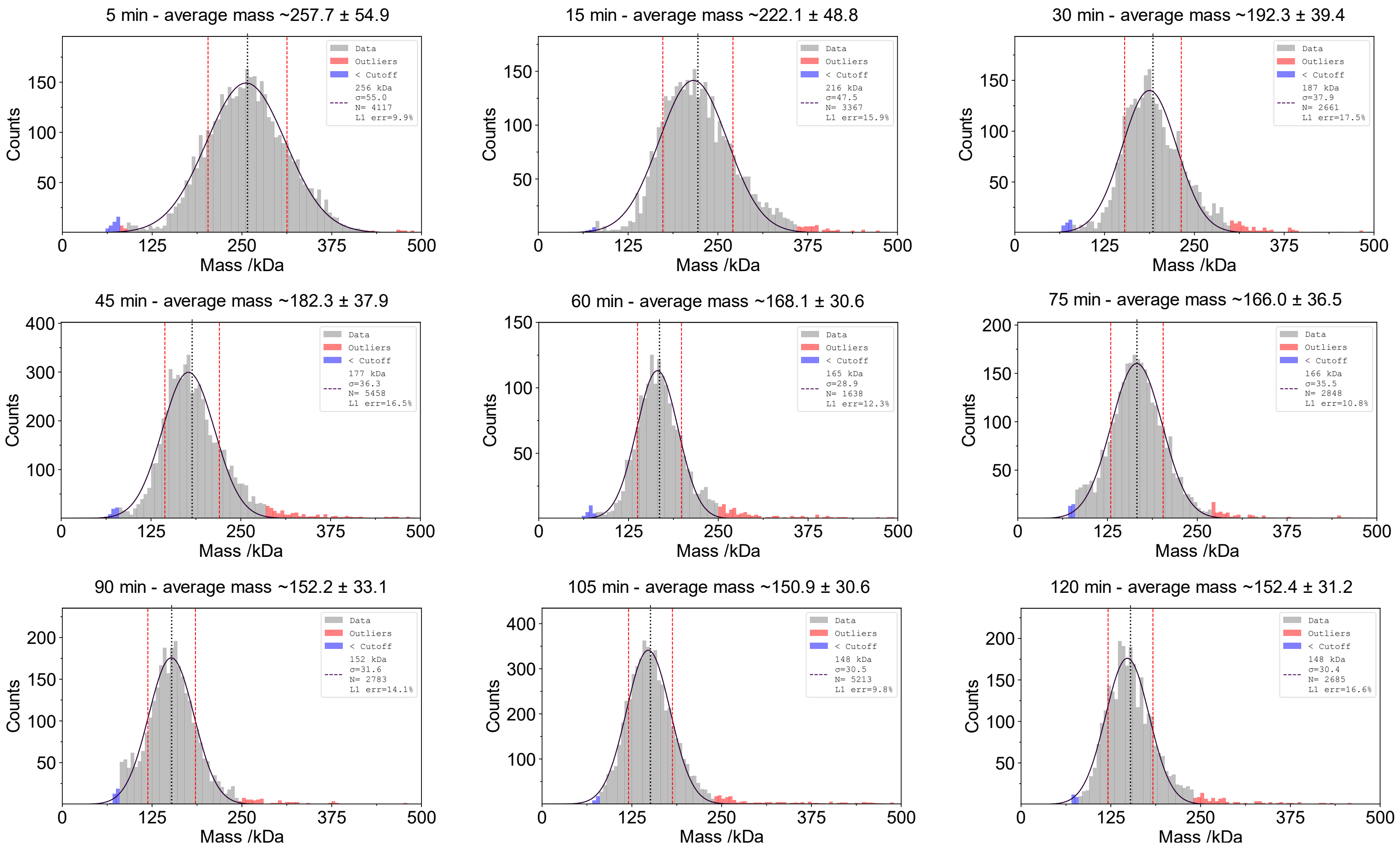

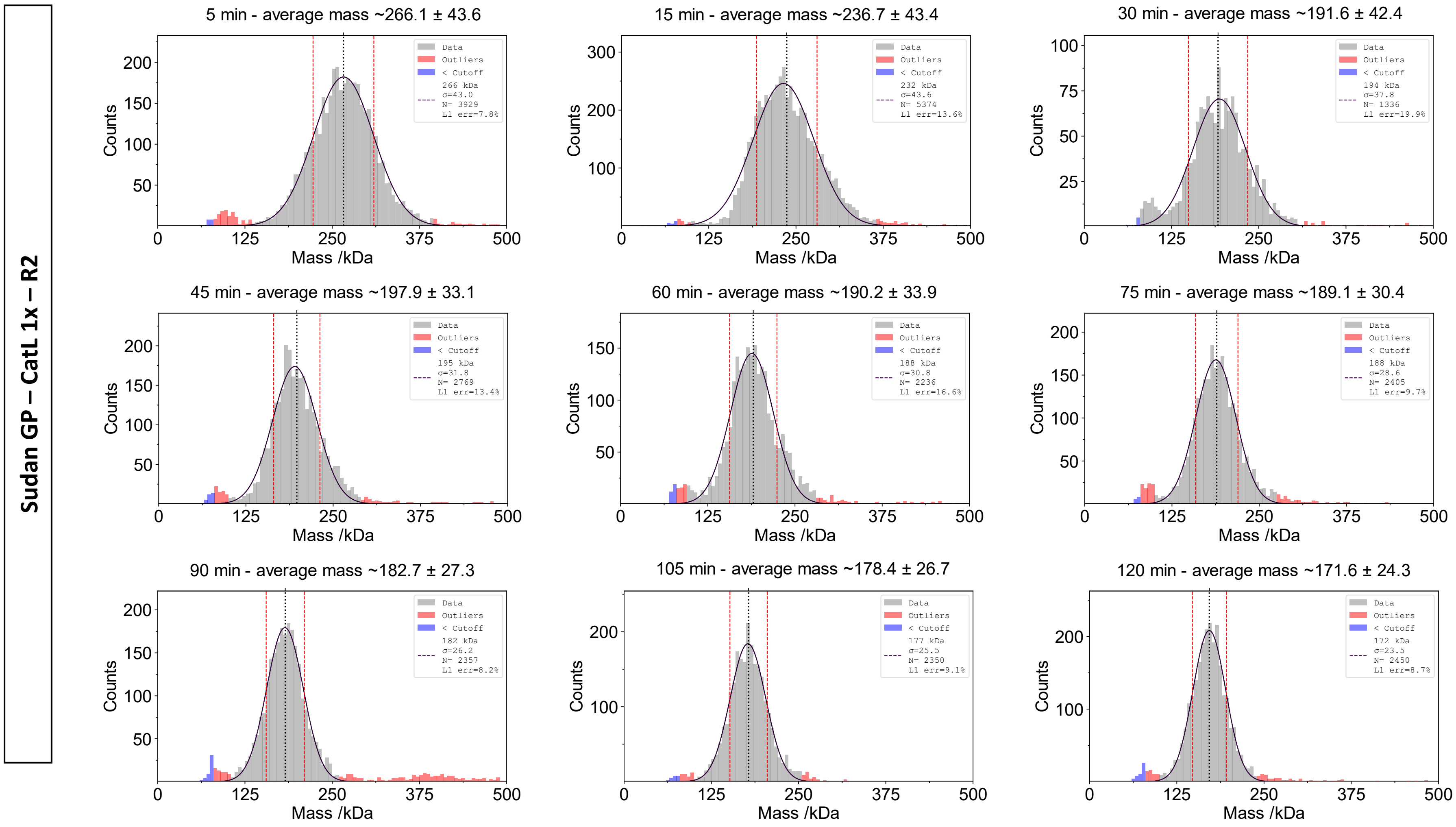

Sudan GP – CatL 1x – R2
5 min - average mass ~266.1 ± 43.6
Data
Outliers
< Cutoff
266 kDa σ=43.0 N= 3929 L1 err=7.8%
15 min - average mass ~236.7 ± 43.4
Data
Outliers
< Cutoff
232 kDa σ=43.6 N= 5374 L1 err=13.6%
30 min - average mass ~191.6 ± 42.4
Data
Outliers
< Cutoff
194 kDa σ=37.8 N= 1336 L1 err=19.9%
45 min - average mass ~197.9 ± 33.1
Data
Outliers
< Cutoff
195 kDa σ=31.8 N= 2769 L1 err=13.4%
60 min - average mass ~190.2 ± 33.9
Data
Outliers
< Cutoff
188 kDa σ=30.8 N= 2236 L1 err=16.6%
75 min - average mass ~189.1 ± 30.4
Data
Outliers
< Cutoff
188 kDa σ=28.6 N= 2405 L1 err=9.7%
90 min - average mass ~182.7 ± 27.3
Data
Outliers
< Cutoff
182 kDa σ=26.2 N= 2357 L1 err=8.2%
105 min - average mass ~178.4 ± 26.7
Data
Outliers
< Cutoff
177 kDa σ=25.5 N= 2350 L1 err=9.1%
120 min - average mass ~171.6 ± 24.3
Data
Outliers
< Cutoff
172 kDa σ=23.5 N= 2450 L1 err=8.7%
Counts
Mass /kDa

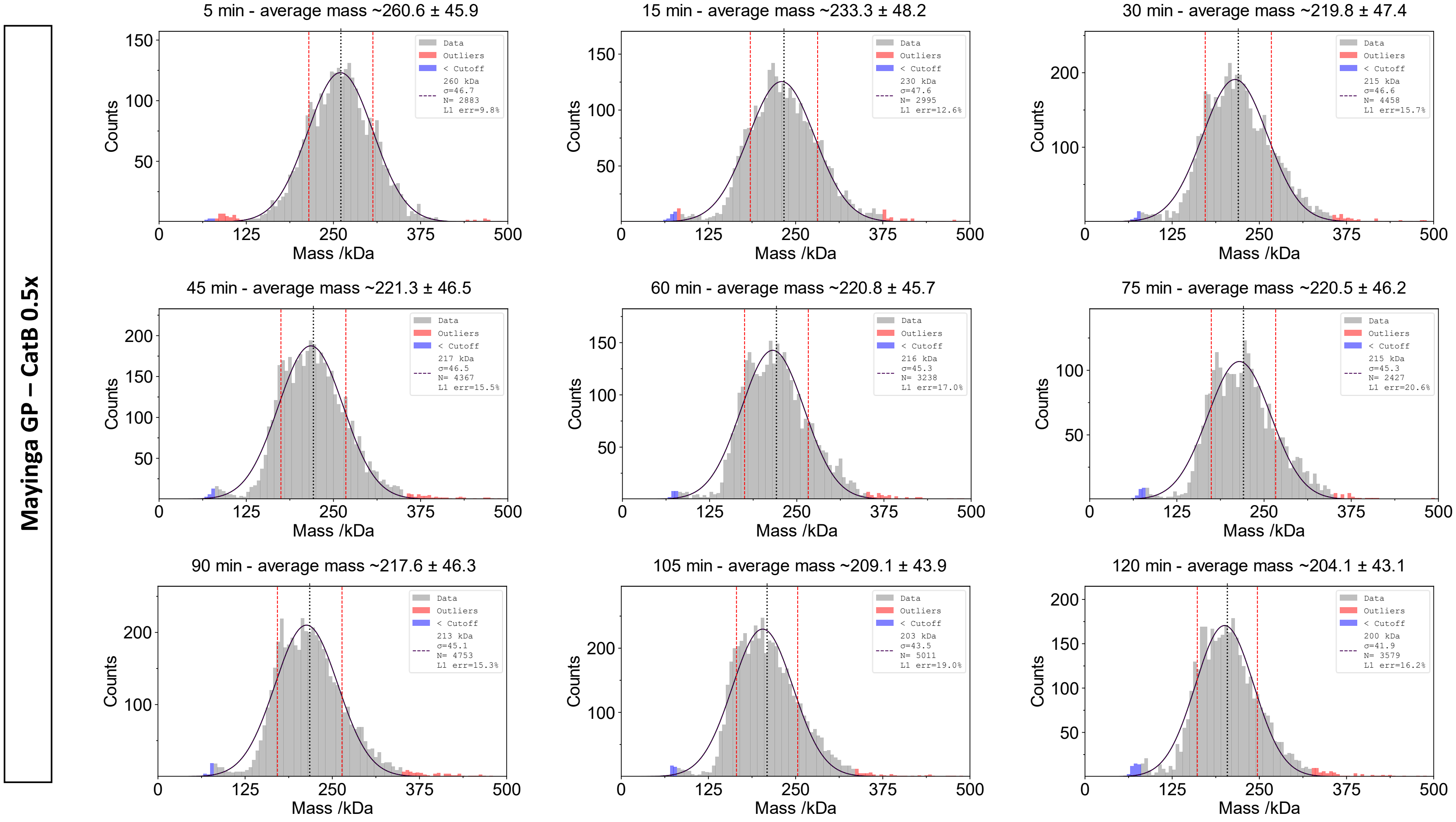
Mayinga GP – CatB 0.5x
5 min - average mass ~260.6 ± 45.9
15 min - average mass ~233.3 ± 48.2
30 min - average mass ~219.8 ± 47.4
45 min - average mass ~221.3 ± 46.5
60 min - average mass ~220.8 ± 45.7
75 min - average mass ~220.5 ± 46.2
90 min - average mass ~217.6 ± 46.3
105 min - average mass ~209.1 ± 43.9
120 min - average mass ~204.1 ± 43.1
Counts
Mass /kDa
Data
Outliers
< Cutoff
260 kDa σ=46.7 N= 2883 L1 err=9.8%
230 kDa σ=47.6 N= 2995 L1 err=12.6%
215 kDa σ=46.6 N= 4458 L1 err=15.7%
217 kDa σ=46.5 N= 4367 L1 err=15.5%
216 kDa σ=45.3 N= 3238 L1 err=17.0%
215 kDa σ=45.3 N= 2427 L1 err=20.6%
213 kDa σ=45.1 N= 4753 L1 err=15.3%
203 kDa σ=43.5 N= 5011 L1 err=19.0%
200 kDa σ=41.9 N= 3579 L1 err=16.2%

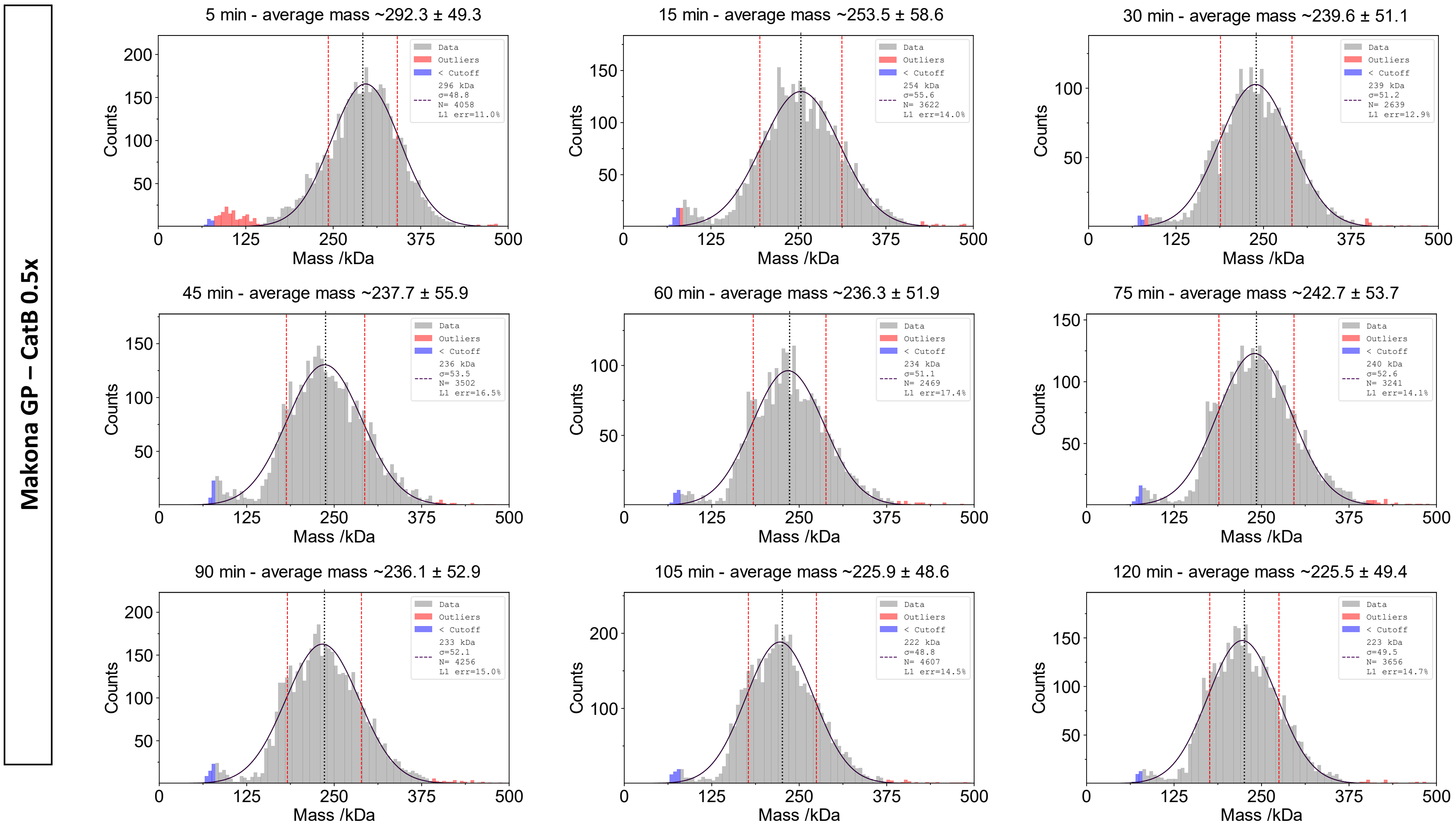
Makona GP – CatB 0.5x
5 min - average mass ~292.3 ± 49.3
Data
Outliers
< Cutoff
296 kDa
σ=48.8
N= 4058
L1 err=11.0%
15 min - average mass ~253.5 ± 58.6
254 kDa
σ=55.6
N= 3622
L1 err=14.0%
30 min - average mass ~239.6 ± 51.1
239 kDa
σ=51.2
N= 2639
L1 err=12.9%
45 min - average mass ~237.7 ± 55.9
236 kDa
σ=53.5
N= 3502
L1 err=16.5%
60 min - average mass ~236.3 ± 51.9
234 kDa
σ=51.1
N= 2469
L1 err=17.4%
75 min - average mass ~242.7 ± 53.7
240 kDa
σ=52.6
N= 3241
L1 err=14.1%
90 min - average mass ~236.1 ± 52.9
233 kDa
σ=52.1
N= 4256
L1 err=15.0%
105 min - average mass ~225.9 ± 48.6
222 kDa
σ=48.8
N= 4607
L1 err=14.5%
120 min - average mass ~225.5 ± 49.4
223 kDa
σ=49.5
N= 3656
L1 err=14.7%
Counts
Mass /kDa
0
125
250
375
500

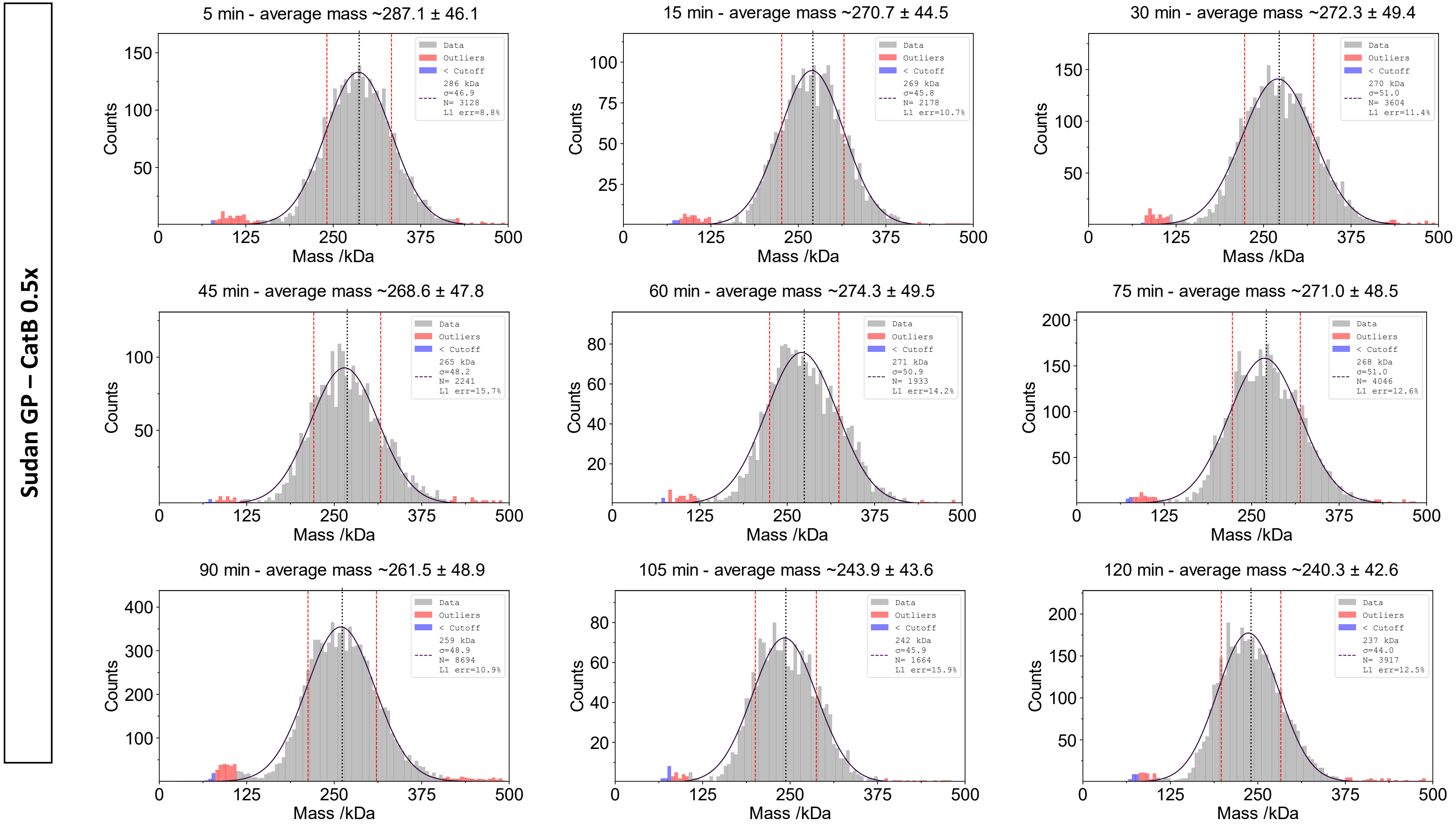

Sudan GP – CatB 0.5x
5 min - average mass ~287.1 ± 46.1
Data
Outliers
< Cutoff
286 kDa
σ=46.9
N= 3128
L1 err=8.8%
15 min - average mass ~270.7 ± 44.5
Data
Outliers
< Cutoff
269 kDa
σ=45.8
N= 2178
L1 err=10.7%
30 min - average mass ~272.3 ± 49.4
Data
Outliers
< Cutoff
270 kDa
σ=51.0
N= 3604
L1 err=11.4%
45 min - average mass ~268.6 ± 47.8
Data
Outliers
< Cutoff
265 kDa
σ=48.2
N= 2241
L1 err=15.7%
60 min - average mass ~274.3 ± 49.5
Data
Outliers
< Cutoff
271 kDa
σ=50.9
N= 1933
L1 err=14.2%
75 min - average mass ~271.0 ± 48.5
Data
Outliers
< Cutoff
268 kDa
σ=51.0
N= 4046
L1 err=12.6%
90 min - average mass ~261.5 ± 48.9
Data
Outliers
< Cutoff
259 kDa
σ=48.9
N= 8694
L1 err=10.9%
105 min - average mass ~243.9 ± 43.6
Data
Outliers
< Cutoff
242 kDa
σ=45.9
N= 1664
L1 err=15.9%
120 min - average mass ~240.3 ± 42.6
Data
Outliers
< Cutoff
237 kDa
σ=44.0
N= 3917
L1 err=12.5%
Counts
Mass /kDa
0
125
250
375
500

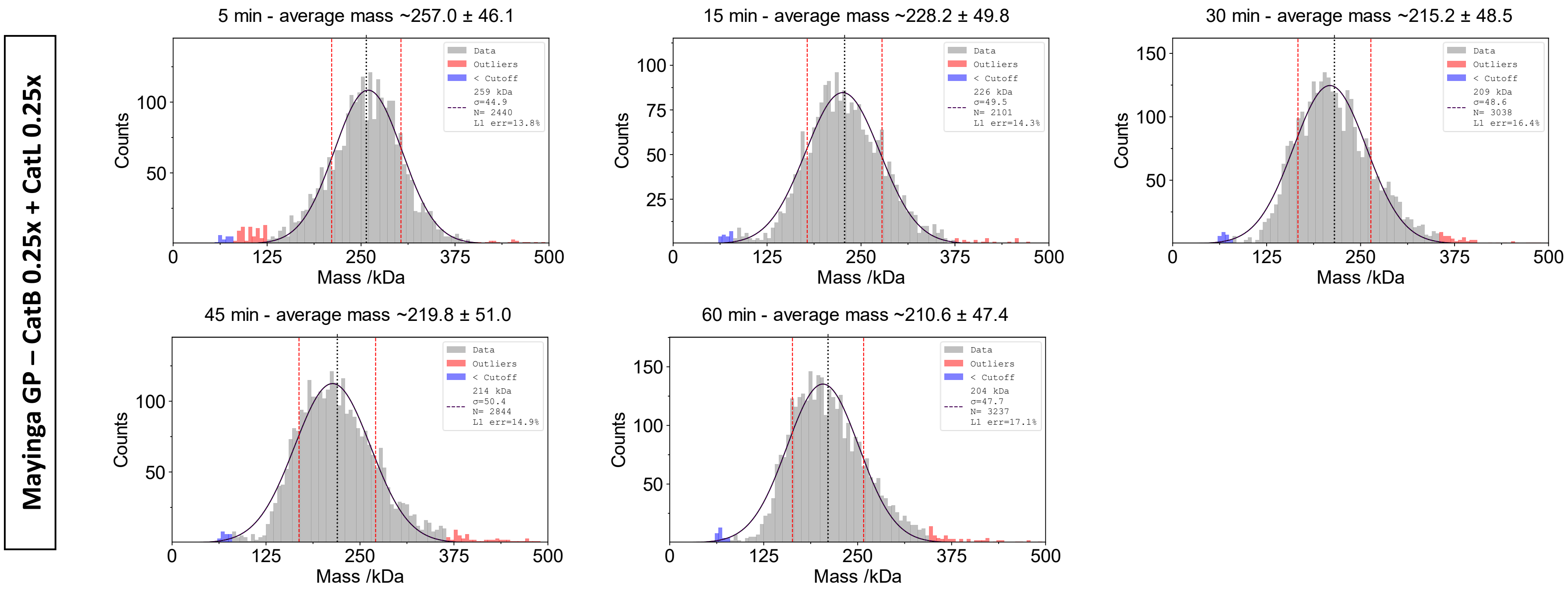
Mayinga GP – CatB 0.25x + CatL 0.25x
5 min - average mass ~257.0 ± 46.1
Data
Outliers
< Cutoff
259 kDa
σ=44.9
N= 2440
L1 err=13.8%
15 min - average mass ~228.2 ± 49.8
Data
Outliers
< Cutoff
226 kDa
σ=49.5
N= 2101
L1 err=14.3%
30 min - average mass ~215.2 ± 48.5
Data
Outliers
< Cutoff
209 kDa
σ=48.6
N= 3038
L1 err=16.4%
45 min - average mass ~219.8 ± 51.0
Data
Outliers
< Cutoff
214 kDa
σ=50.4
N= 2844
L1 err=14.9%
60 min - average mass ~210.6 ± 47.4
Data
Outliers
< Cutoff
204 kDa
σ=47.7
N= 3237
L1 err=17.1%
Counts
Mass /kDa

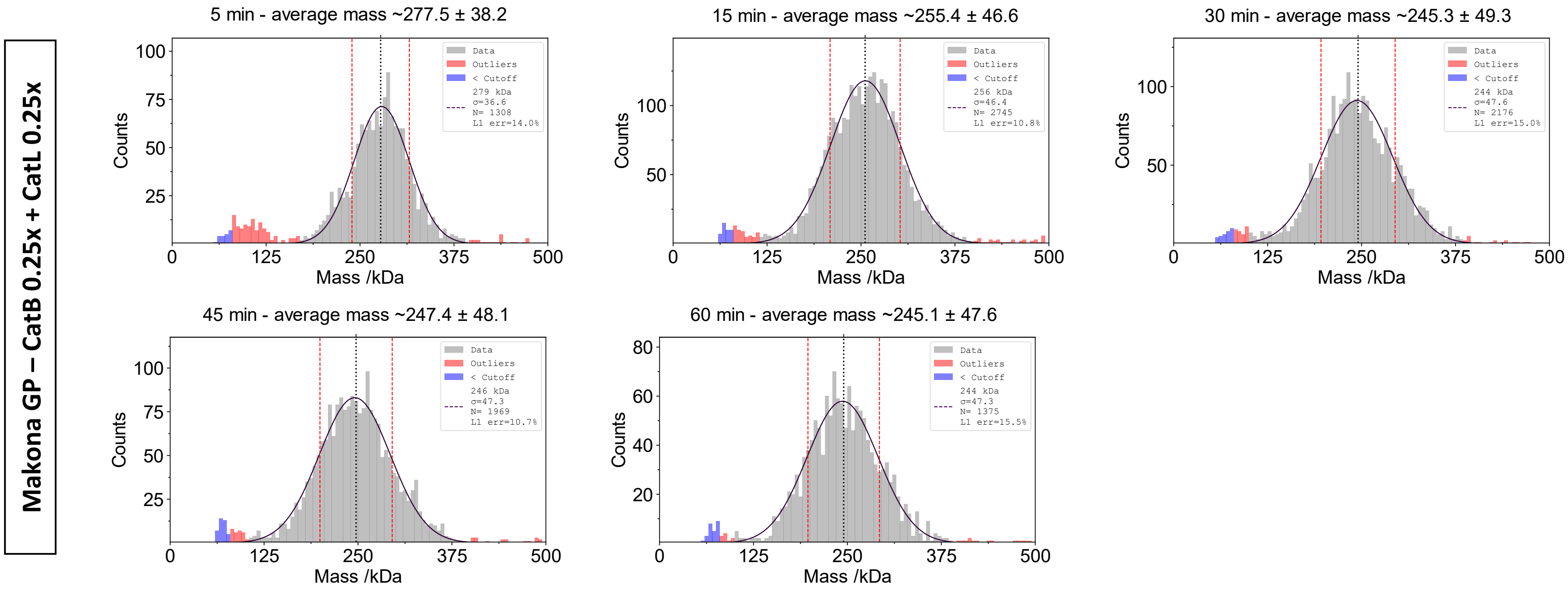
Makona GP – CatB 0.25x + CatL 0.25x
5 min - average mass ~277.5 ± 38.2
Data
Outliers
< Cutoff
279 kDa
σ=36.6
N= 1308
L1 err=14.0%
15 min - average mass ~255.4 ± 46.6
Data
Outliers
< Cutoff
256 kDa
σ=46.4
N= 2745
L1 err=10.8%
30 min - average mass ~245.3 ± 49.3
Data
Outliers
< Cutoff
244 kDa
σ=47.6
N= 2176
L1 err=15.0%
45 min - average mass ~247.4 ± 48.1
Data
Outliers
< Cutoff
246 kDa
σ=47.3
N= 1969
L1 err=10.7%
60 min - average mass ~245.1 ± 47.6
Data
Outliers
< Cutoff
244 kDa
σ=47.3
N= 1375
L1 err=15.5%
Counts
Mass /kDa

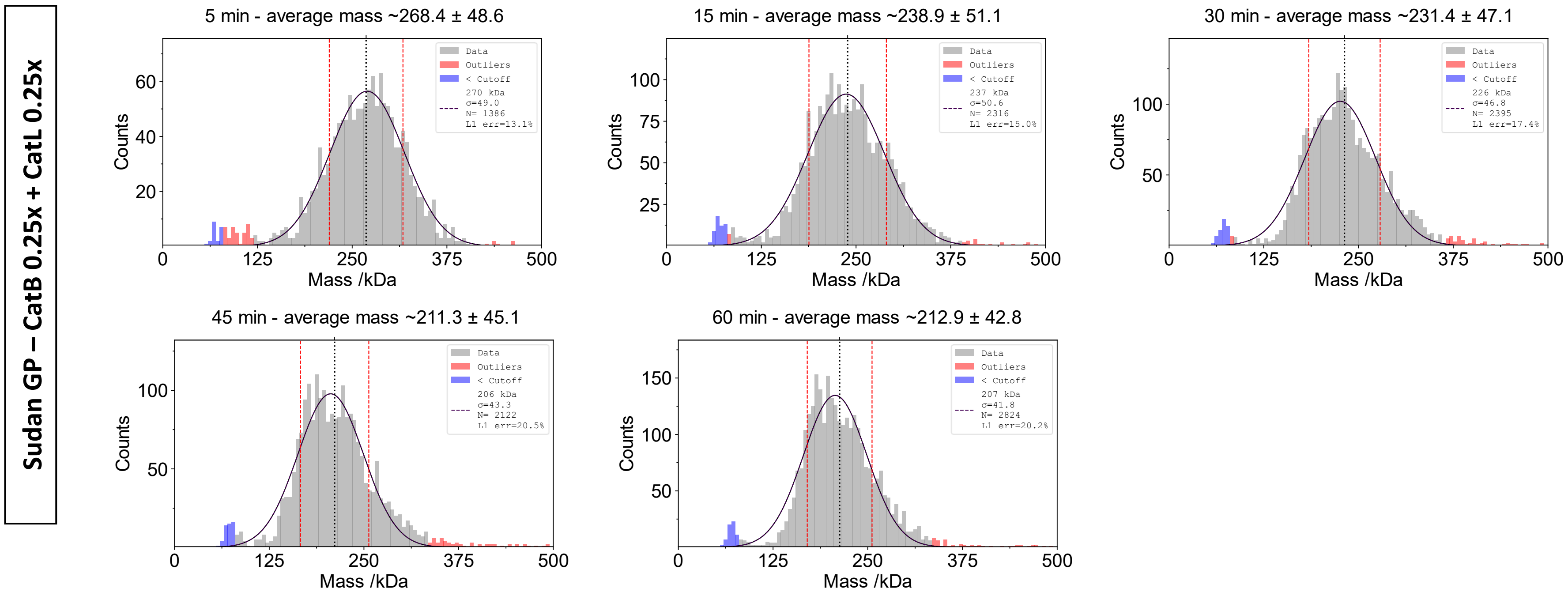
Sudan GP – CatB 0.25x + CatL 0.25x
5 min - average mass ~268.4 ± 48.6
Data
Outliers
< Cutoff
270 kDa
σ=49.0
N= 1386
L1 err=13.1%
Counts
Mass /kDa
15 min - average mass ~238.9 ± 51.1
237 kDa
σ=50.6
N= 2316
L1 err=15.0%
30 min - average mass ~231.4 ± 47.1
226 kDa
σ=46.8
N= 2395
L1 err=17.4%
45 min - average mass ~211.3 ± 45.1
206 kDa
σ=43.3
N= 2122
L1 err=20.5%
60 min - average mass ~212.9 ± 42.8
207 kDa
σ=41.8
N= 2824
L1 err=20.2%

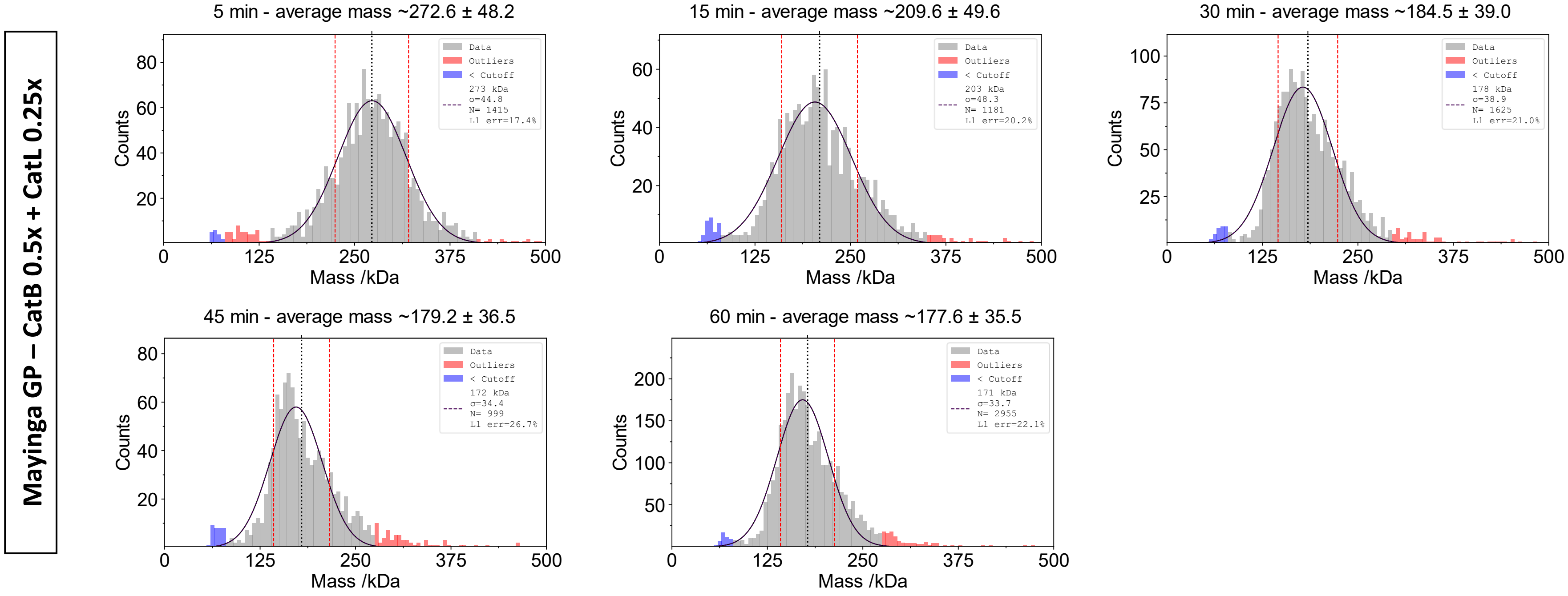
Mayinga GP – CatB 0.5x + CatL 0.25x
5 min - average mass ~272.6 ± 48.2
Data
Outliers
< Cutoff
273 kDa
σ=44.8
N= 1415
L1 err=17.4%
15 min - average mass ~209.6 ± 49.6
Data
Outliers
< Cutoff
203 kDa
σ=48.3
N= 1181
L1 err=20.2%
30 min - average mass ~184.5 ± 39.0
Data
Outliers
< Cutoff
178 kDa
σ=38.9
N= 1625
L1 err=21.0%
45 min - average mass ~179.2 ± 36.5
Data
Outliers
< Cutoff
172 kDa
σ=34.4
N= 999
L1 err=26.7%
60 min - average mass ~177.6 ± 35.5
Data
Outliers
< Cutoff
171 kDa
σ=33.7
N= 2955
L1 err=22.1%
Counts
Mass /kDa

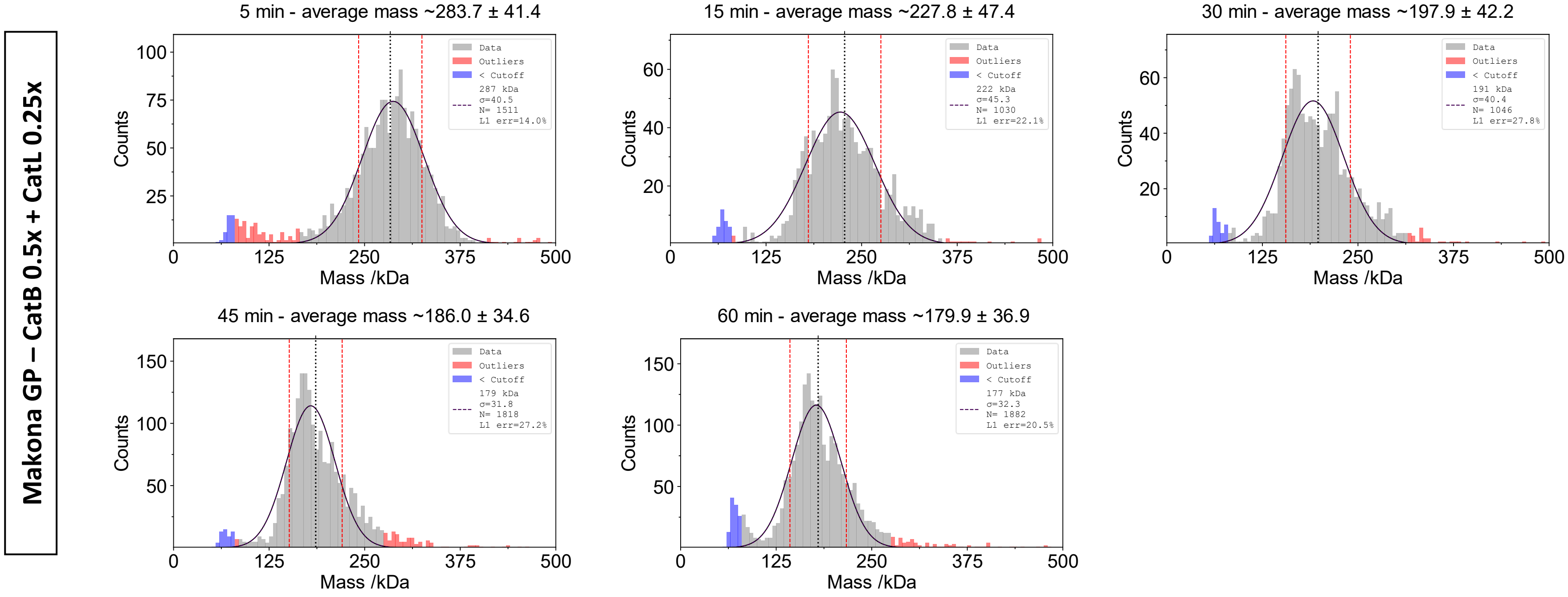

Makona GP – CatB 0.5x + CatL 0.25x
5 min - average mass ~283.7 ± 41.4
Data
Outliers
< Cutoff
287 kDa
σ=40.5
N= 1511
L1 err=14.0%
15 min - average mass ~227.8 ± 47.4
Data
Outliers
< Cutoff
222 kDa
σ=45.3
N= 1030
L1 err=22.1%
30 min - average mass ~197.9 ± 42.2
Data
Outliers
< Cutoff
191 kDa
σ=40.4
N= 1046
L1 err=27.8%
45 min - average mass ~186.0 ± 34.6
Data
Outliers
< Cutoff
179 kDa
σ=31.8
N= 1818
L1 err=27.2%
60 min - average mass ~179.9 ± 36.9
Data
Outliers
< Cutoff
177 kDa
σ=32.3
N= 1882
L1 err=20.5%
Counts
Mass /kDa
0
125
250
375
500

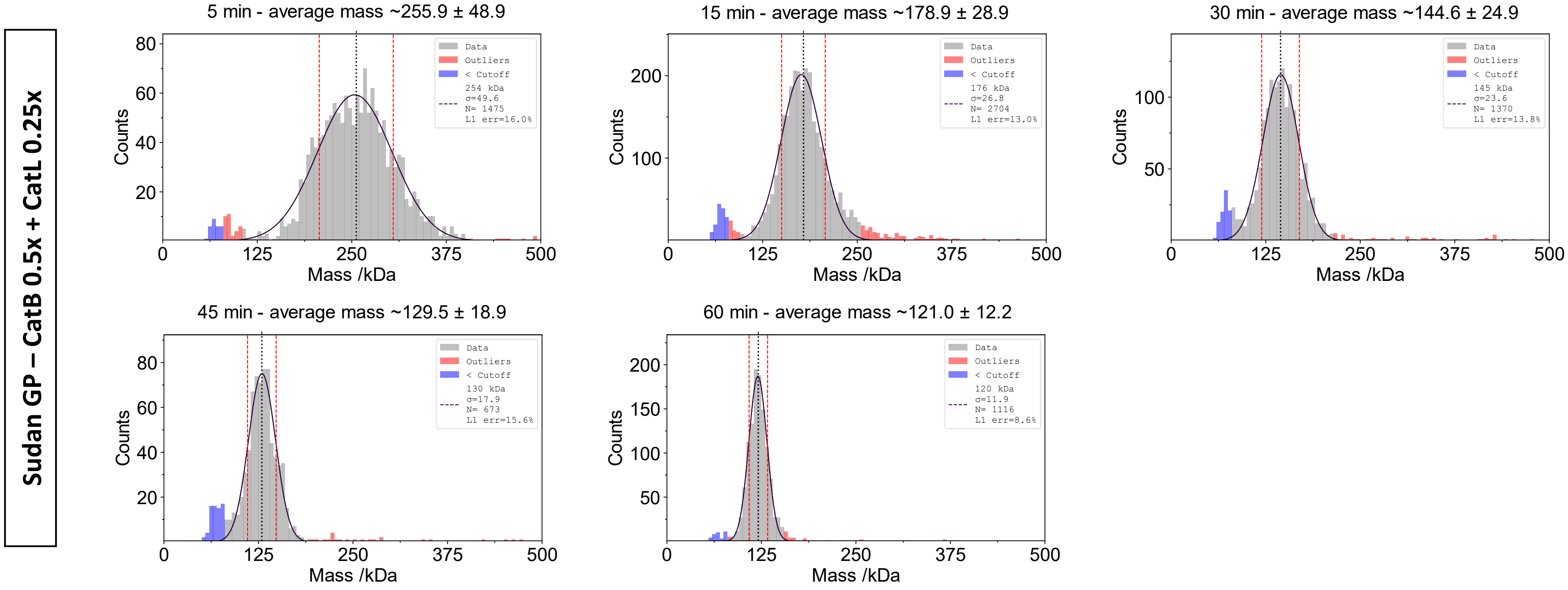

Sudan GP – CatB 0.5x + CatL 0.25x
5 min - average mass ~255.9 ± 48.9
Data
Outliers
< Cutoff
254 kDa
σ=49.6
N= 1475
L1 err=16.0%
15 min - average mass ~178.9 ± 28.9
176 kDa
σ=26.8
N= 2704
L1 err=13.0%
30 min - average mass ~144.6 ± 24.9
145 kDa
σ=23.6
N= 1370
L1 err=13.8%
45 min - average mass ~129.5 ± 18.9
130 kDa
σ=17.9
N= 673
L1 err=15.6%
60 min - average mass ~121.0 ± 12.2
120 kDa
σ=11.9
N= 1116
L1 err=8.6%
Counts
Mass /kDa
0
125
250
375
500

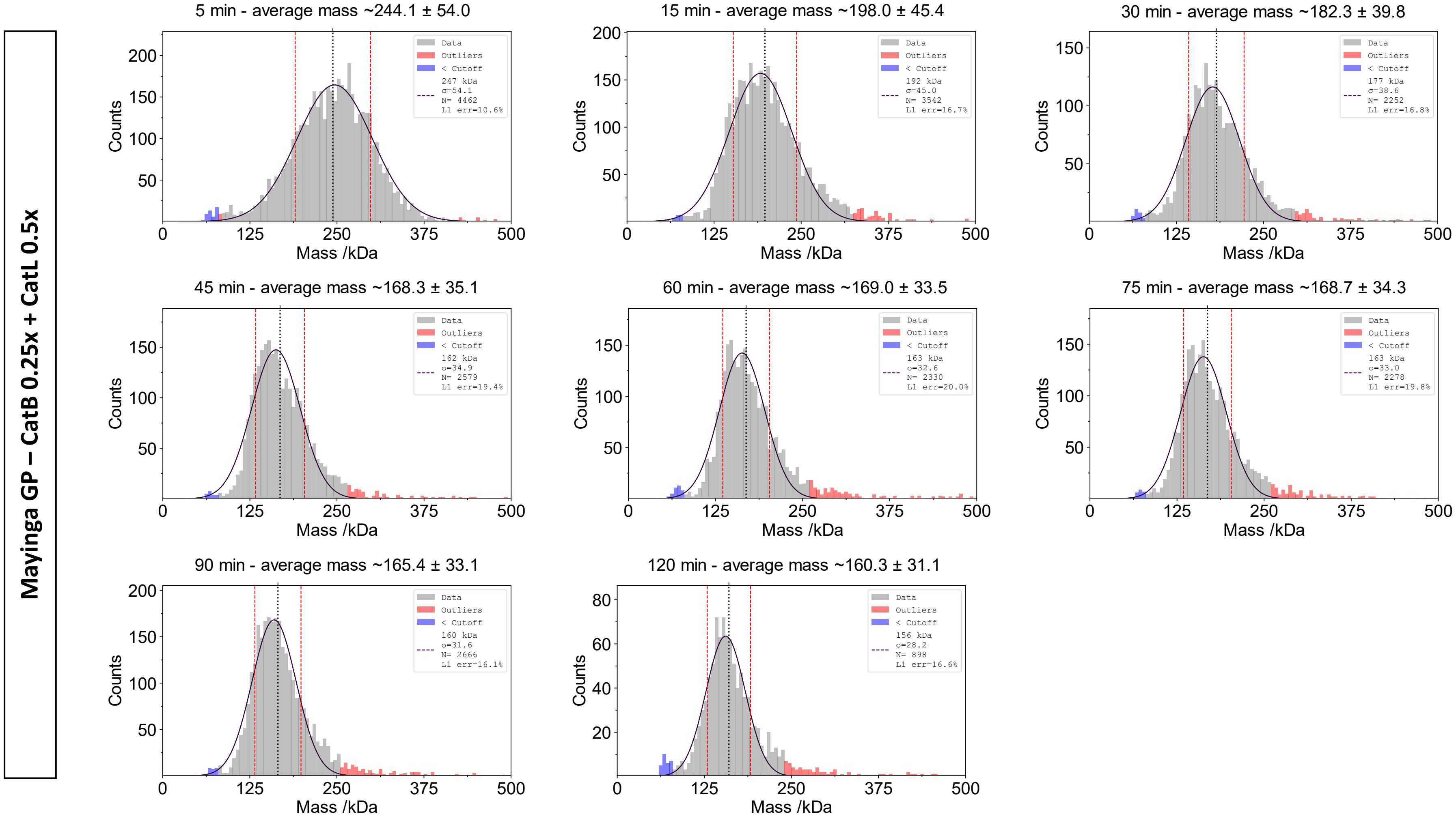
Mayinga GP – CatB 0.25x + CatL 0.5x
5 min - average mass ~244.1 ± 54.0
247 kDa σ=54.1 N= 4462 L1 err=10.6%
15 min - average mass ~198.0 ± 45.4
192 kDa σ=45.0 N= 3542 L1 err=16.7%
30 min - average mass ~182.3 ± 39.8
177 kDa σ=38.6 N= 2252 L1 err=16.8%
45 min - average mass ~168.3 ± 35.1
162 kDa σ=34.9 N= 2579 L1 err=19.4%
60 min - average mass ~169.0 ± 33.5
163 kDa σ=32.6 N= 2330 L1 err=20.0%
75 min - average mass ~168.7 ± 34.3
163 kDa σ=33.0 N= 2278 L1 err=19.8%
90 min - average mass ~165.4 ± 33.1
160 kDa σ=31.6 N= 2666 L1 err=16.1%
120 min - average mass ~160.3 ± 31.1
156 kDa σ=28.2 N= 898 L1 err=16.6%
Data
Outliers
< Cutoff
Counts
Mass /kDa

**Makona GP – CatB 0.25x + CatL 0.5x**

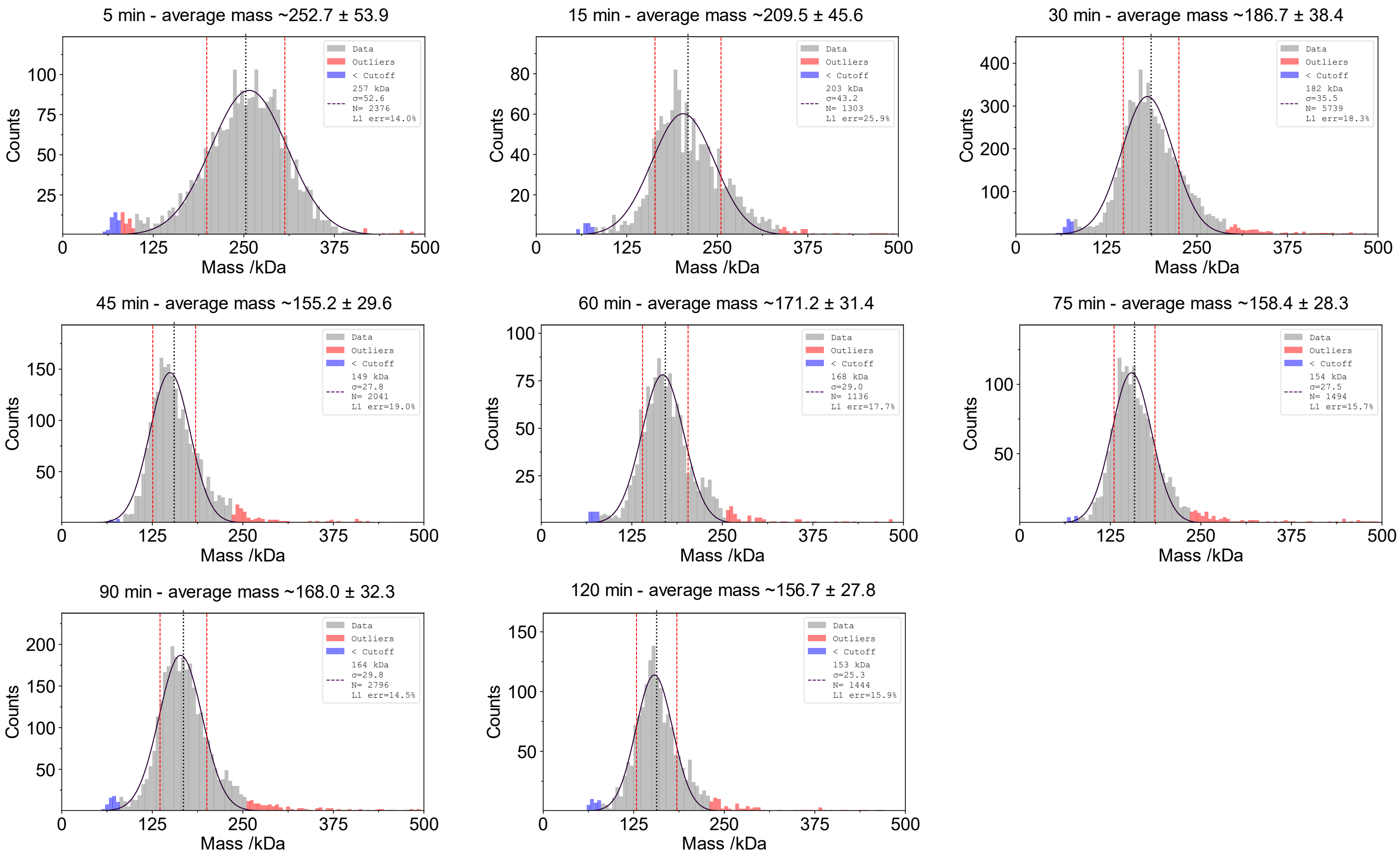

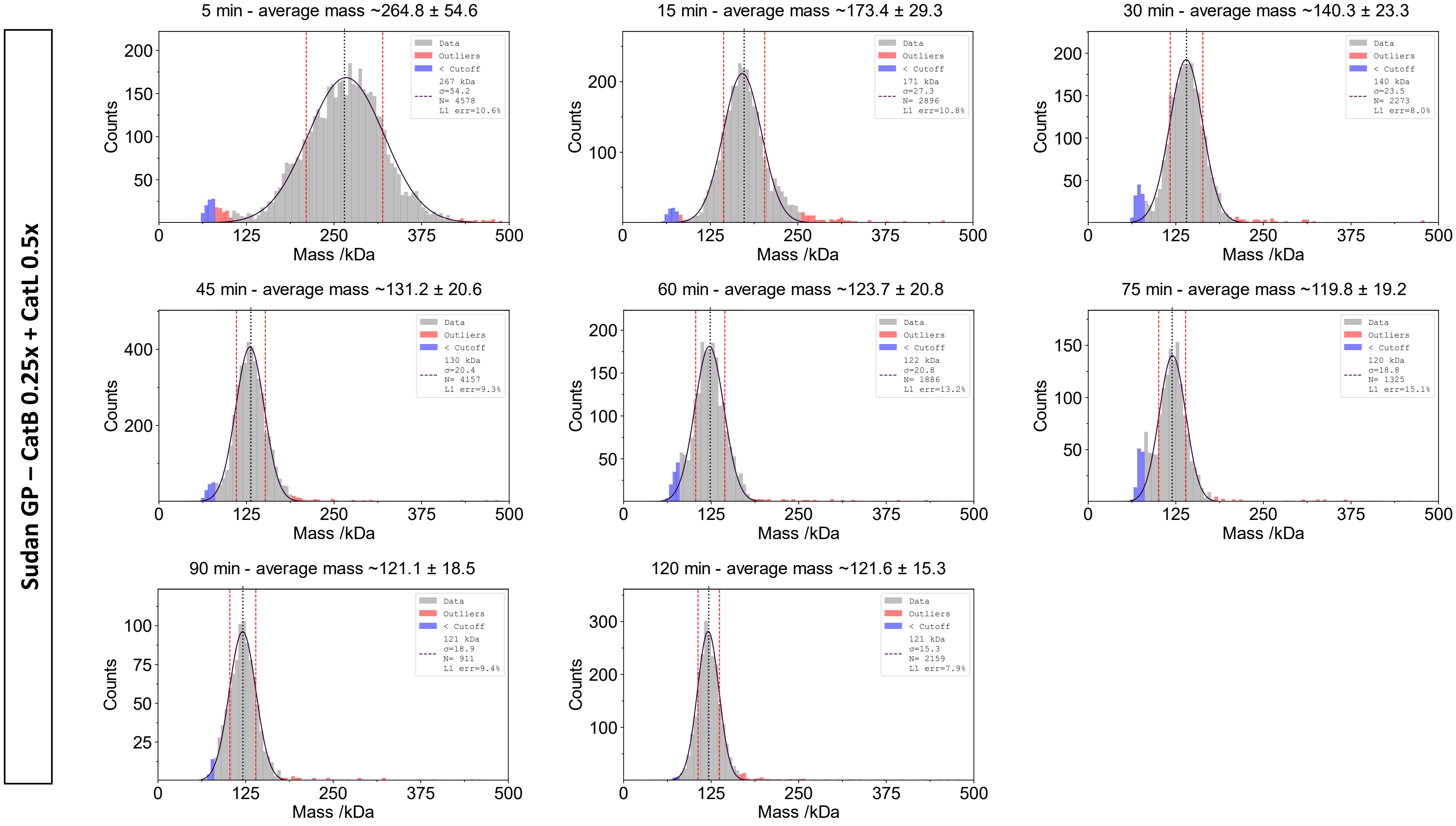
Sudan GP – CatB 0.25x + CatL 0.5x
5 min - average mass ~264.8 ± 54.6
Data
Outliers
< Cutoff
267 kDa
σ=54.2
N= 4578
L1 err=10.6%
15 min - average mass ~173.4 ± 29.3
171 kDa
σ=27.3
N= 2896
L1 err=10.8%
30 min - average mass ~140.3 ± 23.3
140 kDa
σ=23.5
N= 2273
L1 err=8.0%
45 min - average mass ~131.2 ± 20.6
130 kDa
σ=20.4
N= 4157
L1 err=9.3%
60 min - average mass ~123.7 ± 20.8
122 kDa
σ=20.8
N= 1886
L1 err=13.2%
75 min - average mass ~119.8 ± 19.2
120 kDa
σ=18.8
N= 1325
L1 err=15.1%
90 min - average mass ~121.1 ± 18.5
121 kDa
σ=18.9
N= 911
L1 err=9.4%
120 min - average mass ~121.6 ± 15.3
121 kDa
σ=15.3
N= 2159
L1 err=7.9%
Counts
Mass /kDa

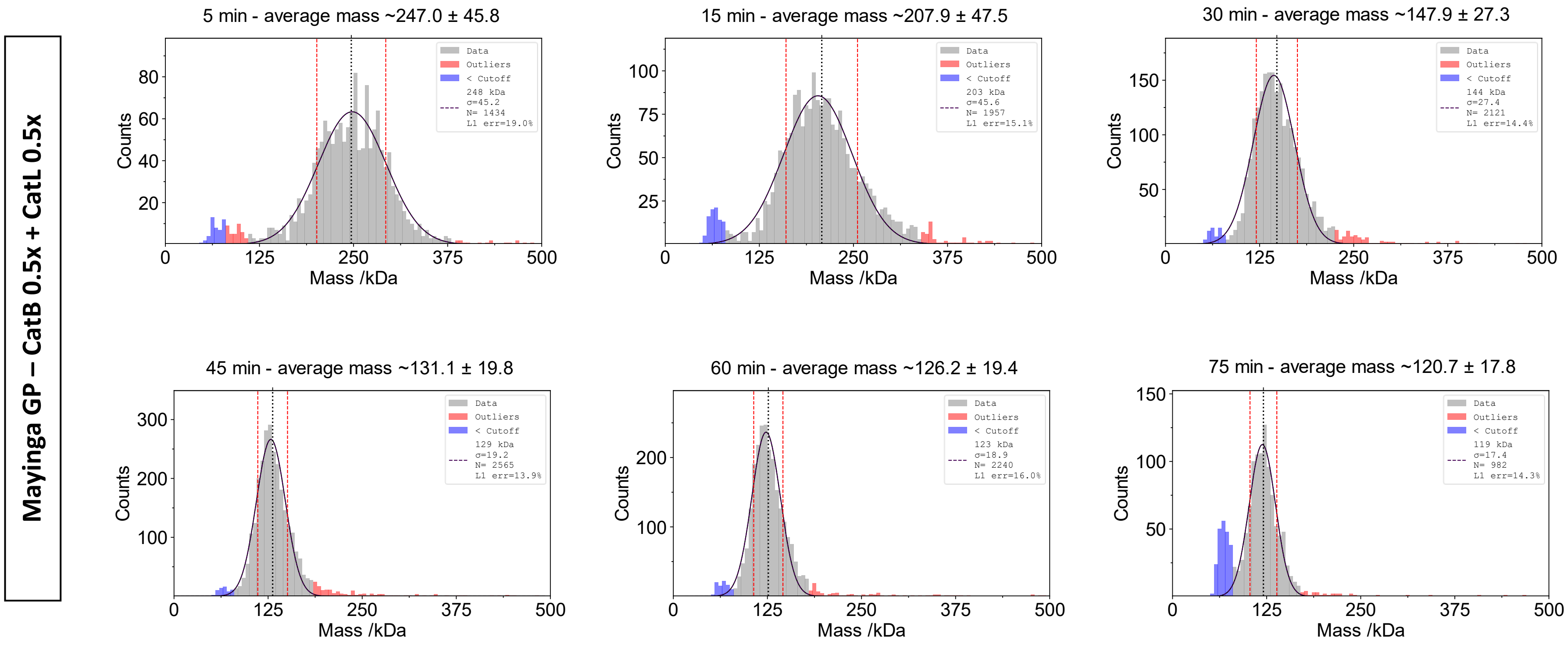

Mayinga GP – CatB 0.5x + CatL 0.5x
5 min - average mass ~247.0 ± 45.8
Data
Outliers
< Cutoff
248 kDa
σ=45.2
N= 1434
L1 err=19.0%
15 min - average mass ~207.9 ± 47.5
Data
Outliers
< Cutoff
203 kDa
σ=45.6
N= 1957
L1 err=15.1%
30 min - average mass ~147.9 ± 27.3
Data
Outliers
< Cutoff
144 kDa
σ=27.4
N= 2121
L1 err=14.4%
45 min - average mass ~131.1 ± 19.8
Data
Outliers
< Cutoff
129 kDa
σ=19.2
N= 2565
L1 err=13.9%
60 min - average mass ~126.2 ± 19.4
Data
Outliers
< Cutoff
123 kDa
σ=18.9
N= 2240
L1 err=16.0%
75 min - average mass ~120.7 ± 17.8
Data
Outliers
< Cutoff
119 kDa
σ=17.4
N= 982
L1 err=14.3%
Counts
Mass /kDa

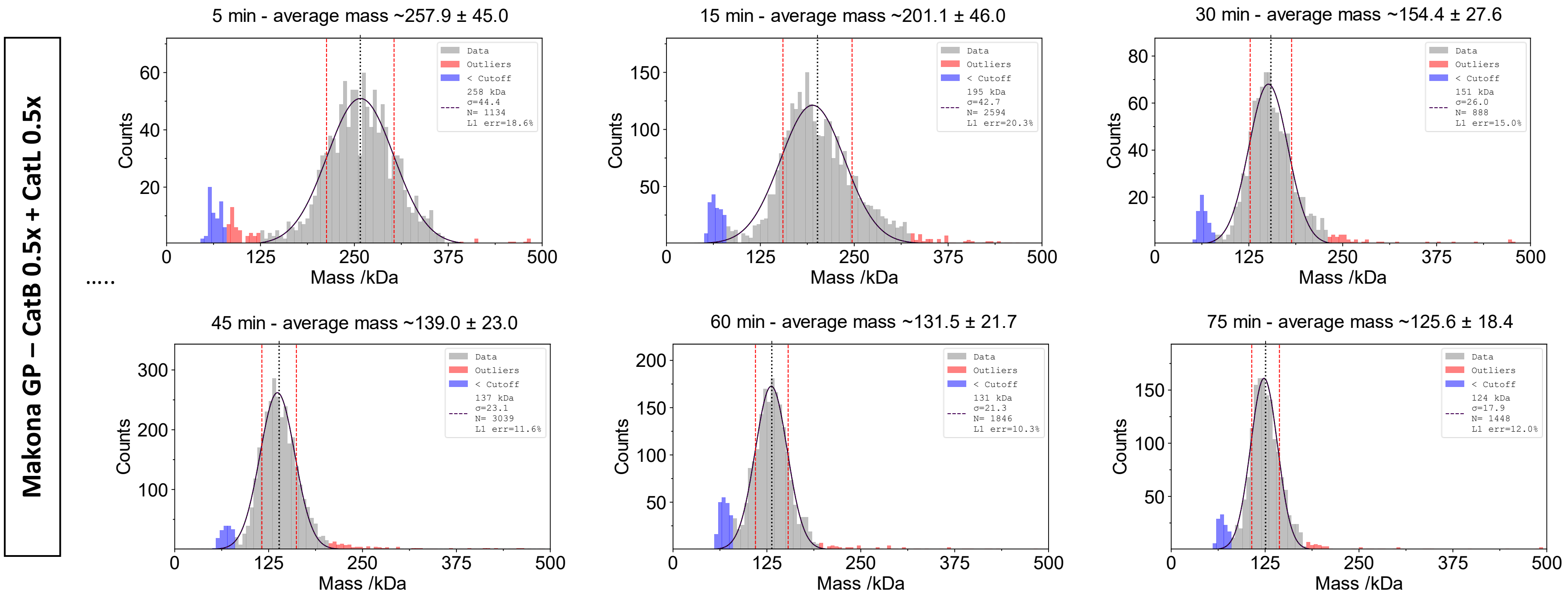
Makona GP – CatB 0.5x + CatL 0.5x
.....
5 min - average mass ~257.9 ± 45.0
Data
Outliers
< Cutoff
258 kDa
σ=44.4
N= 1134
L1 err=18.6%
15 min - average mass ~201.1 ± 46.0
Data
Outliers
< Cutoff
195 kDa
σ=42.7
N= 2594
L1 err=20.3%
30 min - average mass ~154.4 ± 27.6
Data
Outliers
< Cutoff
151 kDa
σ=26.0
N= 888
L1 err=15.0%
45 min - average mass ~139.0 ± 23.0
Data
Outliers
< Cutoff
137 kDa
σ=23.1
N= 3039
L1 err=11.6%
60 min - average mass ~131.5 ± 21.7
Data
Outliers
< Cutoff
131 kDa
σ=21.3
N= 1846
L1 err=10.3%
75 min - average mass ~125.6 ± 18.4
Data
Outliers
< Cutoff
124 kDa
σ=17.9
N= 1448
L1 err=12.0%
Counts
Mass /kDa
0
125
250
375
500

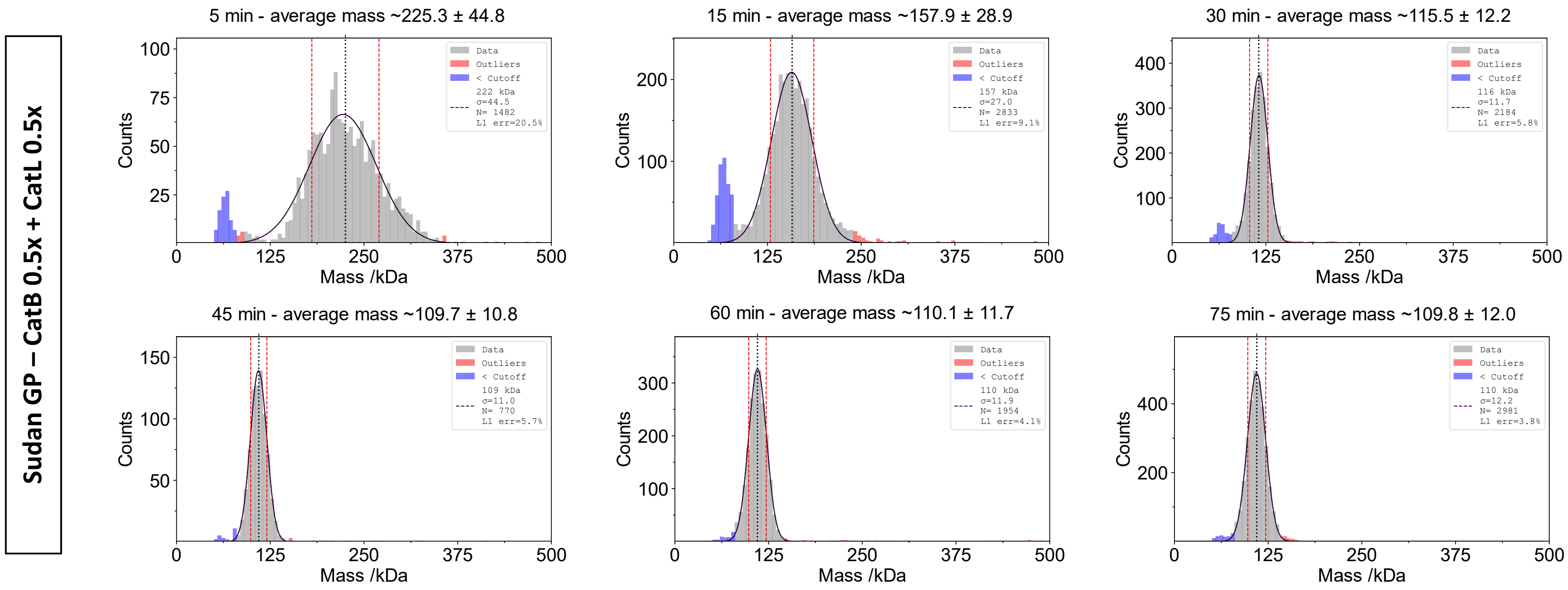
Sudan GP – CatB 0.5x + CatL 0.5x
5 min - average mass ~225.3 ± 44.8
15 min - average mass ~157.9 ± 28.9
30 min - average mass ~115.5 ± 12.2
45 min - average mass ~109.7 ± 10.8
60 min - average mass ~110.1 ± 11.7
75 min - average mass ~109.8 ± 12.0
Data
Outliers
< Cutoff
222 kDa
σ=44.5
N= 1482
L1 err=20.5%
157 kDa
σ=27.0
N= 2833
L1 err=9.1%
116 kDa
σ=11.7
N= 2184
L1 err=5.8%
109 kDa
σ=11.0
N= 770
L1 err=5.7%
110 kDa
σ=11.9
N= 1954
L1 err=4.1%
110 kDa
σ=12.2
N= 2981
L1 err=3.8%
Counts
Mass /kDa

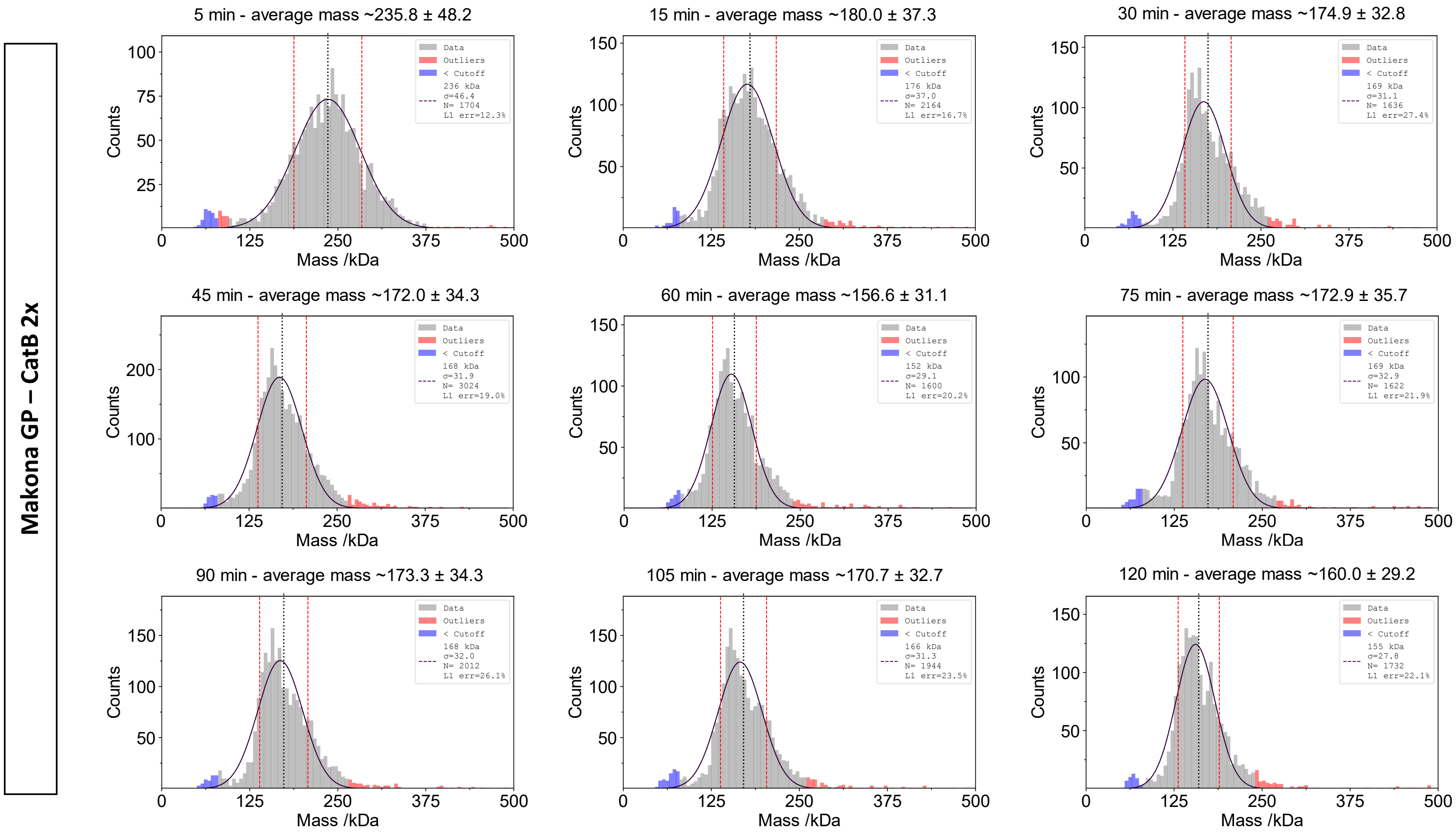
Makona GP – CatB 2x
5 min - average mass ~235.8 ± 48.2
15 min - average mass ~180.0 ± 37.3
30 min - average mass ~174.9 ± 32.8
45 min - average mass ~172.0 ± 34.3
60 min - average mass ~156.6 ± 31.1
75 min - average mass ~172.9 ± 35.7
90 min - average mass ~173.3 ± 34.3
105 min - average mass ~170.7 ± 32.7
120 min - average mass ~160.0 ± 29.2
Data
Outliers
< Cutoff
236 kDa σ=46.4 N= 1704 L1 err=12.3%
176 kDa σ=37.0 N= 2164 L1 err=16.7%
169 kDa σ=31.1 N= 1636 L1 err=27.4%
168 kDa σ=31.9 N= 3024 L1 err=19.0%
152 kDa σ=29.1 N= 1600 L1 err=20.2%
169 kDa σ=32.9 N= 1622 L1 err=21.9%
168 kDa σ=32.0 N= 2012 L1 err=26.1%
166 kDa σ=31.3 N= 1944 L1 err=23.5%
155 kDa σ=27.8 N= 1732 L1 err=22.1%
Counts
Mass /kDa

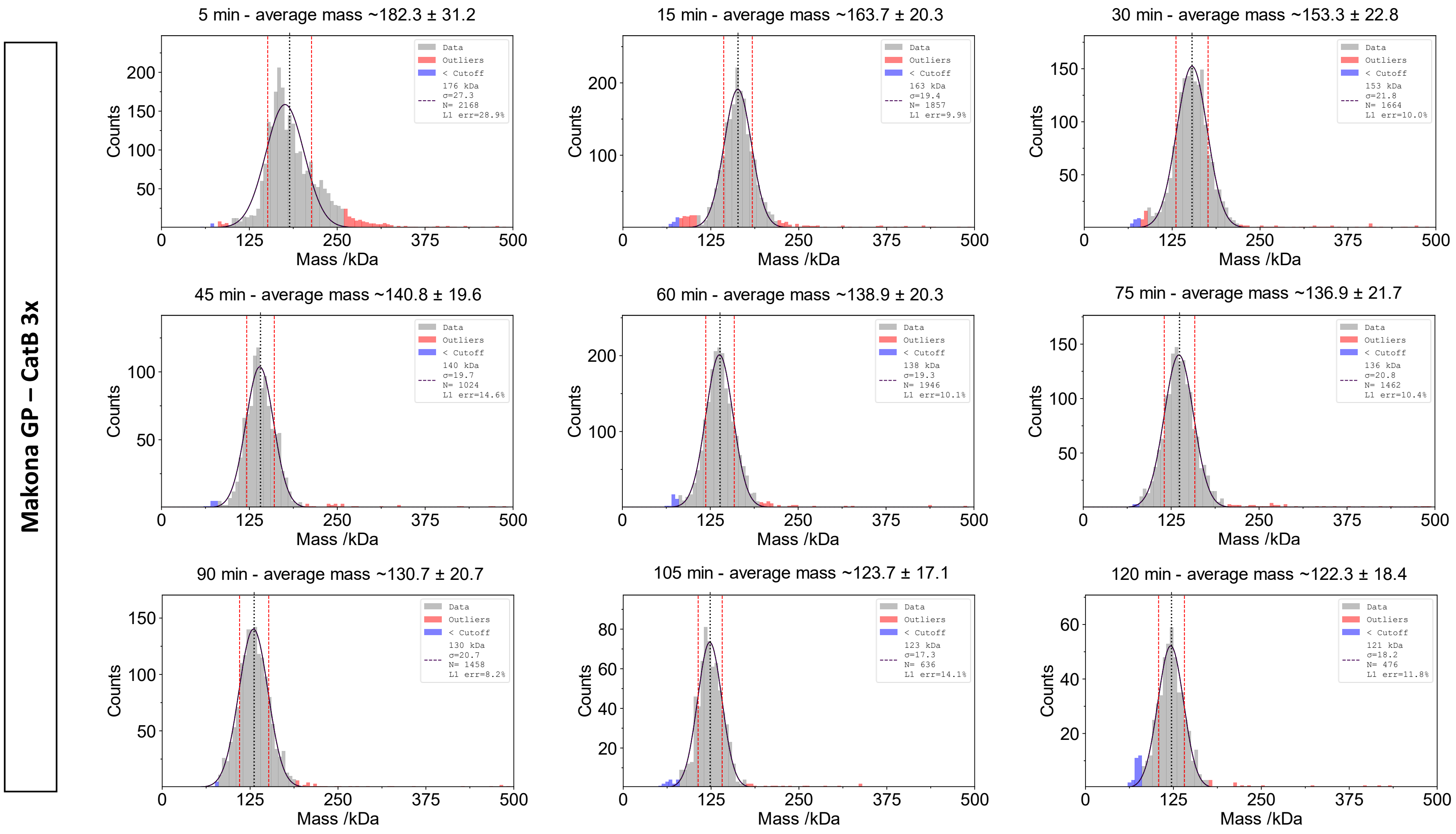

Makona GP – CatB 3x
5 min - average mass ~182.3 ± 31.2
Data
Outliers
< Cutoff
176 kDa
σ=27.3
N= 2168
L1 err=28.9%
15 min - average mass ~163.7 ± 20.3
Data
Outliers
< Cutoff
163 kDa
σ=19.4
N= 1857
L1 err=9.9%
30 min - average mass ~153.3 ± 22.8
Data
Outliers
< Cutoff
153 kDa
σ=21.8
N= 1664
L1 err=10.0%
45 min - average mass ~140.8 ± 19.6
Data
Outliers
< Cutoff
140 kDa
σ=19.7
N= 1024
L1 err=14.6%
60 min - average mass ~138.9 ± 20.3
Data
Outliers
< Cutoff
138 kDa
σ=19.3
N= 1946
L1 err=10.1%
75 min - average mass ~136.9 ± 21.7
Data
Outliers
< Cutoff
136 kDa
σ=20.8
N= 1462
L1 err=10.4%
90 min - average mass ~130.7 ± 20.7
Data
Outliers
< Cutoff
130 kDa
σ=20.7
N= 1458
L1 err=8.2%
105 min - average mass ~123.7 ± 17.1
Data
Outliers
< Cutoff
123 kDa
σ=17.3
N= 636
L1 err=14.1%
120 min - average mass ~122.3 ± 18.4
Data
Outliers
< Cutoff
121 kDa
σ=18.2
N= 476
L1 err=11.8%
Counts
Mass /kDa
0
125
250
375
500

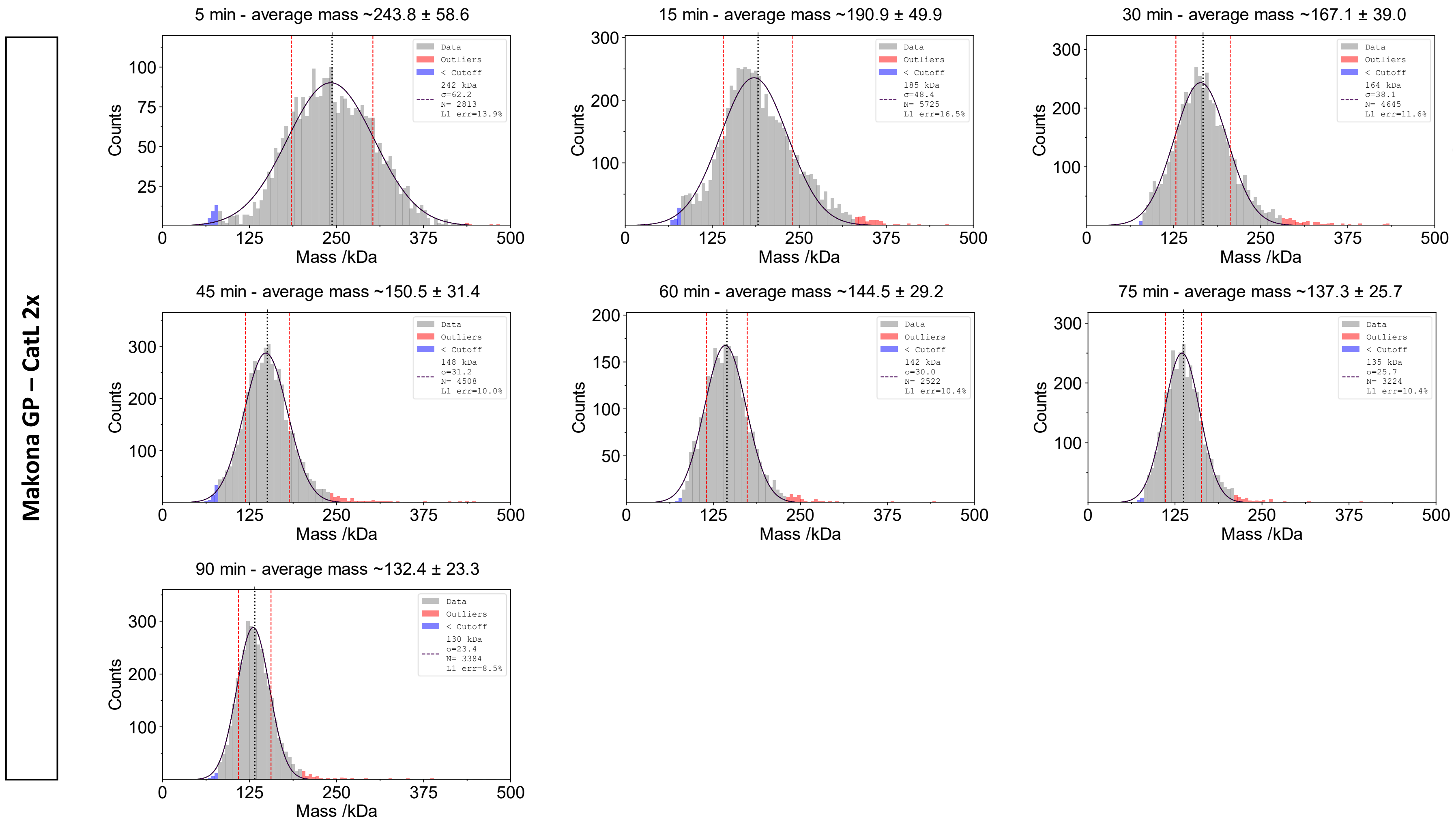

Makona GP – CatL 2x
5 min - average mass ~243.8 ± 58.6
242 kDa σ=62.2 N= 2813 L1 err=13.9%
15 min - average mass ~190.9 ± 49.9
185 kDa σ=48.4 N= 5725 L1 err=16.5%
30 min - average mass ~167.1 ± 39.0
164 kDa σ=38.1 N= 4645 L1 err=11.6%
45 min - average mass ~150.5 ± 31.4
148 kDa σ=31.2 N= 4508 L1 err=10.0%
60 min - average mass ~144.5 ± 29.2
142 kDa σ=30.0 N= 2522 L1 err=10.4%
75 min - average mass ~137.3 ± 25.7
135 kDa σ=25.7 N= 3224 L1 err=10.4%
90 min - average mass ~132.4 ± 23.3
130 kDa σ=23.4 N= 3384 L1 err=8.5%
Data
Outliers
< Cutoff
Counts
Mass /kDa

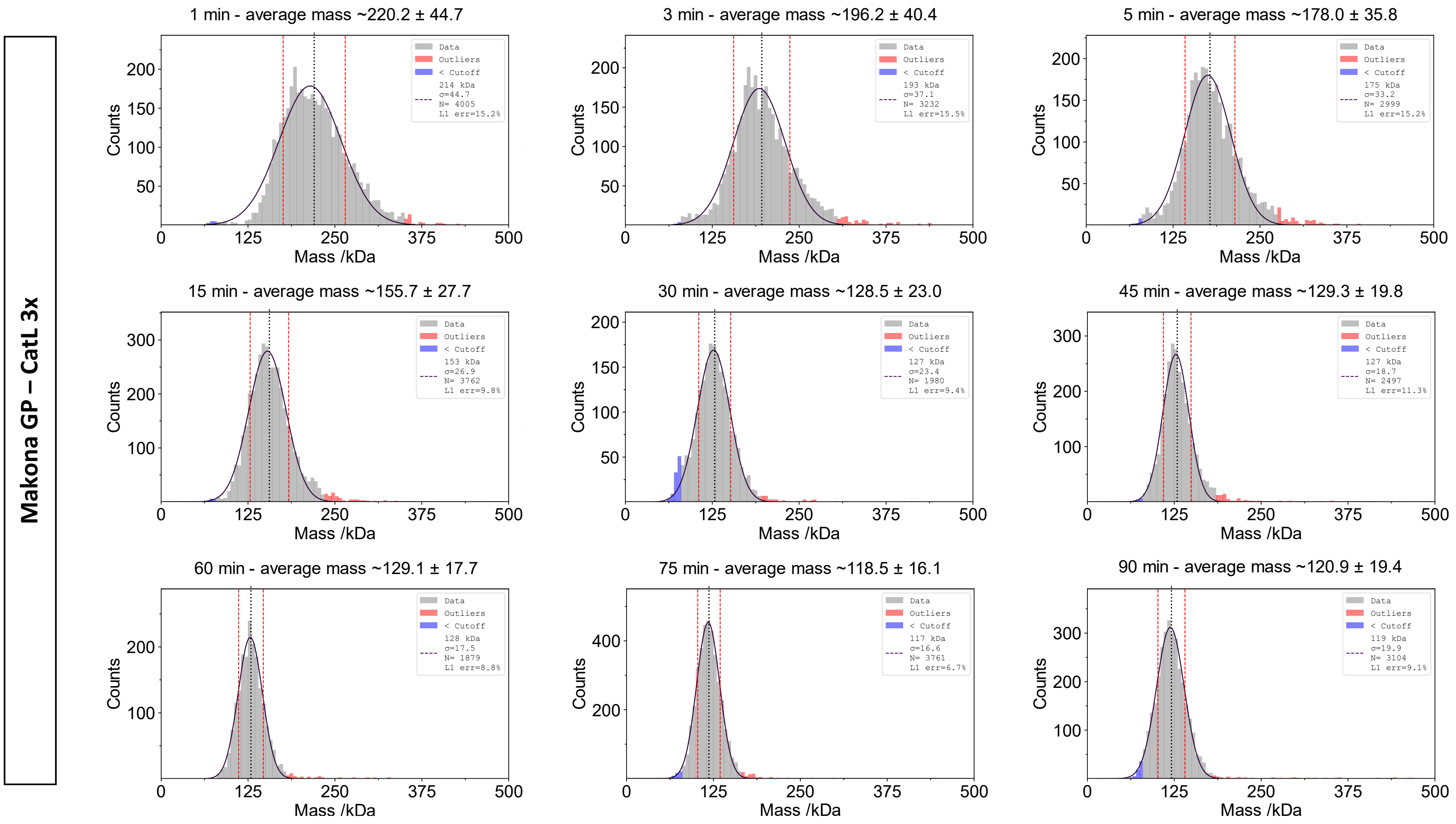
Makona GP – CatL 3x
1 min - average mass ~220.2 ± 44.7
Data
Outliers
< Cutoff
214 kDa
σ=44.7
N= 4005
L1 err=15.2%
3 min - average mass ~196.2 ± 40.4
Data
Outliers
< Cutoff
193 kDa
σ=37.1
N= 3232
L1 err=15.5%
5 min - average mass ~178.0 ± 35.8
Data
Outliers
< Cutoff
175 kDa
σ=33.2
N= 2999
L1 err=15.2%
15 min - average mass ~155.7 ± 27.7
Data
Outliers
< Cutoff
153 kDa
σ=26.9
N= 3762
L1 err=9.8%
30 min - average mass ~128.5 ± 23.0
Data
Outliers
< Cutoff
127 kDa
σ=23.4
N= 1980
L1 err=9.4%
45 min - average mass ~129.3 ± 19.8
Data
Outliers
< Cutoff
127 kDa
σ=18.7
N= 2497
L1 err=11.3%
60 min - average mass ~129.1 ± 17.7
Data
Outliers
< Cutoff
128 kDa
σ=17.5
N= 1879
L1 err=8.8%
75 min - average mass ~118.5 ± 16.1
Data
Outliers
< Cutoff
117 kDa
σ=16.6
N= 3761
L1 err=6.7%
90 min - average mass ~120.9 ± 19.4
Data
Outliers
< Cutoff
119 kDa
σ=19.9
N= 3104
L1 err=9.1%
Counts
Mass /kDa
0
125
250
375
500

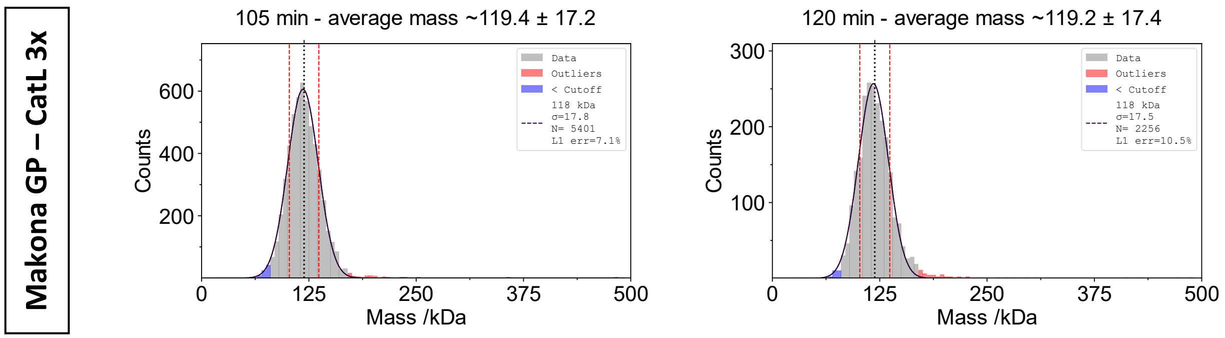
Makona GP – CatL 3x
105 min - average mass ~119.4 ± 17.2
Data
Outliers
< Cutoff
118 kDa
σ=17.8
N= 5401
L1 err=7.1%
Counts
600
400
200
0
125
250
375
500
Mass /kDa
120 min - average mass ~119.2 ± 17.4
Data
Outliers
< Cutoff
118 kDa
σ=17.5
N= 2256
L1 err=10.5%
Counts
300
200
100
0
125
250
375
500
Mass /kDa

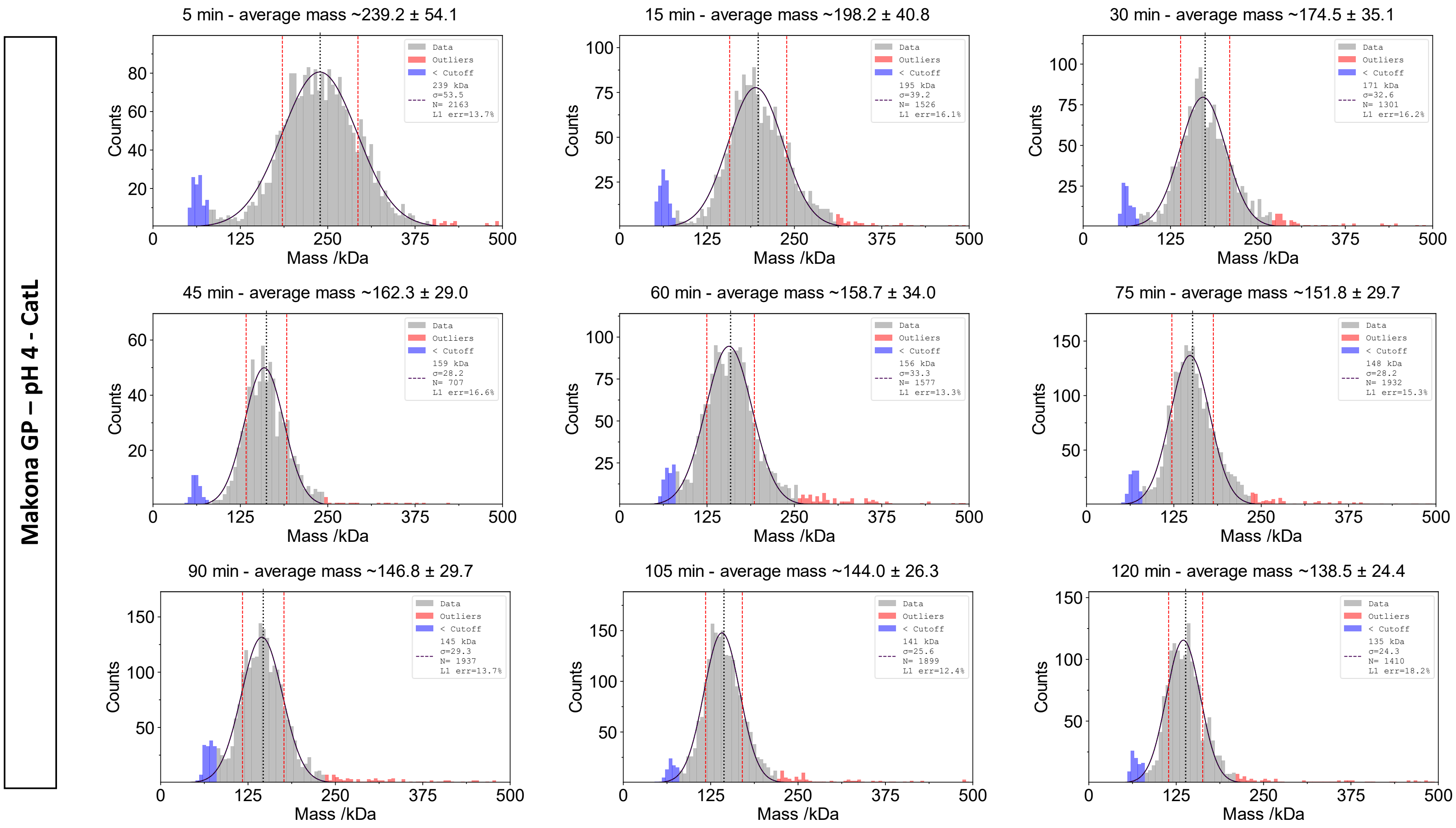

Makona GP – pH 4 - CatL
5 min - average mass ~239.2 ± 54.1
15 min - average mass ~198.2 ± 40.8
30 min - average mass ~174.5 ± 35.1
45 min - average mass ~162.3 ± 29.0
60 min - average mass ~158.7 ± 34.0
75 min - average mass ~151.8 ± 29.7
90 min - average mass ~146.8 ± 29.7
105 min - average mass ~144.0 ± 26.3
120 min - average mass ~138.5 ± 24.4
Data
Outliers
< Cutoff
239 kDa σ=53.5 N= 2163 L1 err=13.7%
195 kDa σ=39.2 N= 1526 L1 err=16.1%
171 kDa σ=32.6 N= 1301 L1 err=16.2%
159 kDa σ=28.2 N= 707 L1 err=16.6%
156 kDa σ=33.3 N= 1577 L1 err=13.3%
148 kDa σ=28.2 N= 1932 L1 err=15.3%
145 kDa σ=29.3 N= 1937 L1 err=13.7%
141 kDa σ=25.6 N= 1899 L1 err=12.4%
135 kDa σ=24.3 N= 1410 L1 err=18.2%
Counts
Mass /kDa

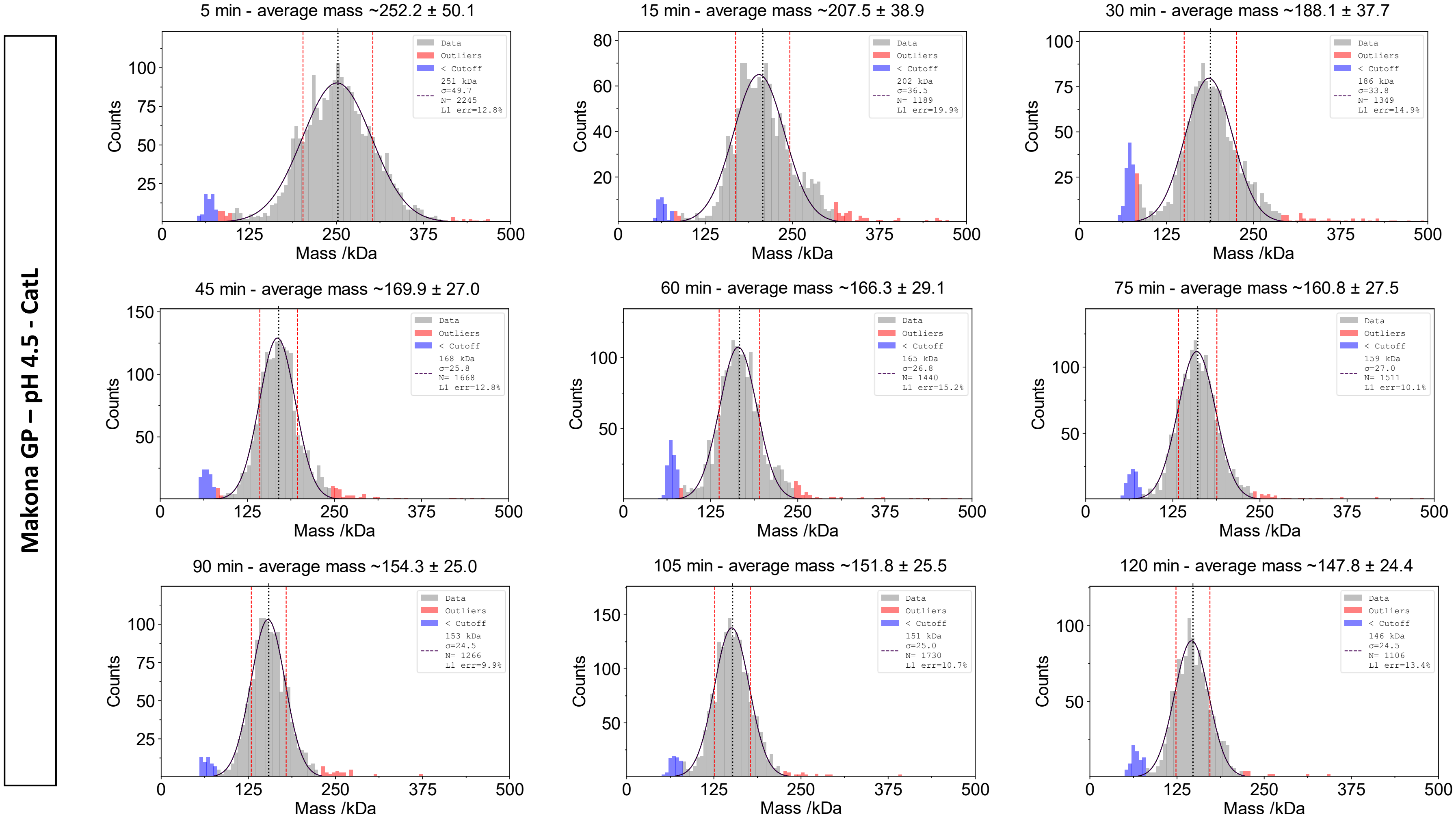
Makona GP – pH 4.5 - CatL
5 min - average mass ~252.2 ± 50.1
Data
Outliers
< Cutoff
251 kDa
σ=49.7
N= 2245
L1 err=12.8%
15 min - average mass ~207.5 ± 38.9
Data
Outliers
< Cutoff
202 kDa
σ=36.5
N= 1189
L1 err=19.9%
30 min - average mass ~188.1 ± 37.7
Data
Outliers
< Cutoff
186 kDa
σ=33.8
N= 1349
L1 err=14.9%
45 min - average mass ~169.9 ± 27.0
Data
Outliers
< Cutoff
168 kDa
σ=25.8
N= 1668
L1 err=12.8%
60 min - average mass ~166.3 ± 29.1
Data
Outliers
< Cutoff
165 kDa
σ=26.8
N= 1440
L1 err=15.2%
75 min - average mass ~160.8 ± 27.5
Data
Outliers
< Cutoff
159 kDa
σ=27.0
N= 1511
L1 err=10.1%
90 min - average mass ~154.3 ± 25.0
Data
Outliers
< Cutoff
153 kDa
σ=24.5
N= 1266
L1 err=9.9%
105 min - average mass ~151.8 ± 25.5
Data
Outliers
< Cutoff
151 kDa
σ=25.0
N= 1730
L1 err=10.7%
120 min - average mass ~147.8 ± 24.4
Data
Outliers
< Cutoff
146 kDa
σ=24.5
N= 1106
L1 err=13.4%
Counts
Mass /kDa
0
125
250
375
500

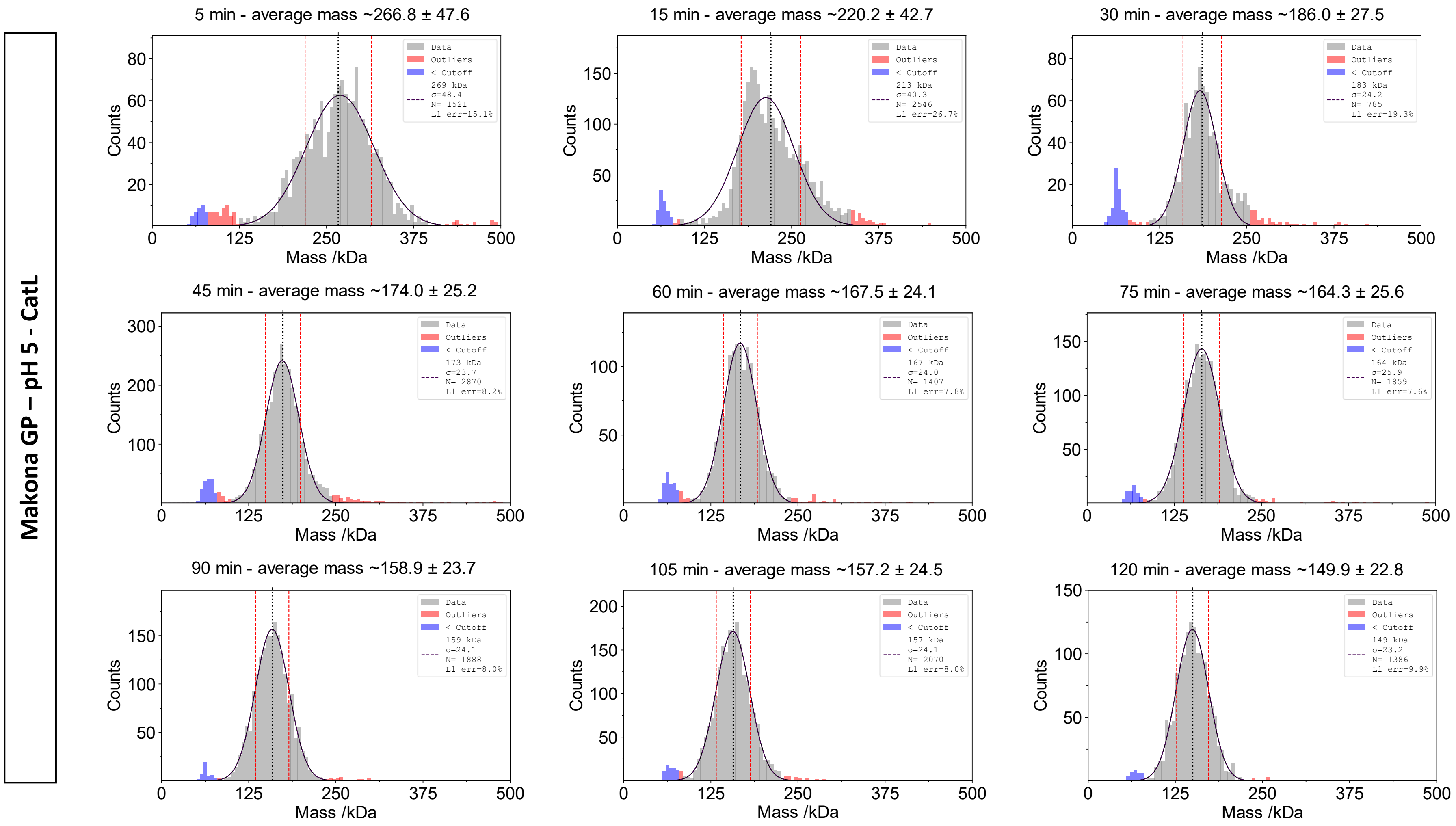

Makona GP – pH 5 - CatL
5 min - average mass ~266.8 ± 47.6
Data
Outliers
< Cutoff
269 kDa
σ=48.4
N= 1521
L1 err=15.1%
15 min - average mass ~220.2 ± 42.7
Data
Outliers
< Cutoff
213 kDa
σ=40.3
N= 2546
L1 err=26.7%
30 min - average mass ~186.0 ± 27.5
Data
Outliers
< Cutoff
183 kDa
σ=24.2
N= 785
L1 err=19.3%
45 min - average mass ~174.0 ± 25.2
Data
Outliers
< Cutoff
173 kDa
σ=23.7
N= 2870
L1 err=8.2%
60 min - average mass ~167.5 ± 24.1
Data
Outliers
< Cutoff
167 kDa
σ=24.0
N= 1407
L1 err=7.8%
75 min - average mass ~164.3 ± 25.6
Data
Outliers
< Cutoff
164 kDa
σ=25.9
N= 1859
L1 err=7.6%
90 min - average mass ~158.9 ± 23.7
Data
Outliers
< Cutoff
159 kDa
σ=24.1
N= 1888
L1 err=8.0%
105 min - average mass ~157.2 ± 24.5
Data
Outliers
< Cutoff
157 kDa
σ=24.1
N= 2070
L1 err=8.0%
120 min - average mass ~149.9 ± 22.8
Data
Outliers
< Cutoff
149 kDa
σ=23.2
N= 1386
L1 err=9.9%
Counts
Mass /kDa

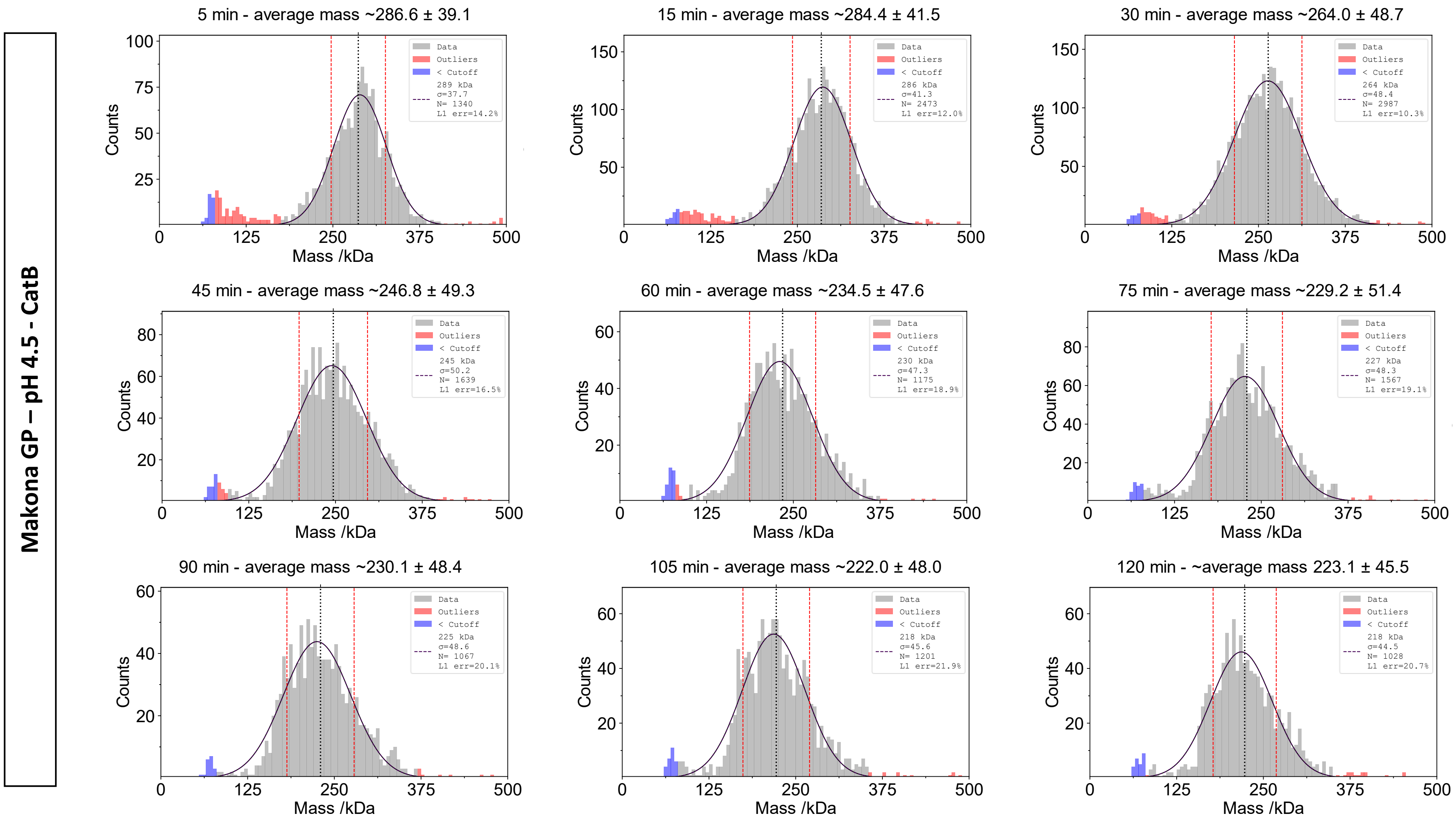
Makona GP – pH 4.5 - CatB
5 min - average mass ~286.6 ± 39.1
289 kDa
σ=37.7
N= 1340
L1 err=14.2%
15 min - average mass ~284.4 ± 41.5
286 kDa
σ=41.3
N= 2473
L1 err=12.0%
30 min - average mass ~264.0 ± 48.7
264 kDa
σ=48.4
N= 2987
L1 err=10.3%
45 min - average mass ~246.8 ± 49.3
245 kDa
σ=50.2
N= 1639
L1 err=16.5%
60 min - average mass ~234.5 ± 47.6
230 kDa
σ=47.3
N= 1175
L1 err=18.9%
75 min - average mass ~229.2 ± 51.4
227 kDa
σ=48.3
N= 1567
L1 err=19.1%
90 min - average mass ~230.1 ± 48.4
225 kDa
σ=48.6
N= 1067
L1 err=20.1%
105 min - average mass ~222.0 ± 48.0
218 kDa
σ=45.6
N= 1201
L1 err=21.9%
120 min - ~average mass 223.1 ± 45.5
218 kDa
σ=44.5
N= 1028
L1 err=20.7%
Data
Outliers
< Cutoff
Counts
Mass /kDa

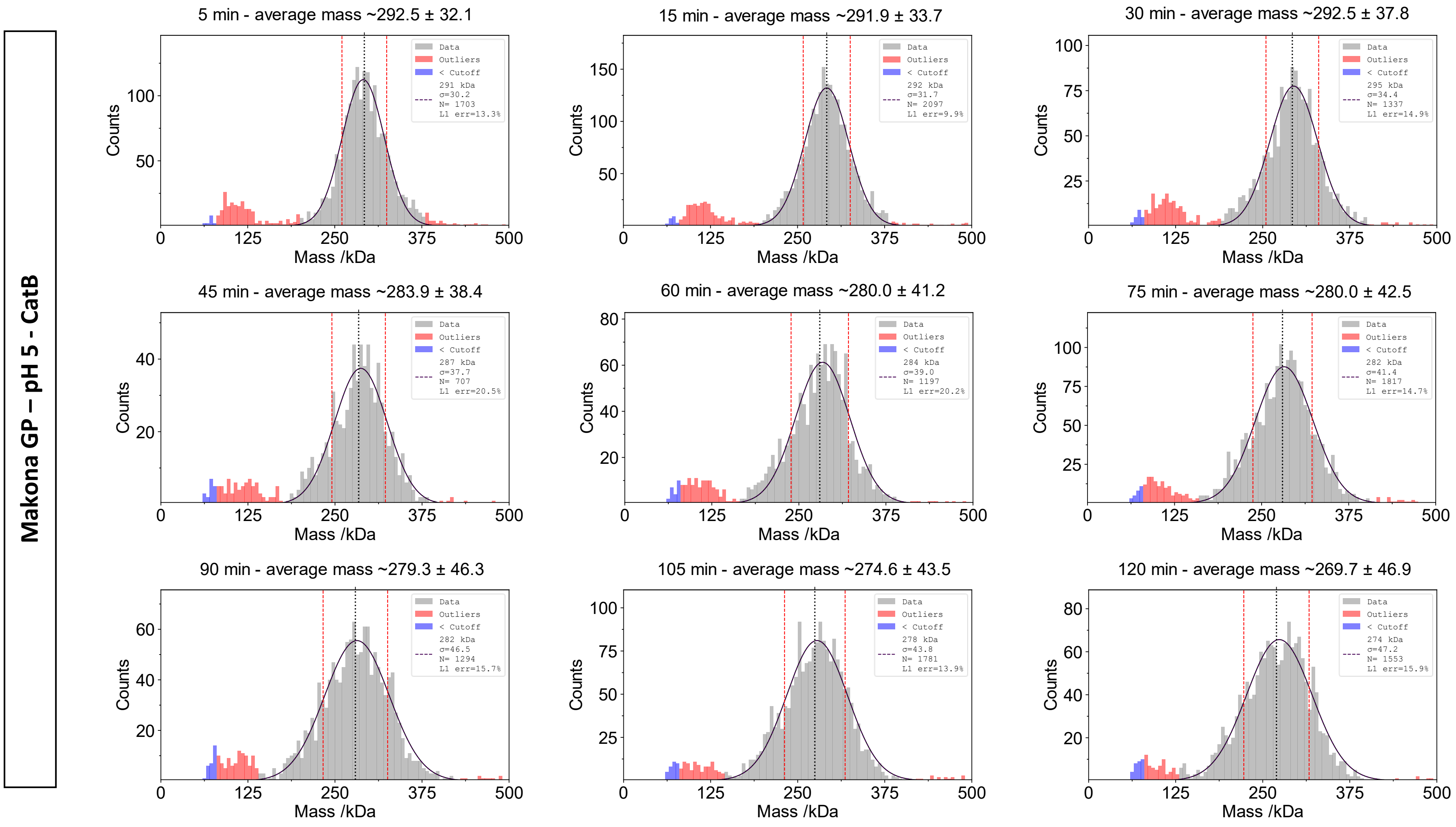
Makona GP – pH 5 - CatB
5 min - average mass ~292.5 ± 32.1
Data
Outliers
< Cutoff
291 kDa
σ=30.2
N= 1703
L1 err=13.3%
15 min - average mass ~291.9 ± 33.7
Data
Outliers
< Cutoff
292 kDa
σ=31.7
N= 2097
L1 err=9.9%
30 min - average mass ~292.5 ± 37.8
Data
Outliers
< Cutoff
295 kDa
σ=34.4
N= 1337
L1 err=14.9%
45 min - average mass ~283.9 ± 38.4
Data
Outliers
< Cutoff
287 kDa
σ=37.7
N= 707
L1 err=20.5%
60 min - average mass ~280.0 ± 41.2
Data
Outliers
< Cutoff
284 kDa
σ=39.0
N= 1197
L1 err=20.2%
75 min - average mass ~280.0 ± 42.5
Data
Outliers
< Cutoff
282 kDa
σ=41.4
N= 1817
L1 err=14.7%
90 min - average mass ~279.3 ± 46.3
Data
Outliers
< Cutoff
282 kDa
σ=46.5
N= 1294
L1 err=15.7%
105 min - average mass ~274.6 ± 43.5
Data
Outliers
< Cutoff
278 kDa
σ=43.8
N= 1781
L1 err=13.9%
120 min - average mass ~269.7 ± 46.9
Data
Outliers
< Cutoff
274 kDa
σ=47.2
N= 1553
L1 err=15.9%
Counts
Mass /kDa
0
125
250
375
500

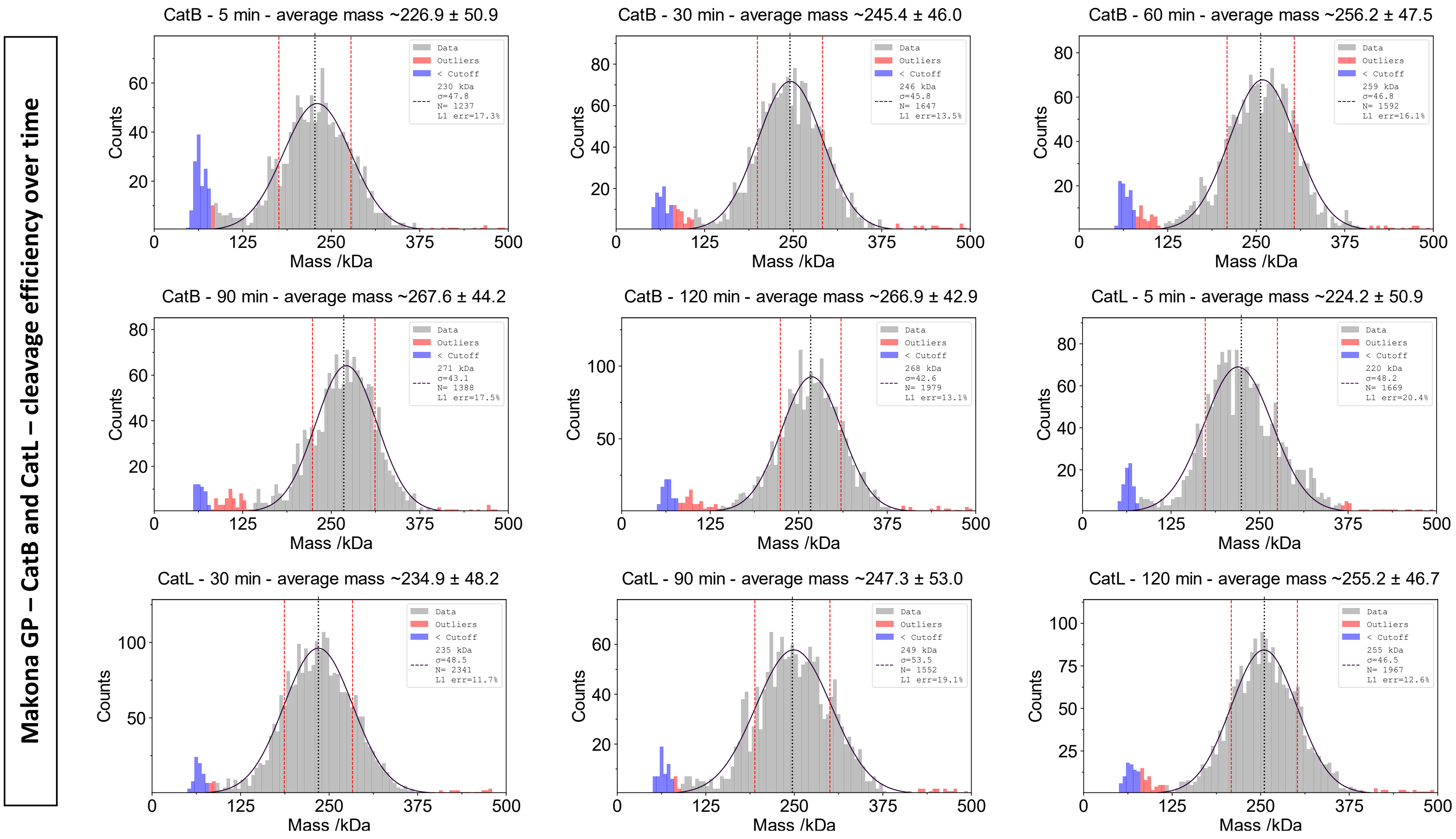

Makona GP – CatB and CatL – cleavage efficiency over time
CatB - 5 min - average mass ~226.9 ± 50.9
Data
Outliers
< Cutoff
230 kDa
σ=47.8
N= 1237
L1 err=17.3%
CatB - 30 min - average mass ~245.4 ± 46.0
246 kDa
σ=45.8
N= 1647
L1 err=13.5%
CatB - 60 min - average mass ~256.2 ± 47.5
259 kDa
σ=46.8
N= 1592
L1 err=16.1%
CatB - 90 min - average mass ~267.6 ± 44.2
271 kDa
σ=43.1
N= 1388
L1 err=17.5%
CatB - 120 min - average mass ~266.9 ± 42.9
268 kDa
σ=42.6
N= 1979
L1 err=13.1%
CatL - 5 min - average mass ~224.2 ± 50.9
220 kDa
σ=48.2
N= 1669
L1 err=20.4%
CatL - 30 min - average mass ~234.9 ± 48.2
235 kDa
σ=48.5
N= 2341
L1 err=11.7%
CatL - 90 min - average mass ~247.3 ± 53.0
249 kDa
σ=53.5
N= 1552
L1 err=19.1%
CatL - 120 min - average mass ~255.2 ± 46.7
255 kDa
σ=46.5
N= 1967
L1 err=12.6%
Counts
Mass /kDa

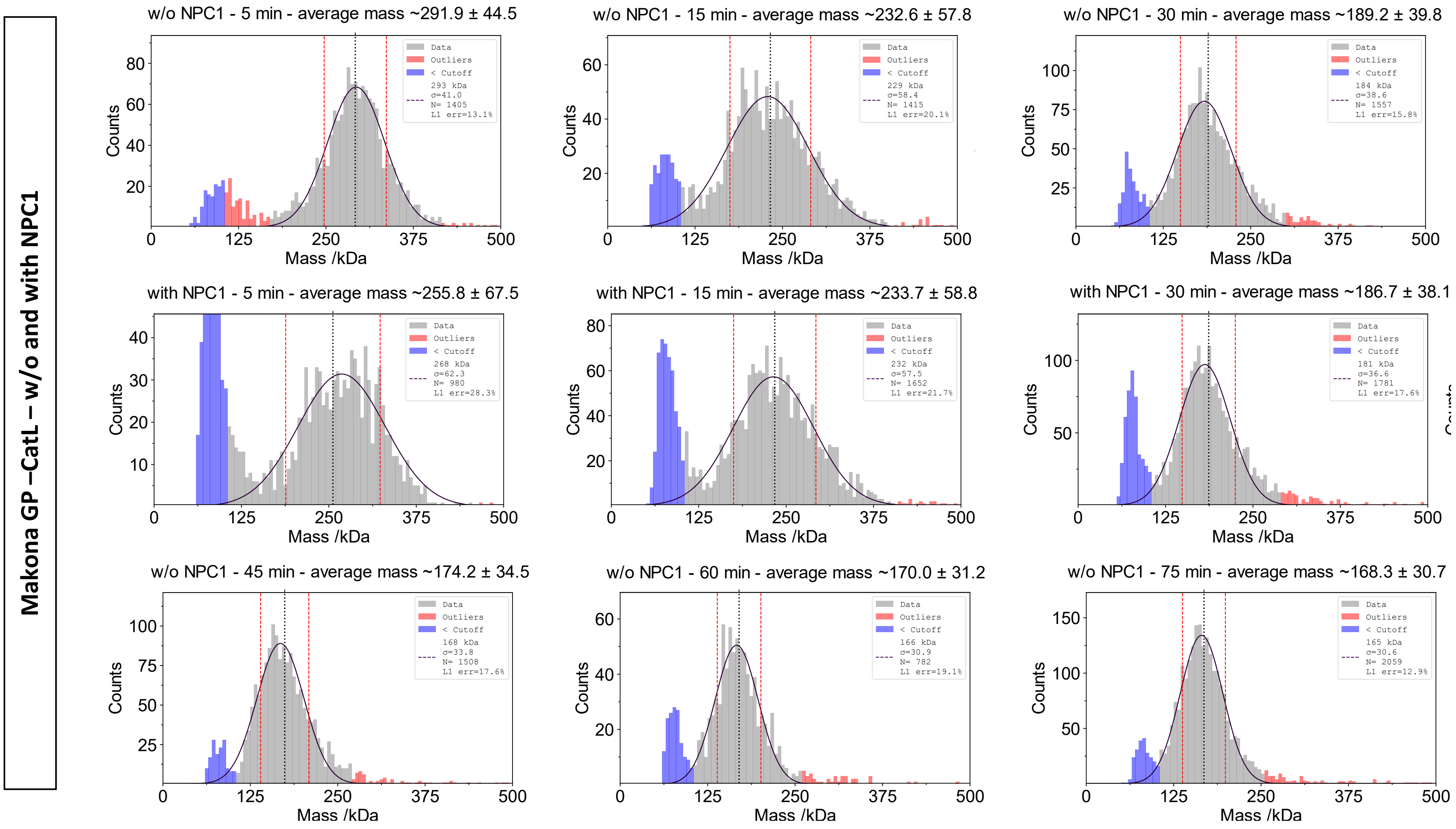
Makona GP –CatL – w/o and with NPC1
w/o NPC1 - 5 min - average mass ~291.9 ± 44.5
Data
Outliers
< Cutoff
293 kDa
σ=41.0
N= 1405
L1 err=13.1%
w/o NPC1 - 15 min - average mass ~232.6 ± 57.8
Data
Outliers
< Cutoff
229 kDa
σ=58.4
N= 1415
L1 err=20.1%
w/o NPC1 - 30 min - average mass ~189.2 ± 39.8
Data
Outliers
< Cutoff
184 kDa
σ=38.6
N= 1557
L1 err=15.8%
with NPC1 - 5 min - average mass ~255.8 ± 67.5
Data
Outliers
< Cutoff
268 kDa
σ=62.3
N= 980
L1 err=28.3%
with NPC1 - 15 min - average mass ~233.7 ± 58.8
Data
Outliers
< Cutoff
232 kDa
σ=57.5
N= 1652
L1 err=21.7%
with NPC1 - 30 min - average mass ~186.7 ± 38.1
Data
Outliers
< Cutoff
181 kDa
σ=36.6
N= 1781
L1 err=17.6%
w/o NPC1 - 45 min - average mass ~174.2 ± 34.5
Data
Outliers
< Cutoff
168 kDa
σ=33.8
N= 1508
L1 err=17.6%
w/o NPC1 - 60 min - average mass ~170.0 ± 31.2
Data
Outliers
< Cutoff
166 kDa
σ=30.9
N= 782
L1 err=19.1%
w/o NPC1 - 75 min - average mass ~168.3 ± 30.7
Data
Outliers
< Cutoff
165 kDa
σ=30.6
N= 2059
L1 err=12.9%
Counts
Mass /kDa

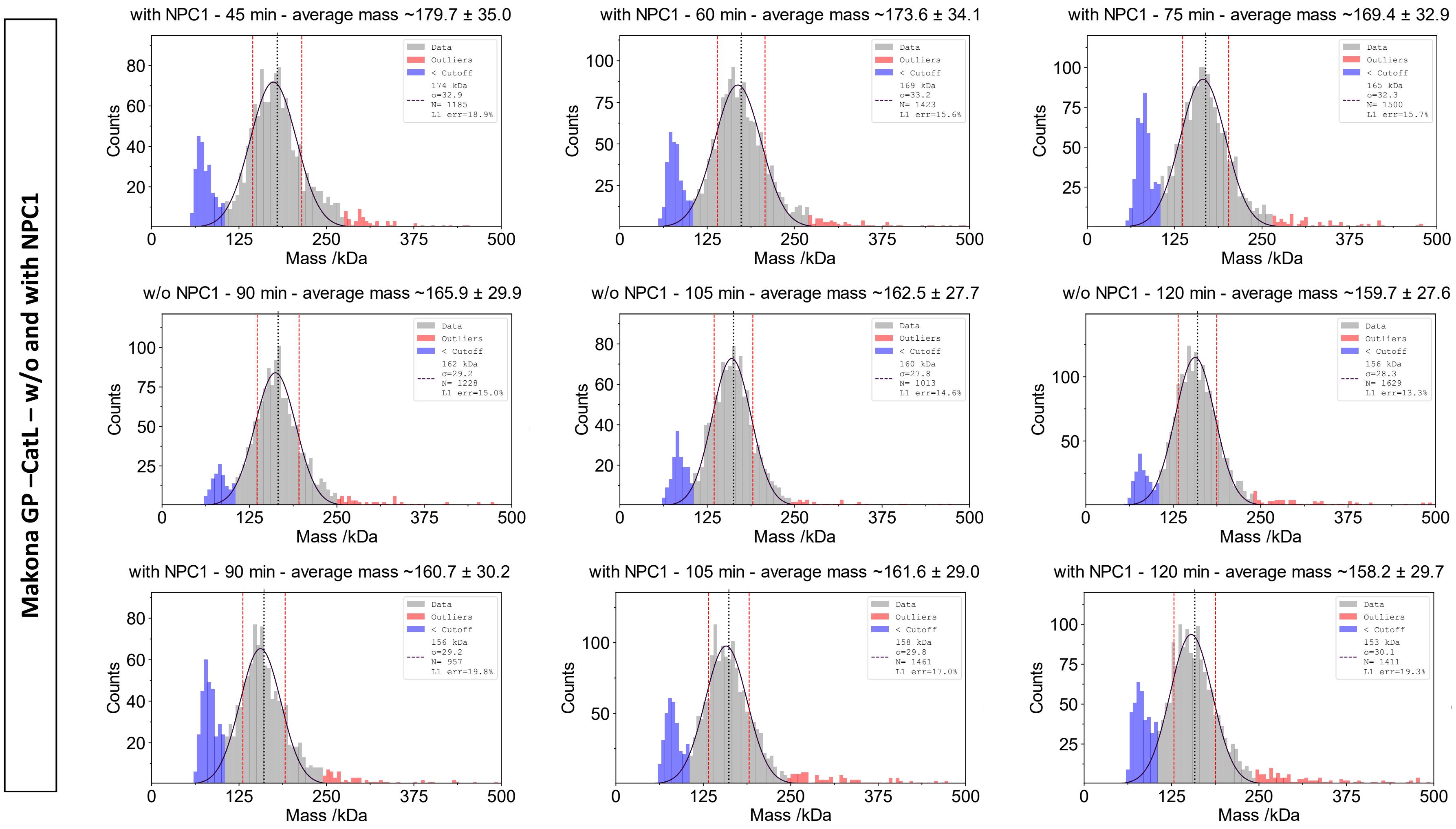
Makona GP –CatL – w/o and with NPC1
with NPC1 - 45 min - average mass ~179.7 ± 35.0
174 kDa σ=32.9 N= 1185 L1 err=18.9%
with NPC1 - 60 min - average mass ~173.6 ± 34.1
169 kDa σ=33.2 N= 1423 L1 err=15.6%
with NPC1 - 75 min - average mass ~169.4 ± 32.9
165 kDa σ=32.3 N= 1500 L1 err=15.7%
w/o NPC1 - 90 min - average mass ~165.9 ± 29.9
162 kDa σ=29.2 N= 1228 L1 err=15.0%
w/o NPC1 - 105 min - average mass ~162.5 ± 27.7
160 kDa σ=27.8 N= 1013 L1 err=14.6%
w/o NPC1 - 120 min - average mass ~159.7 ± 27.6
156 kDa σ=28.3 N= 1629 L1 err=13.3%
with NPC1 - 90 min - average mass ~160.7 ± 30.2
156 kDa σ=29.2 N= 957 L1 err=19.8%
with NPC1 - 105 min - average mass ~161.6 ± 29.0
158 kDa σ=29.8 N= 1461 L1 err=17.0%
with NPC1 - 120 min - average mass ~158.2 ± 29.7
153 kDa σ=30.1 N= 1411 L1 err=19.3%
Data
Outliers
< Cutoff
Counts
Mass /kDa

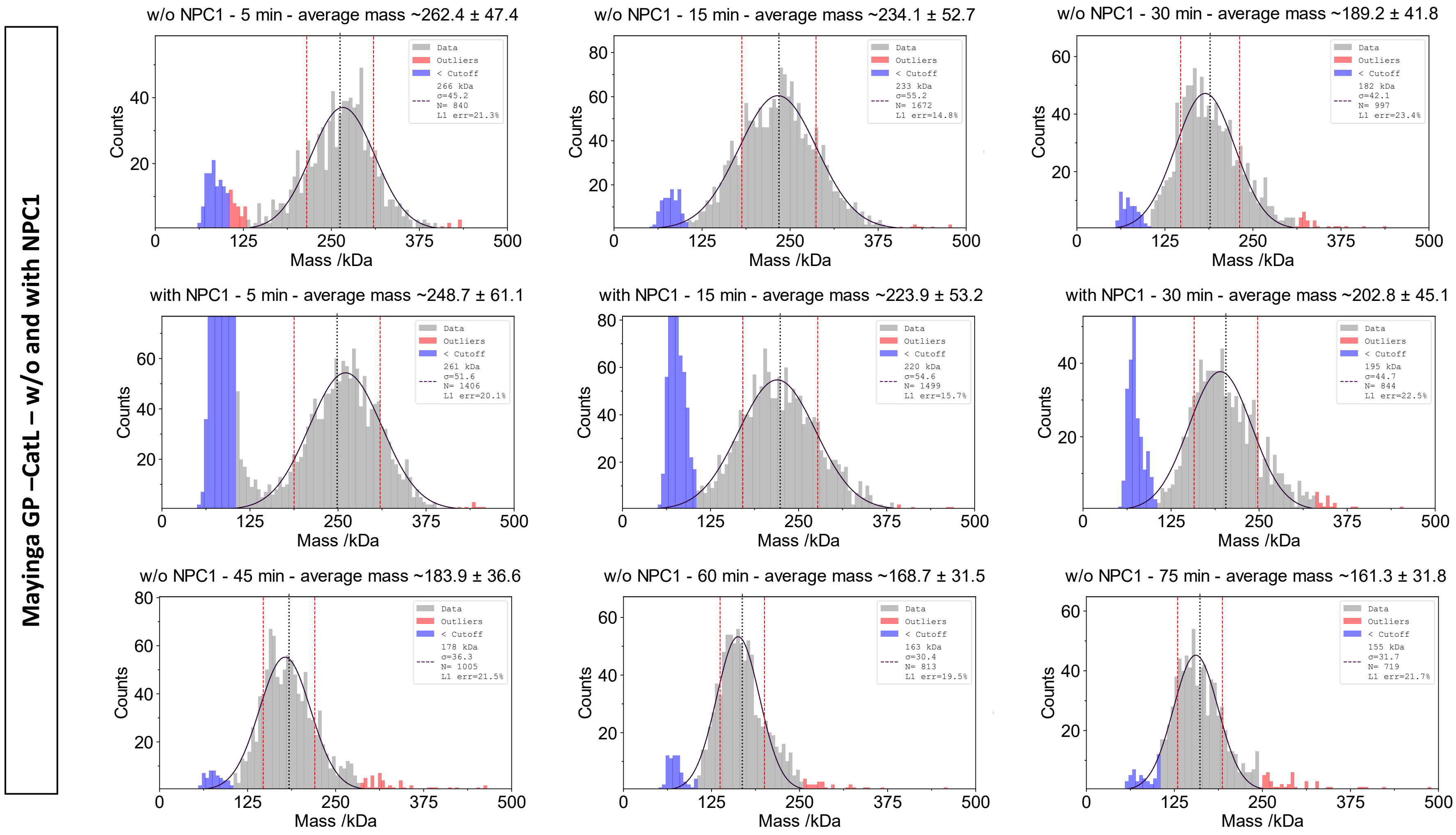
Mayinga GP –CatL – w/o and with NPC1
w/o NPC1 - 5 min - average mass ~262.4 ± 47.4
Data
Outliers
< Cutoff
266 kDa
σ=45.2
N= 840
L1 err=21.3%
w/o NPC1 - 15 min - average mass ~234.1 ± 52.7
233 kDa
σ=55.2
N= 1672
L1 err=14.8%
w/o NPC1 - 30 min - average mass ~189.2 ± 41.8
182 kDa
σ=42.1
N= 997
L1 err=23.4%
with NPC1 - 5 min - average mass ~248.7 ± 61.1
261 kDa
σ=51.6
N= 1406
L1 err=20.1%
with NPC1 - 15 min - average mass ~223.9 ± 53.2
220 kDa
σ=54.6
N= 1499
L1 err=15.7%
with NPC1 - 30 min - average mass ~202.8 ± 45.1
195 kDa
σ=44.7
N= 844
L1 err=22.5%
w/o NPC1 - 45 min - average mass ~183.9 ± 36.6
178 kDa
σ=36.3
N= 1005
L1 err=21.5%
w/o NPC1 - 60 min - average mass ~168.7 ± 31.5
163 kDa
σ=30.4
N= 813
L1 err=19.5%
w/o NPC1 - 75 min - average mass ~161.3 ± 31.8
155 kDa
σ=31.7
N= 719
L1 err=21.7%
Counts
Mass /kDa
0
125
250
375
500

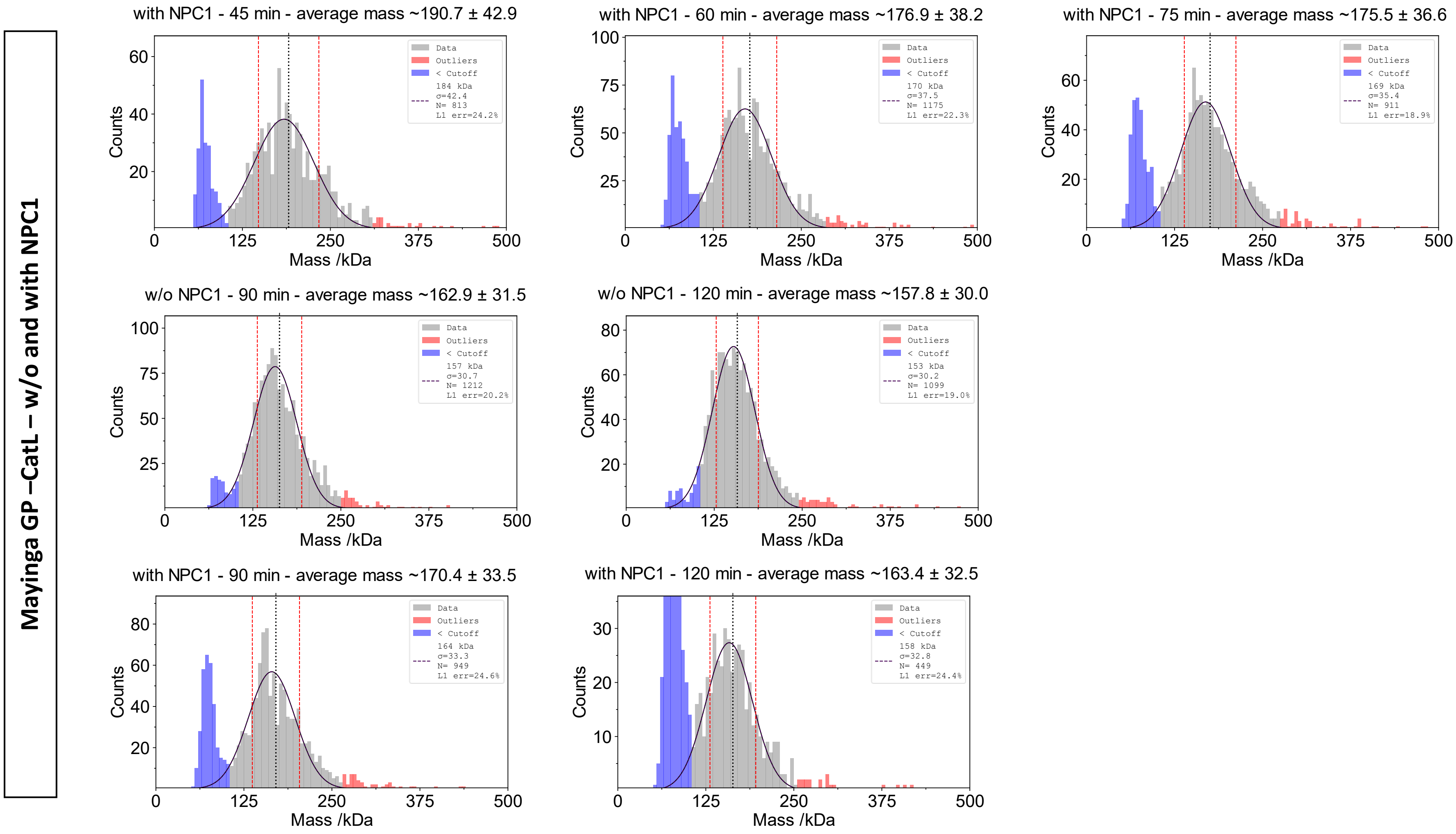
Mayinga GP –CatL – w/o and with NPC1
with NPC1 - 45 min - average mass ~190.7 ± 42.9
Data
Outliers
< Cutoff
184 kDa
σ=42.4
N= 813
L1 err=24.2%
with NPC1 - 60 min - average mass ~176.9 ± 38.2
Data
Outliers
< Cutoff
170 kDa
σ=37.5
N= 1175
L1 err=22.3%
with NPC1 - 75 min - average mass ~175.5 ± 36.6
Data
Outliers
< Cutoff
169 kDa
σ=35.4
N= 911
L1 err=18.9%
w/o NPC1 - 90 min - average mass ~162.9 ± 31.5
Data
Outliers
< Cutoff
157 kDa
σ=30.7
N= 1212
L1 err=20.2%
w/o NPC1 - 120 min - average mass ~157.8 ± 30.0
Data
Outliers
< Cutoff
153 kDa
σ=30.2
N= 1099
L1 err=19.0%
with NPC1 - 90 min - average mass ~170.4 ± 33.5
Data
Outliers
< Cutoff
164 kDa
σ=33.3
N= 949
L1 err=24.6%
with NPC1 - 120 min - average mass ~163.4 ± 32.5
Data
Outliers
< Cutoff
158 kDa
σ=32.8
N= 449
L1 err=24.4%
Counts
Mass /kDa

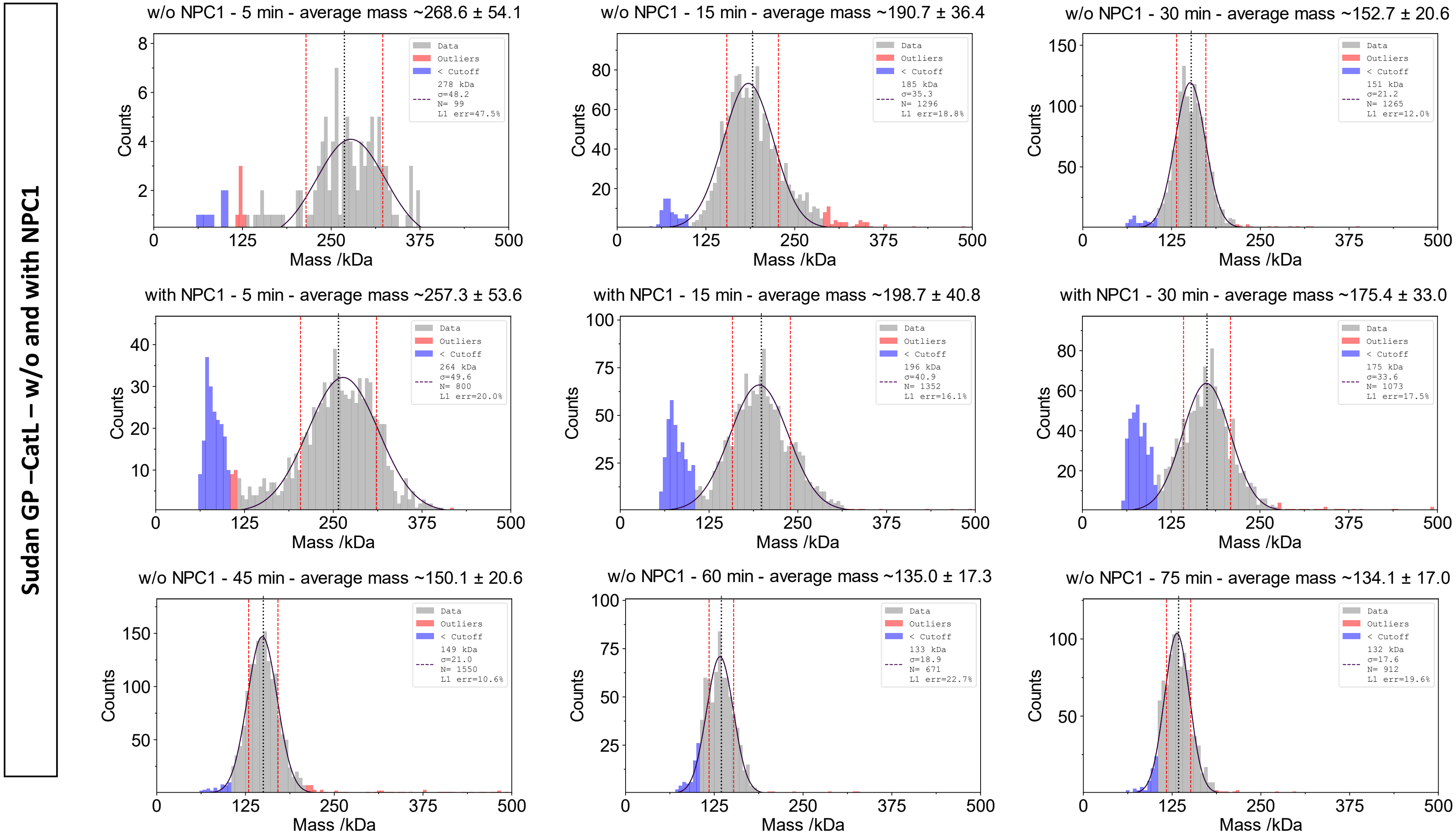
Sudan GP –CatL – w/o and with NPC1
w/o NPC1 - 5 min - average mass ~268.6 ± 54.1
Data
Outliers
< Cutoff
278 kDa
σ=48.2
N= 99
L1 err=47.5%
Counts
Mass /kDa
w/o NPC1 - 15 min - average mass ~190.7 ± 36.4
Data
Outliers
< Cutoff
185 kDa
σ=35.3
N= 1296
L1 err=18.8%
Counts
Mass /kDa
w/o NPC1 - 30 min - average mass ~152.7 ± 20.6
Data
Outliers
< Cutoff
151 kDa
σ=21.2
N= 1265
L1 err=12.0%
Counts
Mass /kDa
with NPC1 - 5 min - average mass ~257.3 ± 53.6
Data
Outliers
< Cutoff
264 kDa
σ=49.6
N= 800
L1 err=20.0%
Counts
Mass /kDa
with NPC1 - 15 min - average mass ~198.7 ± 40.8
Data
Outliers
< Cutoff
196 kDa
σ=40.9
N= 1352
L1 err=16.1%
Counts
Mass /kDa
with NPC1 - 30 min - average mass ~175.4 ± 33.0
Data
Outliers
< Cutoff
175 kDa
σ=33.6
N= 1073
L1 err=17.5%
Counts
Mass /kDa
w/o NPC1 - 45 min - average mass ~150.1 ± 20.6
Data
Outliers
< Cutoff
149 kDa
σ=21.0
N= 1550
L1 err=10.6%
Counts
Mass /kDa
w/o NPC1 - 60 min - average mass ~135.0 ± 17.3
Data
Outliers
< Cutoff
133 kDa
σ=18.9
N= 671
L1 err=22.7%
Counts
Mass /kDa
w/o NPC1 - 75 min - average mass ~134.1 ± 17.0
Data
Outliers
< Cutoff
132 kDa
σ=17.6
N= 912
L1 err=19.6%
Counts
Mass /kDa

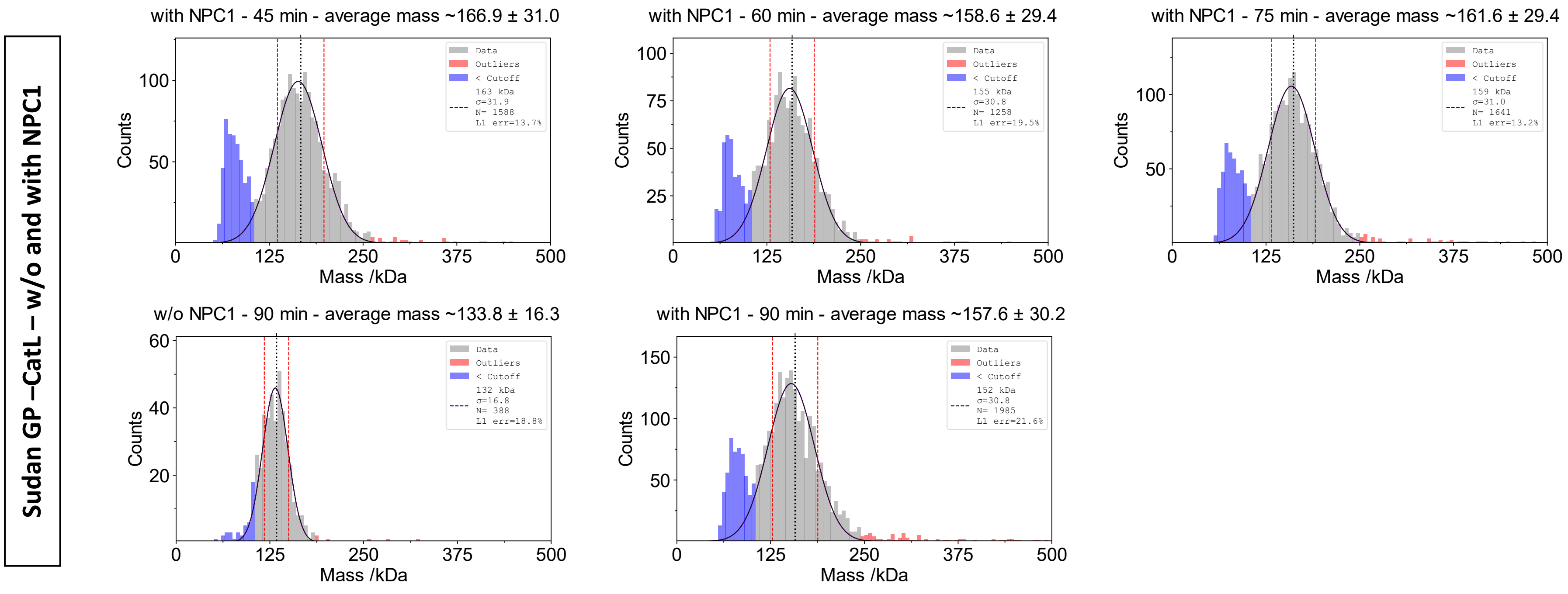

Sudan GP –CatL – w/o and with NPC1
with NPC1 - 45 min - average mass ~166.9 ± 31.0
Data
Outliers
< Cutoff
163 kDa
σ=31.9
N= 1588
L1 err=13.7%
with NPC1 - 60 min - average mass ~158.6 ± 29.4
Data
Outliers
< Cutoff
155 kDa
σ=30.8
N= 1258
L1 err=19.5%
with NPC1 - 75 min - average mass ~161.6 ± 29.4
Data
Outliers
< Cutoff
159 kDa
σ=31.0
N= 1641
L1 err=13.2%
w/o NPC1 - 90 min - average mass ~133.8 ± 16.3
Data
Outliers
< Cutoff
132 kDa
σ=16.8
N= 388
L1 err=18.8%
with NPC1 - 90 min - average mass ~157.6 ± 30.2
Data
Outliers
< Cutoff
152 kDa
σ=30.8
N= 1985
L1 err=21.6%
Counts
Mass /kDa
0
125
250
375
500

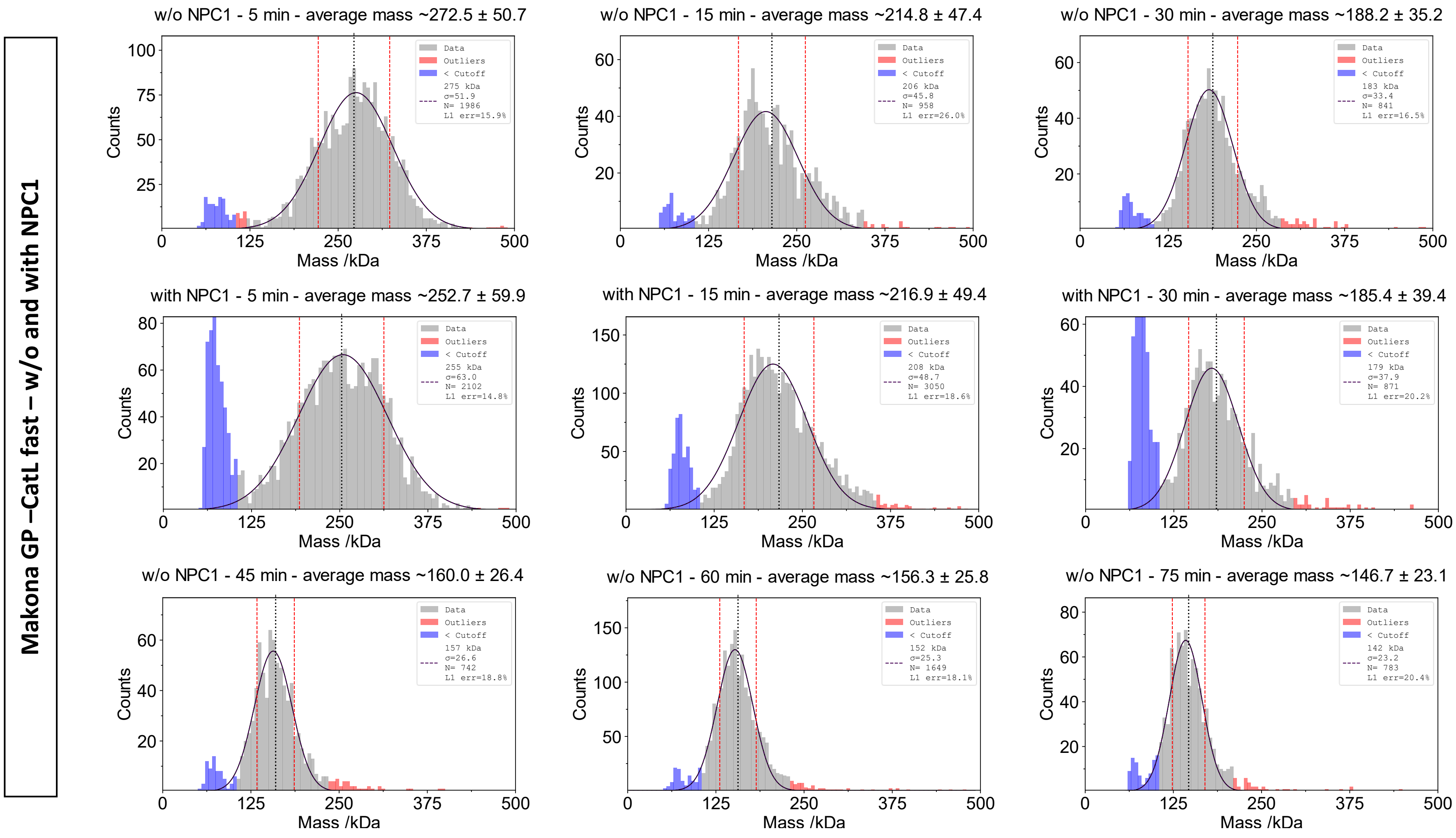
Makona GP –CatL fast – w/o and with NPC1
w/o NPC1 - 5 min - average mass ~272.5 ± 50.7
275 kDa
σ=51.9
N= 1986
L1 err=15.9%
w/o NPC1 - 15 min - average mass ~214.8 ± 47.4
206 kDa
σ=45.8
N= 958
L1 err=26.0%
w/o NPC1 - 30 min - average mass ~188.2 ± 35.2
183 kDa
σ=33.4
N= 841
L1 err=16.5%
with NPC1 - 5 min - average mass ~252.7 ± 59.9
255 kDa
σ=63.0
N= 2102
L1 err=14.8%
with NPC1 - 15 min - average mass ~216.9 ± 49.4
208 kDa
σ=48.7
N= 3050
L1 err=18.6%
with NPC1 - 30 min - average mass ~185.4 ± 39.4
179 kDa
σ=37.9
N= 871
L1 err=20.2%
w/o NPC1 - 45 min - average mass ~160.0 ± 26.4
157 kDa
σ=26.6
N= 742
L1 err=18.8%
w/o NPC1 - 60 min - average mass ~156.3 ± 25.8
152 kDa
σ=25.3
N= 1649
L1 err=18.1%
w/o NPC1 - 75 min - average mass ~146.7 ± 23.1
142 kDa
σ=23.2
N= 783
L1 err=20.4%
Data
Outliers
< Cutoff
Counts
Mass /kDa

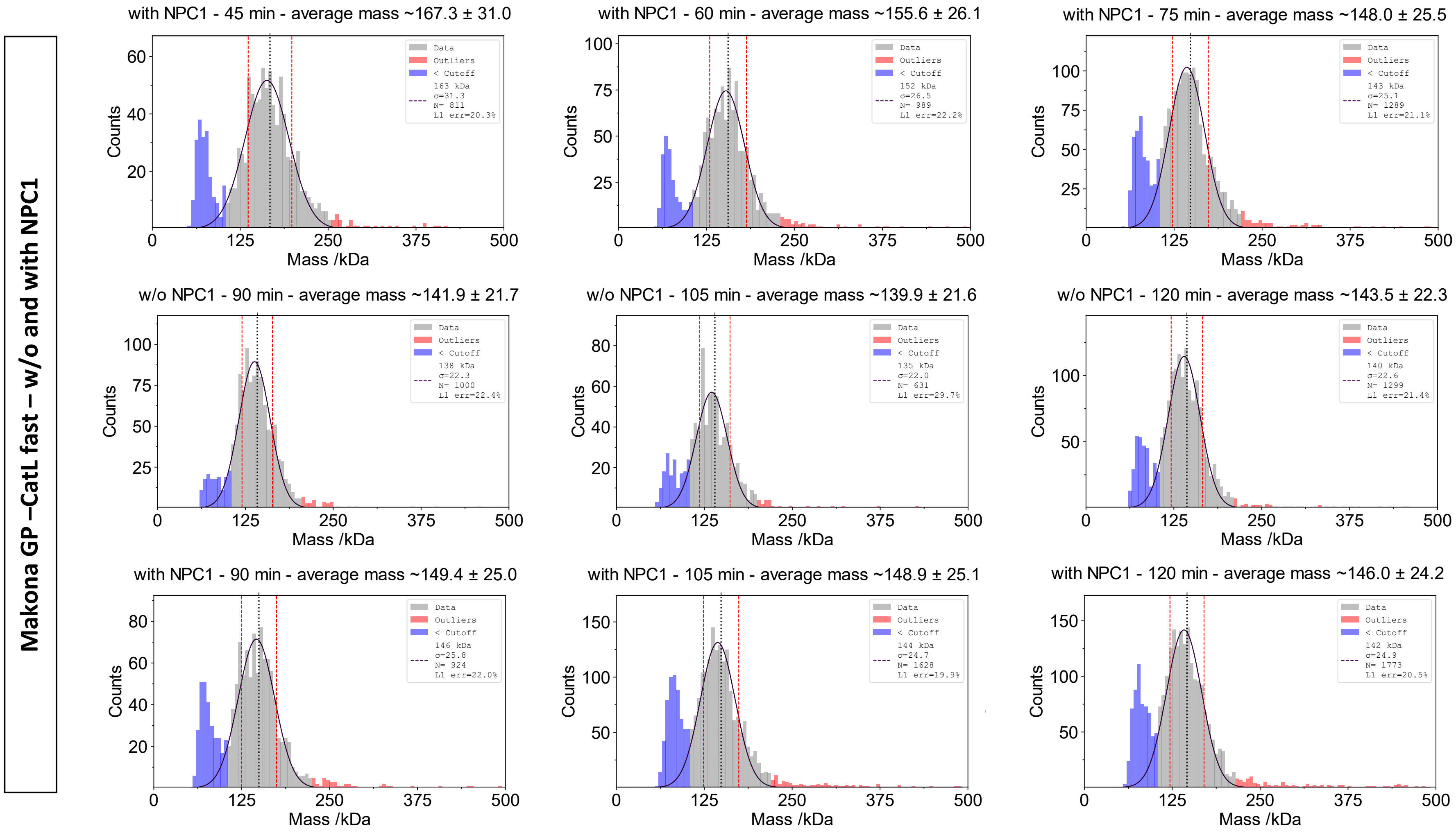
Makona GP –CatL fast – w/o and with NPC1
with NPC1 - 45 min - average mass ~167.3 ± 31.0
Data
Outliers
< Cutoff
163 kDa
σ=31.3
N= 811
L1 err=20.3%
with NPC1 - 60 min - average mass ~155.6 ± 26.1
Data
Outliers
< Cutoff
152 kDa
σ=26.5
N= 989
L1 err=22.2%
with NPC1 - 75 min - average mass ~148.0 ± 25.5
Data
Outliers
< Cutoff
143 kDa
σ=25.1
N= 1289
L1 err=21.1%
w/o NPC1 - 90 min - average mass ~141.9 ± 21.7
Data
Outliers
< Cutoff
138 kDa
σ=22.3
N= 1000
L1 err=22.4%
w/o NPC1 - 105 min - average mass ~139.9 ± 21.6
Data
Outliers
< Cutoff
135 kDa
σ=22.0
N= 631
L1 err=29.7%
w/o NPC1 - 120 min - average mass ~143.5 ± 22.3
Data
Outliers
< Cutoff
140 kDa
σ=22.6
N= 1299
L1 err=21.4%
with NPC1 - 90 min - average mass ~149.4 ± 25.0
Data
Outliers
< Cutoff
146 kDa
σ=25.8
N= 924
L1 err=22.0%
with NPC1 - 105 min - average mass ~148.9 ± 25.1
Data
Outliers
< Cutoff
144 kDa
σ=24.7
N= 1628
L1 err=19.9%
with NPC1 - 120 min - average mass ~146.0 ± 24.2
Data
Outliers
< Cutoff
142 kDa
σ=24.9
N= 1773
L1 err=20.5%
Counts
Mass /kDa
0
125
250
375
500

**Makona GP –CatL slow – w/o and with NPC1**

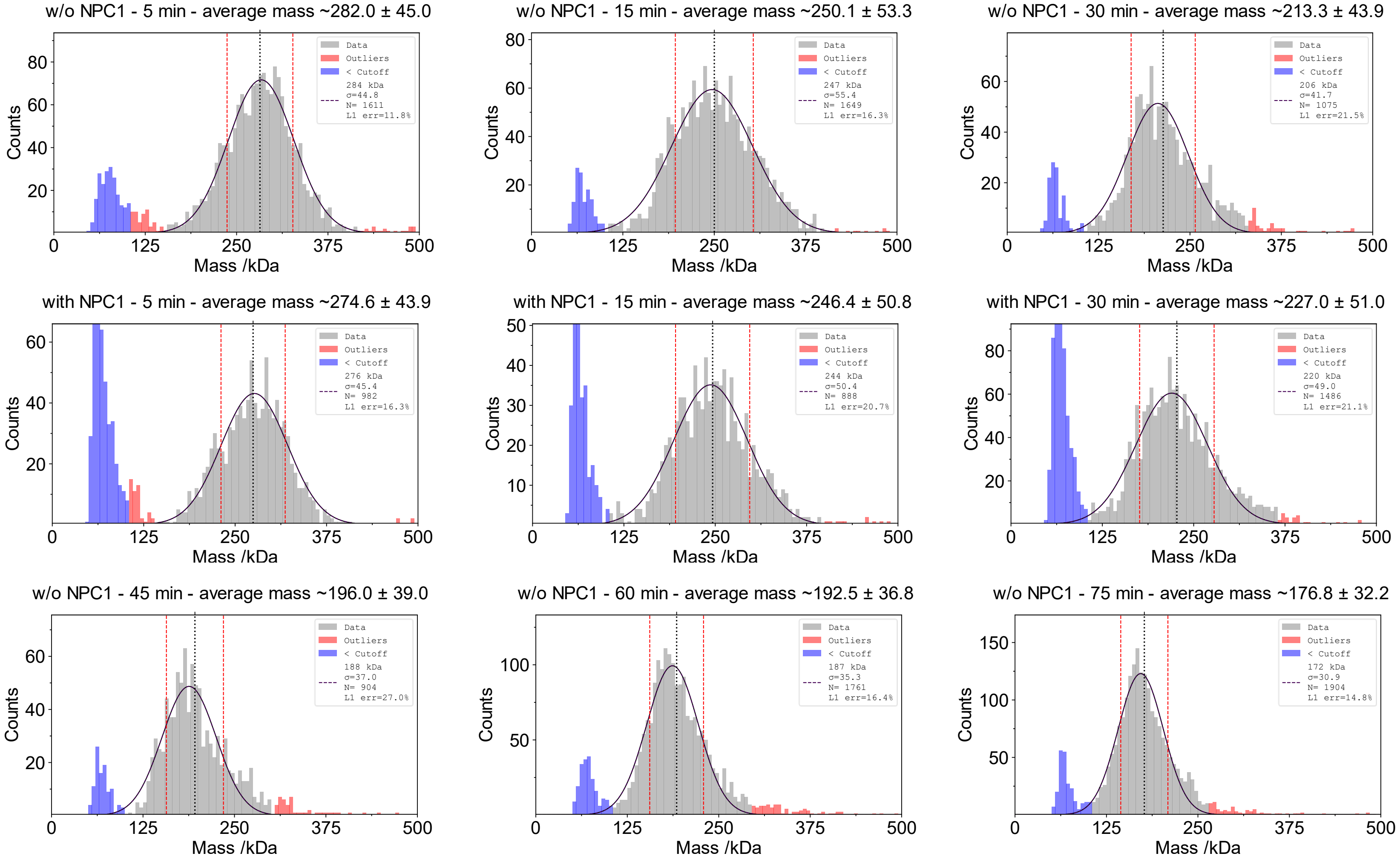

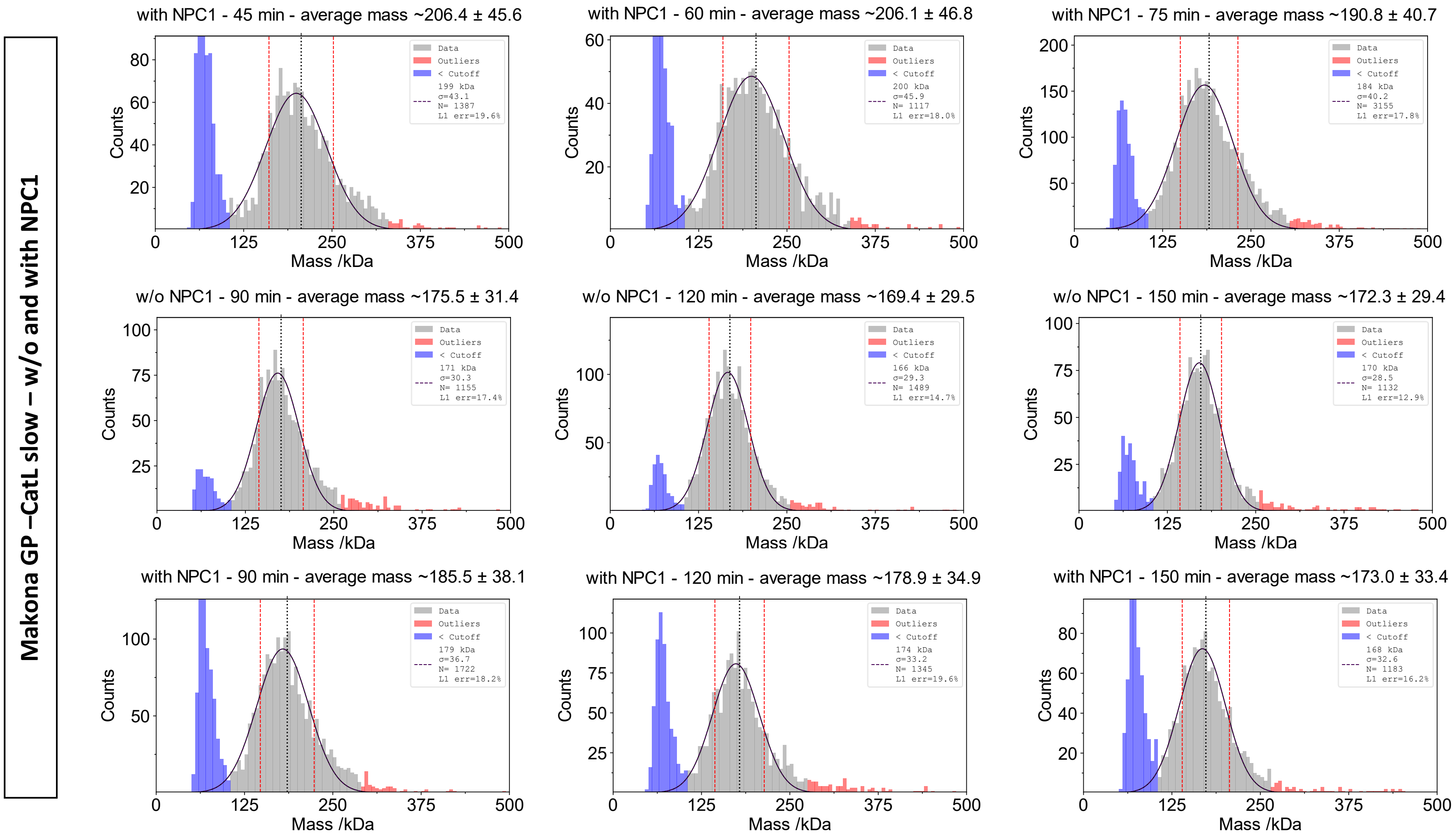
Makona GP –CatL slow – w/o and with NPC1
with NPC1 - 45 min - average mass ~206.4 ± 45.6
Data
Outliers
< Cutoff
199 kDa
σ=43.1
N= 1387
L1 err=19.6%
with NPC1 - 60 min - average mass ~206.1 ± 46.8
Data
Outliers
< Cutoff
200 kDa
σ=45.9
N= 1117
L1 err=18.0%
with NPC1 - 75 min - average mass ~190.8 ± 40.7
Data
Outliers
< Cutoff
184 kDa
σ=40.2
N= 3155
L1 err=17.8%
w/o NPC1 - 90 min - average mass ~175.5 ± 31.4
Data
Outliers
< Cutoff
171 kDa
σ=30.3
N= 1155
L1 err=17.4%
w/o NPC1 - 120 min - average mass ~169.4 ± 29.5
Data
Outliers
< Cutoff
166 kDa
σ=29.3
N= 1489
L1 err=14.7%
w/o NPC1 - 150 min - average mass ~172.3 ± 29.4
Data
Outliers
< Cutoff
170 kDa
σ=28.5
N= 1132
L1 err=12.9%
with NPC1 - 90 min - average mass ~185.5 ± 38.1
Data
Outliers
< Cutoff
179 kDa
σ=36.7
N= 1722
L1 err=18.2%
with NPC1 - 120 min - average mass ~178.9 ± 34.9
Data
Outliers
< Cutoff
174 kDa
σ=33.2
N= 1345
L1 err=19.6%
with NPC1 - 150 min - average mass ~173.0 ± 33.4
Data
Outliers
< Cutoff
168 kDa
σ=32.6
N= 1183
L1 err=16.2%
Counts
Mass /kDa
0
125
250
375
500
